# Evidence for high-frequency parallel evolution in virulent A. *baumannii* cultures

**DOI:** 10.1101/2024.10.22.618551

**Authors:** Jonathon Mclaughlin, John K. Tobin, Jon Hao, Ruth V. Bushnell, Taralyn J. Wiggins, Daniel V. Zurawski, Gregory J. Tobin, Stephen J. Dollery

## Abstract

*Acinetobacter baumannii*, is a gram-negative opportunistic pathogen notorious for its antibiotic resistance and adaptability. These attributes present significant challenges in clinical infection management and failure to manage infection results in around 1,000,000 global deaths per year. A greater knowledge of the layers of regulation employed by such a versatile pathogen may yield an improved understanding of the factors important for *A. baumannii* survival in diverse conditions and then facilitate the development of countermeasures. This study initially began with the investigation of phenotypic changes in colony opacity under varying environmental conditions with experiments designed to probe aspects of virulence, motility, biofilm formation, and antibiotic resistance. Our initial data also suggest evidence for a phenotype driving mutation system which simultaneously occurred in multiple lineages at the same time. This genetic alteration was observed at higher than expected frequencies, seemingly providing a striking example of parallel evolution. Using proteomic profiling and PacBio sequencing, we characterized lineages of AB5075 which, following changes in culture conditions, grew into colonies of a split translucent/ opaque phenotype which was inherited by translucent and opaque progeny lineages. A genetic alteration in the capsule operon gene *wzy*, marked by an ISAba13 transposon insert, led to the downregulation of the *wzy* gene product. Translucent variants demonstrated denser sedimentation and reduced biofilm formation, whereas both opaque and translucent variants showed unexpectedly similar antibiotic resistance profiles, challenging the assumptions that capsule formation and antibiotic resistance are always linked. Our findings suggest that the adaptability and resistance mechanisms of *A. baumannii* are related but distinct, where capsule loss is part of a broader, adaptive strategy rather than a sole determinant of antibiotic susceptibility. These insights highlight the need for a nuanced therapeutic approach, considering the dynamic interplay of environmental cues, phenotypic changes, and genetic rearrangements. We believe that this study further opens a path for understanding the adaptability of *A. baumannii* and lays the groundwork for developing innovative therapeutic strategies against this resilient pathogen.

**Significance Statement:** This study advances our understanding of the adaptability mechanisms of *Acinetobacter baumannii*, a pathogen notorious for its antibiotic resistance in clinical settings. We initially focused on the role of phenotype switches and identified genetic alterations in the Wzy operon that impact capsule production. Our studies led us to the surprising realization that we were observing multiple identical but independent evolution events. In addition, contrary to the traditional view that capsule variability directly correlates with antibiotic resistance, our findings revealed that susceptibility to aminoglycosides can occur independently of capsule loss, indicating capsule loss may be part of multiple survival strategies. These insights challenge existing paradigms and underscore the necessity for multiple therapeutic and preventative strategies that address the pathogen’s multifaceted adaptability for survival. Ultimately, this research contributes significantly to our knowledge of the biology of *A. baumannii*, paving the way for more effective management of infections caused by this formidable pathogen.

## Introduction

*Acinetobacter baumannii*, a gram-negative opportunistic pathogen, presents a considerable challenge in the clinical management of infections, particularly due to its remarkable ability to develop resistance to multiple antibiotics and persistence in the environment(1-4). Recent research underscores the critical roles of phenotypic variation mechanisms of rapid genetic change, in shaping the pathogen’s response to various environmental stresses. These phenomena are often reversible, high-frequency switches in gene expression and can endow *A. baumannii* with a formidable adaptability which can influence virulence, motility, biofilm formation, and antibiotic resistance (5-8).

Variant isolates of *A. baumannii* can be identified through changes in colony opacity, a visually discernible trait that correlates with alterations in various additional phenotypic characteristics (9-11). For instance, colony opacity has been implicated in traits such as surface motility and antibiotic susceptibility, suggesting a broader implication in the pathogen’s interaction with the host immune system (5, 12, 13). The significance of phase variation is believed to extend beyond the clinic; it represents a survival strategy, enabling the pathogen to adapt and persist in numerous hostile environments (8).

The underlying genetic mechanisms of phase variation often involve genetic modifications, such as slipped-strand mispairing or recombinational switches, which lead to reversible on-off expression of critical genes. These types of mechanisms can result in modulation of surface structures, including capsule formation and the expression of outer membrane proteins, which directly impact interactions with host defenses and antibiotics(13-16). Studies on the role of capsule in evading innate immunity and the transcriptional response of porin Omp33-36 to carbapenems and host cells illustrate the impact of phase variation on the survival and pathogenicity of *A. baumannii* (10, 17). Examination of a stable translucent isolate revealed an insertion of a ISBa13 transposable element in the *itrA* gene in the capsule locus (KL). Capsules and KLs are known to be highly variable (18).

Parallel evolution in microbes is characterized by the independent development of similar traits in distinct evolutionary lineages. Microbial populations, when subjected to similar environmental challenges, have been shown to evolve analogous genetic and phenotypic adaptations irrespective of genotype. This phenomenon is well documented for the recurrent emergence of antibiotic resistance across diverse bacterial species (19, 20) Specifically, in *A. baumannii*, parallel evolution has been reported extensively for multidrug-resistant (MDR) genes across multiple lineages (16, 21). Boarder studies of bacterial immunology and experimental evolution provide more direct evidence for this phenomenon (22-25). Parallel evolution is often rare and population size is considered a key contributor. Models suggest that parallel evolution is estimated to occur on average ∼ every 540 generations (19). The predominant theory holds that under conditions of strong selection, a mutation conferring a significant fitness advantage will increase over generations until it reaches fixation.

Here, we have discovered variants that were visibly discernable after culturing under biofilm conditions, revealing aspects of the organism’s capacity to respond to environmental pressures. Through proteomic profiling and PacBio sequencing, we identified several significant genetic alterations. Full genomic sequencing revealed that, in addition to the variants appearing to arise on the plate independently, they were indeed from genetically distinct lineages. This indicated that the visual phenotype had occurred in parallel in the cultures sublineages rather than in sequential progeny. This observation indicates a case of parallel evolution that mirrors the frequency of epigenetic changes under the specified conditions and that individual organisms can rapidly co evolve the same phenotype at a genetic level, but using different genes.

In efforts to understand the newly arisen visual opacity differences, we identified that a predicted hypothetical gene within the capsule operon (*wzy*-homologous regions) acquired an ISAba13 insert, and this insert led to the corresponding downregulation of the *wzy*-like gene. Notably, the translucent variants sediment more densely and exhibit reduced biofilm formation indicating that the translucence is indicative of a capsule defect. However, the antibiotic resistance profiles between opaque and translucent variants are similar, indicating that the link between capsule formation and antibiotic resistance is not dominated by the presence of capsule in these lineages. Unlike previous variants of a similar nature, a reversion to an opaque phenotype was not readily observed. This indicates that an additional mechanism appears to be “locking” the capsule mutation in place or that the selective pressure for reversion was not achieved.

Understanding these intricacies of phase variation in *A. baumannii* is pivotal for developing effective therapeutic strategies, including the design of vaccines and novel antimicrobial agents.

## Results

*A. baumannii* strain AB5075, a widely studied and highly virulent isolate, was cultured in T182 flasks, under M9 media. The media was refreshed daily and subsequently removed on day 5 to harvest the biofilm. The biofilm was collected by scraping, and subsequently disrupted by vortexing and pipetting in PBS followed by serial dilution and plating on TSB agar for quantitation. A notable frequency of variant colony formation was observed. The colonies depicted in **Figure 1** were all sourced from the same set of plates (e.g. **Figure 1A** and **1B**) which had been incubated for 4 days at 37°C. From these visibly heterogeneous colonies, four opaque and four transparent lineages were selectively propagated form the biphasic regions, as illustrated in **Figure 1C**. The growth kinetics appeared to be similar as can be seen by the serial dilution. This figure highlights the distinct density or color intensity of each isolate lineage. Each of the isolated colonies demonstrated robust growth, reaching a density of approximately 1.0 x 10^9^ CFU/mL overnight in a shaking culture of LB media at 37°C (5 µL spots are shown).

**Figure 1.**
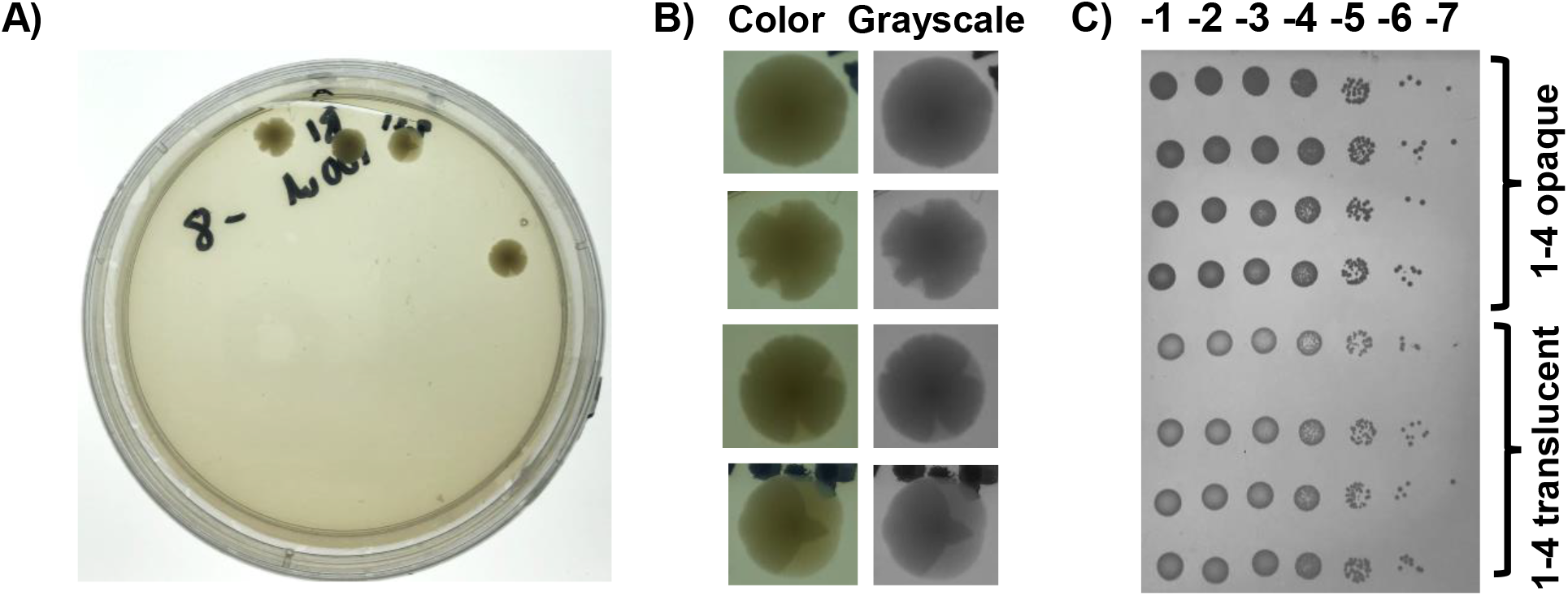
AB5075 were grown as biofilm cultures in T182 flasks containing M9 media and plated onto TSA-II plates. **A)** shows a plate with variant colonies arising **B)** shows individual colonies enlarged digitally from a 1x image which appear to arise from a single colony-forming unit but become morphologically biphasic as growth progresses (left color, right gray scale). Colonies were cultured for 3-4 days to reach approximately 5 mm in diameter. **C)** Translucent and opaque regions from the colonies were streaked 3x, cultured in TSB broth, and serially diluted on a TSB agar plate to clarify differences in morphology(indicated).

The distinct appearances of opaque and translucent variants in *A. baumannii* cultures suggested possible differences in the ability to produce functional capsules. To explore this hypothesis, overnight cultures of the variants were assessed for density using Percoll gradient sedimentation, and compared with parental stock cultures. Notably, the translucent variant demonstrated more rapid migration indicating higher density. A significant portion of the translucent culture settled at the interface between the 100% and 60% Percoll layers. Although mildly disturbed post-centrifugation, this pattern is still prominently visible in **Figure 2A**. In contrast, the opaque variant and the parental cultures remained as a single band at the media Percoll interface.

**Figure 2.**
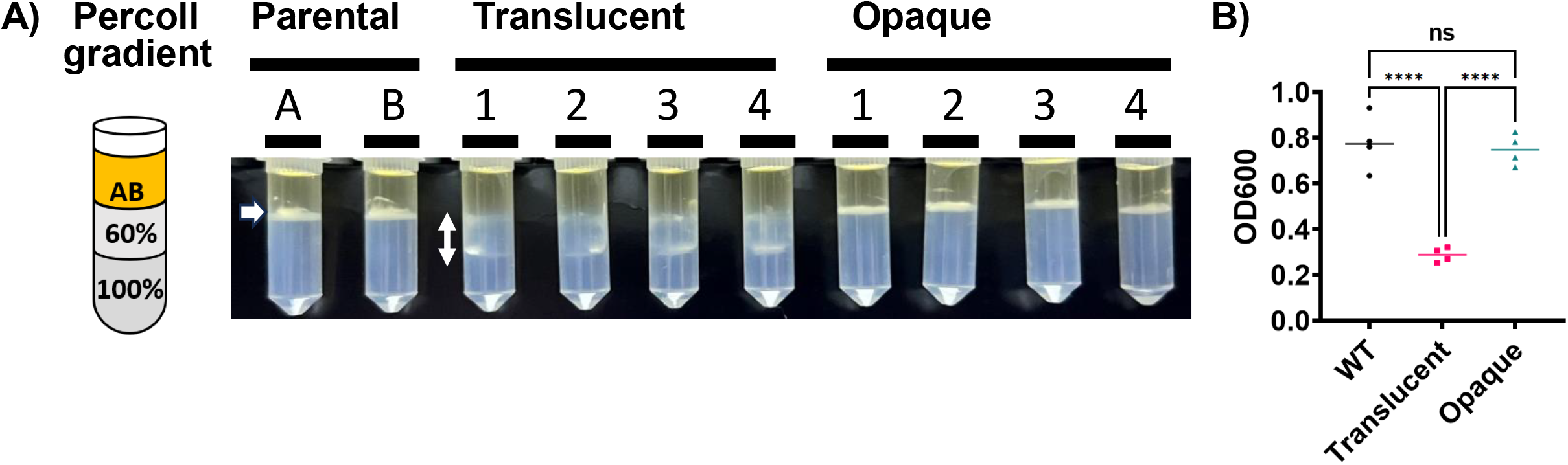
Phenotypic validation of capsule production of A. baumannii and K. pneumoniae WT and mutant strains. **A)** Overnight cultures were layered over Percoll step gradients in transparent 2 ml tubes containing 60% and 100% Percoll in equal volumes. Letters A and B indicate biological duplicate cultures of a parental stock. Numbers 1-4 indicate unique lineages of translucent or opaque morphological variant cultures. An alteration in migration consistent with physiological differences was seen in all translucent lineages. White arrows highlight density changes in translucent samples. None such alteration is seen in the opaque linages in comparison to the parental cultures. **B)** Crystal violate staining of variants or biological replicates of the parental grown on a 96 well plate. Cells were fixed in methanol-acetone (50/50), stained in crystal violate (.05% in PBS), washed 3x, dissolved in 30% glacial acetic acid (transferred to a clean plate) and absorbance read at 600nm. **** denotes an adjusted P value of <0.0001, as derived from an ANOVA with Tukey’s multiple comparisons test applied.

Further investigation into the biofilm formation and adherence properties of these variants was conducted. Cultures were grown for two days in a 96-well plate, then fixed and stained with crystal violate. **Figure 2B** shows OD600 readings indicating less staining in wells with translucent colonies, suggesting potential differences in biofilm stability, biomass quantity, or protein content. This observation was consistently replicated across four sets of parental and opaque preparations, with translucent cultures #1-4 showing less staining indicating a decreased retention of biomass in these samples.

To investigate potential genetic alterations and protein expression differences in the isolates, comprehensive analyses were conducted using PacBio sequencing and mass spectrometry. The detailed sequences and associated findings will be presented in an accompanying paper, which will explore further genetic rearrangements upon subsequent verification although we do wish to make the following observations. The regulation of capsule production genes was a focal point of our analysis. Notably, a region containing a gene predicted to encode a Wzy-like protein exhibited a 1048 nucleotide insertion, as illustrated in **Figure 3A**. The Wzy gene product is located centrally in the chromosome of AB5075 and has been associated with capsular polysaccharide synthesis. However, examination of the adjacent capsule operons did not reveal any other genetic changes. A nucleotide BLAST search identified the insertion as the transposable element ISbA13, which has previously been shown to insert into another K operon gene, *itrA* but appears to be located at a unique insertion site in these lineages (11). The insert described here helps to confirm the potential function of the hypothetical Wzy protein and a potential mechanism for the translucent and opaque phenotypes seen here. The observation that the same genetic alterations appear to occur spontaneously at high frequency suggests parallel evolution can occur at high frequency in the right environment. To display the overall relationships between these variants several analyses are shown. **Figure 3B** shows a phylogenetic tree which indicates that the variants are related but distinct. Although exact lineages were not rigorously tracked during the initial isolation (pick), the analysis revealed that certain pairs, such as O1 and T1, showed close clustering, suggesting genetic similarity between some translucent and opaque variants as would be expected. To further examine global genetic analysis, isolates were compared using the Progressive Mauve system from the Darling Lab on the chromosomal assemblies **Figure 4C** (26). Mauve uses an iterative alignment approach to maximize the accuracy of the alignment. The genomes were then visualized in a color-coded representation of the alignments which allows for rapid identification of identical regions. Please note that *A. baumannii* is known to undergo genetic inversion in a central region. These again indicate that the linages are similar but distinct. Taken together these indicate that the isolates were similar until picked following a capsule mutation but then some further genetic alterations occurred during streaking to ensure clonality before sequencing. Perhaps these occurred as the clones readapted to a rich nutrient source.

**Figure 3:**
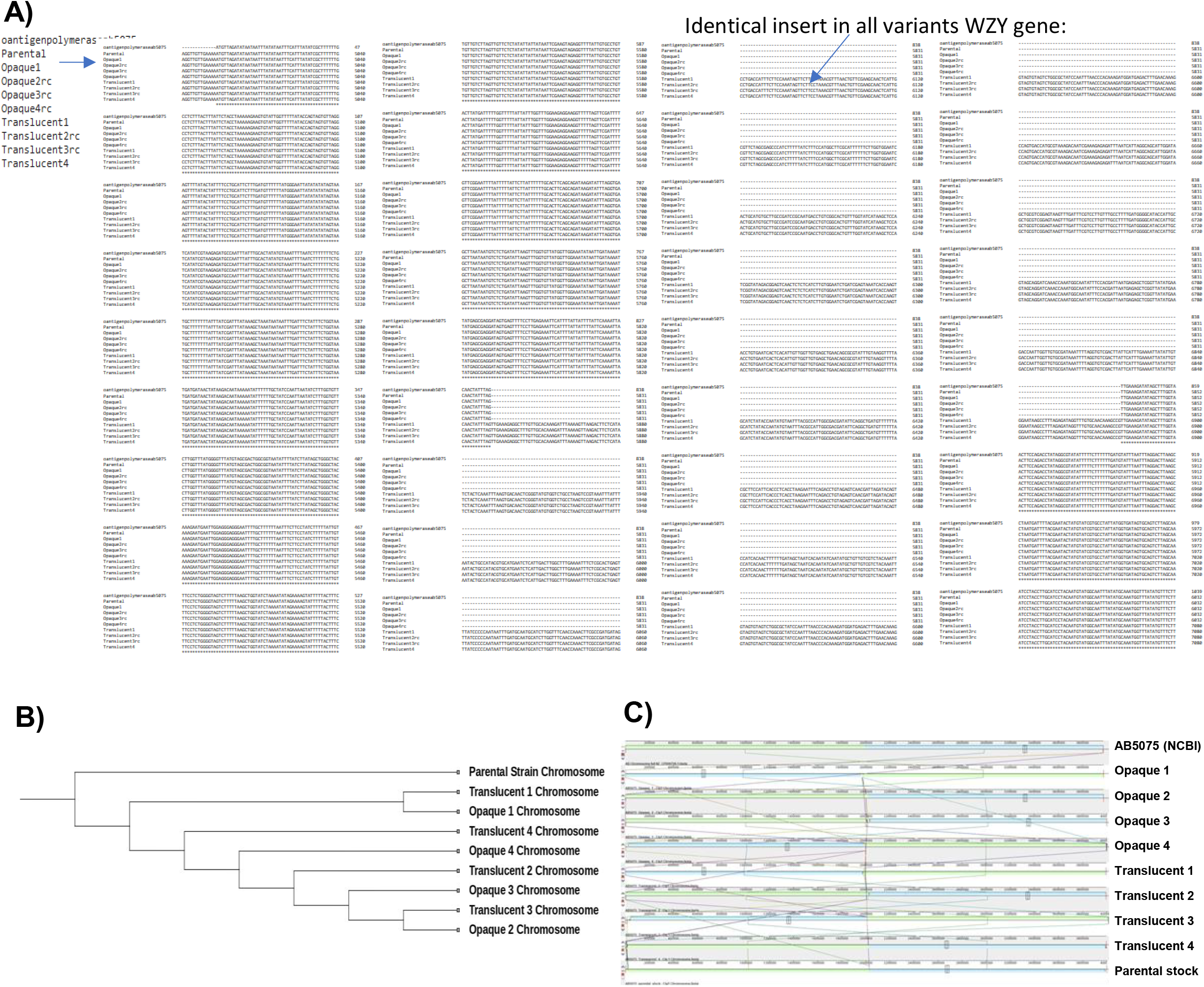
Genomic Analysis of Capsule Production Gene Wzy from Isolate and Parental Cultures. **A)** This figure presents the genomic sequence analysis of the K operon region surrounding the gene predicted to encode the Wzy protein in *Acinetobacter baumannii* isolates and showing the insertion of a 1048 bp ISbA13 transposable element. **B)** Phylogenetic tree of Chromosomes of variants in relationship to the parental. **C**. Progressive Mauve analysis of isolates and Parental AB5075.

**Figure 4:**
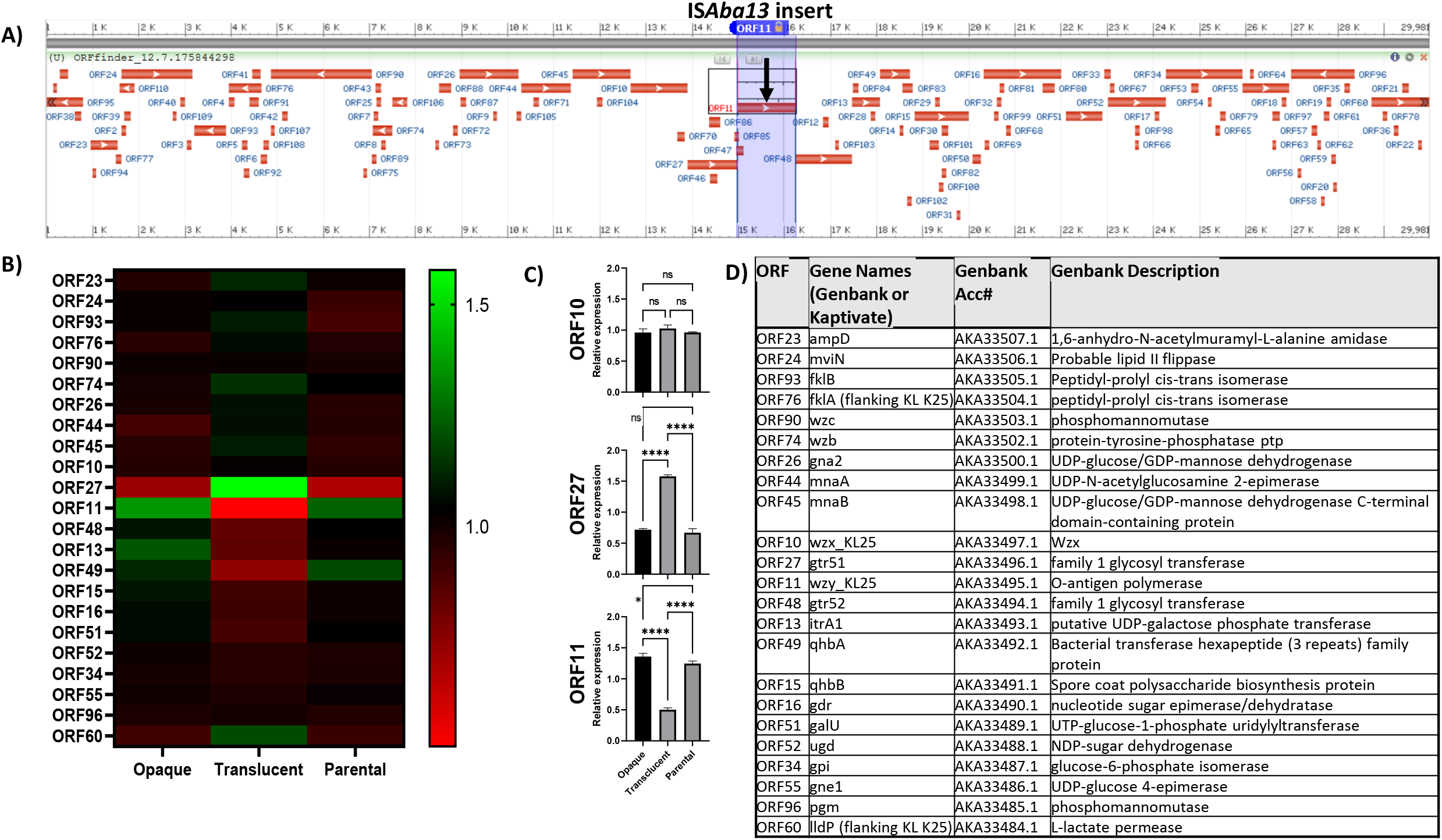
Group Proteomic Analysis of *Acinetobacter baumannii* K locus. **A)** Displays the genomic region around the Wzy-encoding gene which is the site of ISAba13 insertion, highlighting ORF orientations indicate the K operon structure. **B)** A heat map showing relative protein concentration levels in translucent and opaque variants and parental stock, providing a visual comparison across different isolates. **C)** Bar graphs for ORFs 10, 27, and 11, depicting average expression levels with statistical analysis (ANOVA and Tukey’s correction), focusing on expression differences between isolates. **D)** A table listing gene names and functions related to the ORFs, serving as a reference for understanding gene expression patterns and their implications.

**Figure 4A** depicts the genomic region surrounding the insertion site and illustrates the orientation of open reading frames (ORFs), known to be the K locus (capsule) (18). Analysis with Kaptive shows that the K locus type for AB5075 is KL25. Mass spectrometry analysis enabled the quantification of relative abundance levels for the majority of ORF products in the vicinity. This analysis was conducted on isolates 1, 2, and 3 of both translucent and opaque variants, as well as three independent preparations of the parental stock, all of which were cultured overnight on agar plates.

For our assessment, we averaged the relative expression levels within each group. The results are displayed in two formats: a heat map for overall visualization (**Figure 4B**), and, for ORFs 10, 27, and 11, bar graphs depicting the mean expression levels (**Figure 4C**). The bar graphs include statistical analysis with ANOVA and Tukey’s multiple comparison correction. Additionally, the table in **Figure 4D** provides a key to gene names and their associated functions.

Upon comparing protein expression levels, we found that the expression of genes from ORF 23 to ORF 10, encompassing areas outside and within the operon, remained largely unchanged. Notably, the protein encoded by the gene immediately upstream of the insertion site (ORF 27) was significantly upregulated. Conversely, the protein at the insertion site exhibited significant reduction. These alterations are evident in both the heat map and the bar graphs for selected protein levels. Furthermore, the levels of proteins encoded by genes downstream of the insertion site also showed a trend of mild reduction, with the effect generally diminishing progressively with increased distance from the insertion.

### Dysregulation of proteins in the translucent group identified to be involved in organic substance biosynthesis and cellular metabolic pathways

To identify proteins which are significantly different in the translucent group when compared to the parental group, differential expression analysis was performed by mass spectrometry. Results of this analysis, shown in **Figure 5A**, have identified 539 upregulated proteins and 400 downregulated proteins in the translucent group compared to the parental group. The proteins that were most upregulated with the most significance in the translucent group were the flavoprotein subunit of succinate dehydrogenase, subunit B of ATP synthase F0 and another protein belonging to the OmpW family. Of the 539 significantly upregulated, those with the highest fold change that were not hypothetical proteins, included a lysine exporter protein, with a fold change of 1.36, a citrate transporter with a fold change of 1.28 and a family 1 glycosyl transferase with a fold change of 1.24. (Supplemental Table 1)

**Figure 5.**
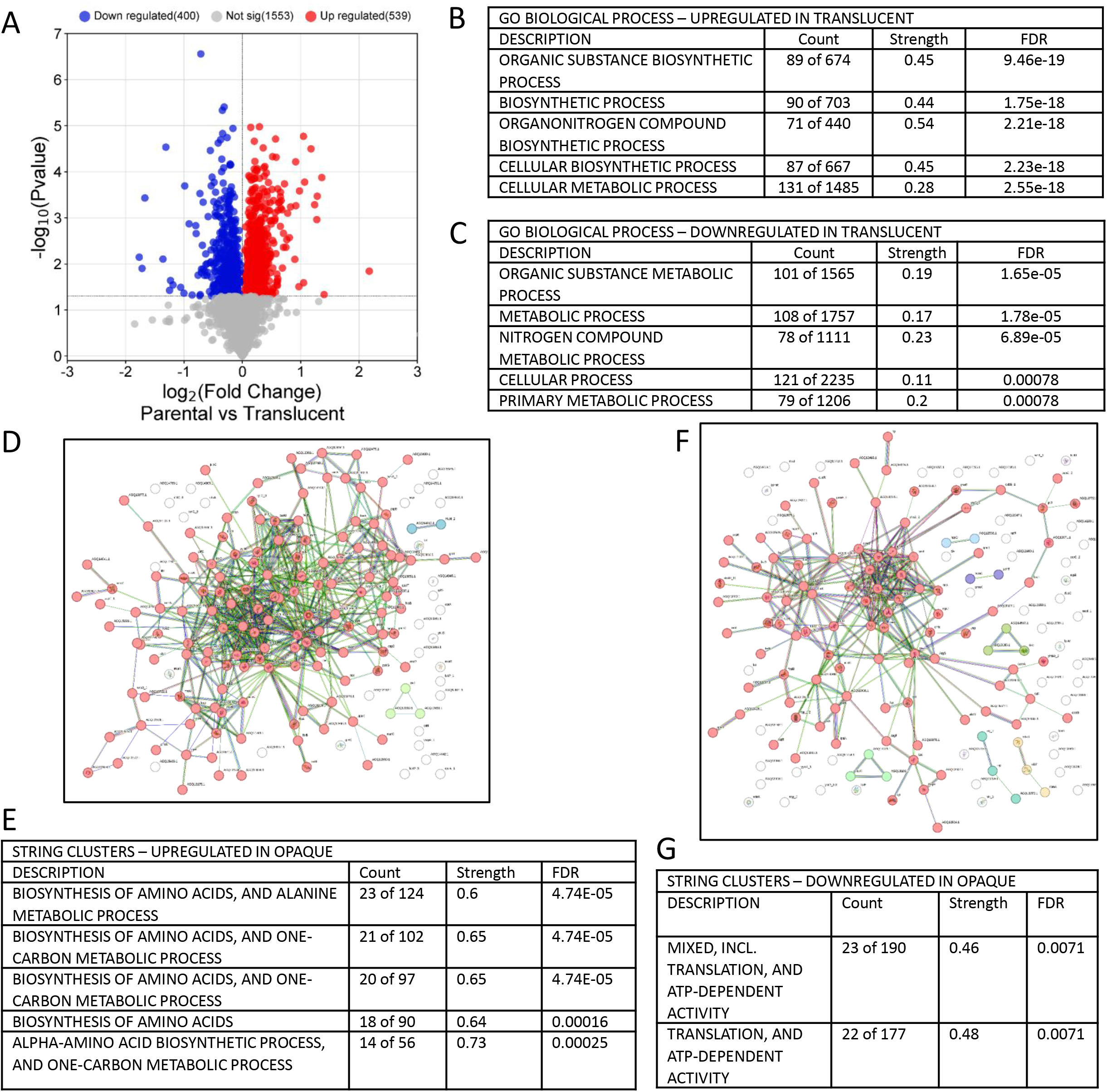
Dysregulated Proteins in Translucent Vs Parental Isolates: GO Enrichment and STRING Analysis. **A)** volcano plot of differentially expressed proteins– blue points represent downregulation, red represents upregulation and grey did not reach statistical significance (p<0.05). **B & C)** gene ontology pathway analysis of upregulated proteins. – count shows the number of proteins identified in an entire pathway list, strength is the level in which the protein signature identified is involved in the pathway and False Discovery Rate (FDR) value which is determined from the observed p-value distribution (FDR<0.05). **D**. STRING network of upregulated proteins (K=3). **E)** Highlights of STRING Clusters with significance scores. **F & G)** STRING analysis (K=7) and highlights of downregulated clusters.

The most significant proteins downregulated in the translucent group included an oxidoreductase short chain dehydrogenase, a nitroreductase family protein and a peptidase belonging to the M48 family. Downregulated proteins with the highest fold change included some hypothetical proteins, and, namely, a hypothetical plasmid protein (AB5075) ABUW_4122. Other proteins with significantly reduced levels included fructose-1,6-biphosphate with a fold change of - 1.67 and the thiamine biosynthesis protein ThiC with a fold change of -1.36. Significant downregulation was also identified in multidrug resistance protein B and modulator of drug activity B.

The dysregulated proteomic signatures were then used for gene ontology (GO) enrichment analysis to identify biological processes which involve these proteins. In **Figures 5B and C** the resulting data shows upregulation in proteins comprising the GO term organic substance biosynthetic process, which is the chemical reactions and pathways involving an organic substance, as well as other biosynthesis processes such as organonitrogen compound and cellular biosynthesis. Downregulated proteins were identified in organic substance metabolic processes and nitrogen compound metabolism. The pattern of up- and down-regulation suggests that there is an upregulation in organic compound biosynthesis and a downregulation of organic compound metabolism.

### Dysregulated proteins share a high number of STRING interactions with a signature of biosynthesis and ATP-dependent activity

STRING analysis of the same differentially expressed proteins provides an alternate look at the interactions within the differentially regulated proteomic signature. The resulting map shows each protein as a node and the known interactions as a line between nodes, with lines representing co-expression, co-mentions in PubMed and proteins falling within the same gene neighborhood to allow for a more in-depth picture of the overall signature. In the upregulated signature **Figure 5D**, there are a small number of outlying protein nodes with no significant connections, however, using k-means clustering with a natural cluster of 3 has shown one major signature highlighted in red. After filtering the proteins to those that are found across the databases available to STRING, there were a total of 174 nodes or proteins with 535 connections. The expected number of edges is 328, showing a higher number of connections than expected in a natural state and resulting in a PPI enrichment score of 1.0e-16. The STRING cluster pathways are shown in **Figure 5E** and consists of proteins involved primarily in the biosynthesis of amino acids with alanine and one-carbon metabolic processes.

The downregulated network of proteins has also shown a significant number of connections between the protein nodes **Figures 5F and G**, with an expected 193 connections but revealing a total number of 266 connections between 156 proteins. This provides a PPI enrichment value of 3.57e-07 deriving from the significantly more interactions than naturally expected. A k value of 7 was used here based on the natural number of clusters in the raw proteomic data, with the largest cluster highlighted again in red. This primary cluster of proteins has revealed STRING cluster pathways including translation and ATP-defendant activity.

### Dysregulation of proteins in opaque isolates are involved in cellular metabolism and biosynthesis of novel compounds

The same differential expression analysis was performed using the opaque isolates and comparing them against the parental samples (**Figure 6A**). Resulting data has shown an upregulated proteomic signature consisting of 255 upregulated and 318 downregulated proteins (Supplemental Table 2). The most significant proteins which were upregulated in the opaque group of isolates included the alpha and beta subunits of NAD(P) transhydrogenase and acyl-CoA dehydrogenase. Two separate alpha subunits of NAD(P) transhydrogenase found at different locations during mass spectrometry were in the top 3 proteins with the largest fold change. Log2 fold changes of 1.74 and 1.47 were identified in both alpha subunits, alongside several hypothetical proteins and a metallo-beta-lactamase family protein with a fold change of 0.92. In terms of downregulation in the opaque isolate group when compared to the parental group, the most significant proteins included a peptidase of the M48 family and a nitroreductase family protein, both of which were also identified as significantly downregulated in the translucent group when compared to parental isolates. Other significant downregulation was identified in adenosylmethionine-8-amino-7-oxononanoate transaminase and a toluene tolerance protein. Downregulation of proteins with the largest fold change was identified as fructose-1,6-biphosphate with a fold change of 1.64 and the thiamine biosynthesis protein ThiC with a fold change of 1.4, - as seen in the translucent samples. This overlap in downregulation may indicate the changes occurring when translucent and opaque isolates are separately compared to the parental group may describe shared genetic alterations, similar environmental adaptations, or the beginning of regulatory signaling cascades.

**Figure 6.**
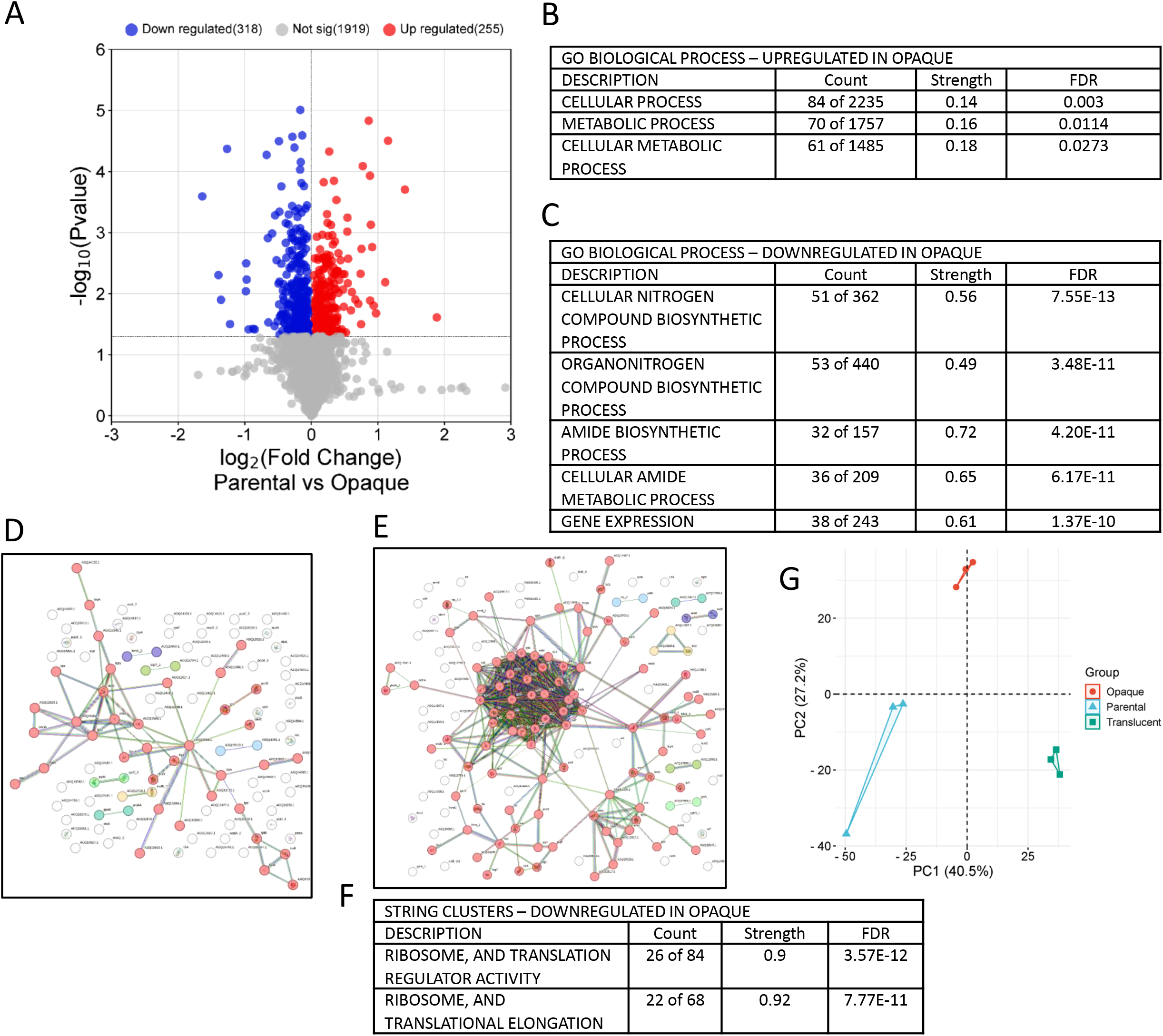
Dysregulated Proteins in Opaque Vs Parental Isolates: GO Enrichment and STRING Analysis. A) volcano plot of differentially expressed proteins– blue points represent downregulation, red represents upregulation and grey did not reach statistical significance (p<0.05). **B & C)** gene ontology pathway analysis of upregulated proteins. – count shows the number of proteins identified in an entire pathway list, strength is the level in which the protein signature identified is involved in the pathway and False Discovery Rate (FDR) value which is determined from the observed p-value distribution (FDR<0.05). **D**. STRING network of upregulated proteins (K=7). **E&F)** String analysis (K=7) and highlights of downregulated clusters. **G)** shows a scatter plot representing the results of the Principal Component Analysis (PCA) – each data point corresponds to an observation within the dataset, with the x-axis and y-axis representing the first 2 principal components (PC1 And PC2) – which collectively explain 67.7% of the total variance in the data.

Dysregulated proteins identified through differential expression were again used as input for gene ontology (GO) enrichment analysis of biological processes implicated by these proteomic changes (**Figure. 6B and C**). Results provide evidence of upregulation primarily in cellular and metabolic processes, however, the strength of this is relatively low but does provide significant false discovery readings indicating significance in a biological setting. Downregulation on the other hand has provided a much more significant list of biological processes, namely, cellular nitrogen compounds, organonitrogen compound and amide biosynthesis processes. Resulting data also shows downregulation of gene expression and a relatively high strength of 0.61 and FDR reading of 1.37e-10.

Genes identified to be downregulated and involved in this pathway include *tyrS* (Tyrosyl-tRNA Synthetase) *rbfA* (Ribosome Binding Factor A), *rpmG* (Ribosomal Protein L33) and *rplY* (Ribosomal Protein L25). Each of these proteins presented a log two-fold change of -0.067, -0.102, -0.141 and -0.221, respectively. The protein TyrS is an enzyme that charges tRNA molecules with tyrosine, meaning its primary role is in translation, however, its activity impacts gene expression by maintaining the efficiency and fidelity of the translation process. Both RpmG and RplY are components of the 50S subunit, with both being integral to the function of the ribosome. Alterations of these can affect translation, thereby influencing overall gene expression. Dysregulation of ribosome binding factor A can also be detrimental to the functionality of translation machinery, RbfA is involved in the maturation of 16S RNA, which is a vital component of the 30S ribosomal subunit (PMID: 12963368). Downregulation of this protein and subsequent deficiencies in ribosome assembly can potentially lead to global downregulation of protein synthesis thus affecting overall gene expression.

### STRING cluster analysis of opaque isolates reveal downregulation in ribosome and translation regulation and elongation

Subsequent STRING analysis of the differentially expressed proteins in the opaque isolates have resulted in distinct network clusters in both upregulated and downregulated signatures. In the upregulated network (**Figure. 6D**), a k value of 7 was implemented based on the natural clusters and a total of 103 protein nodes with 68 connections were observed. The k value of 7 describes the number of clusters the algorithm will partition the data into, 7 was used here based on the distinct clusters already seen in the STRING network without applying any clustering methods. The expected number of edges in this network is 55, resulting in a higher number of edges than expected but a PPI enrichment score of 0.0552 which has not met statistical significance. There is still a significantly large primary cluster, highlighted in red, but due to the lack of significance, there are no relevant STRING clusters to note.

In the downregulated signature however, there is a strong PPI enrichment score of 1.0e-16, with an expected number of 308 connections but an observed number of 539 connections among 145 protein nodes. The natural clustering of this network has provided a k value of 7, with the primary cluster shown in red in **Figure 6E**. STRING cluster analysis of this primary cluster has resulted in 2 main pathways in which downregulated proteins in the opaque isolates are implicated in (**Figure 6F**). Resulting data shows that this cluster is responsible for downregulation in ribosome and translation regulator activity, as well as ribosome and translational elongation, with both pathways presenting a strength of over 0.9 and highly significant false discovery rates.

### Principal component analysis (PCA) depicts a high inter-cluster distance while retaining a low intra-cluster distance

A PCA plot was used to reduce the dimensionality of the proteomic data (**Figure 6G**), this provides a better overview of the high-dimensional data to show the differences between each of the 9 different isolates – 3 parental, 3 opaque and 3 translucent. Using a PCA plot allows for a simplified figure while preserving most of the variability present in the original dataset, this is achieved by transforming the original variables into a new set of orthogonal variables, or principal components. Resulting PCA data of PC1 and PC2 collectively accounted for 67.7% of the total variance in the proteomic data, PC1 explained 40.5% of the variance, while PC2 explained an additional 27.2%. The distribution of the 3 groups along the principal component is depicted in a scatter plot, where each data point represents an observation within the dataset, the separation of these groups along PC1 and PC2 indicates distinct patterns or relationships among each group. Overall, the PCA results highlight the presence of distinct groupings within the dataset and provide a framework for understanding the relationship and structure of the data. Relating this back to large-scale dysregulation in STRING while still showing high levels of interactivity between proteins, shows that the distinct clusters observed through PCA indicate that the phenotypic differences are strongly reflected at the proteomic level. Pathways identified through STRING enrichment analysis can help to explain these differences, in essence, proteins or pathways which are highly dysregulated could be driving the separation seen in PCA. Proteomic analysis on translucent isolates revealed a higher level of dysregulation in pathways primarily related to metabolic processes which could potentially drive the phenotypic differences seen in the translucent PCA grouping. Although there were less protein interactions in the opaque isolates shown through STRING enrichment, a large signature of dysregulated proteins was still evident, further showing that proteomic signatures of dysregulation in both opaque and translucent isolates may be driving the phenotypic differences seen in both groups.

In our investigation of the potential impact of capsule deficiency on antibiotic resistance, duplicate cultures of the isolates were cultivated overnight. These cultures were then used to inoculate a variety of media containing antibiotics from diverse classes. These classes included Penicillins, Glycopeptides, Fluoroquinolones, Aminoglycosides, Tetracyclines, Macrolides, Amphenicols, Polymyxins, and Sulfonamides, each differing in mechanism of action and bacterial specificity, as detailed in **Figure 7**. These are presented as both MIC50 in micrograms per ml and percent increase or decrease in susceptibility. Two proteins potentially involved in the dysregulation are highlighted in C although the mechanisms need to be determined.

**Figure 7:**
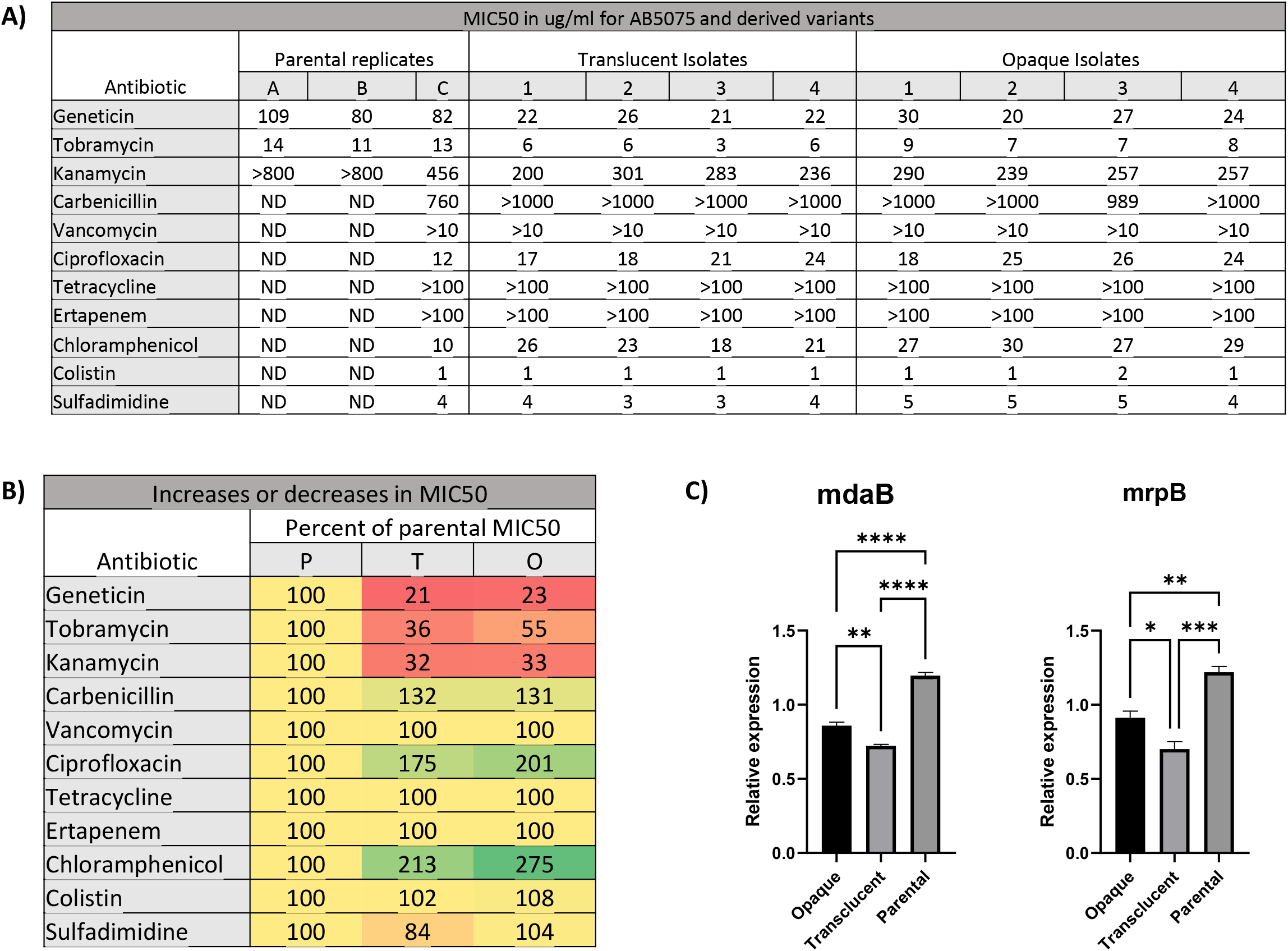
Antibiotic Susceptibility of Acinetobacter baumannii Isolates. **A**) MIC50 for indicated cultures as determined by 96 well plate growth inhibition assay. illustrates the antibiotic resistance profiles of Acinetobacter baumannii isolates with capsule variations. **B**) Numbers are percent MIC50 in relation to parental. **C)** Expression of 2 antibiotic resistance genes (means by group) (mdaB, is modulator of drug activiy B, mrpB, is multi-drug resistance protein B. * indicate one way ANOVA with Tukey’s correction for multiple comparisons.

Predominantly, all isolates demonstrated consistent resistance to carbenicillin, vancomycin, and erythromycin, while they remained susceptible to ciprofloxacin, tetracycline, chloramphenicol, colistin, and sulfadimidine. Interestingly, all translucent isolates exhibited an increased sensitivity to aminoglycosides. Unexpectedly, a similar heightened sensitivity was also observed in the opaque isolates

## Discussion

This study sheds light on the intricate adaptability mechanisms of *A. baumannii*, a pathogen notorious for its resilience and antibiotic resistance (27)(3). While a fairly simple-sounding transposition event may sound commonplace (28-30), the implications of a species evolving to a point where multiple lineages independently make the same genetic mutation simultaneously could be fundamental to our understanding of bacterial adaptation. In the right conditions, parallel evolution may even be inevitable.

These variants arose during our ongoing efforts to improve the immunogenicity of bacterial cultures. A prolonged experiment was being performed as we attempted to generate multiple different bacterial cultures to examine the ability of inactivated whole cell vaccines with unique profiles to protect against challenge in models of pulmonary infection (31-33). From a practical standpoint, this work highlights the need to culture *A. baumannii* under standard conditions to enhance stability and standardize vaccines and experiments. Several promising capsule negative vaccines are also in development.

The concurrent occurrence of multiple genetic events, as unveiled through our genomic and proteomic analyses, underscores the complexity of *A. baumannii’s* response to environmental challenges. These events, far from being isolated, suggest a coordinated network of genetic adaptations that enable the pathogen to rapidly adjust to fluctuating environmental conditions.

Our research further suggests that the pathogen’s antibiotic resistance mechanisms may operate independently of the capsule morphology, a finding that adds another layer to the understanding of its resistance profile. Capsule loss, while significant, is not sole determinant of antibiotic susceptibility. Instead, it may act in concert with, or even mask, other underlying genetic factors that confer resistance. This insight could be crucial, as it indicates that antibiotic susceptibility in *A. baumannii* might often precede or accompany capsule loss, rather than being a direct result of it. Therefore, capsule loss alone, in the absence of other resistance mechanisms, may increase susceptibility to a broader spectrum of antibiotics.

These findings compel us to consider a more nuanced approach to treating *A. baumannii* infections. The dynamic nature of the pathogen’s resistance – influenced by both phenotypic variations and underlying genetic changes – highlights the need for multifaceted therapeutic strategies. Understanding the interplay between environmental cues, phenotypic changes, and genetic rearrangements is pivotal in devising effective treatments and combating the growing challenge of antibiotic resistance in *A. baumannii*

In summary, our study not only contributes to the expanding body of knowledge surrounding *A. baumannii’s* adaptability but also emphasizes the complexity of its response mechanisms, paving the way for innovative approaches in managing infections caused by this adaptable pathogen.

## Materials and Methods

Bacterial culture and generation of isolates. Liquid cultures of AB5075 were cultivated in M9 minimal medium supplemented with 2 mM MgSO4, 0.4% glucose, 0.1 mM CaCl2, and 0.1% casamino acids. This medium is referred to as “M9 medium” throughout the text. 65 mL of M9 minimal medium was inoculated with AB5075 from a freshly streaked plate and incubated overnight at 37°C while shaking at 200 rpm. The following day, the overnight culture was diluted in M9 media to an OD600 of 0.05, and 100 mL was added to 225 cm^2^ polystyrene tissue culture flasks. Media and cultures were mixed and placed in a 37°C incubator without shaking. Media was replaced with fresh M9 media daily while maintaining the biofilm. On day 5, media and floating cells were removed from the flask and replaced with 25 mL of 4°C cold PBS. Cells were scraped into the cold PBS from the bottom of the flask using a plastic cell scraper and collected. Cels were washed twice in cold PBS. These were then titered on 15 cm TSA plates and allowed to grow for 3-5 days. Translucent and opaque regions from the observed biphasic colonies were taken and streaked on TSA. From these, single colonies were obtained and streaked two further times to ensure clonality. For images, agar plates were illuminated from the back using white light (A4 Portable LED Light Box-Amazon) and photographed using an Apple iPhone 8 Plus 12 MP wide camera (28 mm, *f*1.8). Images were obtained in color and rendered grayscale using Microsoft PowerPoint Version 2312.

### Sedimentation Density Gradients

Overnight cultures were layered over Percoll step gradients within transparent 2 mL tubes (Fisher, Hanover, MD). The gradients were composed of 0.75 mL of 60% Percoll layered atop 0.75 mL of 100% Percoll (GE healthcare). Following the preparation, the tubes underwent centrifugation in a microfuge, spinning at 5500 rpm for 5 minutes. To facilitate imaging, the tubes were illuminated from the bottom using white light, with a black background serving as contrast.

### Crystal Violet Staining and Absorbance Measurement

Variants or biological replicates of the parental cells were cultured in 96-well plates (MSP, Atlanta, Georgia). To prepare for staining, cells were first fixed using a mixture of methanol and acetone in a 1:1 ratio (Fisher, Hanover, MD). Subsequently, staining was carried out using a 0.05% crystal violet solution (Fisher, Hanover, MD) in PBS pH 7.4 w/o Ca2+ -Mg2+ (Gibco, Thermo Fisher Scientific, Hanover, MD). After staining, the cells underwent three washes and were then dissolved in 30% glacial acetic acid (Mallinckrodt, St. Louis, MO). The resulting solution was transferred to a clean plate, and absorbance was measured at 600 nm using a SpectraMax 340 plate reader and recorded using SoftPromax software 3.1.

### PacBio sequencing

Libraries made up of barcoded SMRTBells (including DNA barcodes and technical sequences) were created for sequencing using standard protocols. Prepared libraries from several samples were pooled and sequenced on a PacBio Sequel system. The reads were then demultiplexed to separate them by sample and combined if multiple sequencing runs were performed. Subreads were subsequently assembled using Falcon version 1.8.8.

### Antibiotic susceptibility by broth microdilution

Antibiotic susceptibility testing was performed using a modified broth microdilution method in accordance with the Clinical and Laboratory Standards Institute (CLSI) guidelines. Multiple antibiotics were chosen; Carbenicillin (Fisher Scientific 50-213-247), Vancomycin (Millipor-Sigma PHR1732-4X250MG), Ciprofloxacin (Fisher Scientific AC449620050), G418 (Thermo scientific) (Fisher Scientific BP6731), Kanamycin (Fisher Scientific BP906-5), Tobramycin (Thermo scientific) (Fisher Scientific AC455430050), Tetracycline (Fisher Scientific ICN10301125), Ertapenem (Erythromycin) (Fisher Scientific 50-997-767), Chloramphenicol (Fisher Scientific BP904-100), Colistin (Fisher Scientific ICN19415725), and Sulfamethoxazole (Sulfadimidine) (Fisher Scientific AAJ6656522). Overnight cultures of several variants of AB5075 were diluted to approximately 5x10^6 CFU/mL in LB Broth (Fischer bioreagents). 100 uL of the diluted cultures were then combined with 100 uL of antibiotic diluted in LB (final dose specified in figures) in 96-well plates and incubated at 37oC for 18 hours. After incubation, plates were read using a Spectromax 340 plate reader at 600nm to determine optical density.

### Proteomic Analysis via Mass Spectrometry

#### Protein Labeling (for Jon to check)

A TMT labelling set was used to label samples as follows (Thermo Scientific). 100 μg of protein lysate from each sample was digested in trypsin. Pre-digestion and digested samples were run on SDS-PAGE, and the gel was stained to check digestion efficiency. Isobaric labeling was performed using the TMT kit according to the product manual (Table 1). Each set of TMT-labeled peptide mixtures was combined, dried in a vacuum concentrator, and stored at -80 ºC.

#### Fractionation of Labeled Peptides by Basic Reversed-Phase UHPLC

Dried labeled peptides were resuspended in 10 mM TEABC (Thermo Scientific). Labeling efficiency was determined to be above 95%. Fractionation of the TMT labeled peptide mixture was carried out using an Agilent AdvanceBio Column (2.7 μm, 2.1 x 250 mm) and an Agilent UHPLC 1290 system. Separation was performed using a gradient of Solvent B (10 mM TEABC, pH 8.0, 90% ACN) and Solvent A (10 mM TEABC, pH 8.0) at a flow rate of 250 μL/min (Sigma). Eluted fractions were collected into a 96-well plate using a 1260 series auto-sample fraction collector. The 96 eluted fractions were further combined into 12 fractions according to collection time for LC/MS/MS analysis.

#### Nanospray LC/MS/MS Analysis and Database Search

LC/MS/MS analysis was carried out using a Thermo Scientific Q-Exactive hybrid Quadrupole-Orbitrap Mass Spectrometer and a Thermo Dionex UltiMate 3000 RSLCnano System. Each peptide fraction from a set of 12 fractions was loaded onto a peptide trap cartridge at a flow rate of 5 μL/min. Trapped peptides were eluted onto a reversed-phase 20 cm C18 PicoFrit column (New Objective, Woburn, MA) using a linear gradient of acetonitrile (3-36%) in 0.1% formic acid. The elution duration was 110 minutes at a flow rate of 0.3 μL/min. Eluted peptides from the PicoFrit column were ionized and sprayed into the mass spectrometer using a Nanospray Flex Ion Source ES071 (Thermo Scientific) under the following settings: spray voltage, 1.8 kV, capillary temperature, 250°C. Twelve fractions were analyzed sequentially.

The Q Exactive instrument was operated in data-dependent mode to automatically switch between full scan MS and MS/MS acquisition. Survey full scan MS spectra (m/z 350−1800) were acquired in the Orbitrap with 35,000 resolution (m/z 200) after an accumulation of ions to a 3.0 × 10^6^ target value based on predictive automatic gain control (AGC). The maximum injection time was set to 100 ms. The 15 most intense multiply charged ions (z ≥ 2) were sequentially isolated and fragmented in the octopole collision cell by higher-energy collisional dissociation (HCD) using normalized HCD collision energy 35 with an AGC target of 1.0 ×10^5^ and a maximum injection time of 400 ms at 17,500 resolution. The isolation window was set to 2, and the fixed first mass was 120 m/z. Dynamic exclusion was set to 20 seconds. Charge state screening was enabled to reject unassigned and 1+, 7+, 8+, and >8+ ions.

One set of 12 MS raw data files acquired from the analysis of 12 fractions was searched against the Acinetobacter baumannii protein sequences database obtained from the UniprotKB website using the Proteome Discoverer 2.4 software (Thermo, San Jose, CA) based on the SEQUEST and percolator algorithms.

### Kaptive

The Kaptive program and database was used to analyze Acinetobacter K loci (18). Kaptive reports information about surface polysaccharide loci for *Klebsiella pneumoniae* species complex and *Acinetobacter baumannii* genome assemblies.

## Acknowledgments

We would like to thank Garth Ehrlich, Rachel Ehrlich of the Department of Microbiology and Immunology, Drexel University College of Medicine for PacBio sequencing. We would also like to thank Robert Shanks, Department of Microbiology and Molecular Genetics, University of Pittsburg for helpful discussions.

## References

1. CDC. Antibiotic Resistance Threats in the United States, 2019 (2019 AR Threats Report). U.S. Center for Disease Control, The Antibiotic Resistance Coordination and Strategy Unit Do, Healthcare Quality Promotion NCfEaZID, Prevention. CfDCa; 2019.

2. CDC. Acinetobacter in Healthcare Settings 2020 [Available from: https://www.cdc.gov/hai/organisms/acinetobacter.html.

3. Peleg AY, Seifert H, Paterson DL. Acinetobacter baumannii: emergence of a successful pathogen. Clin Microbiol Rev. 2008;21(3):538–82.

4. Harding CM, Hennon SW, Feldman MF. Uncovering the mechanisms of Acinetobacter baumannii virulence. Nature reviews Microbiology. 2018;16(2):91–102.

5. Tipton KA, Dimitrova D, Rather PN. Phase-Variable Control of Multiple Phenotypes in Acinetobacter baumannii Strain AB5075. Journal of bacteriology. 2015;197(15):2593–9.

6. Tipton KA, Rather PN. An ompR-envZ Two-Component System Ortholog Regulates Phase Variation, Osmotic Tolerance, Motility, and Virulence in Acinetobacter baumannii Strain AB5075. Journal of bacteriology. 2017;199(3).

7. Mushtaq F, Nadeem A, Yabrag A, Bala A, Karah N, Zlatkov N, et al. Colony phase variation switch modulates antimicrobial tolerance and biofilm formation in Acinetobacter baumannii. Microbiol Spectr. 2024;12(2):e0295623.

8. Chin CY, Tipton KA, Farokhyfar M, Burd EM, Weiss DS, Rather PN. A high-frequency phenotypic switch links bacterial virulence and environmental survival in Acinetobacter baumannii. Nature microbiology. 2018;3(5):563–9.

9. Valcek A, Collier J, Botzki A, Van der Henst C. Acinetobase: the comprehensive database and repository of Acinetobacter strains. Database (Oxford). 2022;2022.

10. Akoolo L, Pires S, Kim J, Parker D. The Capsule of Acinetobacter baumannii Protects against the Innate Immune Response. J Innate Immun. 2022;14(5):543–54.

11. Whiteway C, Valcek A, Philippe C, Strazisar M, De Pooter T, Mateus I, et al. Scarless excision of an insertion sequence restores capsule production and virulence in Acinetobacter baumannii. The ISME journal. 2022;16(5):1473–7.

12. Anderson SE, Rather PN. Distinguishing Colony Opacity Variants and Measuring Opacity Variation in Acinetobacter baumannii. Methods in molecular biology (Clifton, NJ). 2019;1946:151–7.

13. Dehbanipour R, Ghalavand Z. Acinetobacter baumannii: Pathogenesis, virulence factors, novel therapeutic options and mechanisms of resistance to antimicrobial agents with emphasis on tigecycline. J Clin Pharm Ther. 2022;47(11):1875–84.

14. Weber BS, Harding CM, Feldman MF. Pathogenic Acinetobacter: from the Cell Surface to Infinity and Beyond. Journal of bacteriology. 2015;198(6):880–7.

15. Geisinger E, Huo W, Hernandez-Bird J, Isberg RR. Acinetobacter baumannii: Envelope Determinants That Control Drug Resistance, Virulence, and Surface Variability. Annual review of microbiology. 2019;73:481–506.

16. Doi Y, Murray GL, Peleg AY. Acinetobacter baumannii: evolution of antimicrobial resistance-treatment options. Semin Respir Crit Care Med. 2015;36(1):85–98.

17. Novović K, Mihajlović S, Dinić M, Malešević M, Miljković M, Kojić M, et al. Acinetobacter spp. porin Omp33-36: Classification and transcriptional response to carbapenems and host cells. PLoS One. 2018;13(8):e0201608.

18. Wyres KL, Cahill SM, Holt KE, Hall RM, Kenyon JJ. Identification of Acinetobacter baumannii loci for capsular polysaccharide (KL) and lipooligosaccharide outer core (OCL) synthesis in genome assemblies using curated reference databases compatible with Kaptive. Microbial genomics. 2020;6(3).

19. Bailey SF, Blanquart F, Bataillon T, Kassen R. What drives parallel evolution?: How population size and mutational variation contribute to repeated evolution. BioEssays : news and reviews in molecular, cellular and developmental biology. 2017;39(1):1–9.

20. Toprak E, Veres A, Michel JB, Chait R, Hartl DL, Kishony R. Evolutionary paths to antibiotic resistance under dynamically sustained drug selection. Nat Genet. 2011;44(1):101–5.

21. Scribner MR, Santos-Lopez A, Marshall CW, Deitrick C, Cooper VS. Parallel Evolution of Tobramycin Resistance across Species and Environments. mBio. 2020;11(3).

22. Shabbir MA, Hao H, Shabbir MZ, Wu Q, Sattar A, Yuan Z. Bacteria vs. Bacteriophages: Parallel Evolution of Immune Arsenals. Frontiers in microbiology. 2016;7:1292.

23. Lenski RE. Convergence and Divergence in a Long-Term Experiment with Bacteria. Am Nat. 2017;190(S1):S57–S68.

24. Lenski RE. Experimental evolution and the dynamics of adaptation and genome evolution in microbial populations. The ISME journal. 2017;11(10):2181–94.

25. Maier RM. Biosurfactants: evolution and diversity in bacteria. Adv Appl Microbiol. 2003;52:101–21.

26. Darling AE, Mau B, Perna NT. progressiveMauve: multiple genome alignment with gene gain, loss and rearrangement. PLoS One. 2010;5(6):e11147.

27. Murray CJl, Kevin Shunji Ikuta, Fablina Sharara, Lucien Swetschinski, Gisela Robles Aguilar, Authia Gray, Chieh Han, Catherine Bisignano, Puja Rao, Eve Wool, Sarah C Johnson, Annie J Browne, Michael Give Chipeta, Frederick Fell, Sean Hackett, Georgina Haines-Woodhouse, Bahar H Kashef Hamadani, Emmanuelle A P Kumaran, Barney McManigal, Ramesh Agarwal, Samuel Akech, Samuel Albertson, John Amuasi, Jason Andrews, Aleskandr Aravkin, Elizabeth Ashley, Freddie Bailey, Stephen Baker, Buddha Basnyat, Adrie Bekker, Rose Bender, Adhisivam Bethou, Julia Bielicki, Suppawat Boonkasidecha, James Bukosia, Cristina Carvalheiro, Carlos Castañeda-Orjuela, Vilada Chansamouth, Suman Chaurasia, Sara Chiurchiù, Fazle Chowdhury, Aislinn J Cook, Ben Cooper, Tim R Cressey, Elia Criollo-Mora, Matthew Cunningham, Saffiatou Darboe, Nicholas P J Day, Maia De Luca, Klara Dokova, Angela Dramowski, Susanna J Dunachie, Tim Eckmanns, Daniel Eibach, Amir Emami, Nicholas Feasey, Natasha Fisher-Pearson, Karen Forrest, Denise Garrett, Petra Gastmeier, Ababi Zergaw Giref, Rachel Claire Greer, Vikas Gupta, Sebastian Haller, Andrea Haselbeck, Simon I Hay, Marianne Holm, Susan Hopkins, Kenneth C Iregbu, Jan Jacobs, Daniel Jarovsky, Fatemeh Javanmardi, Meera Khorana, Niranjan Kissoon, Elsa Kobeissi, Tomislav Kostyanev, Fiorella Krapp, Ralf Krumkamp, Ajay Kumar, Hmwe Hmwe Kyu, Cherry Lim, Direk Limmathurotsakul, Michael James Loftus, Miles Lunn, Jianing Ma, Neema Mturi, Tatiana Munera-Huertas, Patrick Musicha, Marisa Marcia Mussi-Pinhata, Tomoka Nakamura, Ruchi Nanavati, Sushma Nangia, Paul Newton, Chanpheaktra Ngoun, Amanda Novotney, Davis Nwakanma, Christina W Obiero, Antonio Olivas-Martinez, Piero Olliaro, Ednah Ooko, Edgar Ortiz-Brizuela, Anton Yariv Peleg, Carlo Perrone, Nishad Plakkal, Alfredo Ponce-de-Leon, Mathieu Raad, Tanusha Ramdin, Amy Riddell, Tamalee Roberts, Julie Victoria Robotham, Anna Roca, Kristina E Rudd, Neal Russell, Jesse Schnall, John Anthony Gerard Scott, Madhusudhan Shivamallappa, Jose Sifuentes-Osornio, Nicolas Steenkeste, Andrew James Stewardson, Temenuga Stoeva, Nidanuch Tasak, Areerat Thaiprakong, Guy Thwaites, Claudia Turner, Paul Turner, H Rogier van Doorn, Sithembiso Velaphi, Avina Vongpradith, Huong Vu, Timothy Walsh, Seymour Waner, Tri Wangrangsimakul, Teresa Wozniak, Peng Zheng, Benn Sartorius, Alan D Lopez, Andy Stergachis, Catrin Moore, Christiane Dolecek, Mohsen Naghavi. Global burden of bacterial antimicrobial resistance in 2019: a systematic analysis. Lancet (London, England). 2022;399(10325):629–55.

28. Hamidian M, Hall RM. The AbaR antibiotic resistance islands found in Acinetobacter baumannii global clone 1 - Structure, origin and evolution. Drug resistance updates : reviews and commentaries in antimicrobial and anticancer chemotherapy. 2018;41:26–39.

29. Peters JE, Fricker AD, Kapili BJ, Petassi MT. Heteromeric transposase elements: generators of genomic islands across diverse bacteria. Molecular microbiology. 2014;93(6):1084–92.

30. Frost LS, Leplae R, Summers AO, Toussaint A. Mobile genetic elements: the agents of open source evolution. Nature reviews Microbiology. 2005;3(9):722–32.

31. Dollery SJ, Zurawski DV, Gaidamakova EK, Matrosova VY, Tobin JK, Wiggins TJ, et al. Radiation-Inactivated Acinetobacter baumannii Vaccine Candidates. Vaccines (Basel). 2021;9(2).

32. Dollery SJ, Harro JM, Wiggins TJ, Wille BP, Kim PC, Tobin JK, et al. Select Whole-Cell Biofilm-Based Immunogens Protect against a Virulent Staphylococcus Isolate in a Stringent Implant Model of Infection. Vaccines (Basel). 2022;10(6):833.

33. Dollery SJ, Zurawski DV, Bushnell RV, Tobin JK, Wiggins TJ, MacLeod DA, et al. Whole-cell vaccine candidates induce a protective response against virulent Acinetobacter baumannii. Front Immunol. 2022;13:941010.

