## Supplemental Table 1 for "Evidence for high-frequency parallel evolution in virulent A. *baumannii* cultures"

| Protein | Parental | Translucent | Log2FC | p-value |
|---|---|---|---|---|
| oxidoreductase short chain dehydrogenase/reductase family. | 1.217599 | 0.742828 | -0.71294 | 2.75E-07 |
| nitroreductase family protein...2257 | 1.077242 | 0.867772 | -0.31195 | 3.91E-06 |
| pirin domain protein...1437 | 1.165933 | 0.919206 | -0.34302 | 4.62E-06 |
| succinate dehydrogenase, flavoprotein subunit | 0.894669 | 1.095668 | 0.292384 | 1.05E-05 |
| ATP synthase F0, B subunit | 0.988179 | 1.09136 | 0.143283 | 1.08E-05 |
| peptidase M48 family protein | 1.053808 | 0.942141 | -0.1616 | 1.14E-05 |
| adenosylmethionine-8-amino-7-oxononanoate transaminase | 1.152658 | 0.911207 | -0.33911 | 1.48E-05 |
| OmpW family protein...1026 | 0.725247 | 1.498813 | 1.047275 | 1.7E-05 |
| peptidase M16 domain protein | 1.123473 | 0.937381 | -0.26126 | 1.81E-05 |
| putative anti-anti-sigma factor | 0.714768 | 1.060766 | 0.569561 | 1.94E-05 |
| hypothetical protein ABUW_2220 | 1.17674 | 0.908673 | -0.37296 | 2.01E-05 |
| imidazoleglycerol-phosphate dehydratase | 0.92792 | 1.069087 | 0.204307 | 2.17E-05 |
| transcriptional regulator, LysR family...1214 | 1.069991 | 0.802684 | -0.41469 | 2.38E-05 |
| hypothetical protein ABUW_3825 | 1.246119 | 0.501992 | -1.31171 | 2.92E-05 |
| aspartate aminotransferase A | 0.916343 | 1.17402 | 0.357498 | 3.02E-05 |
| trehalose-6-phosphate synthase | 0.598205 | 1.352291 | 1.176693 | 3.17E-05 |
| hypothetical protein ABUW_2058 | 1.119204 | 0.764648 | -0.54961 | 3.45E-05 |
| TonB-dependent copper receptor | 0.796835 | 1.140306 | 0.517068 | 4.69E-05 |
| carbamoyl-phosphate synthase, large subunit | 0.855272 | 1.277685 | 0.579076 | 4.83E-05 |
| Protein incC (plasmid) | 1.096769 | 0.858798 | -0.35287 | 5.31E-05 |
| NADH dehydrogenase I chain I 2Fe-2S ferredoxin-related | 0.939555 | 1.090165 | 0.214498 | 5.96E-05 |
| P-hydroxyphenylacetate hydroxylase C1:reductase componer | 0.739702 | 1.393331 | 0.913522 | 6.07E-05 |
| hypothetical protein ABUW_1042 | 1.099006 | 0.950938 | -0.20878 | 6.77E-05 |
| excinuclease ABC, C subunit (EsvL) | 1.08821 | 0.947461 | -0.19982 | 7.07E-05 |
| amidohydrolase...1632 | 1.029269 | 0.887925 | -0.21311 | 7.16E-05 |
| hypothetical protein ABUW_0887 | 0.807227 | 1.054196 | 0.385098 | 7.66E-05 |
| hypothetical protein ABUW_2459 | 1.202753 | 0.963672 | -0.31973 | 8.18E-05 |
| cold-shock DNA-binding domain protein...385 | 0.863415 | 1.113557 | 0.367049 | 8.21E-05 |
| 2,3-bisphosphoglycerate-independent phosphoglycerate mut | 0.924697 | 1.104233 | 0.255992 | 8.54E-05 |
| dihydroxy-acid dehydratase...2283 | 0.939833 | 1.081901 | 0.203092 | 8.7E-05 |
| adenylosuccinate lyase | 0.914237 | 1.08029 | 0.240778 | 9.06E-05 |
| methylisocitrate lyase | 0.897206 | 1.068483 | 0.252053 | 9.56E-05 |
| L-asparaginase 2 | 1.164188 | 0.910093 | -0.35524 | 9.93E-05 |
| nicotinate-nucleotide--dimethylbenzimidazole phosphoribosy | 0.957882 | 1.112347 | 0.215687 | 0.000105 |
| polyprenyl synthetase | 0.868378 | 1.154601 | 0.410999 | 0.000106 |
| phosphoribosylamine--glycine ligase | 0.970952 | 1.047864 | 0.10998 | 0.000122 |
| fumarylacetoacetase | 0.893201 | 1.159142 | 0.376001 | 0.000124 |
| alanyl-tRNA synthetase | 0.954443 | 1.042058 | 0.126704 | 0.00013 |
| glutathione S-transferase...1636 | 1.118223 | 0.780098 | -0.51948 | 0.000131 |
| lysine exporter protein | 0.587175 | 1.508668 | 1.361413 | 0.000133 |
| hypothetical protein ABUW_0435 | 0.968303 | 1.187318 | 0.294175 | 0.000138 |
| 2-oxoglutarate dehydrogenase, E2 component, dihydrolipoan | 0.949615 | 1.1013 | 0.213793 | 0.000139 |
| ribonuclease R | 0.936018 | 1.078632 | 0.204594 | 0.00014 |
| amidophosphoribosyltransferase | 0.945979 | 1.067267 | 0.174041 | 0.000146 |
| phosphoribosylaminoimidazole carboxylase, ATPase subunit | 0.969386 | 1.07799 | 0.1532 | 0.000151 |
| siderophore biosynthesis protein...178 | 0.630699 | 1.321086 | 1.066701 | 0.000167 |

| | | | | |
|---|---|---|---|---|
| hypothetical protein ABUW_3230 | 1.091958 | 0.780778 | -0.48393 | 0.000169 |
| TonB-dependent receptor...2157 | 0.905126 | 1.233656 | 0.446749 | 0.000171 |
| hypothetical protein ABUW_1317 | 1.092307 | 0.87007 | -0.32818 | 0.000176 |
| carbamoyl-phosphate synthase, small subunit | 0.817752 | 1.348346 | 0.721455 | 0.000183 |
| phospholipase C, phosphocholine-specific...768 | 1.187926 | 0.820096 | -0.53458 | 0.000202 |
| hypothetical protein ABUW_3433 | 1.573988 | 0.793586 | -0.98797 | 0.000203 |
| hypothetical protein ABUW_0808 | 1.019514 | 0.8943 | -0.18905 | 0.000213 |
| hypothetical protein ABUW_3828 | 0.939541 | 1.097955 | 0.22479 | 0.00022 |
| homoserine dehydrogenase...2365 | 0.914217 | 1.163066 | 0.347325 | 0.000233 |
| ribonuclease III | 0.987637 | 1.139545 | 0.206406 | 0.000234 |
| TonB protein | 0.798785 | 1.216347 | 0.606677 | 0.000234 |
| outer-membrane receptor for Fe(III)-coprogen,Fe(III)-ferrioxa | 0.853238 | 1.214689 | 0.509567 | 0.000241 |
| hypothetical protein ABUW_1282 | 0.977569 | 1.098889 | 0.168776 | 0.000241 |
| flavin-containing monooxygenase FMO | 1.043404 | 0.903445 | -0.20779 | 0.000258 |
| dethiobiotin synthase | 1.102959 | 0.898626 | -0.29559 | 0.000258 |
| trehalose-phosphatase | 0.679279 | 1.338765 | 0.978825 | 0.00026 |
| isoleucyl-tRNA synthetase | 0.957624 | 1.075918 | 0.168037 | 0.000264 |
| TonB-dependent siderophore receptor...1752 | 0.881042 | 1.183106 | 0.425296 | 0.000267 |
| general substrate transporter:Major facilitator superfamily | 0.848212 | 1.102386 | 0.378132 | 0.000273 |
| C4-dicarboxylate transport protein...365 | 0.906342 | 1.425761 | 0.653605 | 0.000277 |
| xanthine phosphoribosyltransferase | 1.12819 | 0.820736 | -0.45902 | 0.000278 |
| long-chain fatty-acid-CoA ligase | 1.002882 | 1.108494 | 0.144449 | 0.000281 |
| glutathione S-transferase...1840 | 1.048662 | 0.932696 | -0.16907 | 0.000293 |
| modulator of drug activity B | 1.195831 | 0.722184 | -0.72758 | 0.000296 |
| nicotinate-nucleotide diphosphorylase | 0.914368 | 1.117924 | 0.289975 | 0.000302 |
| short-chain dehydrogenase/reductase...542 | 1.142453 | 0.995556 | -0.19856 | 0.00031 |
| phosphoribosylaminoimidazole carboxylase, catalytic subunit | 0.965854 | 1.097128 | 0.183855 | 0.000323 |
| pH adaptation potassium efflux system transmembrane prote | 0.905527 | 1.214963 | 0.424083 | 0.000327 |
| transcriptional regulator, PadR family | 1.070207 | 0.918712 | -0.22021 | 0.000336 |
| malate dehydrogenase...2484 | 0.891935 | 1.086773 | 0.28504 | 0.000338 |
| citrate transporter | 0.702013 | 1.709318 | 1.283851 | 0.000339 |
| urocanate hydratase | 1.092899 | 0.91049 | -0.26345 | 0.000344 |
| thymidylate synthase | 0.939828 | 1.142753 | 0.282045 | 0.000346 |
| two-component system sensor protein KdpD | 0.838775 | 1.316985 | 0.650883 | 0.000348 |
| heat shock protein 33 | 1.040158 | 0.946368 | -0.13633 | 0.000353 |
| fructose-1,6-bisphosphatase | 1.86743 | 0.586313 | -1.67131 | 0.00037 |
| succinate dehydrogenase iron-sulfur subunit | 0.88368 | 1.100566 | 0.31665 | 0.000376 |
| family 1 glycosyl transferase...2077 | 1.032032 | 0.831428 | -0.31182 | 0.000377 |
| hydrolase, alpha/beta fold family | 1.065542 | 0.821684 | -0.37493 | 0.00041 |
| peptidase M24 | 1.082368 | 0.92927 | -0.22002 | 0.000411 |
| preprotein translocase, SecA subunit | 0.942704 | 1.06331 | 0.173685 | 0.000418 |
| glutamyl-tRNA(Gln) amidotransferase, B subunit | 0.94812 | 1.10971 | 0.227042 | 0.000418 |
| major facilitator family transporter...492 | 0.843427 | 1.220364 | 0.532977 | 0.000418 |
| NADH dehydrogenase I chain E | 0.961839 | 1.039303 | 0.111749 | 0.000418 |
| thioesterase domain protein | 0.76874 | 1.21863 | 0.664694 | 0.000423 |
| signal peptide peptidase SppA, 36K type | 0.925729 | 1.101684 | 0.251049 | 0.000423 |
| alkyl hydroperoxide reductase, F subunit | 1.097743 | 0.930466 | -0.23852 | 0.000438 |

| | | | | |
|---|---|---|---|---|
| tryptophanyl-tRNA synthetase | 0.944914 | 1.080564 | 0.19353 | 0.000446 |
| coenzyme PQQ biosynthesis protein B | 1.059647 | 0.754601 | -0.4898 | 0.000447 |
| pyridoxal phosphate biosynthetic protein PdxJ | 0.998153 | 1.102098 | 0.142919 | 0.000451 |
| poly(R)-hydroxyalkanoic acid synthase | 1.135248 | 0.964015 | -0.23588 | 0.000455 |
| hypothetical protein ABUW_0360 | 1.443132 | 0.905154 | -0.67297 | 0.000457 |
| short-chain fatty acid transporter | 0.769682 | 1.197849 | 0.638113 | 0.000457 |
| cell division protein ZipA | 0.830188 | 1.153151 | 0.474072 | 0.000466 |
| transcriptional regulator, MarR-family...738 | 0.860946 | 1.324335 | 0.621274 | 0.000473 |
| hypothetical protein ABUW_0116 | 0.939014 | 1.247511 | 0.409835 | 0.000478 |
| NADH dehydrogenase | 1.067767 | 0.795757 | -0.4242 | 0.00049 |
| copper resistance protein A | 1.076109 | 0.896924 | -0.26277 | 0.000499 |
| methionine synthase | 0.900605 | 1.105139 | 0.295262 | 0.000511 |
| family 1 glycosyl transferase...1349 | 0.668474 | 1.577714 | 1.238892 | 0.000519 |
| hypothetical protein ABUW_1594 | 0.621823 | 1.174351 | 0.917287 | 0.000545 |
| cystathionine beta-lyase PLP-dependent | 0.895122 | 1.056302 | 0.238866 | 0.000574 |
| aspartate carbamoyltransferase | 0.903075 | 1.135026 | 0.329807 | 0.000577 |
| 2-hydroxy-3-oxopropionate reductase | 1.064145 | 0.909185 | -0.22705 | 0.000577 |
| TonB dependent outer membrane siderophore receptor prot | 0.801769 | 1.421817 | 0.826477 | 0.000578 |
| ATP-dependent chaperone ClpB | 1.113424 | 0.81682 | -0.44691 | 0.000585 |
| guanylate kinase | 1.11363 | 0.89591 | -0.31384 | 0.000588 |
| polysaccharide deacetylase...252 | 0.907838 | 1.123831 | 0.307918 | 0.000597 |
| undecaprenyl diphosphate synthase | 1.172047 | 0.897978 | -0.38428 | 0.000614 |
| dimethylmenaquinone methyltransferase | 0.803864 | 1.269533 | 0.659275 | 0.000621 |
| benzoate transport porin BenP | 0.957213 | 1.108634 | 0.211871 | 0.000622 |
| stearoyl-CoA 9-desaturase | 0.868496 | 1.020727 | 0.233006 | 0.000625 |
| hypoxanthine phosphoribosyltransferase | 1.077212 | 0.915489 | -0.23469 | 0.000637 |
| hypothetical protein ABUW_1379 | 1.054933 | 0.91468 | -0.20581 | 0.000642 |
| rhodanese domain protein...1794 | 1.004633 | 1.091675 | 0.119874 | 0.00065 |
| TonB-dependent receptor...2285 | 0.928989 | 1.206929 | 0.377607 | 0.000655 |
| acyl-CoA dehydrogenase, middle domain protein | 0.873693 | 1.398406 | 0.678585 | 0.000668 |
| aromatic-amino-acid aminotransferase | 0.854222 | 1.160673 | 0.442278 | 0.000683 |
| CsuA/B | 0.82877 | 1.269038 | 0.614691 | 0.000685 |
| MotA/TolQ/ExbB proton channel | 0.873977 | 1.206755 | 0.465467 | 0.000694 |
| phenylalanyl-tRNA synthetase, alpha subunit | 0.915623 | 1.063699 | 0.216264 | 0.000732 |
| dihydrolipoamide acetyltransferase | 0.984811 | 0.890361 | -0.14546 | 0.000736 |
| phenylacetic acid degradation protein PaaN | 0.8246 | 1.216745 | 0.561261 | 0.000775 |
| threonyl-tRNA synthetase | 0.98643 | 1.060583 | 0.104569 | 0.000776 |
| ornithine carbamoyltransferase | 1.14481 | 0.829657 | -0.46452 | 0.000776 |
| transcriptional regulator, AraC family...447 | 1.105759 | 0.919565 | -0.26601 | 0.000789 |
| hypothetical protein ABUW_2042 | 0.63754 | 1.091645 | 0.775917 | 0.000793 |
| methionine-R-sulfoxide reductase | 0.84825 | 1.118305 | 0.398753 | 0.000813 |
| short chain dehydrogenase...1126 | 1.085983 | 0.923823 | -0.23331 | 0.000843 |
| chaperonin GroS | 1.084276 | 0.926559 | -0.22678 | 0.000858 |
| biotin synthase | 1.255587 | 0.932297 | -0.4295 | 0.000865 |
| acetolactate synthase, large subunit | 0.949356 | 1.083481 | 0.190653 | 0.000882 |
| aconitate hydratase 2 | 0.958484 | 1.092016 | 0.188169 | 0.000884 |
| histidyl-tRNA synthetase | 0.95053 | 1.036242 | 0.124558 | 0.000911 |

| | | | | |
|---|---|---|---|---|
| ABC transporter, ATP-binding protein...1167 | 1.141803 | 0.827922 | -0.46375 | 0.000915 |
| thiosulfate-binding protein...537 | 1.087024 | 0.869645 | -0.32189 | 0.000923 |
| anti-sigma factor antagonist | 0.984315 | 1.082917 | 0.137731 | 0.000924 |
| aldose 1-epimerase | 1.078087 | 0.816949 | -0.40016 | 0.000928 |
| hypothetical protein ABUW_3292 | 0.979137 | 1.085145 | 0.148305 | 0.000938 |
| biopolymer transport protein | 1.162579 | 0.773986 | -0.58695 | 0.000939 |
| threonine synthase | 0.8924 | 1.11198 | 0.317369 | 0.000993 |
| hypothetical protein ABUW_2655 | 1.071266 | 0.831218 | -0.36602 | 0.001003 |
| glycine hydroxymethyltransferase | 0.879128 | 1.145177 | 0.381426 | 0.001019 |
| putative integrase/recombinase protein | 0.900671 | 1.327617 | 0.559767 | 0.00105 |
| uracil-DNA glycosylase | 1.027409 | 0.948569 | -0.11519 | 0.001085 |
| hypothetical protein ABUW_2985 | 0.636585 | 1.541699 | 1.276097 | 0.001093 |
| acyl coenzyme A reductase | 1.054748 | 0.915906 | -0.20363 | 0.001096 |
| 5'/3'-nucleotidase SurE | 0.906482 | 1.120738 | 0.306099 | 0.001117 |
| delta-aminolevulinic acid dehydratase | 0.894263 | 1.099905 | 0.298609 | 0.00112 |
| type III restriction enzyme, res subunit | 1.019328 | 0.894962 | -0.18772 | 0.001128 |
| hypothetical protein ABUW_3147 | 0.884173 | 1.252846 | 0.502808 | 0.001142 |
| hypothetical protein ABUW_2673 | 0.670662 | 0.984232 | 0.553414 | 0.001145 |
| hypothetical protein ABUW_1987 | 0.953831 | 1.324184 | 0.473298 | 0.001153 |
| aldehyde dehydrogenase...2355 | 1.006084 | 0.914203 | -0.13816 | 0.001155 |
| esterase...2063 | 1.067227 | 0.933634 | -0.19294 | 0.001167 |
| 3-oxoadipate enol-lactonase...835 | 0.893564 | 1.069841 | 0.259754 | 0.001168 |
| hypothetical protein ABUW_0210 | 1.058069 | 0.909249 | -0.21869 | 0.001189 |
| pantoate--beta-alanine ligase | 0.943261 | 1.077545 | 0.192018 | 0.001206 |
| CsuE | 0.873392 | 1.097278 | 0.329227 | 0.001239 |
| phenylalanyl-tRNA synthetase, beta subunit | 0.962825 | 1.067628 | 0.149063 | 0.001255 |
| peptide chain release factor 3 | 0.925536 | 1.142489 | 0.30382 | 0.001269 |
| dihydrolipoamide dehydrogenase...1848 | 1.020929 | 0.938544 | -0.12139 | 0.001277 |
| glycerol-3-phosphate acyltransferase | 0.937391 | 1.113016 | 0.247752 | 0.001284 |
| transcriptional regulator, GntR family...546 | 0.960871 | 1.035793 | 0.10832 | 0.001329 |
| integration host factor, alpha subunit | 1.1308 | 0.895235 | -0.33701 | 0.001337 |
| ribosomal protein L36 | 1.542741 | 0.818866 | -0.9138 | 0.00135 |
| tail tape measure protein...1458 | 1.138547 | 0.922366 | -0.30378 | 0.001355 |
| pH adaptation potassium efflux system protein C | 0.825847 | 1.164514 | 0.495782 | 0.001359 |
| immunoreactive 21 kD antigen PG10 | 0.889502 | 1.095374 | 0.300354 | 0.001397 |
| tyrosyl-tRNA synthetase | 0.963168 | 1.066065 | 0.146436 | 0.001403 |
| maleylacetoacetate isomerase | 0.926042 | 1.096793 | 0.244142 | 0.001437 |
| saccharopine dehydrogenase | 1.059102 | 0.93374 | -0.18175 | 0.001446 |
| multidrug resistance protein B | 1.219702 | 0.701038 | -0.79896 | 0.001478 |
| metallo-beta-lactamase family protein...1433 | 0.670771 | 1.053477 | 0.651266 | 0.001527 |
| 2,3,4,5-tetrahydropyridine-2,6-dicarboxylate N-succinyltransf | 0.928206 | 1.128656 | 0.282089 | 0.001541 |
| putative dihydrodipicolinate synthase | 1.061997 | 0.861994 | -0.30103 | 0.001562 |
| lipid A export permease/ATP-binding protein MsbA | 0.95809 | 0.995052 | 0.054611 | 0.001569 |
| hypothetical protein ABUW_0028 | 0.54855 | 0.456849 | -0.2639 | 0.001573 |
| hydroxypyruvate isomerase | 1.057504 | 0.940827 | -0.16866 | 0.00158 |
| biopolymer transport protein ExbD/TolR | 0.913697 | 1.113022 | 0.284695 | 0.001593 |
| lipoic acid synthetase...1311 | 1.056312 | 0.958156 | -0.1407 | 0.001659 |

| | | | | |
|---|---|---|---|---|
| phenylacetate-CoA oxygenase, PaaG subunit | 0.847207 | 1.039453 | 0.295039 | 0.001659 |
| nudix hydrolase...683 | 0.927435 | 1.120873 | 0.273304 | 0.001669 |
| hypothetical protein ABUW_0469 | 0.93862 | 1.170638 | 0.318683 | 0.001704 |
| sulfate transporter | 0.880236 | 1.177976 | 0.420348 | 0.001716 |
| hypothetical protein ABUW_0699 | 0.928906 | 1.064967 | 0.197203 | 0.001738 |
| acyl-CoA synthase...1459 | 0.927721 | 1.010562 | 0.123395 | 0.001766 |
| ABC transporter ATP-binding protein...2326 | 0.955426 | 1.068685 | 0.16162 | 0.001766 |
| type I secretion outer membrane protein | 1.10756 | 0.987838 | -0.16504 | 0.001817 |
| fatty acid desaturase...479 | 0.89948 | 1.032579 | 0.19909 | 0.001824 |
| short-chain dehydrogenase/reductase...1982 | 0.931343 | 1.200066 | 0.365729 | 0.001834 |
| FAD linked oxidase domain protein | 0.931542 | 1.048739 | 0.170964 | 0.001855 |
| transcriptional regulator, HxlR family | 1.091383 | 0.862244 | -0.33999 | 0.001855 |
| toluene tolerance protein | 1.076256 | 1.033631 | -0.0583 | 0.00188 |
| hypothetical protein ABUW_0882 | 0.889605 | 1.053568 | 0.244046 | 0.00189 |
| GatB/Yqey domain protein | 1.062013 | 0.977252 | -0.12 | 0.001892 |
| 2-methylcitrate synthase | 0.893626 | 1.032637 | 0.208591 | 0.00193 |
| peptidase S1C, Do | 1.058665 | 0.93841 | -0.17395 | 0.001944 |
| single-stranded-DNA-specific exonuclease RecJ | 0.907796 | 1.09876 | 0.275436 | 0.00195 |
| ribosomal large subunit pseudouridine synthase C | 0.885206 | 1.102189 | 0.316286 | 0.001981 |
| Uridylate kinase | 0.964868 | 1.089327 | 0.175034 | 0.001994 |
| coenzyme PQQ synthesis protein D | 1.054973 | 0.951676 | -0.14866 | 0.002035 |
| phosphoenolpyruvate carboxylase | 0.941796 | 1.129846 | 0.262641 | 0.002047 |
| hypothetical protein ABUW_0133 | 1.142787 | 0.895731 | -0.35142 | 0.002049 |
| phospholipase | 0.956759 | 1.055472 | 0.141661 | 0.00207 |
| RNA methyltransferase, TrmH family, group 3 | 1.072012 | 0.947592 | -0.17798 | 0.002082 |
| glutathione S-transferase...859 | 0.974902 | 0.747593 | -0.38301 | 0.002102 |
| putative RND family drug transporter...1660 | 1.055861 | 0.941486 | -0.16541 | 0.002167 |
| heme oxygenase-like protein...1519 | 0.707304 | 1.152089 | 0.70385 | 0.002169 |
| hypothetical protein ABUW_2134 | 1.127609 | 0.907826 | -0.31278 | 0.002183 |
| delta-1-pyrroline-5-carboxylate dehydrogenase | 0.901551 | 1.039156 | 0.204931 | 0.002185 |
| ABC transporter, ATP-binding protein...719 | 1.265129 | 0.730587 | -0.79216 | 0.002193 |
| type IV pilus biogenesis/stability protein PilW | 1.041203 | 0.913366 | -0.18899 | 0.002206 |
| ErfK/YbiS/YcfS/YnhG family | 0.837531 | 1.074816 | 0.359876 | 0.002208 |
| transcriptional Regulator, AraC family | 0.980435 | 1.135041 | 0.211251 | 0.002212 |
| hypothetical protein ABUW_0375 | 0.893518 | 1.202263 | 0.428185 | 0.002266 |
| hypothetical protein ABUW_1360 | 1.146205 | 0.942503 | -0.2823 | 0.002278 |
| glutamine synthetase, type I | 0.821895 | 1.075692 | 0.388238 | 0.00229 |
| peptidyl-prolyl cis-trans isomerase...1578 | 1.052937 | 0.948876 | -0.15013 | 0.002317 |
| GTP-binding protein Obg/CgtA | 1.097876 | 0.868749 | -0.3377 | 0.002336 |
| ribosomal RNA small subunit methyltransferase B | 0.908566 | 1.04575 | 0.202875 | 0.002344 |
| betaine aldehyde dehydrogenase...2038 | 0.929527 | 1.075107 | 0.209912 | 0.002348 |
| transcriptional regulator, Fis-type | 0.872866 | 1.070144 | 0.293973 | 0.002373 |
| dienelactone hydrolase...1877 | 1.032599 | 0.832796 | -0.31025 | 0.002381 |
| hypothetical protein ABUW_3461 | 0.953775 | 1.089079 | 0.191388 | 0.002384 |
| hypothetical protein ABUW_0364 | 0.919551 | 1.097189 | 0.254812 | 0.0024 |
| acyl-CoA dehydrogenase...1728 | 0.889293 | 0.958592 | 0.108258 | 0.002411 |
| membrane spanning export protein | 1.015875 | 0.901667 | -0.17206 | 0.002415 |

| | | | | |
|---|---|---|---|---|
| hypothetical protein ABUW_0128 | 0.879757 | 1.193171 | 0.439623 | 0.002459 |
| major facilitator superfamily MFS_1...228 | 0.884508 | 1.081655 | 0.290294 | 0.00252 |
| ferric uptake regulation protein | 0.928861 | 1.148195 | 0.305832 | 0.002593 |
| phosphoribosylanthranilate isomerase | 0.965052 | 1.10055 | 0.189546 | 0.002597 |
| glutathione S-transferase...1608 | 0.942709 | 1.05824 | 0.166783 | 0.002604 |
| transcriptional regulator, IclR family...705 | 0.884929 | 1.095431 | 0.307865 | 0.002626 |
| 3-oxoadipate CoA-transferase subunit A...1634 | 0.921161 | 1.153079 | 0.323966 | 0.002626 |
| putative acetyltransferase | 1.087737 | 0.850911 | -0.35425 | 0.002654 |
| acyl-CoA dehydrogenase...476 | 0.754832 | 1.319285 | 0.805529 | 0.002687 |
| 3-deoxy-7-phosphoheptulonate synthase...2026 | 0.944988 | 1.097841 | 0.216301 | 0.002757 |
| phosphoribosylaminoimidazolecarboxamide formyltransferas | 0.894247 | 1.157277 | 0.371988 | 0.002759 |
| chaperone protein DnaK | 1.051915 | 0.961758 | -0.12927 | 0.002843 |
| ABC transport system permease protein | 1.082446 | 0.820799 | -0.39919 | 0.00287 |
| translation initiation factor IF-3 | 1.019469 | 0.969397 | -0.07266 | 0.002877 |
| D-3-phosphoglycerate dehydrogenase | 0.915933 | 1.044524 | 0.189532 | 0.002936 |
| HAD-superfamily subfamily IB hydrolase | 0.953289 | 1.143664 | 0.262678 | 0.002962 |
| succinyl-CoA synthase, beta subunit | 0.946556 | 1.129115 | 0.254434 | 0.002976 |
| hypothetical protein ABUW_2663 | 0.766814 | 1.299902 | 0.761453 | 0.00299 |
| leucine-responsive transcriptional regulator, AsnC family | 1.160977 | 1.08581 | -0.09657 | 0.002993 |
| putative benzoate transport porin | 1.006686 | 1.100884 | 0.129049 | 0.003038 |
| outer membrane lipoprotein | 1.058227 | 0.89942 | -0.23458 | 0.003068 |
| CsuD | 0.870939 | 1.248143 | 0.519139 | 0.003092 |
| hypothetical protein ABUW_1367 | 1.09868 | 0.8816 | -0.31757 | 0.003114 |
| exonuclease V alpha subunit | 1.083009 | 0.906548 | -0.25659 | 0.003162 |
| hypothetical protein ABUW_1366 | 0.9573 | 1.061021 | 0.14841 | 0.003175 |
| hypothetical protein ABUW_1723 | 1.076927 | 0.889667 | -0.27558 | 0.003216 |
| peptidyl-prolyl cis-trans isomerase Mip | 0.971343 | 1.060902 | 0.127239 | 0.003218 |
| NADP-specific glutamate dehydrogenase | 0.855478 | 1.142098 | 0.416885 | 0.00322 |
| methionine aminopeptidase, type I...1556 | 0.926828 | 1.086247 | 0.228978 | 0.003235 |
| arabinose 5-phosphate isomerase | 0.976147 | 1.044174 | 0.097192 | 0.003274 |
| hypothetical protein ABUW_2327 | 0.942137 | 1.093062 | 0.214366 | 0.003283 |
| D-serine/D-alanine/glycine transporter...485 | 0.912298 | 1.155041 | 0.340367 | 0.003321 |
| hypothetical protein ABUW_2065 | 0.942296 | 1.102869 | 0.22701 | 0.003341 |
| [protein-PII] uridylyltransferase | 0.94344 | 1.004559 | 0.09056 | 0.003371 |
| hypothetical protein ABUW_0708 | 1.114401 | 0.789105 | -0.49798 | 0.003383 |
| hypothetical protein ABUW_2293 | 1.097305 | 0.833507 | -0.3967 | 0.003405 |
| glutamyl-tRNA(Gln) amidotransferase, A subunit | 0.919838 | 1.124738 | 0.290138 | 0.003464 |
| hypothetical protein ABUW_0046 | 0.889362 | 1.122041 | 0.335283 | 0.003476 |
| hypothetical protein ABUW_3793 | 1.084959 | 0.94575 | -0.19811 | 0.003484 |
| glycosyltransferase...577 | 0.898597 | 1.305467 | 0.538819 | 0.003535 |
| lysyl-tRNA synthetase...2233 | 0.932557 | 1.107445 | 0.247971 | 0.003535 |
| tRNA-i(6)A37 thiotransferase | 1.08176 | 0.931515 | -0.21573 | 0.003542 |
| phosphoserine phosphatase...1617 | 0.932826 | 1.071265 | 0.199635 | 0.00355 |
| hypothetical protein ABUW_3024 | 1.023117 | 0.855513 | -0.25811 | 0.00356 |
| nucleotide sugar epimerase/dehydratase | 1.011019 | 0.909951 | -0.15195 | 0.003571 |
| hypothetical protein ABUW_1066 | 1.005525 | 0.735977 | -0.45022 | 0.003574 |
| trigger factor | 1.095802 | 0.929355 | -0.23769 | 0.003615 |

| | | | | |
|---|---|---|---|---|
| peroxidase | 1.087724 | 0.905751 | -0.26413 | 0.003634 |
| lipid A biosynthesis acyltransferase...870 | 0.860976 | 1.083857 | 0.332129 | 0.003645 |
| cation efflux system protein (EsvF1) | 0.928541 | 1.171348 | 0.335132 | 0.003671 |
| putative RNA methylase | 0.924187 | 1.07497 | 0.218039 | 0.003704 |
| ribosomal large subunit pseudouridine synthase B | 0.944423 | 1.177131 | 0.317769 | 0.003716 |
| transcriptional regulator, TetR family...55 | 0.894907 | 1.062994 | 0.248324 | 0.003757 |
| CsuB | 0.916101 | 1.058505 | 0.208451 | 0.003856 |
| type IV pilin structural subunit | 1.23268 | 0.766553 | -0.68534 | 0.003935 |
| transcriptional regulator NrdR | 1.041362 | 0.963773 | -0.11171 | 0.003952 |
| carboxy- protease | 1.020377 | 0.918969 | -0.15101 | 0.00398 |
| isocitrate dehydrogenase, NADP-dependent...2445 | 0.999556 | 1.08986 | 0.124783 | 0.004012 |
| hypothetical protein ABUW_2180 | 0.884281 | 1.157069 | 0.387898 | 0.004037 |
| imidazole glycerol phosphate synthase, glutamine amidotrans | 0.943585 | 1.143645 | 0.277415 | 0.004037 |
| argininosuccinate lyase | 0.954143 | 1.12736 | 0.240672 | 0.004096 |
| chaperone protein | 0.921488 | 1.124598 | 0.287372 | 0.004116 |
| MFS permease | 0.958007 | 1.215959 | 0.343986 | 0.004135 |
| nucleoprotein/polynucleotide-associated enzyme | 0.86729 | 1.170438 | 0.432461 | 0.004216 |
| transcriptional regulator, TetR family...846 | 0.94525 | 0.998142 | 0.078549 | 0.004263 |
| acyl-CoA dehydrogenase...1936 | 1.02774 | 0.968979 | -0.08494 | 0.004288 |
| transcription elongation factor NusA | 1.056464 | 0.90333 | -0.22592 | 0.004301 |
| tryptophan synthase, beta subunit...1820 | 1.150961 | 0.917712 | -0.32673 | 0.004344 |
| hypothetical protein ABUW_0236 | 0.893496 | 1.083855 | 0.278639 | 0.00435 |
| monooxygenase, flavin-binding family...392 | 0.791529 | 1.328301 | 0.746867 | 0.004357 |
| 3-hydroxyacyl-CoA dehydrogenase...242 | 0.670161 | 1.091759 | 0.704075 | 0.004365 |
| 4-aminobutyrate transaminase | 0.90784 | 1.020576 | 0.168874 | 0.004373 |
| peptide chain release factor 1 | 0.920971 | 1.104774 | 0.262524 | 0.004375 |
| CsuC | 0.929795 | 1.146085 | 0.301729 | 0.004386 |
| alanine racemase...1478 | 0.90357 | 1.112118 | 0.299602 | 0.004425 |
| hypothetical protein ABUW_0673 | 1.142569 | 0.81702 | -0.48384 | 0.00451 |
| outer membrane protein OprE3 | 0.93243 | 1.10185 | 0.240861 | 0.004608 |
| putative adenylyltransferase | 0.983784 | 1.171422 | 0.251847 | 0.004646 |
| hypothetical protein ABUW_0359 | 1.130192 | 0.859768 | -0.39455 | 0.004647 |
| cysteinyl-tRNA synthetase | 1.020471 | 0.908295 | -0.168 | 0.004655 |
| bifunctional protein wax ester synthase/acyl-CoA diacylglycer | 0.946164 | 1.04469 | 0.142913 | 0.004657 |
| peptidase S1 and S6 | 0.920935 | 1.093293 | 0.247509 | 0.004684 |
| hypothetical protein ABUW_3251 | 1.03667 | 0.92482 | -0.16471 | 0.004686 |
| metallo-beta-lactamase family protein...1065 | 0.860074 | 1.091818 | 0.344199 | 0.004741 |
| C4-dicarboxylate transport protein...1033 | 0.973388 | 1.109205 | 0.188439 | 0.004764 |
| OmpA/MotB domain protein...2491 | 1.086668 | 0.879735 | -0.30477 | 0.00484 |
| hypothetical protein ABUW_0402 | 1.059309 | 0.917029 | -0.20808 | 0.004865 |
| acyl-CoA dehydrogenase...892 | 1.170587 | 0.806647 | -0.53722 | 0.004971 |
| leucyl-tRNA synthetase | 0.969764 | 1.076713 | 0.150928 | 0.005024 |
| phenylacetate-CoA oxygenase, PaaI subunit | 0.897409 | 1.072913 | 0.257696 | 0.005068 |
| ribosome recycling factor | 1.013062 | 1.042996 | 0.042012 | 0.005069 |
| preprotein translocase, SecY subunit | 0.828511 | 0.934444 | 0.173589 | 0.005081 |
| GCN5-related N-acetyltransferase | 1.024866 | 0.921525 | -0.15334 | 0.005124 |
| hypothetical protein ABUW_1556 | 0.939225 | 1.096787 | 0.223742 | 0.005125 |

| | | | | |
|---|---|---|---|---|
| phosphate ABC transporter, substrate-binding protein | 1.061436 | 0.848523 | -0.32299 | 0.00513 |
| coenzyme PQQ biosynthesis protein C | 1.073131 | 0.788749 | -0.44419 | 0.005133 |
| outer membrane assembly lipoprotein YfgL | 1.029682 | 0.949826 | -0.11646 | 0.005185 |
| NADPH-dependent fmn reductase...369 | 0.966086 | 1.33416 | 0.465708 | 0.005218 |
| imidazoleglycerol phosphate synthase, cyclase subunit | 1.005406 | 1.071573 | 0.091953 | 0.005285 |
| hypothetical protein ABUW_3805 | 0.919729 | 1.062876 | 0.208692 | 0.005288 |
| peptidase M1, alanyl aminopeptidase | 0.961192 | 1.006371 | 0.066266 | 0.005306 |
| copper-translocating P-type ATPase...1335 | 1.036001 | 0.910647 | -0.18606 | 0.005321 |
| pyridine nucleotide transhydrogenase | 0.943089 | 1.154595 | 0.291922 | 0.005375 |
| NADH dehydrogenase I chain N | 0.879548 | 1.051558 | 0.257696 | 0.00539 |
| pseudouridine synthase, Rsu | 0.922563 | 1.062044 | 0.203123 | 0.005395 |
| aspartate aminotransferase | 0.948551 | 1.055334 | 0.153902 | 0.005461 |
| apolipoprotein N-acyltransferase | 0.943097 | 1.089576 | 0.208289 | 0.005462 |
| phospholipid/glycerol acyltransferase...1329 | 0.972935 | 1.057121 | 0.119726 | 0.005516 |
| efflux transporter, RND family, MFP subunit | 1.1177 | 0.98529 | -0.18191 | 0.005535 |
| surface antigen | 0.891602 | 1.129812 | 0.341611 | 0.005581 |
| NADH dehydrogenase I chain B | 0.9184 | 1.060626 | 0.207723 | 0.005643 |
| hypothetical protein ABUW_1775 | 0.860359 | 1.153621 | 0.423159 | 0.005695 |
| integral membrane protein DUF6 | 1.122916 | 0.86463 | -0.3771 | 0.005731 |
| nonribosomal peptide synthetase BasD | 1.070435 | 0.861656 | -0.31301 | 0.005736 |
| ATP phosphoribosyltransferase | 1.026074 | 0.948071 | -0.11407 | 0.005791 |
| 3-deoxy-manno-octulosonate cyidylyltransferase | 0.966792 | 1.089444 | 0.172314 | 0.005792 |
| integral membrane protein TerC | 1.138597 | 0.838693 | -0.44104 | 0.005901 |
| multidrug efflux protein...1589 | 1.02708 | 0.932035 | -0.14009 | 0.005937 |
| band 7 protein | 0.947679 | 1.088707 | 0.200145 | 0.005962 |
| methenyltetrahydrofolate cyclohydrolase | 0.949914 | 1.057352 | 0.154587 | 0.005966 |
| histidine triad protein...1597 | 1.129093 | 0.873909 | -0.36961 | 0.005986 |
| cytochrome b562 | 0.752825 | 1.121366 | 0.57487 | 0.006008 |
| transcriptional regulator OhrR | 0.972512 | 1.067122 | 0.133937 | 0.006039 |
| riboflavin synthase, alpha subunit | 0.878721 | 1.210417 | 0.462027 | 0.00604 |
| enoyl-CoA hydratase/isomerase...168 | 0.968489 | 1.137213 | 0.231695 | 0.006053 |
| thioesterase superfamily protein...1195 | 1.297893 | 0.839892 | -0.62789 | 0.00607 |
| hypothetical protein ABUW_2125 | 0.793135 | 1.043605 | 0.395936 | 0.006154 |
| site-specific recombinase | 0.926646 | 1.167611 | 0.33347 | 0.006157 |
| peptidase S16, lon domain protein | 0.838223 | 1.175402 | 0.487748 | 0.006183 |
| hypothetical protein ABUW_3796 | 1.044508 | 0.881174 | -0.24532 | 0.0062 |
| D-alanine--D-alanine ligase B | 1.001644 | 1.061725 | 0.084042 | 0.006222 |
| molybdopterin biosynthesis protein...772 | 0.918452 | 1.044719 | 0.185838 | 0.006228 |
| NADPH-dependent fmn reductase...1212 | 0.896084 | 1.021881 | 0.189522 | 0.006231 |
| aspartate-semialdehyde dehydrogenase | 1.04359 | 0.935728 | -0.15739 | 0.00635 |
| chloramphenicol resistance pump cmr | 1.065341 | 0.914755 | -0.21986 | 0.006367 |
| pirin family protein | 1.097847 | 0.888349 | -0.30548 | 0.006369 |
| seryl-tRNA synthetase | 0.98402 | 1.044406 | 0.085923 | 0.006375 |
| phosphatidylserine decarboxylase | 0.923089 | 1.02015 | 0.14424 | 0.006411 |
| UDP-glucose 4-epimerase...2000 | 1.021193 | 0.977119 | -0.06365 | 0.006415 |
| acetyl-CoA carboxylase, biotin carboxylase | 1.091196 | 0.993249 | -0.13568 | 0.006424 |
| UDP-N-acetylmuramoylalanine--D-glutamate ligase | 0.925185 | 1.091412 | 0.238382 | 0.006459 |

| | | | | |
|---|---|---|---|---|
| triose-phosphate isomerase | 1.018117 | 0.950985 | -0.09841 | 0.006466 |
| 17 kDa surface antigen | 1.088703 | 0.931299 | -0.22529 | 0.006504 |
| dienelactone hydrolase...2270 | 0.9813 | 1.064253 | 0.117076 | 0.006552 |
| hypothetical protein ABUW_0589 | 0.98074 | 1.082848 | 0.142888 | 0.006561 |
| iron-containing alcohol dehydrogenase | 1.003541 | 0.926522 | -0.1152 | 0.006611 |
| Glycerate kinase | 0.93691 | 1.016082 | 0.117034 | 0.006646 |
| phosphatidate cytidylyltransferase | 0.876822 | 1.146836 | 0.387303 | 0.006662 |
| alcohol dehydrogenase, zinc-binding...1858 | 0.938088 | 1.156636 | 0.302139 | 0.006682 |
| cysteine synthase B | 0.976592 | 1.04023 | 0.091075 | 0.0067 |
| nitroreductase...1095 | 1.09671 | 0.943425 | -0.2172 | 0.006711 |
| L-lactate dehydrogenase (cytochrome) | 0.96516 | 1.036916 | 0.103458 | 0.006747 |
| DNA repair protein RecO | 0.934059 | 1.06369 | 0.187492 | 0.006748 |
| pyrroline-5-carboxylate reductase | 0.906921 | 1.117171 | 0.300802 | 0.006773 |
| hypothetical protein ABUW_3064 | 1.063129 | 0.903192 | -0.23521 | 0.006879 |
| guanine deaminase...1601 | 0.926895 | 1.08284 | 0.224342 | 0.006996 |
| dihydroorotase | 0.841807 | 1.16302 | 0.466316 | 0.007006 |
| fumarylacetoacetate hydrolase...1574 | 0.939873 | 1.154058 | 0.296179 | 0.007006 |
| FAD-dependent pyridine nucleotide- disulfide oxidoreductase | 0.917815 | 1.047108 | 0.190135 | 0.007115 |
| lipid A biosynthesis acyltransferase...692 | 0.938977 | 1.065886 | 0.182891 | 0.007118 |
| DNA topoisomerase IV, A subunit | 1.039738 | 0.956685 | -0.1201 | 0.007147 |
| hypothetical protein ABUW_2921 | 1.990196 | 0.584401 | -1.76788 | 0.007149 |
| phosphoglycerate kinase | 0.955803 | 1.05804 | 0.146609 | 0.00717 |
| CTP synthase | 0.952097 | 1.028141 | 0.110858 | 0.007192 |
| hypothetical protein ABUW_3718 | 0.86432 | 1.203184 | 0.477219 | 0.007224 |
| PTS system fructose-specific EIIBC component | 1.109621 | 0.814288 | -0.44645 | 0.007271 |
| dihydroxy-acid dehydratase...1555 | 1.156939 | 0.875369 | -0.40235 | 0.007273 |
| YCII-related protein...1512 | 0.906544 | 1.113424 | 0.296555 | 0.007289 |
| hypothetical protein ABUW_1909 | 0.857423 | 1.124177 | 0.39079 | 0.007293 |
| hypothetical protein ABUW_1429 | 1.025828 | 1.305728 | 0.348066 | 0.007363 |
| hypothetical protein ABUW_1727 | 0.998912 | 1.1073 | 0.148617 | 0.007386 |
| multiphosphoryl transfer protein | 1.051715 | 0.947588 | -0.15041 | 0.00747 |
| ggdef domain/eal domain protein | 0.946036 | 1.119128 | 0.242408 | 0.007488 |
| RND type efflux pump | 1.089808 | 0.855941 | -0.34849 | 0.007497 |
| RDD family protein | 0.949147 | 1.055202 | 0.152815 | 0.007653 |
| FeS cluster assembly scaffold IscU | 1.151406 | 0.86228 | -0.41717 | 0.007667 |
| 3-oxoadipate enol-lactonase...842 | 0.986153 | 1.132139 | 0.199169 | 0.007735 |
| ferrous iron transport protein B | 0.880708 | 1.187929 | 0.431713 | 0.007795 |
| ribosomal protein L35 | 1.146016 | 0.890292 | -0.36428 | 0.007851 |
| thiamine biosynthesis protein ThiC | 1.537656 | 0.598531 | -1.36124 | 0.007852 |
| hypothetical protein ABUW_1063 | 0.960644 | 0.78338 | -0.29429 | 0.007921 |
| peptidoglycan glycosyltransferase | 0.95768 | 1.016697 | 0.086275 | 0.007936 |
| hypothetical protein ABUW_0113 | 0.852168 | 1.164296 | 0.450248 | 0.007941 |
| hypothetical protein ABUW_3157 | 0.627074 | 1.169399 | 0.89906 | 0.007953 |
| glutamate-1-semialdehyde-2,1-aminomutase | 0.982233 | 1.077423 | 0.133447 | 0.008101 |
| UDP-N-acetylenolpyruvoylglucosamine reductase | 0.980919 | 1.061891 | 0.11443 | 0.008102 |
| glutamate racemase | 0.858934 | 1.05181 | 0.292256 | 0.008179 |
| acyl-(acyl-carrier-protein)--UDP-N-acetylglucosamine O-acyltr | 0.989778 | 1.081554 | 0.127929 | 0.008179 |

| | | | | |
|---|---|---|---|---|
| dihydrofolate reductase (plasmid) | 0.903813 | 0.984169 | 0.122882 | 0.008193 |
| hypothetical protein ABUW_0111 | 1.026395 | 0.961107 | -0.09482 | 0.008317 |
| 3-deoxy-D-manno-octulosonate 8-phosphate phosphatase | 0.967142 | 1.093017 | 0.176516 | 0.008354 |
| transcriptional regulator, LysR family...342 | 1.010494 | 0.905004 | -0.15906 | 0.008392 |
| hypothetical protein ABUW_0896 | 1.051461 | 0.91024 | -0.20808 | 0.008414 |
| multidrug efflux protein...701 | 1.117471 | 0.917091 | -0.2851 | 0.008498 |
| putative class A beta-lactamase | 1.085756 | 0.88777 | -0.29044 | 0.008529 |
| putative ATP-binding protein | 0.953825 | 1.083376 | 0.183738 | 0.008535 |
| hypothetical protein ABUW_2820 | 0.940576 | 1.114435 | 0.244696 | 0.008591 |
| ABC transporter ATP-binding protein...1689 | 0.980441 | 0.936996 | -0.06539 | 0.008618 |
| poly(A) polymerase | 0.873884 | 1.090457 | 0.319419 | 0.008632 |
| transcriptional regulator, AraC family...2200 | 1.135186 | 1.010979 | -0.16718 | 0.008651 |
| fumarate hydratase | 0.923245 | 1.103737 | 0.257611 | 0.008686 |
| glutathione S-transferase family protein...2177 | 0.999639 | 0.893806 | -0.16145 | 0.008714 |
| 3-oxoacyl-(acyl-carrier-protein) reductase | 1.077636 | 0.966858 | -0.15649 | 0.008715 |
| Cu(I)-responsive transcriptional regulator | 1.142888 | 0.872394 | -0.38963 | 0.008728 |
| exodeoxyribonuclease X, putative | 0.923427 | 1.096955 | 0.248435 | 0.008798 |
| magnesium and cobalt efflux protein CorC | 0.929724 | 1.10791 | 0.252967 | 0.008808 |
| hypothetical protein ABUW_1543 | 1.034285 | 0.962047 | -0.10446 | 0.008861 |
| ribosomal RNA large subunit methyltransferase J | 1.052319 | 0.963738 | -0.12686 | 0.008872 |
| acyl-CoA dehydrogenase...2323 | 0.98918 | 0.801703 | -0.30316 | 0.008929 |
| lipoprotein, putative...1648 | 0.891105 | 1.13468 | 0.348619 | 0.008939 |
| hypothetical protein ABUW_0646 | 1.075896 | 0.741111 | -0.53778 | 0.008954 |
| hypothetical protein ABUW_3866 | 0.885617 | 1.046028 | 0.240167 | 0.008973 |
| RHS family protein | 0.930686 | 1.049365 | 0.17315 | 0.009021 |
| 3-hydroxyacyl-CoA dehydrogenase...1818 | 1.11868 | 0.937392 | -0.25507 | 0.009086 |
| protein-(glutamine-N5) methyltransferase, ribosomal protein | 1.010494 | 0.940848 | -0.10303 | 0.009107 |
| Zn-dependent oligopeptidase | 1.046168 | 0.898635 | -0.21931 | 0.009214 |
| cell division protein FtsA | 0.911436 | 1.048753 | 0.202461 | 0.009215 |
| hypothetical protein ABUW_2570 | 0.904208 | 1.047219 | 0.211836 | 0.009224 |
| hypothetical protein ABUW_3322 | 1.209631 | 0.884769 | -0.4512 | 0.009305 |
| glycosyltransferase...487 | 1.011758 | 0.936152 | -0.11205 | 0.009408 |
| copper amine oxidase | 1.168496 | 0.797929 | -0.55032 | 0.009426 |
| Transcriptional repressor protein (plasmid) | 1.057223 | 0.916868 | -0.20549 | 0.009442 |
| threonine dehydratase...2054 | 0.923132 | 1.145966 | 0.311955 | 0.009506 |
| GTP-binding protein LepA | 0.961305 | 1.035527 | 0.1073 | 0.00954 |
| 3-methyl-2-oxobutanoate hydroxymethyltransferase | 0.946672 | 1.034096 | 0.127435 | 0.009555 |
| hypothetical protein ABUW_0048 | 1.074237 | 1.026275 | -0.0659 | 0.009565 |
| muconate cycloisomerase | 0.839061 | 1.144935 | 0.448419 | 0.009649 |
| hypothetical protein ABUW_0664 | 1.102312 | 0.790053 | -0.48051 | 0.009709 |
| UDP-3-O-(3-hydroxymyristoyl) glucosamine N-acyltransferase | 0.918431 | 1.088416 | 0.244986 | 0.009763 |
| 2-C-methyl-D-erythritol 4-phosphate cytidylyltransferase | 0.99686 | 1.08126 | 0.117251 | 0.009766 |
| L-sorbosone dehydrogenase...1614 | 0.983957 | 0.910774 | -0.1115 | 0.009827 |
| isochorismatase hydrolase...401 | 0.849974 | 1.06715 | 0.328272 | 0.009845 |
| dihydrodipicolinate reductase | 0.979045 | 1.076195 | 0.136492 | 0.009853 |
| hypothetical protein ABUW_1629 | 1.192854 | 0.783674 | -0.60609 | 0.009876 |
| D-amino acid dehydrogenase small subunit | 1.152646 | 1.052584 | -0.13101 | 0.009923 |

| | | | | |
|---|---|---|---|---|
| NADPH:quinone oxidoreductase | 0.976034 | 1.039252 | 0.090541 | 0.009923 |
| malate dehydrogenase...2408 | 0.940396 | 1.124828 | 0.258364 | 0.009974 |
| cytidylate kinase | 1.071963 | 0.910336 | -0.23578 | 0.010023 |
| hypothetical protein ABUW_2610 | 0.901383 | 1.207549 | 0.42187 | 0.01011 |
| CsuA | 0.912856 | 1.195354 | 0.388979 | 0.010126 |
| hypothetical protein ABUW_0710 | 1.055763 | 0.902122 | -0.22689 | 0.010152 |
| Smr protein/MutS2 | 0.934456 | 1.013016 | 0.116459 | 0.010161 |
| NADH pyrophosphatase | 0.978452 | 1.050006 | 0.101825 | 0.010167 |
| glutamate-ammonia-ligase adenylyltransferase | 0.983132 | 1.060806 | 0.109704 | 0.010245 |
| gamma-glutamyltransferase | 1.123894 | 0.796605 | -0.49657 | 0.010288 |
| metalloendopeptidase | 1.022277 | 1.001313 | -0.02989 | 0.01034 |
| Holliday junction DNA helicase RuvB | 1.269212 | 0.822589 | -0.62569 | 0.010381 |
| general secretion pathway protein G | 0.918176 | 1.070103 | 0.220907 | 0.010389 |
| TetR family transcriptional regulator | 1.086367 | 0.976347 | -0.15405 | 0.01048 |
| NAD dependent epimerase/dehydratase family...909 | 0.908244 | 1.070963 | 0.237757 | 0.010525 |
| rod shape-determining protein MreC | 1.083385 | 0.946465 | -0.19493 | 0.010647 |
| phosphoserine phosphatase/homoserine phosphotransferase | 0.909337 | 1.073233 | 0.239076 | 0.010666 |
| deoxyuridine 5'-triphosphate nucleotidohydrolase | 0.95323 | 1.078203 | 0.177732 | 0.010719 |
| phosphoribosyl (-ATP, -AMP) pyrophosphohydrolase/cyclohy | 0.868371 | 1.154195 | 0.410503 | 0.010725 |
| UDP-glucose/GDP-mannose dehydrogenase | 0.957852 | 1.070183 | 0.159984 | 0.010748 |
| aldo/keto reductase...2262 | 1.072052 | 0.916803 | -0.22569 | 0.010786 |
| phosphoribosylglycinamide formyltransferase 2 | 0.997878 | 1.091149 | 0.128913 | 0.010891 |
| hypothetical protein ABUW_2605 | 0.916553 | 1.112874 | 0.28 | 0.011133 |
| hypothetical protein ABUW_3155 | 1.147605 | 0.990563 | -0.21231 | 0.011154 |
| PGAP1 family protein | 0.993177 | 0.880661 | -0.17346 | 0.011158 |
| integrase | 0.811871 | 0.97021 | 0.257046 | 0.011183 |
| hypothetical protein ABUW_0462 | 1.009334 | 1.064971 | 0.077411 | 0.011194 |
| ATP synthase F1, gamma subunit AtpG | 0.947916 | 1.086684 | 0.197102 | 0.011253 |
| phosphoribosylformylglycinamidine synthase | 0.951839 | 1.019108 | 0.098518 | 0.011283 |
| hypothetical protein ABUW_3648 | 1.019925 | 0.951118 | -0.10077 | 0.011376 |
| nudix hydrolase...687 | 0.986904 | 1.043951 | 0.081072 | 0.011377 |
| hypothetical protein ABUW_1534 | 0.995455 | 1.21397 | 0.286305 | 0.01147 |
| hypothetical protein ABUW_3582 | 1.067208 | 0.862657 | -0.30698 | 0.011475 |
| carbapenem-hydrolyzing oxacillinase OXA-69 | 1.028534 | 0.894666 | -0.20117 | 0.011558 |
| hypothetical protein ABUW_1983 | 1.180516 | 0.94966 | -0.31393 | 0.011566 |
| ATP synthase F1, alpha subunit | 0.977394 | 1.076282 | 0.139044 | 0.011591 |
| hypothetical protein ABUW_3310 | 0.882994 | 0.987982 | 0.16208 | 0.011627 |
| glyoxalase/bleomycin resistance protein/dioxygenase | 0.806635 | 1.061205 | 0.395717 | 0.011664 |
| copper-translocating P-type ATPase...2120 | 1.042367 | 0.882526 | -0.24015 | 0.01174 |
| Heavy metal transport/detoxification protein | 1.123177 | 0.899075 | -0.32107 | 0.011815 |
| urea amidolyase | 1.081274 | 0.908529 | -0.25113 | 0.011819 |
| hypothetical protein ABUW_1761 | 0.881929 | 1.113683 | 0.336605 | 0.011865 |
| nicotinamide phosphoribosyltransferase | 1.001993 | 0.891052 | -0.16929 | 0.011898 |
| hypothetical protein ABUW_3179 | 1.036298 | 0.976407 | -0.08588 | 0.011934 |
| ribosome-binding factor A | 1.045305 | 0.947994 | -0.14097 | 0.011947 |
| aldo/keto reductase...2097 | 0.960398 | 1.078643 | 0.167513 | 0.012031 |
| cupin family protein | 0.976546 | 1.112282 | 0.187763 | 0.012038 |

| | | | | |
|---|---|---|---|---|
| hypothetical protein ABUW_1120 | 1.096237 | 0.863978 | -0.34349 | 0.012164 |
| phosphogluconate dehydratase | 0.961232 | 1.039453 | 0.112868 | 0.012178 |
| hypothetical protein ABUW_1541 | 1.039871 | 0.849461 | -0.29178 | 0.012267 |
| hypothetical protein ABUW_2144 | 0.812155 | 1.307215 | 0.686669 | 0.01229 |
| ribosomal large subunit pseudouridine synthase D | 0.867869 | 1.087674 | 0.325697 | 0.012302 |
| sodium/glutamate symporter | 0.846143 | 1.215423 | 0.522486 | 0.012327 |
| general secretion pathway protein L | 1.049995 | 0.956462 | -0.1346 | 0.012355 |
| peptidoglycan-binding LysM...901 | 1.036991 | 0.765176 | -0.43854 | 0.012402 |
| 2-ketogluconate reductase | 0.94749 | 1.096321 | 0.210488 | 0.012456 |
| hypothetical protein ABUW_0452 | 0.951572 | 1.048779 | 0.140326 | 0.012527 |
| chromosome partitioning protein ParB | 0.982806 | 1.061647 | 0.111325 | 0.012531 |
| hypothetical protein ABUW_4122 (plasmid) | 1.802691 | 0.547461 | -1.71932 | 0.012532 |
| hypothetical protein ABUW_1046 | 0.948569 | 1.035715 | 0.126802 | 0.012692 |
| peptidylprolyl isomerase, fkbp-type | 1.051337 | 0.988501 | -0.08891 | 0.012702 |
| hypothetical protein ABUW_0473 | 0.996944 | 1.10329 | 0.146227 | 0.012702 |
| Fe-S protein assembly co-chaperone HscB | 1.038331 | 0.956273 | -0.11877 | 0.012742 |
| putave Na+/H+ antiporter | 0.832487 | 1.016919 | 0.288706 | 0.012743 |
| ribosomal protein S15 | 1.094606 | 0.963047 | -0.18473 | 0.012896 |
| hypothetical protein ABUW_0738 | 1.201828 | 0.753723 | -0.67312 | 0.012937 |
| translation initiation factor IF-2 | 1.024668 | 0.960822 | -0.09282 | 0.012958 |
| UspA domain protein | 1.250018 | 0.768522 | -0.70179 | 0.012997 |
| phosphomannomutase...1567 | 0.885014 | 1.137077 | 0.361557 | 0.013011 |
| hypothetical protein ABUW_0823 | 1.006464 | 0.823246 | -0.2899 | 0.013234 |
| transcriptional regulator, GntR family...1331 | 0.925367 | 1.102713 | 0.25296 | 0.013408 |
| quinoprotein glucose dehydrogenase-B...2440 | 1.06665 | 0.867275 | -0.29852 | 0.013445 |
| putative lysozyme | 1.162399 | 0.932137 | -0.31849 | 0.013466 |
| preprotein translocase iisp family auxillary membrane compo | 0.989236 | 1.153946 | 0.222188 | 0.013528 |
| porphobilinogen deaminase | 1.022203 | 0.999262 | -0.03275 | 0.013703 |
| malate dehydrogenase...1909 | 0.933326 | 1.118996 | 0.261751 | 0.013763 |
| hypothetical protein ABUW_0820 | 0.956726 | 1.264258 | 0.402113 | 0.013775 |
| hypothetical protein ABUW_2364 | 1.008078 | 0.933333 | -0.11114 | 0.013795 |
| invasion protein expression up-regulator | 0.793335 | 1.0018 | 0.336593 | 0.013897 |
| putrescine importer | 1.089506 | 0.863076 | -0.33611 | 0.013907 |
| alginate biosynthesis regulatory protein | 1.077435 | 0.826398 | -0.38269 | 0.013942 |
| peptidase | 0.987955 | 0.887003 | -0.15551 | 0.014029 |
| cysteine desulfurase IscS | 1.200594 | 0.862608 | -0.47697 | 0.014064 |
| hypothetical protein ABUW_2735 | 1.062795 | 0.920505 | -0.20737 | 0.014089 |
| hypothetical protein ABUW_1060 | 1.055092 | 0.916247 | -0.20356 | 0.014137 |
| hypothetical protein ABUW_4077 (plasmid) | 1.104144 | 0.934959 | -0.23995 | 0.014181 |
| hypothetical protein ABUW_1642 | 0.472962 | 2.135451 | 2.174745 | 0.01437 |
| chaperonin GroL | 1.109916 | 0.9412 | -0.23788 | 0.014374 |
| 2-methylisocitrate dehydratase, Fe/S-dependent | 0.887862 | 1.103098 | 0.313154 | 0.014409 |
| hypothetical protein ABUW_1256 | 1.208003 | 0.772994 | -0.6441 | 0.014422 |
| transcription-repair coupling factor | 1.032945 | 0.969308 | -0.09174 | 0.01444 |
| hypothetical protein ABUW_1210 | 1.068475 | 0.837222 | -0.35187 | 0.014518 |
| hypothetical protein ABUW_2434 | 0.954226 | 1.03202 | 0.113067 | 0.014737 |
| GTP cyclohydrolase II | 0.994005 | 1.041753 | 0.067688 | 0.014745 |

| | | | | |
|---|---|---|---|---|
| hypothetical protein ABUW_2616 | 0.931077 | 1.043202 | 0.164045 | 0.014771 |
| hypothetical protein ABUW_2349 | 0.871427 | 0.973387 | 0.159634 | 0.014792 |
| high affinity gluconate permease | 0.938148 | 1.098103 | 0.227125 | 0.014809 |
| OmpA/MotB...2467 | 1.085329 | 0.787509 | -0.46276 | 0.014839 |
| dihydropteroate synthase | 0.930709 | 1.091717 | 0.230198 | 0.014858 |
| dihydroneopterin aldolase | 0.953072 | 1.086489 | 0.189017 | 0.015003 |
| Mating pair stabilization TraN precursor (plasmid) | 1.169797 | 0.773101 | -0.59753 | 0.01517 |
| chaperone protein DnaJ | 0.938542 | 1.052016 | 0.164663 | 0.015225 |
| outer membrane lipoprotein Blc | 1.090837 | 0.952769 | -0.19524 | 0.015261 |
| glutathione import ATP-binding protein GsiA | 1.088615 | 0.870535 | -0.32252 | 0.015318 |
| hypothetical protein ABUW_0666 | 0.951518 | 1.105358 | 0.216211 | 0.015356 |
| Bacterial transferase hexapeptide (three repeats) family prote | 1.192861 | 0.732924 | -0.70269 | 0.01539 |
| chaperone protein HtpG | 1.154684 | 0.895157 | -0.36728 | 0.015435 |
| ribosomal protein L29 | 1.055713 | 0.994437 | -0.08626 | 0.015453 |
| transcriptional regulator, AraC family...468 | 0.965189 | 1.158372 | 0.263215 | 0.015469 |
| transcriptional regulator, TetR family...1744 | 0.980854 | 1.069683 | 0.125073 | 0.015563 |
| hypothetical protein ABUW_2061 | 0.765213 | 1.065729 | 0.477907 | 0.015624 |
| 3-deoxy-D-manno-2-octulosonate transferase | 1.018125 | 0.983114 | -0.05048 | 0.015641 |
| 2-octaprenylphenol hydroxylase of ubiquinone biosynthetic p | 0.978221 | 1.016436 | 0.055287 | 0.01565 |
| hypothetical protein ABUW_5007 (plasmid) | 1.143786 | 0.949323 | -0.26885 | 0.01572 |
| transcriptional factor | 1.048674 | 0.887301 | -0.24107 | 0.015834 |
| phosphate starvation-inducible protein | 0.933639 | 1.047467 | 0.165969 | 0.015896 |
| hypothetical protein ABUW_4120 (plasmid) | 0.915555 | 1.110674 | 0.278718 | 0.015913 |
| phosphopantothenoylcysteine decarboxylase/phosphopantot | 0.826164 | 1.094324 | 0.405539 | 0.015918 |
| ribosomal-protein-alanine acetyltransferase | 0.911792 | 1.028986 | 0.174446 | 0.015942 |
| UDP-N-acetylglucosamine diphosphorylase/glucosamine-1- pl | 0.882478 | 1.114329 | 0.336543 | 0.015955 |
| Zeta toxin family protein (plasmid) | 1.053423 | 1.007192 | -0.06475 | 0.016037 |
| N-acetylmuramoyl-L-alanine amidase, family 2 | 1.01033 | 1.118447 | 0.146671 | 0.016381 |
| branched-chain amino acid aminotransferase | 0.956367 | 1.054864 | 0.141422 | 0.01639 |
| transcriptional regulator, LysR family...1804 | 0.982157 | 0.926628 | -0.08396 | 0.016391 |
| alpha/beta hydrolase fold protein...554 | 0.926029 | 1.151268 | 0.314094 | 0.016426 |
| 3,4-dihydroxy-2-butanone 4-phosphate synthase | 0.992387 | 1.029289 | 0.052674 | 0.016536 |
| hypothetical protein ABUW_1918 | 0.770213 | 1.160687 | 0.591651 | 0.01662 |
| high-affinity choline transport protein | 0.90847 | 1.109023 | 0.287778 | 0.016626 |
| putative outer membrane protein W | 0.895706 | 1.076539 | 0.265304 | 0.016651 |
| Spore coat polysaccharide biosynthesis protein spsC | 1.014099 | 0.900994 | -0.17061 | 0.016653 |
| NADH:flavin oxidoreductase/nadh oxidase | 1.127243 | 0.941281 | -0.2601 | 0.016841 |
| hypothetical protein ABUW_1633 | 1.009785 | 0.914152 | -0.14354 | 0.016919 |
| glycerol-3-phosphate dehydrogenase (NAD(P)+) | 0.97471 | 1.013744 | 0.056649 | 0.01695 |
| hypothetical protein ABUW_3265 | 1.125486 | 0.924505 | -0.2838 | 0.016954 |
| copper resistance D | 0.69177 | 0.545849 | -0.34179 | 0.016968 |
| hypothetical protein ABUW_0749 | 1.077544 | 0.904677 | -0.25227 | 0.017204 |
| hypothetical protein ABUW_1587 | 1.060625 | 1.185546 | 0.160637 | 0.017229 |
| inorganic diphosphatase | 0.97373 | 1.083312 | 0.153855 | 0.017269 |
| NADH dehydrogenase I chain J | 0.926112 | 1.082878 | 0.225612 | 0.01733 |
| transcriptional regulator, TetR family...817 | 0.992069 | 1.067522 | 0.105753 | 0.01743 |
| 6-O-methylguanine-DNA methyltransferase | 1.277543 | 0.754357 | -0.76005 | 0.017615 |

| | | | | |
|---|---|---|---|---|
| lipoprotein, putative...1053 | 1.227401 | 0.782732 | -0.64902 | 0.017732 |
| TonB-dependent receptor protein | 0.952743 | 1.088597 | 0.19231 | 0.017837 |
| hypothetical protein ABUW_0792 | 1.007158 | 0.6608 | -0.608 | 0.017863 |
| uroporphyrin-III C-methyltransferase | 0.96303 | 1.055415 | 0.132158 | 0.017964 |
| 2-octaprenyl-6-methoxyphynol hydroxylase, FAD/NAD UbiH | 0.912624 | 1.038123 | 0.185886 | 0.018012 |
| phosphoribosylformylglycinamidine cyclo-ligase | 0.922183 | 1.101866 | 0.256823 | 0.018036 |
| hypothetical protein ABUW_2317 | 1.043415 | 0.927943 | -0.16921 | 0.018115 |
| glutathione-dependent formaldehyde dehydrogenase | 1.150197 | 0.860224 | -0.4191 | 0.018135 |
| toluene tolerance efflux transporter...2098 | 0.970545 | 1.017997 | 0.068866 | 0.018141 |
| hypothetical protein ABUW_2199 | 0.864857 | 1.153388 | 0.415345 | 0.018179 |
| hypothetical protein ABUW_2834 | 1.070433 | 0.852622 | -0.32822 | 0.018195 |
| multidrug efflux protein AdeK | 1.010304 | 0.945961 | -0.09494 | 0.018205 |
| D-amino acid dehydrogenase 3 small subunit | 0.868933 | 1.110025 | 0.353274 | 0.018352 |
| dimethyladenosine transferase | 1.067655 | 0.971818 | -0.13569 | 0.018394 |
| 1-phosphofructokinase | 1.031842 | 0.865881 | -0.25298 | 0.018486 |
| uroporphyrinogen-III synthase | 0.945053 | 1.104932 | 0.22549 | 0.018551 |
| glucose dehydrogenase | 0.985426 | 0.857952 | -0.19985 | 0.018634 |
| ferrichrome-iron receptor | 0.902536 | 1.085627 | 0.266472 | 0.018732 |
| UDP-N-acetylglucosamine 1-carboxyvinyltransferase | 1.024718 | 0.93946 | -0.12532 | 0.018732 |
| ribonucleoside-diphosphate reductase alpha subunit | 1.059536 | 0.932781 | -0.18382 | 0.01878 |
| sporulation initiation inhibitor protein soj | 1.053586 | 0.971858 | -0.11649 | 0.018835 |
| hypothetical protein ABUW_3079 | 0.974936 | 0.837354 | -0.21947 | 0.018959 |
| short-chain enoyl-CoA hydratase | 1.014545 | 0.933131 | -0.12068 | 0.019026 |
| hypothetical protein ABUW_2859 | 1.072566 | 0.945963 | -0.18121 | 0.019148 |
| ribosomal L25 | 1.058089 | 0.941553 | -0.16835 | 0.019531 |
| ATP-dependent Clp protease ATP-binding subunit | 1.066336 | 0.93422 | -0.19083 | 0.019638 |
| tRNA-dihydrouridine synthase C | 1.183536 | 0.818164 | -0.53264 | 0.019675 |
| uracil phosphoribosyltransferase | 0.916312 | 0.996724 | 0.121355 | 0.019756 |
| hypothetical protein ABUW_3656 | 0.934609 | 1.007136 | 0.107823 | 0.019988 |
| hypothetical protein ABUW_0389 | 0.987963 | 1.034545 | 0.066467 | 0.020043 |
| transcriptional regulator, LysR family...1961 | 0.948182 | 1.085777 | 0.195492 | 0.020256 |
| O-succinylhomoserine sulfhydrylase | 0.878694 | 1.087148 | 0.307115 | 0.020276 |
| two-component system sensory histidine kinase | 0.9723 | 1.057264 | 0.120861 | 0.020464 |
| YecA family protein | 0.923682 | 1.190402 | 0.36598 | 0.020468 |
| amino-acid permease | 0.896495 | 1.181562 | 0.398328 | 0.020506 |
| polyphosphate kinase...2069 | 0.979615 | 1.095937 | 0.161877 | 0.020634 |
| diaminopimelate decarboxylase | 0.972457 | 1.084043 | 0.156716 | 0.020814 |
| diaminopimelate epimerase | 1.079174 | 0.987689 | -0.1278 | 0.020832 |
| hypothetical protein ABUW_2145 | 1.079382 | 0.825947 | -0.38609 | 0.02087 |
| succinate dehydrogenase, hydrophobic membrane anchor pro | 0.780919 | 1.189689 | 0.60734 | 0.020878 |
| acetylglutamate kinase | 0.963726 | 1.03676 | 0.105386 | 0.021042 |
| DEAD/DEAH box helicase...854 | 0.733783 | 0.885952 | 0.271875 | 0.021117 |
| hypothetical protein ABUW_2932 | 0.936191 | 1.045786 | 0.159713 | 0.021153 |
| phosphomannomutase...2141 | 0.965412 | 0.995323 | 0.04402 | 0.021287 |
| adenosylhomocysteinase | 1.105074 | 0.897933 | -0.29946 | 0.021289 |
| endonuclease/exonuclease/phosphatase...1155 | 0.872165 | 0.960552 | 0.139262 | 0.0213 |
| hypothetical protein ABUW_1293 | 1.088382 | 0.92623 | -0.23274 | 0.021381 |

| | | | | |
|---|---|---|---|---|
| queuine tRNA-ribosyltransferase | 1.032864 | 0.933477 | -0.14596 | 0.021627 |
| adenosine deaminase...1836 | 1.128149 | 0.899606 | -0.32659 | 0.021773 |
| hypothetical protein ABUW_1123 | 0.904092 | 1.008235 | 0.157291 | 0.02178 |
| hypothetical protein ABUW_0851 | 1.034212 | 0.993118 | -0.0585 | 0.021799 |
| hypothetical protein ABUW_0909 | 1.020639 | 0.958996 | -0.08988 | 0.021805 |
| activator of HSP90 ATPase...2129 | 1.112042 | 0.833894 | -0.41528 | 0.021876 |
| hypothetical protein ABUW_2240 | 1.118835 | 0.851225 | -0.39438 | 0.021913 |
| hypothetical protein ABUW_0466 | 1.052733 | 1.275925 | 0.277404 | 0.021944 |
| Rhs element Vgr protein, putative | 0.968425 | 1.050379 | 0.117197 | 0.022049 |
| ribosomal protein S6 | 1.029077 | 0.992714 | -0.0519 | 0.022076 |
| D-lactate dehydrogenase | 0.862607 | 1.041517 | 0.27191 | 0.0221 |
| ATP-dependent Clp protease adaptor protein | 1.082352 | 0.969401 | -0.159 | 0.022226 |
| universal stress protein family...2453 | 1.077372 | 0.900599 | -0.25856 | 0.022321 |
| transcriptional repressor BetI | 0.918049 | 1.034576 | 0.172398 | 0.022328 |
| hypothetical protein ABUW_3349 | 0.90194 | 1.094823 | 0.279595 | 0.02234 |
| RNA binding S1 domain protein | 1.037773 | 0.910778 | -0.18832 | 0.022395 |
| alpha/beta hydrolase...2175 | 1.034529 | 0.919804 | -0.16958 | 0.022415 |
| DNA repair protein RadC | 0.904155 | 1.176428 | 0.379771 | 0.022507 |
| transcriptional regulator, TetR family...915 | 0.894553 | 1.127196 | 0.333499 | 0.02253 |
| hypothetical protein ABUW_1031 | 1.049375 | 0.936661 | -0.16393 | 0.022589 |
| ureidoglycolate hydrolase | 1.006761 | 1.133186 | 0.170664 | 0.022683 |
| argininosuccinate synthase | 0.967256 | 1.055551 | 0.126027 | 0.022713 |
| 2-isopropylmalate synthase | 0.92617 | 1.010453 | 0.125653 | 0.022789 |
| putative transcriptional regulator...505 | 1.500494 | 0.639309 | -1.23085 | 0.022933 |
| copper/zinc superoxide dismutase | 1.147285 | 0.896363 | -0.35607 | 0.022943 |
| fatty oxidation complex, alpha subunit FadB | 1.074211 | 0.98862 | -0.11979 | 0.02299 |
| Fe-S protein assembly chaperone HscA | 1.004061 | 1.061786 | 0.080646 | 0.023048 |
| primosomal protein N` | 0.90902 | 1.130318 | 0.314345 | 0.023209 |
| hypothetical protein ABUW_1592 | 1.057714 | 0.988874 | -0.09709 | 0.023255 |
| DNA polymerase III, beta subunit | 0.97471 | 1.021533 | 0.067691 | 0.023344 |
| NADH dehydrogenase I chain G | 0.899889 | 1.055399 | 0.229969 | 0.023356 |
| GGDEF domain protein | 0.982866 | 0.836761 | -0.23218 | 0.023384 |
| transglutaminase | 0.933592 | 1.285951 | 0.461973 | 0.023462 |
| type VI secretion-associated protein, ImpA family | 0.888693 | 1.04014 | 0.227019 | 0.023481 |
| hypothetical protein ABUW_3070 | 0.860321 | 1.058546 | 0.299137 | 0.023611 |
| hypothetical protein ABUW_2894 | 0.947916 | 1.07824 | 0.185848 | 0.02362 |
| site-specific DNA-methyltransferase | 1.039139 | 1.32591 | 0.351594 | 0.023645 |
| hypothetical protein ABUW_2442 | 0.616487 | 0.936864 | 0.603769 | 0.023704 |
| hypothetical protein ABUW_1065 | 1.045897 | 0.889475 | -0.23371 | 0.023796 |
| putative metal-dependent hydrolase | 0.873425 | 0.742059 | -0.23515 | 0.023867 |
| integration host factor, beta subunit | 1.121744 | 0.904846 | -0.31 | 0.023882 |
| hypothetical protein ABUW_3493 | 1.008133 | 0.910521 | -0.14692 | 0.024021 |
| UDP-3-O-[3-hydroxymyristoyl] N- acetylglucosamine deacetyl | 0.957738 | 1.074397 | 0.165824 | 0.024121 |
| hypothetical protein ABUW_1262 | 0.845315 | 1.269449 | 0.586641 | 0.024393 |
| anhydro-N-acetylmuramic acid kinase | 1.092597 | 0.988275 | -0.14478 | 0.024447 |
| protein YegH | 0.985358 | 1.037274 | 0.074078 | 0.024461 |
| aldehyde dehydrogenase...2252 | 0.939793 | 1.084361 | 0.20643 | 0.024476 |

| | | | | |
|---|---|---|---|---|
| two-component system response regulator protein...219 | 0.981062 | 0.832809 | -0.23636 | 0.024676 |
| orotate phosphoribosyltransferase | 0.942755 | 1.097411 | 0.21915 | 0.024699 |
| putative UDP-galactose phosphate transferase (WeeH) | 1.015869 | 0.845081 | -0.26555 | 0.024851 |
| hypothetical protein ABUW_0596 | 0.978332 | 1.115805 | 0.189689 | 0.025036 |
| homoserine O-acetyltransferase | 0.988197 | 0.958391 | -0.04418 | 0.025069 |
| RNA polymerase sigma factor sigma-70 | 1.024387 | 0.961504 | -0.0914 | 0.025182 |
| NADH dehydrogenase I chain CD | 0.955669 | 1.03919 | 0.120877 | 0.025322 |
| amino acid transport protein | 0.832522 | 1.026639 | 0.302368 | 0.025363 |
| succinylglutamate desuccinylase | 1.03159 | 0.941937 | -0.13117 | 0.025438 |
| transporter, anion:cation symporter (ACS) family | 1.063806 | 0.799185 | -0.41263 | 0.025463 |
| alkaline phosphatase | 0.907345 | 1.150695 | 0.342782 | 0.025521 |
| stringent starvation protein A | 1.076152 | 0.890972 | -0.27243 | 0.025582 |
| NAD(P) transhydrogenase subunit alpha...2399 | 0.654728 | 0.843615 | 0.36569 | 0.025604 |
| signal peptidase I | 0.950176 | 0.993665 | 0.064564 | 0.025623 |
| hypothetical protein ABUW_2022 | 1.002701 | 0.970715 | -0.04677 | 0.025701 |
| transporter, LysE family | 0.747656 | 1.547249 | 1.049259 | 0.025707 |
| L-lactate permease | 0.918932 | 1.19452 | 0.378402 | 0.025723 |
| hypothetical protein ABUW_1601 | 1.12378 | 0.944696 | -0.25044 | 0.025787 |
| auxin efflux Carrier | 1.206231 | 0.89794 | -0.42582 | 0.025791 |
| aconitate hydratase 1 | 1.091914 | 0.932772 | -0.22726 | 0.025882 |
| acyl-CoA dehydrogenase...2006 | 1.01632 | 0.967753 | -0.07064 | 0.026048 |
| iscRSUA operon repressor | 1.103666 | 0.964045 | -0.19513 | 0.026063 |
| 16S rRNA processing protein RimM | 1.046963 | 0.931215 | -0.16903 | 0.026185 |
| MscS Mechanosensitive ion channel...923 | 0.911772 | 1.068894 | 0.229373 | 0.026227 |
| penicillin-binding protein 6 (D-alanyl-D-alanine carboxypeptid | 0.971672 | 1.117176 | 0.201315 | 0.026277 |
| resolvase (plasmid) | 1.115754 | 0.901586 | -0.30748 | 0.026279 |
| guanosine-3,5-bis(diphosphate) 3-pyrophosphohydrolase | 1.033476 | 0.930871 | -0.15085 | 0.026451 |
| transcriptional regulator, XRE family...93 | 1.126609 | 1.021806 | -0.14087 | 0.026508 |
| twitching motility protein | 1.214004 | 0.958847 | -0.3404 | 0.026671 |
| UDP-N-acetylmuramate--alanine ligase | 0.974953 | 1.052402 | 0.110281 | 0.026923 |
| NAD(+) kinase | 0.910679 | 1.061096 | 0.220541 | 0.026997 |
| hypothetical protein ABUW_3304 | 0.942125 | 1.057416 | 0.166554 | 0.027169 |
| alpha/beta fold family hydrolase...856 | 1.009184 | 0.917227 | -0.13784 | 0.027178 |
| 1-acyl-sn-glycerol-3-phosphate acyltransferase | 0.908682 | 1.040717 | 0.19573 | 0.027401 |
| hypothetical protein ABUW_0181 | 1.150575 | 0.96806 | -0.24919 | 0.027532 |
| hypothetical protein ABUW_2809 | 0.865065 | 1.060149 | 0.293387 | 0.027554 |
| GTP pyrophosphokinase (ppGpp synthetase I) | 1.02261 | 0.905783 | -0.17502 | 0.027564 |
| hypothetical protein ABUW_1426 | 0.876593 | 1.014555 | 0.210867 | 0.027766 |
| glutamate/aspartate transport system permease protein GltJ | 1.008581 | 0.888437 | -0.18299 | 0.028037 |
| Ser/Thr protein phosphatase family protein | 0.857302 | 1.076311 | 0.328218 | 0.028112 |
| glutamate/aspartate ABC transporter, periplasmic glutamate/ | 1.181256 | 0.837952 | -0.49538 | 0.028133 |
| homocysteine S-methyltransferase family protein | 1.044475 | 0.945931 | -0.14297 | 0.028146 |
| hypothetical protein ABUW_2685 | 0.797855 | 0.914591 | 0.197 | 0.028199 |
| hypothetical protein ABUW_0441 | 1.079899 | 0.778745 | -0.47167 | 0.028215 |
| UDP-2,3-diacylglucosamine hydrolase | 1.258769 | 0.933031 | -0.43202 | 0.02831 |
| curved DNA-binding protein | 1.022321 | 0.94934 | -0.10685 | 0.028339 |
| putative two-component response regulator | 1.144695 | 1.021645 | -0.16407 | 0.028369 |

| | | | | |
|---|---|---|---|---|
| transthyretin | 0.801833 | 1.208739 | 0.592129 | 0.028383 |
| type VI secretion system effector, Hcp1 family | 0.707765 | 0.811128 | 0.196659 | 0.028548 |
| hypothetical protein ABUW_2658 | 1.585364 | 0.695282 | -1.18914 | 0.028583 |
| succinate-semialdehyde dehydrogenase (NADP+)...2103 | 1.018183 | 0.929181 | -0.13196 | 0.028692 |
| multidrug efflux protein AdeB | 1.041857 | 0.958774 | -0.1199 | 0.028813 |
| enoyl-CoA hydratase/isomerase...1314 | 1.031641 | 0.955561 | -0.11052 | 0.028834 |
| cell division protein FtsQ | 0.93781 | 1.055278 | 0.170256 | 0.028932 |
| ribosomal protein L11 methyltransferase...1676 | 1.049335 | 0.940007 | -0.15873 | 0.02897 |
| transcriptional regulator, TetR family...1022 | 1.04677 | 0.903229 | -0.21278 | 0.02898 |
| hypothetical protein ABUW_3271 | 0.931236 | 1.09377 | 0.23209 | 0.029082 |
| ferric acinetobactin transport system periplasmic binding prot | 0.956488 | 1.125829 | 0.23517 | 0.029206 |
| ribosomal protein L19 | 1.052054 | 0.953411 | -0.14204 | 0.029269 |
| TrwC protein (plasmid) | 0.767476 | 1.16474 | 0.601815 | 0.029273 |
| phosphate transport system regulatory protein PhoU | 1.021815 | 0.943806 | -0.11457 | 0.029373 |
| hypothetical protein ABUW_0931 | 1.049286 | 0.88927 | -0.23871 | 0.0294 |
| GntP family transporter | 0.923759 | 1.113527 | 0.269548 | 0.029431 |
| shikimate transporter | 0.749687 | 1.000472 | 0.41632 | 0.029461 |
| aminoacyl-histidine dipeptidase | 1.018078 | 0.919777 | -0.14649 | 0.029488 |
| aminotransferase | 0.934842 | 1.098563 | 0.232823 | 0.029557 |
| nucleoside-diphosphate-sugar epimerase...1621 | 1.051963 | 0.937758 | -0.1658 | 0.02977 |
| putative hemolysin | 1.11415 | 0.795204 | -0.48655 | 0.029785 |
| transcriptional regulator, LysR family...1463 | 0.995828 | 1.043558 | 0.067541 | 0.030107 |
| ribosomal protein S14 | 1.149 | 0.946003 | -0.28046 | 0.030452 |
| lipoprotein, putative...997 | 0.896675 | 0.996084 | 0.151682 | 0.030576 |
| acyl-CoA dehydrogenase...1248 | 1.187247 | 0.905502 | -0.39083 | 0.030849 |
| CBS domain containing protein | 1.111049 | 0.875558 | -0.34365 | 0.030938 |
| acetyl-CoA carboxylase, carboxyl transferase, beta subunit | 1.065488 | 0.99069 | -0.10501 | 0.03104 |
| cold-shock DNA-binding domain protein...1533 | 0.399187 | 0.781045 | 0.96834 | 0.031162 |
| alternative sigma factor RpoH (EsvH) | 1.000977 | 1.092972 | 0.126848 | 0.0313 |
| malonyl CoA-acyl carrier protein transacylase | 1.095974 | 0.976588 | -0.16639 | 0.031599 |
| hypothetical protein ABUW_2679 | 1.197706 | 0.844105 | -0.50478 | 0.031687 |
| ABC-1 domain protein...2113 | 1.031496 | 0.972855 | -0.08444 | 0.031739 |
| 4-hydroxyphenylpyruvate dioxygenase | 1.075581 | 0.943496 | -0.18903 | 0.031885 |
| molybdenum cofactor biosynthesis protein C | 0.90742 | 1.01607 | 0.163157 | 0.032073 |
| hypothetical protein ABUW_2443 | 1.226387 | 0.589229 | -1.05751 | 0.032279 |
| arginyl-tRNA synthetase | 1.00998 | 0.954925 | -0.08087 | 0.032291 |
| short-chain dehydrogenase/reductase...2238 | 1.058313 | 0.954663 | -0.1487 | 0.032427 |
| chorismate mutase...2047 | 0.952594 | 1.09425 | 0.200009 | 0.032803 |
| hypothetical protein ABUW_1828 | 0.799372 | 1.152316 | 0.527598 | 0.032983 |
| ABC transporter, periplasmic binding protein | 1.065 | 0.974086 | -0.12873 | 0.033373 |
| heme-binding protein A...945 | 1.010528 | 0.937936 | -0.10755 | 0.033446 |
| ATP-dependent Clp protease, proteolytic subunit ClpP | 1.034848 | 0.909932 | -0.18559 | 0.033632 |
| aspartate kinase | 0.921749 | 1.011651 | 0.134265 | 0.0337 |
| phospholipase D | 0.943003 | 1.225403 | 0.377922 | 0.033752 |
| hypothetical protein ABUW_0527 | 0.795035 | 1.013545 | 0.35032 | 0.033764 |
| ribosomal protein S16 | 1.065097 | 0.978732 | -0.122 | 0.033946 |
| prophage LambdaCh01, coat protein | 1.086881 | 0.952163 | -0.19091 | 0.034084 |

| | | | | |
|---|---|---|---|---|
| hypothetical protein ABUW_3434 | 1.09035 | 1.006048 | -0.11609 | 0.034157 |
| crispr-associated protein Cas1 | 1.076821 | 0.845407 | -0.34906 | 0.034164 |
| chorismate mutase family protein | 0.886812 | 1.080642 | 0.285188 | 0.034187 |
| Holliday junction DNA helicase RuvA | 0.974996 | 1.020003 | 0.065106 | 0.034311 |
| beta-ketoadipyl CoA thiolase...1559 | 0.913491 | 1.069342 | 0.22726 | 0.034319 |
| C4-dicarboxylate transport protein...1698 | 0.920926 | 1.092488 | 0.246459 | 0.034434 |
| hypothetical protein ABUW_3234 | 0.864873 | 1.1277 | 0.382824 | 0.034445 |
| hypothetical protein ABUW_2143 | 0.881123 | 1.127324 | 0.355487 | 0.034481 |
| transferase hexapeptide repeat protein | 0.997711 | 1.139199 | 0.191326 | 0.034515 |
| amidase | 0.949664 | 1.108723 | 0.223409 | 0.034662 |
| hypothetical protein ABUW_3516 | 1.00137 | 0.92544 | -0.11376 | 0.034676 |
| transcriptional regulator AcrR family | 1.053792 | 0.995167 | -0.08258 | 0.034717 |
| ThiJ/PfpI domain protein | 1.166786 | 0.838587 | -0.47651 | 0.034747 |
| cytochrome O ubiquinol oxidase, subunit I | 0.900745 | 1.048909 | 0.2197 | 0.034802 |
| ATP synthase F0, A subunit | 0.881702 | 1.070037 | 0.279299 | 0.034848 |
| glutamyl-tRNA reductase | 0.943732 | 1.066108 | 0.175903 | 0.034882 |
| succinyl-CoA synthetase, alpha subunit | 0.960245 | 1.1127 | 0.212589 | 0.035069 |
| 3-dehydroquinate dehydratase, type I | 0.980444 | 1.040165 | 0.085304 | 0.035103 |
| transcriptional regulator, TetR family...1418 | 1.028843 | 0.878917 | -0.22722 | 0.035211 |
| nucleoside-diphosphate-sugar epimerase...1892 | 0.99556 | 0.899159 | -0.14693 | 0.035222 |
| rubredoxin reductase | 0.969938 | 1.112666 | 0.198056 | 0.035472 |
| phenylacetate-CoA ligase | 0.898457 | 1.032782 | 0.201015 | 0.035504 |
| hypothetical protein ABUW_1651 | 0.680815 | 0.978482 | 0.523282 | 0.03556 |
| glutamate synthase, small subunit | 0.960011 | 1.063635 | 0.14788 | 0.03567 |
| VacJ family lipoprotein | 0.974026 | 1.085443 | 0.156252 | 0.035735 |
| 3-oxoacyl-(acyl carrier protein) synthase | 0.939054 | 1.079769 | 0.201442 | 0.035793 |
| hypothetical protein ABUW_0153 | 0.955287 | 1.089446 | 0.189588 | 0.035823 |
| hypothetical protein ABUW_1860 | 0.803873 | 0.993309 | 0.305274 | 0.035953 |
| hypothetical protein ABUW_1879 | 0.952886 | 1.06892 | 0.165778 | 0.036053 |
| hypothetical protein ABUW_0248 | 1.127417 | 1.007297 | -0.16253 | 0.036131 |
| hypothetical protein ABUW_5012 (plasmid) | 0.977218 | 0.911598 | -0.10028 | 0.036149 |
| acetolactate synthase, small subunit | 0.901886 | 1.083888 | 0.265199 | 0.036284 |
| hypothetical protein ABUW_3282 | 0.906822 | 1.106708 | 0.287384 | 0.036589 |
| diaminobutyrate decarboxylase | 0.985212 | 1.010033 | 0.035895 | 0.036624 |
| hypothetical protein ABUW_0593 | 0.956142 | 1.156205 | 0.2741 | 0.036753 |
| YD repeat protein | 0.905947 | 1.071899 | 0.242671 | 0.036972 |
| lipoprotein-releasing system transmembrane protein LolE | 0.905773 | 1.017173 | 0.167344 | 0.03703 |
| ErfK/YbiS/YcfS/YnhG family protein | 0.931227 | 1.053204 | 0.177579 | 0.037105 |
| putative transferase | 1.240404 | 0.998972 | -0.31229 | 0.037419 |
| hypothetical protein ABUW_0134 | 0.974558 | 1.045395 | 0.101229 | 0.037426 |
| hypothetical protein ABUW_2439 | 1.65802 | 0.698081 | -1.24799 | 0.037455 |
| flavodoxin/nitric oxide synthase...1244 | 0.937673 | 1.019782 | 0.121103 | 0.03783 |
| hypothetical protein ABUW_3470 | 1.026988 | 0.901101 | -0.18866 | 0.037898 |
| 3-oxoadipate CoA-transferase subunit B...213 | 0.894045 | 0.945962 | 0.081434 | 0.037991 |
| hypothetical protein ABUW_0398 | 0.836456 | 0.996045 | 0.251922 | 0.038116 |
| putative periplasmic carboxyl-terminal protease | 0.92032 | 1.229678 | 0.418073 | 0.038296 |
| hypothetical protein ABUW_1746 | 1.095716 | 0.928299 | -0.23921 | 0.038433 |

| | | | | |
|---|---|---|---|---|
| ribulose-phosphate 3-epimerase | 0.969583 | 1.060969 | 0.129945 | 0.038439 |
| non-ribosomal peptide synthetase...420 | 1.154765 | 0.918767 | -0.32983 | 0.038593 |
| hypothetical protein ABUW_4006 (plasmid) | 0.944663 | 1.05311 | 0.156785 | 0.03879 |
| glutaredoxin-related protein | 0.953933 | 1.125589 | 0.23872 | 0.039129 |
| NADP-dependent fatty aldehyde dehydrogenase...786 | 0.997662 | 1.053881 | 0.079089 | 0.039149 |
| flavodoxin/nitric oxide synthase...989 | 1.017433 | 0.867328 | -0.23028 | 0.039151 |
| transcriptional regulator TetR/AcrR family | 1.175362 | 0.910854 | -0.36781 | 0.039152 |
| D-serine ammonia-lyase | 0.968198 | 1.041205 | 0.104881 | 0.039264 |
| hypothetical protein ABUW_0986 | 1.051037 | 1.157539 | 0.139247 | 0.039521 |
| hypothetical protein ABUW_0490 | 1.14916 | 0.826064 | -0.47625 | 0.039634 |
| aminodeoxychorismate lyase...1570 | 0.899374 | 1.010816 | 0.168527 | 0.039635 |
| 2-amino-4-hydroxy-6-hydroxymethyldihydropteridine pyroph | 0.918992 | 1.043343 | 0.183088 | 0.039654 |
| hypothetical protein ABUW_1536 | 1.215921 | 0.822432 | -0.56408 | 0.039939 |
| tRNA--hydroxylase | 0.914309 | 1.091147 | 0.255092 | 0.040013 |
| hypothetical protein ABUW_3235 | 0.789308 | 1.117231 | 0.501268 | 0.040148 |
| hypothetical protein ABUW_3458 | 1.213987 | 0.885063 | -0.4559 | 0.040304 |
| ribonucleoside-diphosphate reductase, beta subunit | 1.044135 | 0.918884 | -0.18435 | 0.040372 |
| ATP synthase F0, C subunit | 0.918348 | 1.154215 | 0.329799 | 0.040456 |
| hypothetical protein ABUW_3360 | 1.00654 | 0.951056 | -0.0818 | 0.040796 |
| hypothetical protein ABUW_2886 | 1.003919 | 1.117549 | 0.154694 | 0.040905 |
| hypothetical protein ABUW_0096 | 0.865878 | 1.017992 | 0.233491 | 0.040978 |
| monooxygenase | 1.027612 | 0.975242 | -0.07546 | 0.041028 |
| hypothetical protein ABUW_2599 | 0.842502 | 1.154059 | 0.453964 | 0.0411 |
| hypothetical protein ABUW_1784 | 0.929129 | 0.811643 | -0.19503 | 0.041215 |
| chromosome segregation protein SMC | 0.804839 | 1.192675 | 0.567429 | 0.041255 |
| phosphate acetyltransferase | 1.165716 | 0.9193 | -0.34261 | 0.041293 |
| ribonuclease | 1.110316 | 0.853203 | -0.38001 | 0.04133 |
| transcriptional regulatory protein | 1.163567 | 0.919753 | -0.33924 | 0.041418 |
| magnesium and cobalt transport protein | 0.848026 | 1.084977 | 0.355485 | 0.041588 |
| permease...646 | 0.946128 | 1.045819 | 0.144526 | 0.041638 |
| ThiF family protein | 1.046942 | 1.152101 | 0.138086 | 0.041686 |
| ATP-dependent protease La | 1.023273 | 0.960538 | -0.09128 | 0.041787 |
| endonuclease/exonuclease/phosphatase...484 | 0.95389 | 1.080363 | 0.179622 | 0.041991 |
| putative ATP-dependent DNA helicase | 1.097417 | 0.961109 | -0.19134 | 0.04202 |
| glutaminyl-tRNA synthetase | 0.968234 | 1.090997 | 0.172219 | 0.042265 |
| hypothetical protein ABUW_0895 | 0.878696 | 0.988372 | 0.169691 | 0.042873 |
| porin...414 | 1.111584 | 0.910195 | -0.28837 | 0.042888 |
| riboflavin biosynthesis protein RibD | 0.981855 | 1.02645 | 0.064081 | 0.043181 |
| hypothetical protein ABUW_0760 | 1.396131 | 0.696914 | -1.00238 | 0.043293 |
| phosphoenolpyruvate carboxykinase | 0.871378 | 0.997024 | 0.19433 | 0.043367 |
| Na+/H+ antiporter NhaC | 0.992031 | 1.131869 | 0.190251 | 0.043489 |
| hypothetical protein ABUW_0668 | 1.041978 | 0.895778 | -0.21811 | 0.043555 |
| short chain dehydrogenase...2341 | 1.105596 | 0.927011 | -0.25416 | 0.043682 |
| lipoprotein, putative...942 | 0.995161 | 1.154366 | 0.214098 | 0.043929 |
| isochorismatase hydrolase...2114 | 0.942734 | 1.151288 | 0.288327 | 0.04406 |
| hypothetical protein ABUW_0027 | 0.96403 | 1.045697 | 0.117316 | 0.044095 |
| 3-oxoacid CoA-transferase, subunit A | 0.989196 | 1.116585 | 0.174765 | 0.044365 |

| | | | | |
|---|---|---|---|---|
| putative aminotransferase | 1.007377 | 0.971808 | -0.05186 | 0.044453 |
| ABC transporter ATP-binding protein uup | 0.915036 | 1.137032 | 0.313373 | 0.044482 |
| metallo-beta-lactamase family protein...1683 | 0.96627 | 1.02334 | 0.082788 | 0.04456 |
| amino acid adenylation | 0.844838 | 1.116944 | 0.402809 | 0.044562 |
| integral membrane protein MviN | 0.918421 | 1.035075 | 0.172507 | 0.044605 |
| histidine kinase | 0.956423 | 1.029496 | 0.106217 | 0.044607 |
| hypothetical protein ABUW_2363 | 0.702711 | 0.943459 | 0.425029 | 0.044863 |
| hypothetical protein ABUW_0040 | 1.10456 | 0.780939 | -0.50019 | 0.044997 |
| glyceraldehyde 3-phosphate dehydrogenase | 0.947003 | 1.011013 | 0.094362 | 0.045249 |
| transcriptional regulator, DUF24 family | 0.49312 | 0.415542 | -0.24695 | 0.045306 |
| Putative DNA binding protein (plasmid) | 1.058825 | 0.890647 | -0.24954 | 0.045503 |
| cold-shock DNA-binding domain protein...522 | 0.647389 | 0.926674 | 0.517428 | 0.045519 |
| X-Pro aminopeptidase | 0.877069 | 1.058364 | 0.271075 | 0.045729 |
| DNA polymerase III subunit alpha | 0.942613 | 1.010631 | 0.10052 | 0.045869 |
| acetyltransferase, gnat family...1091 | 1.029391 | 0.964743 | -0.09358 | 0.045878 |
| acetyltransferase, gnat family...850 | 0.942752 | 1.045263 | 0.148916 | 0.046105 |
| hypothetical protein ABUW_2511 | 0.987769 | 1.073714 | 0.120364 | 0.046133 |
| aldehyde dehydrogenase...1581 | 0.893126 | 1.013441 | 0.182327 | 0.046259 |
| hypothetical protein ABUW_2512 | 0.286708 | 0.757049 | 1.400803 | 0.046319 |
| acetoin:2,6-dichlorophenolindophenol oxidoreductase alpha s | 0.942821 | 0.833152 | -0.1784 | 0.046383 |
| enoyl-CoA hydratase/isomerase...1946 | 1.238045 | 0.750822 | -0.72152 | 0.046443 |
| pilus assembly protein tip-associated adhesin PilY1 | 1.061709 | 1.251082 | 0.236788 | 0.046673 |
| hypothetical protein ABUW_2647 | 0.892205 | 1.025553 | 0.200955 | 0.046843 |
| ferric acinetobactin receptor | 0.986306 | 1.073688 | 0.122468 | 0.046852 |
| hypothetical protein ABUW_1812 | 1.071182 | 0.954378 | -0.16657 | 0.047121 |
| short chain dehydrogenase...458 | 1.023345 | 0.918627 | -0.15574 | 0.047218 |
| hypothetical protein ABUW_2158 | 1.408924 | 0.775469 | -0.86145 | 0.047312 |
| acetyltransferase, GNAT family | 1.12276 | 0.894492 | -0.32791 | 0.047314 |
| hypothetical protein ABUW_2085 | 0.964818 | 1.237172 | 0.358717 | 0.047335 |
| histidinol-phosphate aminotransferase | 0.962513 | 1.032729 | 0.101584 | 0.047337 |
| hypothetical protein ABUW_2831 | 0.919819 | 1.104986 | 0.264606 | 0.047525 |
| methionyl-tRNA synthetase | 1.014647 | 0.946921 | -0.09966 | 0.047592 |
| adenosine deaminase...1044 | 1.054086 | 0.851602 | -0.30774 | 0.047693 |
| transcriptional regulator, TetR family...627 | 1.024338 | 0.962685 | -0.08956 | 0.048082 |
| lipid-A-disaccharide synthase | 0.813652 | 0.99853 | 0.295394 | 0.048106 |
| hypothetical protein ABUW_1084 | 1.017028 | 0.854821 | -0.25066 | 0.048246 |
| transcriptional regulator, LysR family...1435 | 1.068137 | 0.987337 | -0.11348 | 0.048312 |
| S-adenosylmethionine:tRNA ribosyltransferase-isomerase | 0.972175 | 1.05028 | 0.111486 | 0.048636 |
| ATP-dependent helicase HrpA | 0.87179 | 1.026271 | 0.235359 | 0.048813 |
| esterase...1086 | 0.918502 | 1.06566 | 0.214392 | 0.049084 |
| N-acetyl-gamma-glutamyl-phosphate reductase | 0.929218 | 1.08805 | 0.227657 | 0.049154 |
| hypothetical protein ABUW_3158 | 1.259476 | 0.760245 | -0.72829 | 0.049403 |
| transcription termination/antitermination factor NusG | 0.965845 | 1.092184 | 0.177353 | 0.049586 |
| NADH dehydrogenase fad-containing subunit | 1.077623 | 0.744744 | -0.53304 | 0.049649 |
| choline dehydrogenase | 0.958869 | 1.00356 | 0.065722 | 0.049714 |
| hypothetical protein ABUW_1734 | 0.966664 | 1.070433 | 0.147108 | 0.049789 |
| dioxygenase alpha subunit | 1.093321 | 0.92278 | -0.24466 | 0.049997 |

| | | | | |
|---|---|---|---|---|
| uroporphyrinogen decarboxylase | 0.945034 | 1.049348 | 0.151055 | 0.050049 |
| zinc-binding alcohol dehydrogenase | 0.999709 | 0.902419 | -0.14771 | 0.050332 |
| transcriptional regulator, TetR family...1076 | 1.088766 | 0.929597 | -0.22802 | 0.050507 |
| NAD dependent epimerase/dehydratase family...39 | 1.122741 | 0.872809 | -0.36329 | 0.050788 |
| transcriptional regulator, LysR family...273 | 1.049512 | 0.902254 | -0.21811 | 0.050837 |
| luciferase family monooxygenase | 1.011833 | 0.939607 | -0.10684 | 0.051032 |
| non-ribosomal peptide synthetase...503 | 0.937254 | 1.032305 | 0.139357 | 0.051085 |
| isocitrate lyase | 1.049813 | 0.958948 | -0.13061 | 0.051135 |
| transcriptional regulator, LysR family...1353 | 0.952273 | 1.04854 | 0.138934 | 0.051302 |
| UTP-glucose-1-phosphate uridylyltransferase | 1.039747 | 0.891762 | -0.2215 | 0.051305 |
| 23S rRNA (uracil-5-)-methyltransferase RumA | 0.910316 | 1.056645 | 0.215053 | 0.051563 |
| putative L-asparaginase I | 1.022361 | 0.968185 | -0.07855 | 0.051946 |
| ferredoxin, 2Fe-2S type | 1.091077 | 0.986487 | -0.14538 | 0.051962 |
| melanin biosynthesis protein TyrA | 0.949267 | 1.036776 | 0.127218 | 0.052015 |
| UDP-N-acetylmuramoylalanyl-D-glutamyl-2,6-diaminopimelat | 0.984597 | 1.079009 | 0.132101 | 0.052095 |
| glutamate/aspartate transport ATP-binding protein GltL | 1.092335 | 0.855894 | -0.35191 | 0.052121 |
| transcriptional regulator, Fur family | 1.019184 | 0.968198 | -0.07404 | 0.052198 |
| NAD(P) transhydrogenase subunit alpha...605 | 0.661442 | 0.759886 | 0.200168 | 0.052227 |
| excinuclease ABC, B subunit | 0.995695 | 1.026138 | 0.043449 | 0.052366 |
| transcriptional regulator, AraC family...996 | 1.042841 | 0.951037 | -0.13295 | 0.05238 |
| formamidopyrimidine-DNA glycosylase | 1.070624 | 0.996403 | -0.10365 | 0.05256 |
| glutamyl-tRNA synthetase...2153 | 0.956258 | 1.043394 | 0.125813 | 0.052647 |
| heme-binding protein A...344 | 0.969814 | 0.794713 | -0.28727 | 0.052878 |
| two component signal transduction system response regulatc | 1.000097 | 1.066836 | 0.093199 | 0.053027 |
| DNA-directed RNA polymerase, alpha subunit | 1.027656 | 0.980496 | -0.06777 | 0.053129 |
| secretion protein HlyD...1783 | 1.012101 | 0.947627 | -0.09496 | 0.053375 |
| hypothetical protein ABUW_3829 | 0.966931 | 1.066418 | 0.141289 | 0.053403 |
| hypothetical protein ABUW_2200 | 1.017275 | 0.890939 | -0.19131 | 0.053645 |
| nucleotide-binding protein | 0.95297 | 1.025913 | 0.106405 | 0.053981 |
| pirin domain protein...1396 | 0.970012 | 1.027106 | 0.08251 | 0.054161 |
| aminoglycoside phosphotransferase...1494 | 0.952918 | 1.052786 | 0.143788 | 0.054298 |
| hypothetical protein ABUW_0498 | 1.060668 | 0.816139 | -0.37809 | 0.054339 |
| hypothetical protein ABUW_1051 | 1.06998 | 0.975937 | -0.13272 | 0.054691 |
| succinyl-CoA:coenzyme A transferase | 1.154605 | 1.079909 | -0.09649 | 0.054897 |
| hypothetical protein ABUW_3688 | 0.901837 | 1.029041 | 0.190362 | 0.054912 |
| hypothetical protein ABUW_3375 | 0.950228 | 1.002941 | 0.077891 | 0.055056 |
| heme oxygenase-like protein...150 | 1.040097 | 0.856382 | -0.28039 | 0.055195 |
| iron-sulfur cluster assembly accessory protein | 0.794804 | 1.222454 | 0.621109 | 0.055427 |
| porin...28 | 0.71365 | 0.910863 | 0.352018 | 0.055436 |
| nitroreductase family protein...1859 | 1.066646 | 0.926659 | -0.20297 | 0.055564 |
| DNA polymerase III subunit tau | 1.01067 | 0.964432 | -0.06756 | 0.055573 |
| N-ethylmaleimide reductase | 1.167987 | 0.84617 | -0.465 | 0.055916 |
| glutamine-dependent NAD+ synthetase | 0.960334 | 0.99512 | 0.051334 | 0.056025 |
| outer membrane protein A | 1.057225 | 0.919352 | -0.20159 | 0.056119 |
| hypothetical protein ABUW_3577 | 1.082525 | 0.868394 | -0.31798 | 0.056308 |
| ATP-dependent dsDNA exonuclease...229 | 0.891998 | 1.187847 | 0.413237 | 0.056309 |
| DNA mismatch repair protein MutS | 0.981615 | 1.054776 | 0.103707 | 0.056388 |

| | | | | |
|---|---|---|---|---|
| fumarate hydratase, class II | 0.971878 | 1.012392 | 0.05892 | 0.056439 |
| 2-nitropropane dioxygenase...1525 | 1.068761 | 0.961896 | -0.15199 | 0.056473 |
| putative enoyl-CoA hydratase/isomerase | 1.037431 | 0.911846 | -0.18615 | 0.056523 |
| septum site-determining protein MinD | 1.038479 | 0.963285 | -0.10844 | 0.056568 |
| methyltransferase GidB | 0.985826 | 1.029924 | 0.063133 | 0.056608 |
| rubredoxin | 0.863503 | 1.191078 | 0.463995 | 0.056664 |
| amino-acid N-acetyltransferase | 0.941317 | 1.050802 | 0.158738 | 0.056896 |
| beta-lactamase domain protein | 1.051113 | 0.948784 | -0.14777 | 0.056902 |
| phenylacetic acide degradation protein PaaY | 0.944798 | 1.118379 | 0.243332 | 0.056929 |
| hypothetical protein ABUW_2868 | 0.999501 | 1.077327 | 0.108175 | 0.057356 |
| major facilitator family transporter...98 | 0.788579 | 1.280343 | 0.699203 | 0.057491 |
| peptidyl-prolyl cis-trans isomerase...1400 | 1.059987 | 0.999866 | -0.08424 | 0.057534 |
| ribosomal protein S21 | 1.192555 | 0.93804 | -0.34633 | 0.057541 |
| nonribosomal peptide synthetase BasC | 1.017675 | 0.879032 | -0.21129 | 0.057581 |
| hypothetical protein ABUW_1767 | 0.645648 | 1.058848 | 0.713676 | 0.057634 |
| hypothetical protein ABUW_0786 | 0.912143 | 1.036722 | 0.184696 | 0.057713 |
| ribosomal protein S3 | 1.047746 | 0.959652 | -0.12671 | 0.057786 |
| transcriptional regulator, GntR family...908 | 1.011234 | 0.869235 | -0.2183 | 0.05784 |
| sodium:dicarboxylate symporter | 1.003974 | 0.909509 | -0.14256 | 0.057988 |
| Hca operon transcriptional activator | 0.98128 | 1.049012 | 0.096295 | 0.05799 |
| hypothetical protein ABUW_2043 | 0.956491 | 1.190726 | 0.316019 | 0.058065 |
| exonuclease V beta chain | 1.085741 | 0.94761 | -0.19632 | 0.05846 |
| FAD/FMN-binding/pyridine nucleotide- disulfide oxidoreducta | 1.043316 | 0.944473 | -0.14359 | 0.058498 |
| transglycosylase | 0.982477 | 1.033159 | 0.072567 | 0.058761 |
| hypothetical protein ABUW_0334 | 1.285855 | 0.829842 | -0.63182 | 0.058977 |
| tRNA/rRNA methyltransferase | 1.128068 | 0.892748 | -0.33753 | 0.059023 |
| hypothetical protein ABUW_3051 | 1.028328 | 0.913747 | -0.17043 | 0.059149 |
| outer membrane protein assembly complex BamA | 0.993555 | 1.028638 | 0.050064 | 0.059207 |
| glutamate-5-semialdehyde dehydrogenase | 1.034425 | 0.974095 | -0.08669 | 0.059426 |
| DNA topoisomerase IV, B subunit | 1.024795 | 0.948749 | -0.11124 | 0.059443 |
| hypothetical protein ABUW_0971 | 1.000421 | 1.051313 | 0.071585 | 0.059851 |
| hypothetical protein ABUW_3378 | 0.949455 | 1.066437 | 0.167626 | 0.059949 |
| sensor kinase CusS | 0.945196 | 1.033995 | 0.129545 | 0.060049 |
| 6-pyruvoyl tetrahydrobiopterin synthase | 0.987123 | 0.95464 | -0.04827 | 0.060086 |
| YCII-related protein...752 | 0.870076 | 1.071232 | 0.300057 | 0.060259 |
| 4-carboxymuconolactone decarboxylase...1303 | 1.170995 | 0.90536 | -0.37117 | 0.060352 |
| lipoprotein, putative...469 | 0.803062 | 1.084541 | 0.4335 | 0.060354 |
| transcriptional regulator AraC family | 0.792483 | 1.021483 | 0.366213 | 0.060428 |
| AAA ATPase superfamily protein | 0.940743 | 1.021357 | 0.118615 | 0.060445 |
| fimbrial protein | 1.082212 | 0.885163 | -0.28997 | 0.060707 |
| aminopeptidase N | 0.883055 | 0.754526 | -0.22693 | 0.06073 |
| NAD(P) transhydrogenase subunit beta | 0.815779 | 0.89197 | 0.128817 | 0.060775 |
| DNA polymerase I | 0.961296 | 1.003843 | 0.06248 | 0.060863 |
| bacterioferritin comigratory protein | 0.875579 | 1.140971 | 0.381954 | 0.061214 |
| alpha-methylacyl-CoA racemase | 1.120763 | 0.913729 | -0.29464 | 0.06127 |
| 4-hydroxy-3-methylbut-2-en-1-yl diphosphate synthase | 0.931183 | 1.000059 | 0.102949 | 0.061354 |
| hypothetical protein ABUW_2923 | 0.9585 | 1.102449 | 0.201862 | 0.061404 |

| | | | | |
|---|---|---|---|---|
| exodeoxyribonuclease VII | 0.94532 | 1.061935 | 0.167821 | 0.061482 |
| hypothetical protein ABUW_1034 | 0.840419 | 1.194699 | 0.507467 | 0.061712 |
| acyl-CoA dehydrogenase...1674 | 1.059447 | 0.980421 | -0.11184 | 0.062001 |
| heme oxygenase-like protein...523 | 1.082753 | 0.807189 | -0.42373 | 0.062057 |
| DNA ligase, NAD-dependent | 0.990525 | 1.073903 | 0.116598 | 0.062165 |
| ribosomal protein L27 | 1.113706 | 0.904227 | -0.30061 | 0.062351 |
| GTP-binding protein YchF | 0.972649 | 1.085771 | 0.158729 | 0.062714 |
| penicillin-binding protein 1A | 0.97598 | 1.048379 | 0.103237 | 0.062734 |
| cobyrinic acid a,c-diamide synthase | 0.946462 | 1.115821 | 0.237489 | 0.06292 |
| D-alanyl-D-alanine carboxypeptidase family protein | 0.990921 | 0.852878 | -0.21643 | 0.063084 |
| transcriptional regulator...957 | 1.077555 | 0.989523 | -0.12296 | 0.063156 |
| acetyltransferase...922 | 0.909167 | 0.984165 | 0.114354 | 0.063237 |
| benzoate 1,2-dioxygenase, large subunit | 0.85191 | 0.960138 | 0.172542 | 0.063448 |
| phospho-N-acetylmuramoyl-pentapeptide-transferase | 1.028866 | 1.247714 | 0.278232 | 0.063457 |
| GTP cyclohydrolase I | 0.996605 | 0.94385 | -0.07846 | 0.063619 |
| hypothetical protein ABUW_0118 | 1.002941 | 1.045773 | 0.060333 | 0.063741 |
| alcohol dehydrogenase, iron-containing | 1.135811 | 0.872402 | -0.38066 | 0.063748 |
| hypothetical protein ABUW_0264 | 1.01534 | 1.140982 | 0.168313 | 0.063758 |
| multidrug efflux protein AdeC | 1.005302 | 1.117216 | 0.152279 | 0.063839 |
| preprotein translocase, SecE subunit | 0.923999 | 1.015295 | 0.135935 | 0.063856 |
| feruloyl-CoA synthetase | 1.118703 | 0.882135 | -0.34276 | 0.063906 |
| dihydrofolate reductase | 0.885734 | 1.072067 | 0.27545 | 0.064372 |
| intracellular protease, PfpI family | 1.00836 | 0.961826 | -0.06816 | 0.064395 |
| glycolate/propanediol utilization protein | 1.024117 | 0.932368 | -0.13541 | 0.064684 |
| dihydroxy-acid dehydratase...316 | 0.98845 | 1.140265 | 0.206129 | 0.064896 |
| hypothetical protein ABUW_0099 | 1.085551 | 0.903183 | -0.26534 | 0.065339 |
| hypothetical protein ABUW_2898 | 1.060829 | 0.976773 | -0.1191 | 0.065449 |
| hypothetical protein ABUW_1405 | 1.641942 | 4.066786 | 1.308486 | 0.06566 |
| glutathione S-transferase family protein...1900 | 1.050524 | 0.86105 | -0.28694 | 0.065893 |
| methyltransferase superfamily | 0.822977 | 0.963927 | 0.228072 | 0.066032 |
| phosphate ABC transporter, ATP-binding protein | 0.891057 | 1.168413 | 0.39096 | 0.066207 |
| glutamine-fructose-6-phosphate transaminase (isomerizing) | 0.969993 | 1.034276 | 0.092574 | 0.066407 |
| flavin binding monooxygenase | 0.927561 | 0.997114 | 0.104316 | 0.0665 |
| hypothetical protein ABUW_2607 | 0.965945 | 1.102219 | 0.190398 | 0.066649 |
| hypothetical protein ABUW_3526 | 0.935162 | 0.999551 | 0.096063 | 0.066785 |
| protein-export chaperone SecB | 0.964835 | 1.05683 | 0.13139 | 0.066833 |
| hypothetical protein ABUW_0515 | 1.151263 | 0.92279 | -0.31914 | 0.06692 |
| hypothetical protein ABUW_2858 | 1.05651 | 0.981392 | -0.1064 | 0.067011 |
| phage/plasmid-related protein | 0.783668 | 1.207459 | 0.62366 | 0.067068 |
| hypothetical protein ABUW_1056 | 0.9083 | 0.97181 | 0.097505 | 0.067092 |
| replicative DNA helicase...1112 | 0.940426 | 1.055899 | 0.167085 | 0.067799 |
| outer membrane lipoprotein carrier protein LolA | 1.046658 | 0.966373 | -0.11514 | 0.068146 |
| DNA polymerase IV | 0.998461 | 0.90084 | -0.14843 | 0.068693 |
| glutamine amidotransferase, class-II | 0.970571 | 0.989172 | 0.027389 | 0.068877 |
| acetyl-CoA acyltransferase | 0.997947 | 0.967408 | -0.04484 | 0.068878 |
| fumarylacetoacetate hydrolase...1192 | 1.177415 | 0.838174 | -0.4903 | 0.068979 |
| phospholipid/glycerol acyltransferase...1687 | 0.87493 | 1.035122 | 0.242562 | 0.069263 |

| | | | | |
|---|---|---|---|---|
| dihydrodipicolinate synthase | 0.975573 | 1.073899 | 0.138536 | 0.069436 |
| beta-ribbon domain peptidase S24/S26A/S26B/S26C | 0.993648 | 1.062482 | 0.096632 | 0.069443 |
| peptide deformylase...517 | 0.983343 | 0.944676 | -0.05788 | 0.069558 |
| two-component heavy metal sensor histidine kinase | 0.9965 | 0.752847 | -0.40451 | 0.069598 |
| hypothetical protein ABUW_0187 | 0.945259 | 1.018762 | 0.108036 | 0.069751 |
| hypothetical protein ABUW_2815 | 0.927062 | 1.111673 | 0.261995 | 0.06993 |
| hypothetical protein ABUW_2752 | 0.968404 | 1.029193 | 0.087833 | 0.069961 |
| high affinity Zn transport protein | 1.017436 | 0.926252 | -0.13546 | 0.070049 |
| hypothetical protein ABUW_0274 | 0.941117 | 1.082602 | 0.202057 | 0.070103 |
| protein-tyrosine-phosphatase ptp | 1.03671 | 1.138256 | 0.134812 | 0.070159 |
| helix-turn-helix- domain containing protein, AraC type | 1.054918 | 0.944328 | -0.15977 | 0.070307 |
| thiol:disulfide interchange protein...1845 | 1.003879 | 1.047516 | 0.061387 | 0.070468 |
| hypothetical protein ABUW_1005 | 1.06232 | 0.975533 | -0.12296 | 0.070538 |
| hypothetical protein ABUW_0112 | 0.948379 | 1.04731 | 0.143152 | 0.070587 |
| aminodeoxychorismate lyase...1080 | 1.139285 | 0.889652 | -0.35681 | 0.071194 |
| DsrC family protein | 0.893404 | 0.987863 | 0.144998 | 0.071287 |
| hypothetical protein ABUW_3575 | 1.076299 | 0.955983 | -0.17102 | 0.071496 |
| thiamine biosynthesis protein ThiS | 0.9047 | 1.04164 | 0.203344 | 0.071584 |
| hypothetical protein ABUW_1688 | 0.958824 | 0.910085 | -0.07526 | 0.071793 |
| phosphate regulon transcriptional regulatory protein PhoB | 1.024484 | 0.97777 | -0.06733 | 0.071801 |
| helix-turn-helix domain protein | 1.086687 | 1.02352 | -0.0864 | 0.071959 |
| RNA polymerase sigma-54 factor | 1.016956 | 0.993121 | -0.03422 | 0.072077 |
| ferredoxin-1 | 0.919193 | 1.23797 | 0.429537 | 0.072584 |
| protein YrdC | 1.043012 | 1.161903 | 0.155734 | 0.072584 |
| RNA methyltransferase, TrmH family | 1.059675 | 1.172533 | 0.146006 | 0.072664 |
| transcriptional regulator, GntR family...486 | 0.960351 | 1.047736 | 0.125641 | 0.072714 |
| hypothetical protein ABUW_0923 | 1.007784 | 0.870786 | -0.2108 | 0.072755 |
| copper resistance protein B...592 | 1.032064 | 0.928796 | -0.1521 | 0.072902 |
| hemerythrin | 0.933171 | 1.023683 | 0.133555 | 0.073207 |
| 2,3-dihydro-2,3-dihydroxybenzoate dehydrogenase | 1.068449 | 0.949449 | -0.17036 | 0.07325 |
| alpha/beta hydrolase...1292 | 1.03658 | 0.929531 | -0.15726 | 0.073549 |
| glucose sorbosone dehydrogenase | 1.033567 | 0.97509 | -0.08402 | 0.073682 |
| type IV pilus signal transduction protein PilI | 1.046911 | 1.174911 | 0.166412 | 0.073821 |
| glyoxalase I | 1.244735 | 0.883677 | -0.49425 | 0.07384 |
| Arc domain protein DNA binding domain protein | 0.925403 | 1.021202 | 0.142115 | 0.073862 |
| transcriptional regulator, LysR family...222 | 1.060909 | 0.962194 | -0.1409 | 0.073998 |
| hypothetical protein ABUW_3068 | 0.632069 | 0.51163 | -0.30498 | 0.074127 |
| two-component system response regulator protein...1811 | 1.004377 | 1.042851 | 0.054232 | 0.074296 |
| ribosomal protein L6 | 1.101186 | 0.950617 | -0.21212 | 0.074403 |
| chorismate mutase...711 | 1.130213 | 0.951103 | -0.24892 | 0.074442 |
| peptidoglycan-binding LysM...2472 | 1.04872 | 0.916144 | -0.19498 | 0.074518 |
| hypothetical protein ABUW_1606 | 1.034058 | 1.082368 | 0.065873 | 0.07471 |
| OmpA/MotB domain protein...556 | 0.869968 | 1.102505 | 0.341752 | 0.075068 |
| lipid A phosphoethanolamine transferase | 1.107891 | 0.985066 | -0.16952 | 0.075287 |
| peptide chain release factor 2 | 0.907948 | 1.092816 | 0.267368 | 0.075521 |
| ribosomal protein L4/L1e | 1.095656 | 0.953243 | -0.20088 | 0.075706 |
| protein YaaA | 1.073743 | 0.917383 | -0.22705 | 0.075763 |

| | | | | |
|---|---|---|---|---|
| hypothetical protein ABUW_2637 | 0.973612 | 1.082289 | 0.152666 | 0.076089 |
| hypothetical protein ABUW_0769 | 0.970591 | 1.031793 | 0.088217 | 0.076136 |
| outer membrane protein E | 0.896641 | 1.103786 | 0.299858 | 0.076345 |
| 3-oxoacyl-[acyl-carrier-protein] reductase...964 | 1.28211 | 0.758176 | -0.75792 | 0.076346 |
| transport protein | 1.057782 | 1.002377 | -0.07762 | 0.076485 |
| hypothetical protein ABUW_0554 | 0.94198 | 1.261622 | 0.421511 | 0.076491 |
| hypothetical protein ABUW_3839 | 0.892337 | 1.214584 | 0.444801 | 0.076524 |
| hypothetical protein ABUW_0934 | 1.173844 | 0.894411 | -0.39223 | 0.077023 |
| hypothetical protein ABUW_0373 | 1.016693 | 0.950727 | -0.09678 | 0.07719 |
| hypothetical protein ABUW_3756 | 0.97183 | 0.856129 | -0.18288 | 0.077262 |
| transcriptional regulator, GntR family...1077 | 1.016457 | 0.907884 | -0.16297 | 0.077551 |
| hypothetical protein ABUW_0607 | 1.11394 | 0.951829 | -0.2269 | 0.077955 |
| aldehyde dehydrogenase...2209 | 1.073605 | 0.916285 | -0.2286 | 0.078184 |
| 3-oxoadipate CoA-transferase subunit B...1258 | 1.003727 | 1.129103 | 0.16981 | 0.078284 |
| hypothetical protein ABUW_3496 | 0.988759 | 0.911564 | -0.11727 | 0.078354 |
| UDP-glucose 4-epimerase...2351 | 0.995481 | 0.891845 | -0.1586 | 0.078401 |
| putative DNA helicase | 0.946348 | 1.044072 | 0.141779 | 0.078596 |
| ribosomal protein L34 | 1.481878 | 0.614819 | -1.26919 | 0.078647 |
| entericidin EcnAB | 1.085927 | 0.826692 | -0.3935 | 0.07911 |
| aldehyde dehydrogenase...311 | 1.186816 | 0.805181 | -0.55971 | 0.079905 |
| (di)nucleoside polyphosphate hydrolase | 1.00945 | 0.941301 | -0.10084 | 0.080245 |
| hypothetical protein ABUW_2699 | 0.997783 | 0.89512 | -0.15664 | 0.080764 |
| shikimate 5-dehydrogenase | 0.973275 | 1.061983 | 0.125841 | 0.080797 |
| hypothetical protein ABUW_0344 | 0.996759 | 1.07269 | 0.105916 | 0.080926 |
| hypothetical protein ABUW_0612 | 1.099798 | 0.949851 | -0.21146 | 0.080998 |
| peptidase S49 | 0.91008 | 1.0562 | 0.214818 | 0.081097 |
| malonate decarboxylase, epsilon subunit | 0.987018 | 0.668729 | -0.56165 | 0.081537 |
| homoserine dehydrogenase...804 | 0.989753 | 1.064696 | 0.1053 | 0.08202 |
| 2-oxoglutarate dehydrogenase, E1 component | 0.966809 | 1.059223 | 0.131703 | 0.082179 |
| putative arsenate reductase | 1.190941 | 0.894101 | -0.41359 | 0.082366 |
| ring hydroxylating dioxygenase, Rieske | 0.945293 | 1.043608 | 0.142745 | 0.08264 |
| hypothetical protein ABUW_2906 | 0.92269 | 0.75574 | -0.28796 | 0.082829 |
| aldehyde dehydrogenase...1362 | 0.960541 | 1.061597 | 0.144317 | 0.083097 |
| hypothetical protein ABUW_2729 | 0.955761 | 1.025754 | 0.101963 | 0.08311 |
| acetate--CoA ligase | 1.039109 | 0.945959 | -0.1355 | 0.08313 |
| hypothetical protein ABUW_1437 | 0.847314 | 0.999684 | 0.238575 | 0.083485 |
| ribosomal protein L15 | 1.181659 | 0.888672 | -0.41109 | 0.083527 |
| phosphatase domain-containing protein | 0.855965 | 1.038991 | 0.279561 | 0.083547 |
| L-aspartate oxidase | 1.005136 | 0.966332 | -0.0568 | 0.083754 |
| phosphoglycolate phosphatase, bacterial | 0.894121 | 1.107028 | 0.30815 | 0.083765 |
| biofilm-associated protein...2215 | 0.946417 | 0.818027 | -0.21033 | 0.083973 |
| transcriptional regulator, TetR family...2014 | 1.135837 | 0.955379 | -0.24961 | 0.084389 |
| hypothetical protein ABUW_3804 | 0.891107 | 1.012195 | 0.183817 | 0.084434 |
| thiol:disulfide interchange protein DsbA | 0.995114 | 0.942246 | -0.07876 | 0.08446 |
| lipolytic enzyme | 1.075474 | 0.914712 | -0.23358 | 0.084585 |
| rieske 2Fe-2S family protein | 1.078728 | 0.857158 | -0.3317 | 0.084854 |
| hypothetical protein ABUW_1659 | 1.371443 | 0.78746 | -0.80042 | 0.084943 |

| | | | | |
|---|---|---|---|---|
| GTP-binding protein...896 | 0.941001 | 1.015432 | 0.109825 | 0.085238 |
| monofunctional biosynthetic peptidoglycan transglycosylase | 0.908477 | 1.090522 | 0.263497 | 0.08537 |
| NADH dehydrogenase I chain L | 0.920285 | 1.002093 | 0.122863 | 0.085485 |
| YaeQ protein | 0.893864 | 1.039291 | 0.217472 | 0.085509 |
| copper resistance protein B...1710 | 1.079056 | 0.876487 | -0.29996 | 0.086315 |
| glutathione peroxidase...2311 | 1.097276 | 0.903309 | -0.28064 | 0.086323 |
| tRNA pseudouridine synthase B | 0.948645 | 1.072031 | 0.176407 | 0.08633 |
| transcriptional regulator, LysR family...911 | 0.872815 | 1.078018 | 0.304634 | 0.086388 |
| electron transport complex, rnfaBcdge type, B subunit | 0.982793 | 1.143677 | 0.21872 | 0.086491 |
| efflux ABC transporter, permease protein | 1.041701 | 0.926105 | -0.16969 | 0.086579 |
| phage protein...602 | 1.034247 | 0.697864 | -0.56756 | 0.086654 |
| hypothetical protein ABUW_3760 | 0.964583 | 1.025012 | 0.087664 | 0.086706 |
| amidohydrolase...1413 | 1.044113 | 0.985718 | -0.08303 | 0.086727 |
| O-acetylhomoserine/O-acetylserine sulfhydrylase | 0.842268 | 1.021657 | 0.27856 | 0.086816 |
| LemA | 1.110973 | 0.818673 | -0.44047 | 0.086988 |
| tRNA nucleotidyl transferase | 0.955851 | 1.039582 | 0.121146 | 0.086998 |
| hypothetical protein ABUW_2724 | 1.310812 | 0.731776 | -0.84099 | 0.087056 |
| DNA replication protein (plasmid)...1232 | 0.973359 | 0.935603 | -0.05708 | 0.08727 |
| hypothetical protein ABUW_1287 | 1.107292 | 0.89498 | -0.30711 | 0.087335 |
| 3-dehydroshikimate dehydratase | 1.265001 | 1.022099 | -0.3076 | 0.087467 |
| hypothetical protein ABUW_1427 | 0.802614 | 0.986688 | 0.297888 | 0.087601 |
| 3-isopropylmalate dehydratase, large subunit | 1.007982 | 0.931617 | -0.11366 | 0.087645 |
| hypothetical protein ABUW_2674 | 1.201345 | 0.706751 | -0.76538 | 0.087925 |
| hypothetical protein ABUW_0839 | 1.07818 | 0.921095 | -0.22718 | 0.087926 |
| aspartyl-tRNA synthetase | 0.996909 | 1.018713 | 0.031214 | 0.08815 |
| enoyl-CoA hydratase/isomerase family protein | 0.985319 | 1.016726 | 0.045268 | 0.08823 |
| recombination protein RecR | 0.960395 | 0.929512 | -0.04716 | 0.088478 |
| general secretory pathway protein E | 1.186145 | 0.976059 | -0.28124 | 0.088571 |
| CDP-diacylglycerol--glycerol-3-phosphate 3-phosphatidyltrans | 0.703656 | 1.075221 | 0.611691 | 0.088731 |
| OsmC family protein...1516 | 1.047345 | 1.009367 | -0.05329 | 0.089049 |
| carbohydrate kinase family | 0.948355 | 0.997767 | 0.073275 | 0.089186 |
| indole-3-pyruvate decarboxylase | 0.942092 | 1.041114 | 0.144189 | 0.089259 |
| phosphoadenosine phosphosulfate reductase | 1.073152 | 0.972749 | -0.14171 | 0.089464 |
| glutaryl-CoA dehydrogenase | 0.937998 | 1.028802 | 0.133309 | 0.089517 |
| porin B...946 | 1.025105 | 0.939445 | -0.12589 | 0.089674 |
| cell division topological specificity factor MinE | 1.094413 | 0.930481 | -0.23411 | 0.089874 |
| ribonuclease T | 0.96639 | 1.223795 | 0.340685 | 0.090036 |
| enoyl-coA hydratase | 0.905328 | 1.046619 | 0.209224 | 0.090201 |
| lysyl-tRNA synthetase...1441 | 0.971087 | 1.018453 | 0.068708 | 0.090255 |
| phosphoribosylglycinamide formyltransferase | 1.044287 | 1.147633 | 0.136143 | 0.090285 |
| methyltransferase type 11 | 0.967339 | 1.04549 | 0.112087 | 0.090372 |
| cyclic nucleotide-binding domain protein | 0.989726 | 1.019359 | 0.042561 | 0.090866 |
| hypothetical protein ABUW_2244 | 0.898074 | 0.991954 | 0.14344 | 0.091249 |
| hypothetical protein ABUW_3141 | 1.207923 | 0.7715 | -0.64679 | 0.091337 |
| acetyl-CoA carboxylase, biotin carboxyl carrier protein | 1.157332 | 0.976018 | -0.24582 | 0.091554 |
| hypothetical protein ABUW_1059 | 0.969198 | 1.155877 | 0.254125 | 0.091581 |
| pantothenate kinase, type III | 0.972805 | 0.909605 | -0.09691 | 0.091625 |

| | | | | |
|---|---|---|---|---|
| universal stress protein family...2294 | 1.057441 | 0.922564 | -0.19686 | 0.091865 |
| pH adaptation potassium efflux system transmembrane prote | 0.940965 | 1.095019 | 0.218743 | 0.092038 |
| cell division protein FtsH | 0.950882 | 1.04012 | 0.129412 | 0.092051 |
| hypothetical protein ABUW_0308 | 0.979867 | 1.06922 | 0.1259 | 0.092105 |
| hypothetical protein ABUW_1358 | 0.935164 | 1.045706 | 0.161185 | 0.093004 |
| hypothetical protein ABUW_2853 | 0.997052 | 0.89373 | -0.15783 | 0.093527 |
| hypothetical protein ABUW_2703 | 1.107037 | 0.942506 | -0.23213 | 0.093654 |
| allophanate hydrolase | 0.977223 | 1.033515 | 0.0808 | 0.093675 |
| hypothetical protein ABUW_2783 | 1.126003 | 0.816195 | -0.46423 | 0.093852 |
| peptidase M23B | 1.017701 | 0.948878 | -0.10102 | 0.094139 |
| D-serine/D-alanine/glycine transporter...676 | 1.197565 | 0.895199 | -0.41982 | 0.094212 |
| hypothetical protein ABUW_1314 | 0.932234 | 1.054196 | 0.177379 | 0.094304 |
| fatty acid desaturase...1337 | 1.026666 | 0.934062 | -0.13638 | 0.094337 |
| enoyl-CoA hydratase...2343 | 0.916578 | 1.040217 | 0.182555 | 0.094389 |
| thioesterase family protein | 1.126023 | 0.866563 | -0.37786 | 0.094488 |
| transcriptional regulator, LysR family...199 | 1.062521 | 1.653258 | 0.637821 | 0.094515 |
| radical SAM domain protein | 1.017753 | 0.952702 | -0.09529 | 0.09463 |
| coproporphyrinogen III oxidase, aerobic | 0.968262 | 1.034985 | 0.09614 | 0.094742 |
| ribosomal protein L22 | 1.101845 | 0.935984 | -0.23536 | 0.094807 |
| hypothetical protein ABUW_0449 | 1.005605 | 1.090928 | 0.117493 | 0.095367 |
| UDP-N-acetylmuramyl-tripeptide synthetase | 0.951762 | 0.989536 | 0.056151 | 0.095424 |
| histidine utilization repressor | 0.94382 | 1.043696 | 0.145117 | 0.09568 |
| khg/kdpg aldolase | 0.973483 | 1.055464 | 0.11665 | 0.096082 |
| senescence marker protein-30 | 1.069104 | 0.929524 | -0.20184 | 0.096446 |
| protein TolQ | 0.964337 | 0.870545 | -0.14762 | 0.096545 |
| hypothetical protein ABUW_3001 | 0.89149 | 1.017727 | 0.191061 | 0.096618 |
| KGG domain-containing protein | 0.99181 | 0.942605 | -0.07341 | 0.096989 |
| monooxygenase, NtaA/SnaA/SoxA family | 1.084861 | 0.956355 | -0.18189 | 0.097746 |
| putative VGR-related protein | 0.911683 | 0.981919 | 0.107071 | 0.098134 |
| hypothetical protein ABUW_0812 | 1.118552 | 0.974549 | -0.19883 | 0.098506 |
| hypothetical protein ABUW_3882 | 1.143126 | 0.880518 | -0.37656 | 0.09931 |
| exopolyphosphatase | 0.955645 | 1.012164 | 0.082896 | 0.099677 |
| UDP-N-acetylmuramate:L-alanyl-gamma-D-glutamyl-meso-dia | 0.963983 | 1.058955 | 0.135562 | 0.09969 |
| hypothetical protein ABUW_0263 | 0.991089 | 1.066637 | 0.105984 | 0.099875 |
| septum formation initiator | 1.173882 | 1.300808 | 0.14812 | 0.100191 |
| acetyl-CoA acetyltransferase...2094 | 0.986896 | 1.034738 | 0.068295 | 0.100455 |
| hypothetical protein ABUW_0021 | 1.03923 | 0.942365 | -0.14116 | 0.100749 |
| transcriptional regulator, TetR family...1149 | 1.055953 | 1.100494 | 0.059606 | 0.10084 |
| AAA ATPase family protein | 1.080142 | 0.945964 | -0.19136 | 0.100856 |
| hypothetical protein ABUW_0297 | 0.936368 | 1.009501 | 0.108495 | 0.101105 |
| hypothetical protein ABUW_0183 | 0.942378 | 1.045321 | 0.149568 | 0.101221 |
| rhodanese domain protein...1847 | 1.152369 | 0.920188 | -0.3246 | 0.101558 |
| glycerol kinase | 1.033485 | 0.967314 | -0.09546 | 0.102144 |
| multidrug resistance protein VceB | 0.992483 | 1.111774 | 0.16375 | 0.102279 |
| protocatechuate 3,4-dioxygenase, beta subunit | 0.949369 | 1.018673 | 0.101651 | 0.102451 |
| glycosyltransferase...1269 | 0.910091 | 0.978794 | 0.104994 | 0.103164 |
| hypothetical protein ABUW_2493 | 1.383999 | 0.940762 | -0.55694 | 0.103271 |

| | | | | |
|---|---|---|---|---|
| shikimate kinase...825 | 0.926907 | 1.06628 | 0.20209 | 0.103652 |
| LmbE-like protein | 1.100447 | 0.955802 | -0.20331 | 0.103676 |
| bifunctional succinylornithine transaminase/acetylornithine tı | 1.018953 | 0.944048 | -0.11016 | 0.103968 |
| ribosomal protein L16 | 1.14352 | 0.857974 | -0.41447 | 0.104036 |
| acyl carrier protein...2241 | 0.944701 | 1.078997 | 0.191761 | 0.104197 |
| TonB-dependent receptor...1322 | 1.017897 | 0.938556 | -0.11708 | 0.104763 |
| glycyl-tRNA synthetase, beta subunit | 1.014842 | 0.960636 | -0.07919 | 0.104777 |
| fmn-dependent NADH-azoreductase | 1.22617 | 0.874593 | -0.48747 | 0.1048 |
| cytochrome D ubiquinol oxidase subunit I | 0.813384 | 1.012721 | 0.316228 | 0.105036 |
| UDP-N-acetylglucosamine 2-epimerase...2004 | 1.019934 | 0.962768 | -0.08322 | 0.105126 |
| P-aminobenzoate synthetase PabB | 1.197987 | 0.859232 | -0.47949 | 0.105444 |
| prolyl-tRNA synthetase | 0.958513 | 1.020854 | 0.090905 | 0.105514 |
| DNA repair protein RadA | 0.924257 | 1.052075 | 0.186871 | 0.106079 |
| histidine acid phosphatase family protein | 1.063796 | 0.968903 | -0.1348 | 0.106258 |
| ribosomal protein L20 | 1.090418 | 0.898515 | -0.27927 | 0.106349 |
| hypothetical protein ABUW_0746 | 1.078889 | 1.014576 | -0.08867 | 0.106539 |
| tRNA delta(2)-isopentenylpyrophosphate transferase | 0.927907 | 1.03717 | 0.160599 | 0.10699 |
| transcriptional regulator, TetR family...971 | 1.029084 | 0.976281 | -0.07599 | 0.107215 |
| 5'-nucleotidase | 0.990889 | 1.047127 | 0.079641 | 0.107259 |
| ribosomal protein S9 | 1.110847 | 0.954892 | -0.21825 | 0.107587 |
| shikimate kinase...820 | 0.97762 | 1.074842 | 0.136778 | 0.107958 |
| FilD | 0.804554 | 1.111706 | 0.466514 | 0.108179 |
| sensory box protein | 1.008824 | 1.123971 | 0.155931 | 0.108372 |
| DNA repair protein RecN | 0.994818 | 0.947381 | -0.07049 | 0.108719 |
| ribosomal protein S20 | 1.100139 | 0.939812 | -0.22724 | 0.108732 |
| cytochrome B561...551 | 1.086154 | 1.036507 | -0.0675 | 0.109063 |
| hypothetical protein ABUW_2845 | 1.04709 | 0.968133 | -0.11311 | 0.109173 |
| thiamine pyrophosphate enzyme domain protein TPP-binding | 1.021176 | 0.936901 | -0.12426 | 0.109454 |
| ribosomal protein L23 | 1.090241 | 0.994519 | -0.13258 | 0.109585 |
| ribosomal protein L10 | 1.05832 | 0.988695 | -0.09818 | 0.109759 |
| sulfurtransferase TusA | 1.006619 | 1.097598 | 0.124832 | 0.110143 |
| excinuclease ABC, A subunit | 0.958797 | 1.034289 | 0.109343 | 0.110412 |
| hypothetical protein ABUW_3005 | 0.940214 | 1.027793 | 0.128487 | 0.110431 |
| hypothetical protein ABUW_2142 | 1.082904 | 1.139304 | 0.073248 | 0.110881 |
| putative RND family drug transporter...1993 | 1.036204 | 0.936387 | -0.14613 | 0.111199 |
| tannase/feruloyl esterase family protein | 1.106188 | 0.83726 | -0.40185 | 0.111388 |
| redox-sensitive transcriptional activator SoxR | 0.946656 | 0.896552 | -0.07845 | 0.111468 |
| hypothetical protein ABUW_2156 | 0.887137 | 1.013183 | 0.191665 | 0.111658 |
| large conductance mechanosensitive channel protein | 1.131607 | 0.880647 | -0.36174 | 0.111897 |
| biotin biosynthesis protein BioC | 1.282987 | 1.016072 | -0.3365 | 0.111903 |
| iron transporter | 1.00264 | 1.062728 | 0.083968 | 0.111975 |
| 3-oxoadipate CoA-transferase, subunit B | 1.046474 | 0.915607 | -0.19274 | 0.112033 |
| hypothetical protein ABUW_3348 | 1.037098 | 0.969375 | -0.09743 | 0.112089 |
| dephospho-CoA kinase | 0.958325 | 1.010067 | 0.075864 | 0.112132 |
| ribosomal protein L7/L12 | 1.052941 | 0.978657 | -0.10555 | 0.112211 |
| TonB-dependent siderophore receptor...1261 | 0.968629 | 1.028702 | 0.086809 | 0.112354 |
| thioesterase superfamily protein...750 | 0.98224 | 1.082006 | 0.13956 | 0.112491 |

| | | | | |
|---|---|---|---|---|
| transcriptional regulator, ArsR family...1206 | 1.045238 | 0.933927 | -0.16245 | 0.112654 |
| transcriptional regulator, GntR family...343 | 0.897548 | 1.142676 | 0.348356 | 0.112913 |
| alpha/beta hydrolase fold protein EstB | 0.899062 | 0.9868 | 0.134336 | 0.112939 |
| enoyl-CoA hydratase...223 | 0.726258 | 1.379455 | 0.925544 | 0.112961 |
| hypothetical protein ABUW_1192 | 1.08434 | 0.804822 | -0.43007 | 0.113036 |
| cytochrome B561...383 | 1.001602 | 0.892056 | -0.1671 | 0.113182 |
| transcriptional regulator, LysR family...209 | 1.019288 | 1.123275 | 0.140149 | 0.113306 |
| DNA-binding protein H-NS | 1.0647 | 0.984944 | -0.11233 | 0.11407 |
| glycoprotease | 0.973739 | 1.060497 | 0.123134 | 0.114215 |
| inner membrane protein OxaA | 1.08513 | 0.930598 | -0.22164 | 0.114246 |
| hypothetical protein ABUW_3489 | 0.952062 | 1.040779 | 0.128536 | 0.114323 |
| transporter, major facilitator family | 0.963128 | 0.766684 | -0.3291 | 0.114527 |
| phage small terminase subunit | 0.829788 | 0.964812 | 0.217505 | 0.114641 |
| hypothetical protein ABUW_0840 | 1.101468 | 0.904855 | -0.28367 | 0.115126 |
| arsenate reductase...113 | 0.728609 | 1.240362 | 0.767545 | 0.115238 |
| methylcrotonoyl-CoA carboxylase subunit alpha | 1.048185 | 0.998354 | -0.07027 | 0.115861 |
| 5'-methylthioadenosine/S-adenosylhomocysteine nucleosidas | 1.017912 | 0.841909 | -0.27388 | 0.117222 |
| ExsB protein | 1.224224 | 0.816734 | -0.58393 | 0.117419 |
| hypothetical protein ABUW_0233 | 1.11878 | 0.878185 | -0.34933 | 0.118298 |
| hypothetical protein ABUW_0709 | 0.895801 | 0.975307 | 0.122678 | 0.118425 |
| glutamate synthase, large subunit | 0.984085 | 1.032395 | 0.06914 | 0.11857 |
| ribosomal protein L33 | 1.062882 | 1.001935 | -0.08519 | 0.119833 |
| phosphoribosylformimino-5-aminoimidazole carboxamide rib | 0.985224 | 1.015233 | 0.043288 | 0.12001 |
| thiol:disulfide interchange protein...225 | 0.888623 | 1.02621 | 0.207683 | 0.120087 |
| oxidoreductase, short chain dehydrogenase/reductase family | 0.891955 | 1.030602 | 0.208444 | 0.120722 |
| methyltransferase | 0.956924 | 0.988278 | 0.046512 | 0.121085 |
| ribosomal protein L18 | 1.045129 | 0.978881 | -0.09448 | 0.12143 |
| small GTP-binding protein | 0.970729 | 1.026383 | 0.080428 | 0.121479 |
| acyl-CoA dehydrogenase...2410 | 1.075413 | 0.969508 | -0.14957 | 0.12161 |
| NAD(P)H dehydrogenase, quinone family | 0.801685 | 1.173111 | 0.549232 | 0.121683 |
| ribosomal protein L13 | 1.149074 | 0.88756 | -0.37256 | 0.122111 |
| O-methyl transferase | 1.012437 | 0.978315 | -0.04946 | 0.122223 |
| transcriptional regulator, BadM/Rrf2 family | 1.053924 | 0.954023 | -0.14367 | 0.122254 |
| transcriptional regulator, IclR family...1332 | 1.042989 | 1.002317 | -0.05738 | 0.122333 |
| ribosomal protein S10 | 1.051028 | 1.005268 | -0.06422 | 0.122583 |
| phosphoglycolate phosphatase 2 | 0.963986 | 0.88731 | -0.11957 | 0.123262 |
| putative ATP binding site | 0.999845 | 0.887348 | -0.1722 | 0.123329 |
| ribosomal protein L11 | 1.091607 | 0.974566 | -0.16362 | 0.123535 |
| zinc-binding dehydrogenase | 1.042489 | 0.956133 | -0.12475 | 0.123537 |
| ribosomal protein S17 | 1.189543 | 0.862002 | -0.46465 | 0.123744 |
| hypothetical protein ABUW_3059 | 0.927316 | 0.830467 | -0.15914 | 0.123827 |
| TonB-dependent siderophore receptor...1175 | 1.053456 | 1.134479 | 0.1069 | 0.123886 |
| sulfur relay protein TusD/DsrE | 0.956984 | 0.87716 | -0.12565 | 0.123903 |
| phospholipid/glycerol acyltransferase...1189 | 1.077021 | 1.002935 | -0.10282 | 0.12392 |
| putative peroxidase | 1.004526 | 0.89444 | -0.16746 | 0.124018 |
| translation initiation factor IF-1 | 1.174549 | 0.938252 | -0.32406 | 0.124161 |
| DNA-binding protein HU | 1.054303 | 0.954381 | -0.14365 | 0.124229 |

| | | | | |
|---|---|---|---|---|
| NADH dehydrogenase I chain F | 0.912504 | 1.004192 | 0.138133 | 0.124268 |
| hydroxymethylglutaryl-CoA lyase | 0.985358 | 1.031436 | 0.065935 | 0.124648 |
| phenazine biosynthesis protein PhzF family...1423 | 1.0624 | 0.960946 | -0.1448 | 0.124652 |
| ribosomal protein S13 | 1.11322 | 0.909528 | -0.29155 | 0.124985 |
| aldehyde dehydrogenase...672 | 1.011653 | 0.909593 | -0.15342 | 0.12501 |
| DNA-directed RNA polymerase, beta' subunit | 1.016678 | 0.969241 | -0.06893 | 0.125144 |
| hypothetical protein ABUW_1399 | 1.007284 | 1.0589 | 0.072096 | 0.125227 |
| hypothetical protein ABUW_1511 | 1.296679 | 0.786085 | -0.72206 | 0.125387 |
| transcriptional regulator, LysR family...587 | 1.117299 | 0.939991 | -0.2493 | 0.12547 |
| hypothetical protein ABUW_3435 | 0.85907 | 1.280033 | 0.575332 | 0.12549 |
| threonine ammonia-lyase | 1.005624 | 0.984263 | -0.03098 | 0.125556 |
| glutamyl-tRNA synthetase...433 | 1.104957 | 0.766914 | -0.52685 | 0.125592 |
| ADP-ribose pyrophosphatase | 1.056479 | 0.895316 | -0.2388 | 0.125631 |
| aminoglycoside phosphotransferase...1302 | 0.98525 | 1.095188 | 0.152617 | 0.125722 |
| tetratricopeptide repeat domain protein | 1.042208 | 0.922806 | -0.17554 | 0.126444 |
| hypothetical protein ABUW_0630 | 1.141041 | 0.675442 | -0.75645 | 0.12658 |
| hypothetical protein ABUW_2612 | 1.004413 | 1.050355 | 0.064524 | 0.126696 |
| hypothetical protein ABUW_0798 | 1.019138 | 0.89583 | -0.18605 | 0.126797 |
| quinolinate synthetase complex, A subunit | 0.978801 | 1.0356 | 0.081379 | 0.126858 |
| hypothetical protein ABUW_2463 | 1.059578 | 0.923427 | -0.19842 | 0.127226 |
| SsrA-binding protein | 0.961353 | 1.027411 | 0.095875 | 0.127415 |
| transcriptional regulator, TetR family...675 | 1.033362 | 0.899995 | -0.19936 | 0.127423 |
| 4-hydroxybenzoate transporter | 0.843624 | 0.999525 | 0.244642 | 0.127948 |
| glucose-6-phosphate isomerase | 0.981079 | 0.955605 | -0.03795 | 0.128129 |
| hydrolase...287 | 1.003137 | 1.096429 | 0.128293 | 0.128168 |
| fumarate lyase | 1.147777 | 1.037127 | -0.14625 | 0.128208 |
| major facilitator superfamily MFS_1...550 | 0.861627 | 1.125141 | 0.384971 | 0.128244 |
| hypothetical protein ABUW_2290 | 1.073909 | 0.841698 | -0.3515 | 0.128844 |
| undecaprenyldiphospho-muramoylpentapeptide beta-N-acet | 0.953285 | 1.001437 | 0.071092 | 0.12923 |
| stringent starvation protein B | 0.87022 | 1.038099 | 0.254493 | 0.12925 |
| deoxycytidine triphosphate deaminase | 0.953422 | 1.068857 | 0.164882 | 0.129485 |
| FilF | 1.001741 | 0.945777 | -0.08294 | 0.129722 |
| hypothetical protein ABUW_3631 | 1.16466 | 0.955584 | -0.28545 | 0.129774 |
| ribosomal protein L32 | 1.061148 | 0.87831 | -0.27282 | 0.129828 |
| hypothetical protein ABUW_3658 | 0.918846 | 0.96765 | 0.074663 | 0.129851 |
| helicase domain-containing terminase-like protein | 1.139249 | 0.994685 | -0.19577 | 0.130025 |
| hypothetical protein ABUW_2670 | 1.05514 | 0.864434 | -0.28761 | 0.13003 |
| hypothetical protein ABUW_0139 | 1.03442 | 0.934048 | -0.14725 | 0.130073 |
| transcriptional regulator, TetR family...677 | 0.933391 | 1.084979 | 0.217114 | 0.13036 |
| sodium-and chloride-dependent transporter | 1.035709 | 1.00114 | -0.04898 | 0.130435 |
| hypothetical protein ABUW_2272 | 0.93433 | 1.010342 | 0.112839 | 0.130666 |
| Sec-independent protein translocase protein | 1.40014 | 0.842695 | -0.73249 | 0.13097 |
| TonB-dependent receptor...1528 | 0.974411 | 1.070099 | 0.135143 | 0.131311 |
| short chain dehydrogenase...1109 | 1.031137 | 0.922336 | -0.16087 | 0.13153 |
| transcriptional regulator, AraC family...681 | 0.947961 | 1.025261 | 0.113091 | 0.131575 |
| hypothetical protein ABUW_0507 | 1.017865 | 0.938723 | -0.11677 | 0.131836 |
| RNA methyltransferase, TrmH family, group 1 | 0.937436 | 0.966527 | 0.044089 | 0.132008 |

| | | | | |
|---|---|---|---|---|
| D-tyrosyl-tRNA(Tyr) deacylase | 0.950902 | 1.125035 | 0.2426 | 0.132235 |
| universal stress protein | 1.029707 | 0.957626 | -0.1047 | 0.132733 |
| two-component system sensor kinase protein | 0.923907 | 1.011615 | 0.130841 | 0.133277 |
| ABC transporter, ATP-binding protein...963 | 0.985267 | 1.062428 | 0.108778 | 0.133361 |
| acetyl-coenzyme A synthetase | 1.045478 | 0.999239 | -0.06526 | 0.133453 |
| site-specific recombinase, phage integrase family | 0.941077 | 1.082437 | 0.201898 | 0.13351 |
| hypothetical protein ABUW_0102 | 1.045752 | 0.950835 | -0.13727 | 0.133681 |
| Sua5/YciO/YrdC/YwlC family protein | 1.051805 | 0.987271 | -0.09135 | 0.133768 |
| hypothetical protein ABUW_0523 | 0.990181 | 1.094961 | 0.145115 | 0.134348 |
| hypothetical protein ABUW_3006 | 1.086656 | 1.14794 | 0.079152 | 0.13452 |
| general secretion pathway protein D | 0.943749 | 1.082569 | 0.197984 | 0.134767 |
| transcriptional regulator, MerR family | 0.898801 | 1.057684 | 0.234835 | 0.134914 |
| glycyl-tRNA synthetase, alpha subunit | 0.99687 | 0.977099 | -0.0289 | 0.135128 |
| 1-deoxy-D-xylulose-5-phosphate synthase | 0.978389 | 1.046976 | 0.097748 | 0.135166 |
| vitamin B12 receptor | 0.967024 | 1.049409 | 0.117955 | 0.135363 |
| hypothetical protein ABUW_2997 | 0.935652 | 0.997957 | 0.093005 | 0.135544 |
| ATPase, AAA family | 1.019447 | 0.984592 | -0.05019 | 0.135559 |
| pyruvate dehydrogenase complex dihydrolipoamide acetyltra | 1.041071 | 0.975214 | -0.09428 | 0.135759 |
| Na+/solute symporter | 1.00948 | 1.097027 | 0.119987 | 0.135835 |
| biotin-[acetyl-CoA-carboxylase] ligase | 0.957259 | 1.069776 | 0.160328 | 0.136426 |
| AcnD-accessory protein PrpF | 1.00849 | 1.023095 | 0.020743 | 0.136867 |
| ribosomal protein S5 | 1.060063 | 1.005838 | -0.07575 | 0.137193 |
| transcriptional regulator...16 | 0.991821 | 1.29046 | 0.379734 | 0.137223 |
| DNA mismatch repair protein MutL | 0.958479 | 0.989517 | 0.045978 | 0.137259 |
| MaoC domain protein dehydratase | 1.032229 | 0.959537 | -0.10535 | 0.138629 |
| DSBA oxidoreductase | 0.984118 | 0.915188 | -0.10476 | 0.138714 |
| adenylate cyclase | 1.022068 | 1.061197 | 0.054201 | 0.13907 |
| competence/damage-inducible protein CinA...180 | 1.029205 | 0.916109 | -0.16794 | 0.139594 |
| exodeoxyribonuclease III | 1.01475 | 0.991002 | -0.03416 | 0.13982 |
| BolA family protein | 1.140101 | 0.977815 | -0.22153 | 0.140287 |
| hypothetical protein ABUW_0217 | 0.983336 | 0.894562 | -0.1365 | 0.140291 |
| protein tyrosine phosphatase | 0.974197 | 1.107977 | 0.185643 | 0.140378 |
| hypothetical protein ABUW_0135 | 1.000569 | 1.063266 | 0.087683 | 0.140687 |
| hypothetical protein ABUW_3687 | 0.900693 | 1.019713 | 0.179055 | 0.140748 |
| quinoprotein glucose dehydrogenase-B...2242 | 0.929492 | 1.025892 | 0.142365 | 0.141332 |
| lipoprotein-releasing system ATP-binding protein LolD | 0.998703 | 1.03809 | 0.055804 | 0.141339 |
| septum formation protein Maf | 0.983433 | 1.067973 | 0.118976 | 0.142013 |
| Glu-tRNA amidotransferase | 1.033175 | 0.975365 | -0.08307 | 0.142042 |
| ribosomal protein L21 | 1.076242 | 0.956748 | -0.16979 | 0.142095 |
| protein secretion ABC efflux system, permease and ATP-bindi | 0.978095 | 0.934937 | -0.06511 | 0.142371 |
| protein-export membrane protein SecF | 0.985491 | 0.926623 | -0.08886 | 0.142692 |
| ribosomal protein L24 | 1.095683 | 0.967856 | -0.17897 | 0.142878 |
| hypothetical protein ABUW_1960 | 1.093481 | 1.007666 | -0.11791 | 0.143373 |
| methionine biosynthesis protein MetW | 0.960246 | 0.997145 | 0.054398 | 0.143454 |
| hypothetical protein ABUW_0468 | 1.096386 | 0.925689 | -0.24416 | 0.143526 |
| hypothetical protein ABUW_0271 | 0.883851 | 1.010702 | 0.193483 | 0.143719 |
| thioredoxin...1904 | 1.141112 | 0.951249 | -0.26254 | 0.14405 |

| | | | | |
|---|---|---|---|---|
| ribosomal protein L1 | 1.057948 | 0.988321 | -0.09822 | 0.144251 |
| hypothetical protein ABUW_2595 | 1.071025 | 0.956441 | -0.16324 | 0.144277 |
| hypothetical protein ABUW_0914 | 0.925666 | 0.972137 | 0.070667 | 0.144445 |
| hypothetical protein ABUW_2624 | 1.332384 | 0.979166 | -0.44438 | 0.14455 |
| hypothetical protein ABUW_0897 | 1.046719 | 0.92683 | -0.1755 | 0.144787 |
| hypothetical protein ABUW_0225 | 0.792954 | 0.903129 | 0.187694 | 0.144994 |
| acyl-CoA dehydrogenase...2373 | 1.067868 | 0.941783 | -0.18127 | 0.145401 |
| sulfate adenylyltransferase subunit 2 | 1.001324 | 1.033623 | 0.045801 | 0.14813 |
| hypothetical protein ABUW_2862 | 0.894781 | 1.010069 | 0.174847 | 0.148206 |
| 2-nitropropane dioxygenase...1406 | 1.117793 | 0.937237 | -0.25417 | 0.148266 |
| 3-oxoadipate CoA-transferase subunit A...634 | 1.385965 | 0.71475 | -0.95538 | 0.148616 |
| ribosomal protein L30 | 1.047051 | 0.998816 | -0.06804 | 0.1488 |
| transcriptional regulator, LysR family...654 | 0.850983 | 1.171784 | 0.461505 | 0.148802 |
| glutamyl-tRNA(Gln) amidotransferase, C subunit | 1.186941 | 0.916894 | -0.37242 | 0.149283 |
| ABC transporter, ATP-binding protein...1215 | 1.024 | 0.953704 | -0.1026 | 0.149294 |
| homoserine kinase | 1.080129 | 0.905616 | -0.25423 | 0.149329 |
| ATP synthase F1, beta subunit | 0.983767 | 1.051562 | 0.096144 | 0.149768 |
| hypothetical protein ABUW_2519 | 0.948735 | 1.045203 | 0.139707 | 0.150956 |
| proline-specific permease ProY | 1.001927 | 0.797986 | -0.32834 | 0.151279 |
| hypothetical protein ABUW_0076 | 0.8824 | 0.991076 | 0.167563 | 0.1513 |
| citrate synthase I | 0.946059 | 1.08026 | 0.191377 | 0.151836 |
| hypothetical protein ABUW_5010 (plasmid) | 1.001971 | 0.94877 | -0.07871 | 0.151857 |
| beta-lactamase...5 | 1.179046 | 1.046655 | -0.17183 | 0.152137 |
| hypothetical protein ABUW_3555 | 1.013986 | 0.97448 | -0.05733 | 0.152608 |
| hypothetical protein ABUW_1787 | 0.992812 | 1.0176 | 0.035578 | 0.153037 |
| ATP-dependent dsDNA exonuclease...1565 | 0.952467 | 0.990685 | 0.056758 | 0.154122 |
| ribosomal protein S18 | 1.145549 | 0.889145 | -0.36555 | 0.154585 |
| hypothetical protein ABUW_2625 | 1.0364 | 0.832727 | -0.31566 | 0.154985 |
| NADH-dependent enoyl-ACP reductase | 1.057397 | 1.017446 | -0.05557 | 0.155199 |
| single-strand binding protein | 0.982631 | 1.016943 | 0.049517 | 0.155919 |
| regulatory protein RecX | 1.452751 | 0.610383 | -1.251 | 0.156478 |
| regulator of ribonuclease activity A | 1.006336 | 1.071578 | 0.090625 | 0.157184 |
| thioesterase superfamily | 1.173841 | 0.959406 | -0.29102 | 0.157363 |
| phenylacetic acid degradation operon negative regulatory prc | 1.090415 | 1.000984 | -0.12346 | 0.157571 |
| transcriptional regulator-like protein | 1.078365 | 0.966253 | -0.15837 | 0.15762 |
| ribosomal protein L11 methyltransferase...416 | 1.084269 | 1.04239 | -0.05683 | 0.157687 |
| glyceropghosphoryl diester phosphodiesterase | 1.202574 | 1.058494 | -0.18411 | 0.157893 |
| oxygen-insensitive NAD(P)H nitroreductase | 1.069018 | 0.979189 | -0.12663 | 0.158052 |
| ribosomal protein S1 | 1.059758 | 0.958656 | -0.14465 | 0.15879 |
| transcriptional regulator SoxR-family | 1.095024 | 1.167059 | 0.091914 | 0.159133 |
| transcriptional regulator, MarR-family...1999 | 1.055846 | 0.928163 | -0.18595 | 0.159202 |
| S-formylglutathione hydrolase | 1.154991 | 0.92108 | -0.32648 | 0.159257 |
| threonine dehydratase...1260 | 0.985646 | 1.021768 | 0.051926 | 0.159868 |
| type IV pilus methyl-accepting chemotaxis sensory transducer | 1.166755 | 1.454885 | 0.318404 | 0.160012 |
| pseudouridine synthase A | 1.030764 | 0.912088 | -0.17647 | 0.160109 |
| putative cold shock protein | 0.974421 | 0.952454 | -0.0329 | 0.160191 |
| type VI secretion protein, family...1186 | 1.047542 | 0.999974 | -0.06705 | 0.160317 |

| | | | | |
|---|---|---|---|---|
| hypothetical protein ABUW_2797 | 0.920801 | 0.851398 | -0.11306 | 0.160354 |
| vanillate O-demethylase oxygenase subunit | 0.905912 | 1.07992 | 0.253482 | 0.160405 |
| thiamine-phosphate pyrophosphorylase | 0.968602 | 1.074937 | 0.150275 | 0.160552 |
| FHA domain protein | 1.163866 | 0.868884 | -0.42169 | 0.160714 |
| allantoicase | 0.962343 | 1.027959 | 0.09516 | 0.161045 |
| peptidyl-prolyl cis-trans isomerase, cyclophilin type | 1.054756 | 0.972807 | -0.11668 | 0.161398 |
| hypothetical protein ABUW_2016 | 0.971788 | 1.068257 | 0.136545 | 0.161758 |
| hypothetical protein ABUW_0879 | 1.04812 | 0.897398 | -0.22398 | 0.161878 |
| acetyl-CoA acetyltransferase...2447 | 0.991446 | 1.093005 | 0.140694 | 0.161884 |
| 2,3-dihydroxybenzoate-AMP ligase | 1.093796 | 0.9601 | -0.18809 | 0.16214 |
| phosphoribosylpyrophosphate synthetase | 0.965911 | 1.021912 | 0.081308 | 0.162347 |
| transcriptional regulator, GntR family...584 | 0.997484 | 0.943333 | -0.08053 | 0.162639 |
| transcriptional regulator, TenA family | 1.028551 | 1.051985 | 0.032502 | 0.162843 |
| nicotinamidase | 0.933215 | 1.052559 | 0.17362 | 0.16351 |
| peptidyl-prolyl cis-trans isomerase, FKBP-type | 1.030391 | 0.941293 | -0.13048 | 0.163923 |
| hypothetical protein ABUW_1450 | 1.053713 | 0.945819 | -0.15585 | 0.164558 |
| ribosomal protein L5 | 1.062911 | 0.955928 | -0.15305 | 0.164778 |
| hypothetical protein ABUW_2675 | 0.979108 | 1.107991 | 0.178408 | 0.164815 |
| hypothetical protein ABUW_0208 | 1.028427 | 0.984763 | -0.06259 | 0.164868 |
| transcriptional regulator...858 | 1.045431 | 0.943495 | -0.14801 | 0.164935 |
| 3-dehydroquinate dehydratase, type II | 1.256793 | 0.946698 | -0.40877 | 0.164951 |
| hypothetical protein ABUW_0300 | 1.809611 | 0.62109 | -1.54281 | 0.165119 |
| hypothetical protein ABUW_2565 | 1.155474 | 1.020544 | -0.17915 | 0.165233 |
| hypothetical protein ABUW_0700 | 1.026366 | 0.917555 | -0.16168 | 0.165233 |
| ATP-dependent DNA helicase | 0.988897 | 1.104263 | 0.159192 | 0.165413 |
| RNA polymerase sigma-70 factor | 0.984641 | 1.047248 | 0.088934 | 0.166217 |
| patatin family phospholipase | 1.044895 | 0.935759 | -0.15915 | 0.166528 |
| ribosomal protein S12 | 1.501345 | 0.755558 | -0.99064 | 0.166707 |
| phage head morphogenesis protein | 1.028568 | 0.935885 | -0.13623 | 0.166762 |
| hypothetical protein ABUW_1561 | 1.048038 | 0.923003 | -0.18328 | 0.167277 |
| nitrate transport ATP-binding protein NrtC | 1.074654 | 0.938411 | -0.19558 | 0.167576 |
| 3-isopropylmalate dehydratase, small subunit | 0.987195 | 0.929986 | -0.08613 | 0.16791 |
| fimbrial assembly protein PilQ | 0.957786 | 0.885985 | -0.11242 | 0.168124 |
| hypothetical protein ABUW_0480 | 1.017229 | 0.893495 | -0.18711 | 0.168318 |
| electron transfer flavoprotein subunit beta | 1.079598 | 0.946895 | -0.18922 | 0.168346 |
| hypothetical protein ABUW_2810 | 0.95072 | 0.987872 | 0.055304 | 0.16843 |
| protocatechuate 3,4-dioxygenase, alpha subunit | 0.966222 | 1.039827 | 0.105917 | 0.168519 |
| OsmC family protein...1447 | 0.959731 | 0.987838 | 0.041644 | 0.168715 |
| endonuclease III | 1.086466 | 1.013874 | -0.09976 | 0.169019 |
| quinate/shikimate dehydrogenase | 0.994692 | 0.931429 | -0.0948 | 0.169201 |
| RND efflux transporter | 1.035108 | 0.954632 | -0.11677 | 0.169319 |
| hypothetical protein ABUW_1128 | 0.928664 | 1.032313 | 0.152652 | 0.169865 |
| AdeR | 0.951807 | 1.107754 | 0.218895 | 0.171922 |
| hypothetical protein ABUW_3586 | 1.011162 | 0.916141 | -0.14237 | 0.172076 |
| glycine cleavage system H protein | 1.083839 | 0.962848 | -0.17077 | 0.172255 |
| oxidoreductase...1707 | 1.044809 | 1.005584 | -0.05521 | 0.172593 |
| transcriptional regulator, LysR family...848 | 0.827539 | 1.041707 | 0.33205 | 0.172803 |

| | | | | |
|---|---|---|---|---|
| secretion protein HlyD...2182 | 0.981533 | 0.994071 | 0.018313 | 0.173288 |
| glutathione peroxidase...2012 | 1.082592 | 0.978253 | -0.14621 | 0.174795 |
| Type IV secretory pathway, VirB4 component (plasmid) | 0.989025 | 1.181332 | 0.256335 | 0.174813 |
| glutaredoxin 3 | 1.021958 | 0.961498 | -0.08798 | 0.174889 |
| transcriptional regulator, LysR family...467 | 1.099677 | 0.976272 | -0.17173 | 0.175258 |
| hypothetical protein ABUW_2576 | 0.961132 | 1.074489 | 0.160844 | 0.175308 |
| hypothetical protein ABUW_2965 | 1.455798 | 0.587534 | -1.30907 | 0.175755 |
| endonuclease | 0.980307 | 1.042982 | 0.089408 | 0.17661 |
| hypothetical protein ABUW_2702 | 0.928129 | 1.027186 | 0.146301 | 0.177116 |
| NADPH-dependent 7-cyano-7-deazaguanine reductase (EsvE1 | 0.956032 | 1.022032 | 0.09631 | 0.177243 |
| hypothetical protein ABUW_1318 | 1.021419 | 0.865332 | -0.23925 | 0.17734 |
| hypothetical protein ABUW_0522 | 0.929513 | 1.184271 | 0.349453 | 0.177531 |
| hypothetical protein ABUW_0120 | 1.01177 | 1.079279 | 0.093186 | 0.177986 |
| biofilm-associated protein...2229 | 1.045381 | 0.916542 | -0.18976 | 0.177997 |
| DNA-directed RNA polymerase, beta subunit | 1.002099 | 0.977604 | -0.0357 | 0.178084 |
| 2'-aminoglycoside nucleotidyltransferase AadB (plasmid) | 1.050112 | 0.833548 | -0.33321 | 0.178124 |
| Transposase (plasmid) | 1.350473 | 0.568985 | -1.247 | 0.178989 |
| hypothetical protein ABUW_1649 | 0.927751 | 1.013698 | 0.127819 | 0.17913 |
| hypothetical protein ABUW_4107 (plasmid) | 1.704631 | 0.707051 | -1.26957 | 0.179252 |
| sulfite reductase | 1.018278 | 0.958446 | -0.08736 | 0.179994 |
| ATP phosphoribosyltransferase regulatory subunit | 0.970583 | 1.030156 | 0.085938 | 0.181035 |
| peptidyl-prolyl cis-trans isomerase...2181 | 0.977512 | 1.02854 | 0.073412 | 0.181109 |
| DNA topoisomerase I | 0.984804 | 0.994153 | 0.01363 | 0.181149 |
| methionyl-tRNA formyltransferase | 1.005627 | 1.044556 | 0.054795 | 0.181459 |
| hypothetical protein ABUW_1431 | 1.229697 | 0.824038 | -0.57752 | 0.182071 |
| replication initiation inhibitor | 1.05255 | 0.959653 | -0.13331 | 0.182231 |
| cytochrome D ubiquinol oxidase, subunit II...669 | 0.89159 | 0.9833 | 0.141251 | 0.182672 |
| adenylosuccinate synthetase | 0.97859 | 1.023037 | 0.064082 | 0.182823 |
| ribosomal protein L28 | 1.081188 | 0.980669 | -0.14078 | 0.183296 |
| hypothetical protein ABUW_3047 | 0.985343 | 1.020109 | 0.050025 | 0.183777 |
| hypothetical protein ABUW_0251 | 0.963575 | 1.00419 | 0.059564 | 0.183897 |
| hypothetical protein ABUW_3441 | 1.065041 | 0.979787 | -0.12037 | 0.18456 |
| GTP-binding protein Era | 0.983607 | 0.960781 | -0.03387 | 0.184904 |
| methylmalonate-semialdehyde dehydrogenase | 0.981778 | 1.01114 | 0.042514 | 0.18506 |
| Glutamate synthase (NADPH) | 0.974688 | 1.014696 | 0.058034 | 0.185309 |
| hypothetical protein ABUW_3768 | 0.996121 | 0.819421 | -0.28172 | 0.185608 |
| DNA polymerase III, delta subunit | 0.951843 | 0.875566 | -0.12051 | 0.185738 |
| arginine N-succinyltransferase...2048 | 0.938537 | 0.99681 | 0.086905 | 0.185882 |
| short chain dehydrogenase...132 | 1.107316 | 0.769843 | -0.52443 | 0.186097 |
| hypothetical protein ABUW_1376 | 0.846395 | 0.907514 | 0.100589 | 0.186852 |
| putative methyltransferase | 0.970324 | 1.014457 | 0.064169 | 0.186911 |
| para-aminobenzoate/anthranilate synthase glutamine amidot | 0.989853 | 1.051274 | 0.086853 | 0.188522 |
| intracellular septation protein A | 0.991738 | 1.071186 | 0.111177 | 0.189316 |
| hypothetical protein ABUW_2805 | 1.218773 | 0.821695 | -0.56876 | 0.190099 |
| hemerythrin/HHE cation-binding motif- containing protein | 0.936222 | 1.107434 | 0.242299 | 0.190108 |
| hypothetical protein ABUW_2875 | 1.069874 | 0.950876 | -0.17011 | 0.19027 |
| 5,10-methylenetetrahydrofolate reductase | 1.033588 | 0.987593 | -0.06567 | 0.190687 |

| | | | | |
|---|---|---|---|---|
| ferredoxin-NADP reductase | 0.990876 | 1.05071 | 0.084587 | 0.190801 |
| DNA polymerase V component | 0.998446 | 0.968549 | -0.04386 | 0.191041 |
| hydroxyacylglutathione hydrolase | 1.067822 | 0.983691 | -0.11839 | 0.191937 |
| ribosomal RNA large subunit methytransferase A | 1.039034 | 0.993556 | -0.06457 | 0.192116 |
| hypothetical protein ABUW_3827 | 0.96245 | 1.025679 | 0.091796 | 0.19231 |
| hypothetical protein ABUW_1885 | 1.136269 | 0.893495 | -0.34677 | 0.192403 |
| glyceraldehyde-3-phosphate dehydrogenase | 0.924179 | 0.971956 | 0.072718 | 0.192609 |
| ribosomal protein L14 | 1.054056 | 0.998903 | -0.07754 | 0.193547 |
| hypothetical protein ABUW_0783 | 1.052668 | 0.959079 | -0.13433 | 0.193716 |
| glutathione S-transferase...1378 | 1.000761 | 0.940498 | -0.0896 | 0.194404 |
| hypothetical protein ABUW_2915 | 0.998656 | 1.056612 | 0.081386 | 0.195373 |
| histidine triad protein...1359 | 1.020412 | 0.988651 | -0.04562 | 0.195658 |
| peroxiredoxin | 1.092348 | 0.938059 | -0.21968 | 0.195942 |
| hypothetical protein ABUW_2329 | 1.107373 | 0.993153 | -0.15705 | 0.195978 |
| protease | 1.013618 | 1.165662 | 0.201636 | 0.196233 |
| ribosomal protein L3 | 1.030325 | 0.996356 | -0.04837 | 0.196445 |
| aquaporin Z | 1.080746 | 0.915179 | -0.2399 | 0.19645 |
| hypothetical protein ABUW_4073 (plasmid) | 1.048452 | 1.024142 | -0.03385 | 0.196456 |
| hypothetical protein ABUW_3472 | 1.072241 | 1.218583 | 0.184576 | 0.196542 |
| hypothetical protein ABUW_2665 | 1.016695 | 0.85631 | -0.24768 | 0.197283 |
| two-component system response regulator protein...2318 | 1.022221 | 0.994677 | -0.03941 | 0.199035 |
| hypothetical protein ABUW_1011 | 1.008175 | 0.966506 | -0.0609 | 0.199593 |
| Csy3 family crispr-associated protein | 0.966035 | 0.998512 | 0.047704 | 0.200526 |
| lipopolysaccharide transport periplasmic protein LptA | 0.977036 | 0.998729 | 0.031681 | 0.200537 |
| hypothetical protein ABUW_3850 | 1.142315 | 0.912591 | -0.32392 | 0.20106 |
| hypothetical protein ABUW_0927 | 1.205641 | 0.89451 | -0.43063 | 0.201125 |
| multidrug ABC transporter permease | 2.084595 | 0.579809 | -1.84612 | 0.201412 |
| toluene tolerance protein Ttg2B | 0.918961 | 0.980905 | 0.09411 | 0.201617 |
| thiol:disulfide interchange protein DsbC | 1.002361 | 0.974548 | -0.0406 | 0.201924 |
| DNA strand exchange and recombination protein RecA | 1.024685 | 0.959471 | -0.09487 | 0.202623 |
| lipoprotein releasing system, ATP-binding protein | 0.942257 | 1.022272 | 0.117586 | 0.202701 |
| organic solvent tolerance protein | 1.021822 | 0.99952 | -0.03184 | 0.203647 |
| hypothetical protein ABUW_0780 | 1.07095 | 0.823 | -0.37993 | 0.204073 |
| histidine ammonia-lyase | 1.00048 | 0.969008 | -0.04611 | 0.204195 |
| hypothetical protein ABUW_2803 | 1.053191 | 1.131642 | 0.103651 | 0.204418 |
| ParA family protein | 0.995574 | 1.015504 | 0.028595 | 0.205095 |
| hypothetical protein ABUW_2297 | 0.952975 | 1.010266 | 0.084226 | 0.205834 |
| DNA translocase FtsK | 0.933675 | 1.000734 | 0.100066 | 0.206439 |
| hypothetical protein ABUW_1590 | 0.954132 | 1.017128 | 0.092241 | 0.207242 |
| permease...1066 | 1.058367 | 0.909927 | -0.21802 | 0.207795 |
| transcriptional regulator, GntR family...609 | 0.99184 | 0.841175 | -0.2377 | 0.208073 |
| guanine deaminase...565 | 0.985661 | 1.135302 | 0.203912 | 0.208355 |
| 3-hydroxybutyrate dehydrogenase | 1.025169 | 1.055293 | 0.041781 | 0.20837 |
| carbon storage regulator | 0.980481 | 0.995485 | 0.021911 | 0.208433 |
| hypothetical protein ABUW_0338 | 0.432131 | 0.361793 | -0.2563 | 0.209266 |
| transcriptional activator protein CusR | 0.955809 | 0.927159 | -0.04391 | 0.210912 |
| TonB family protein | 0.845189 | 0.98642 | 0.222929 | 0.211885 |

| | | | | |
|---|---|---|---|---|
| alpha/beta hydrolase fold protein...112 | 1.136326 | 1.045034 | -0.12083 | 0.212275 |
| putative esterase | 1.126231 | 0.953747 | -0.23983 | 0.212325 |
| ribosomal subunit interface protein | 1.022095 | 0.903972 | -0.17718 | 0.212493 |
| 3-hydroxyisobutyrate dehydrogenase...1438 | 1.058121 | 1.036897 | -0.02923 | 0.212548 |
| rhodanese domain protein...194 | 0.904942 | 0.964543 | 0.09202 | 0.212588 |
| mechanosensitive ion channel family protein | 0.906443 | 0.985565 | 0.120735 | 0.212888 |
| hypothetical protein ABUW_2893 | 0.934907 | 0.98133 | 0.069915 | 0.213154 |
| dihydrodipicolinate synthetase | 0.983747 | 1.045239 | 0.087473 | 0.213163 |
| bifunctional hydroxy-methylpyrimidine kinase/hydroxy-phosp | 0.984991 | 1.009167 | 0.034983 | 0.213582 |
| DNA gyrase, A subunit | 1.002608 | 0.992753 | -0.01425 | 0.213725 |
| acetyltransferase...1307 | 0.99138 | 1.031804 | 0.057659 | 0.214338 |
| thiazole biosynthesis protein ThiG | 1.038561 | 1.099561 | 0.082341 | 0.215172 |
| protein in PtsN-ptsO intergenic region | 0.980818 | 0.990442 | 0.014087 | 0.215479 |
| glycoside hydrolase/deacetylase family protein | 1.26922 | 0.790718 | -0.68271 | 0.215758 |
| hypothetical protein ABUW_1868 | 1.023688 | 0.951606 | -0.10534 | 0.216105 |
| translation elongation factor G | 1.056409 | 0.952153 | -0.1499 | 0.216198 |
| hypothetical protein ABUW_0771 | 0.998013 | 1.061273 | 0.088664 | 0.216687 |
| folylpolyglutamate synthetase/dihydrofolate synthase | 0.963827 | 0.996054 | 0.04745 | 0.216695 |
| hypothetical protein ABUW_3138 | 0.677104 | 0.62948 | -0.10522 | 0.217086 |
| oligopeptidase A | 0.95992 | 0.917547 | -0.06513 | 0.217575 |
| alcohol dehydrogenase, class IV | 0.981891 | 1.015264 | 0.04822 | 0.217604 |
| hypothetical protein ABUW_1988 | 1.246507 | 1.371424 | 0.137784 | 0.217968 |
| imidazolonepropionase | 1.047461 | 0.932462 | -0.16778 | 0.217992 |
| polyribonucleotide nucleotidyltransferase | 1.01788 | 0.976081 | -0.0605 | 0.218224 |
| hypothetical protein ABUW_1319 | 1.061603 | 0.893853 | -0.24813 | 0.218336 |
| response regulator protein | 1.033265 | 1.093932 | 0.082313 | 0.219548 |
| transcriptional regulator, LysR family...873 | 1.036863 | 1.093985 | 0.077368 | 0.220265 |
| sodium/proline symporter | 0.955023 | 1.060283 | 0.150842 | 0.220689 |
| hypothetical protein ABUW_0270 | 0.982051 | 1.057722 | 0.10709 | 0.220711 |
| hypothetical protein ABUW_4002 (plasmid) | 1.218697 | 0.898698 | -0.43943 | 0.220933 |
| hypothetical protein ABUW_0742 | 1.060245 | 0.97708 | -0.11785 | 0.221305 |
| general secretion pathway protein F | 0.919164 | 1.016117 | 0.144673 | 0.221317 |
| aromatic-ring-hydroxylating dioxygenase beta subunit | 0.985869 | 1.029763 | 0.062844 | 0.221637 |
| hypothetical protein ABUW_0781 | 1.021615 | 0.950987 | -0.10335 | 0.221845 |
| tRNA-dihydrouridine synthase B | 1.043275 | 1.127492 | 0.111998 | 0.222104 |
| nicotinate phosphoribosyltransferase | 0.957951 | 0.993348 | 0.052346 | 0.222791 |
| dioxygenase beta subunit | 1.10494 | 0.97852 | -0.17529 | 0.222916 |
| nucleotidyl transferase family protein | 1.044531 | 1.006095 | -0.05409 | 0.223074 |
| acyl-CoA thioesterase | 1.20694 | 0.824945 | -0.54898 | 0.223167 |
| magnesium Mg(2+)/cobalt Co(2+) transport protein | 1.048189 | 0.828594 | -0.33916 | 0.223199 |
| acetyl-CoA C-acyltransferase FadA | 1.03656 | 1.027405 | -0.0128 | 0.22351 |
| LemA family | 0.983358 | 1.082123 | 0.138076 | 0.224174 |
| transcriptional regulator, GntR family...598 | 1.044954 | 0.956801 | -0.12715 | 0.225006 |
| isochorismate hydrolase | 1.086188 | 0.963684 | -0.17264 | 0.225224 |
| hypothetical protein ABUW_1920 | 1.000962 | 1.034301 | 0.047269 | 0.226528 |
| O-methyltransferase | 0.99517 | 1.061721 | 0.09339 | 0.228108 |
| inner membrane protein | 1.056562 | 0.953468 | -0.14812 | 0.228115 |

| | | | | |
|---|---|---|---|---|
| ammonium transporter...748 | 0.924299 | 0.990614 | 0.099964 | 0.228365 |
| hypothetical protein ABUW_1466 | 0.95025 | 1.000066 | 0.073715 | 0.22882 |
| ankyrin repeat protein | 0.880352 | 0.823764 | -0.09585 | 0.229177 |
| transcriptional regulator, Crp/Fnr family | 1.152618 | 1.626141 | 0.496538 | 0.229425 |
| catalase/peroxidase HPI | 0.943161 | 0.972726 | 0.044529 | 0.229515 |
| arsenate reductase...599 | 0.973737 | 1.082636 | 0.152945 | 0.230067 |
| translation elongation factor Tu | 1.04776 | 0.972078 | -0.10817 | 0.230188 |
| phage tail completion protein | 0.917258 | 1.067205 | 0.218438 | 0.231351 |
| 6,7-dimethyl-8-ribityllumazine synthase | 1.039825 | 0.950232 | -0.12999 | 0.231685 |
| OmpA/MotB...1765 | 1.011386 | 0.984135 | -0.03941 | 0.23187 |
| ketol-acid reductoisomerase | 1.005912 | 1.065467 | 0.082983 | 0.232471 |
| cold-shock domain protein | 0.924108 | 0.971437 | 0.07206 | 0.232989 |
| hypothetical protein ABUW_1441 | 1.012599 | 0.95025 | -0.09168 | 0.233331 |
| hypothetical protein ABUW_2238 | 1.067311 | 1.019998 | -0.06542 | 0.233394 |
| transcriptional regulator, AsnC family...799 | 1.02703 | 0.982938 | -0.06331 | 0.234408 |
| molybdopterin oxidoreductase...141 | 1.277316 | 1.015337 | -0.33116 | 0.235002 |
| hypothetical protein ABUW_2211 | 0.933556 | 1.081195 | 0.211819 | 0.235098 |
| pyridoxamine 5'-phosphate oxidase | 0.959291 | 1.021997 | 0.091351 | 0.235345 |
| S-adenosyl-L-methionine-dependent methyltransferase famil | 1.02435 | 1.073455 | 0.067552 | 0.23576 |
| transcriptional regulator, TetR family...887 | 0.993005 | 1.031454 | 0.054806 | 0.235846 |
| transcription elongation factor GreA | 1.03794 | 0.999843 | -0.05395 | 0.235939 |
| cell division protein FtsZ | 0.979863 | 1.016371 | 0.052776 | 0.236938 |
| host factor Hfq | 0.979755 | 1.015306 | 0.051422 | 0.237061 |
| esterase...903 | 0.98599 | 0.951445 | -0.05145 | 0.238483 |
| ribosomal protein S11 | 1.044064 | 1.006768 | -0.05248 | 0.238767 |
| aldehyde dehydrogenase...2483 | 1.046713 | 0.944747 | -0.14787 | 0.238962 |
| ABC-type transporter, periplasmic component: TauT family | 0.999532 | 0.921134 | -0.11784 | 0.239056 |
| hypothetical protein ABUW_3559 | 0.940444 | 1.006683 | 0.098196 | 0.239452 |
| putative prophage protein | 1.012668 | 0.946239 | -0.09788 | 0.239556 |
| 3-oxoacid CoA-transferase, subunit B | 1.036377 | 1.087387 | 0.069316 | 0.23967 |
| transcriptional regulator, LysR family...350 | 1.019336 | 0.931436 | -0.1301 | 0.240823 |
| chromosome segregation and condensation protein ScpA | 0.956381 | 1.026663 | 0.102305 | 0.241058 |
| hypothetical protein ABUW_1780 | 1.091782 | 0.925555 | -0.23829 | 0.241436 |
| hypothetical protein ABUW_0602 | 0.978352 | 0.936309 | -0.06337 | 0.241607 |
| outer membrane protein...2190 | 1.002659 | 0.969343 | -0.04875 | 0.242096 |
| oxidoreductase...2485 | 1.034595 | 0.95991 | -0.1081 | 0.242186 |
| hypothetical protein ABUW_0619 | 0.9236 | 1.028883 | 0.155739 | 0.242346 |
| beta-hydroxyacyl-(acyl-carrier-protein) dehydratase | 1.001242 | 1.080255 | 0.109582 | 0.243135 |
| ATPase | 0.969006 | 1.003268 | 0.05013 | 0.243231 |
| DNA replication protein (plasmid)...905 | 0.913502 | 0.999567 | 0.129895 | 0.244104 |
| two-component system DNA-binding response regulator | 0.932004 | 0.942904 | 0.016775 | 0.244389 |
| ribosomal protein S2 | 1.057078 | 1.002859 | -0.07596 | 0.244808 |
| NADH dehydrogenase I chain H | 0.922348 | 1.01973 | 0.144804 | 0.244919 |
| 4-(cytidine 5'-diphospho)-2-C-methyl-D-erythritol kinase | 0.973248 | 0.948705 | -0.03685 | 0.245712 |
| alpha/beta fold family hydrolase...195 | 1.144449 | 1.062965 | -0.10656 | 0.245983 |
| type VI secretion ATPase, ClpV1 family | 0.971633 | 0.933498 | -0.05776 | 0.246321 |
| iron uptake factor | 0.973857 | 1.053562 | 0.113493 | 0.247479 |

| | | | | |
|---|---|---|---|---|
| radical SAM enzyme, Cfr family | 0.905048 | 0.946117 | 0.064024 | 0.247843 |
| ribosomal protein L31 | 1.111366 | 0.963994 | -0.20524 | 0.247932 |
| crispr-associated helicase Cas3 | 1.157343 | 1.001052 | -0.2093 | 0.248839 |
| tyrosine-protein kinase ptk | 0.989479 | 1.014765 | 0.036404 | 0.249078 |
| chromosomal replication initiator protein DnaA | 0.987372 | 1.01274 | 0.036599 | 0.249582 |
| 3-isopropylmalate dehydrogenase | 0.961799 | 1.037096 | 0.108742 | 0.250203 |
| ubiquinone biosynthesis hydroxylase, UbiH/UbiF/VisC/COQ6 f | 0.965521 | 0.992894 | 0.040332 | 0.250436 |
| alanine racemase...1968 | 0.997361 | 1.112049 | 0.157033 | 0.250961 |
| peptidase S45, penicillin amidase | 1.014454 | 1.032018 | 0.024765 | 0.251649 |
| anthranilate phosphoribosyltransferase | 0.978481 | 1.009195 | 0.04459 | 0.252318 |
| multi-sensor hybrid histidine kinase | 0.97046 | 0.995919 | 0.037361 | 0.252731 |
| hypothetical protein ABUW_3696 | 0.977385 | 1.031309 | 0.077478 | 0.253018 |
| 5-carboxymethyl-2-hydroxymuconate isomerase | 0.940266 | 1.016609 | 0.112625 | 0.253114 |
| putative transcriptional regulator...1137 | 0.932596 | 1.024371 | 0.135415 | 0.253878 |
| hypothetical protein ABUW_0216 | 0.775289 | 0.924188 | 0.253451 | 0.254356 |
| type VI secretion protein IcmF | 0.994688 | 0.964675 | -0.0442 | 0.25542 |
| major facilitator superfamily MFS_1...375 | 1.04632 | 0.966372 | -0.11467 | 0.256148 |
| transcriptional regulator, LysR family...1879 | 0.954714 | 1.016894 | 0.09103 | 0.256301 |
| transcriptional regulator, AraC family...9 | 1.114046 | 1.292887 | 0.214788 | 0.256632 |
| hypothetical protein ABUW_2190 | 0.989726 | 1.052619 | 0.088882 | 0.256904 |
| polyphosphate kinase...928 | 0.967087 | 1.001541 | 0.050504 | 0.257114 |
| heat shock protein htpX | 1.000904 | 0.950227 | -0.07496 | 0.257848 |
| hypothetical protein ABUW_1108 | 1.029107 | 1.010844 | -0.02583 | 0.258039 |
| putative MTA/SAH nucleosidase | 1.005266 | 0.981262 | -0.03487 | 0.258631 |
| general secretion pathway protein...136 | 0.99211 | 1.13349 | 0.1922 | 0.259217 |
| urease accessory protein F | 0.848452 | 1.075842 | 0.342562 | 0.259983 |
| hydrolase, NUDIX family | 0.928786 | 0.79249 | -0.22895 | 0.260712 |
| hydroxymethylbutenyl pyrophosphate reductase | 1.006004 | 0.969167 | -0.05382 | 0.261315 |
| 1,6-dihydroxycyclohexa-2,4-diene-1-carboxylate dehydrogena | 0.846691 | 1.015222 | 0.261887 | 0.261886 |
| tolerance to group A colicins single-stranded filamentous DN/ | 0.943341 | 0.986384 | 0.064371 | 0.261909 |
| nitrite reductase [NAD(P)H], large subunit | 1.04329 | 0.997823 | -0.06428 | 0.262199 |
| taurine ABC transporter, periplasmic binding protein | 1.122664 | 0.957743 | -0.22921 | 0.263062 |
| phospholipase C, phosphocholine-specific...47 | 1.050852 | 0.922637 | -0.18772 | 0.263589 |
| DnaA family protein | 0.958875 | 0.9886 | 0.044044 | 0.263969 |
| tryptophan synthase, beta subunit...2055 | 0.991142 | 1.033243 | 0.060016 | 0.26408 |
| lipoprotein | 0.977585 | 1.005681 | 0.040878 | 0.26613 |
| hypothetical protein ABUW_0745 | 1.051092 | 1.000364 | -0.07136 | 0.266134 |
| methylenetetrahydrofolate reductase domain protein | 1.045854 | 1.006004 | -0.05605 | 0.266916 |
| serine acetyltransferase | 0.923484 | 1.014318 | 0.135352 | 0.268009 |
| TatD-related deoxyribonuclease | 1.040793 | 1.004232 | -0.05159 | 0.269222 |
| hypothetical protein ABUW_2900 | 1.110505 | 1.044776 | -0.08802 | 0.270352 |
| adenylate/guanylate cyclase | 1.036514 | 1.118023 | 0.109211 | 0.270624 |
| cysteine desulfurase | 0.842552 | 1.060748 | 0.332243 | 0.270894 |
| hypothetical protein ABUW_4004 (plasmid) | 0.970966 | 1.030178 | 0.085401 | 0.27115 |
| short-chain dehydrogenase/reductase...2261 | 1.032414 | 0.985959 | -0.06642 | 0.271676 |
| ribonuclease Z | 1.02501 | 1.105453 | 0.108999 | 0.272295 |
| putative polysaccharide deacetylase | 0.93536 | 0.985019 | 0.07463 | 0.272623 |

| | | | | |
|---|---|---|---|---|
| peptidyl-tRNA hydrolase | 1.079267 | 0.945588 | -0.19077 | 0.272686 |
| paraquat-inducible protein B | 0.971949 | 1.0038 | 0.04652 | 0.27446 |
| hypothetical protein ABUW_0519 | 1.025557 | 0.991614 | -0.04856 | 0.274806 |
| hypothetical protein ABUW_1172 | 1.017178 | 0.957117 | -0.0878 | 0.275534 |
| IscR-regulated protein YhgI | 0.987763 | 1.040867 | 0.07555 | 0.275677 |
| tail tape measure protein...509 | 1.231295 | 1.097653 | -0.16575 | 0.276586 |
| 2-amino-4-hydroxy-6-hydroxymethyldihydropteridine pyroph | 0.955541 | 1.104204 | 0.208617 | 0.276765 |
| hypothetical protein ABUW_0644 | 1.027292 | 0.974753 | -0.07574 | 0.277131 |
| glutathione S-transferase...596 | 1.011351 | 0.979392 | -0.04633 | 0.277326 |
| alcohol dehydrogenase, zinc-binding...1843 | 0.983416 | 1.029039 | 0.065423 | 0.27799 |
| transaldolase | 0.991097 | 0.975828 | -0.0224 | 0.278205 |
| hypothetical protein ABUW_1652 | 1.041545 | 0.982903 | -0.0836 | 0.278713 |
| beta-lactamase...991 | 1.042172 | 0.919289 | -0.181 | 0.279398 |
| hypothetical protein ABUW_1725 | 0.957722 | 1.022915 | 0.095009 | 0.279426 |
| 4-carboxymuconolactone decarboxylase...656 | 1.000657 | 1.058794 | 0.081474 | 0.27952 |
| sugar kinase, ribokinase family | 0.948166 | 0.831882 | -0.18876 | 0.279723 |
| MscS Mechanosensitive ion channel...477 | 0.932054 | 0.956708 | 0.037666 | 0.280037 |
| hypothetical protein ABUW_2847 | 0.985775 | 1.022556 | 0.05285 | 0.280831 |
| hypothetical protein ABUW_1295 | 0.976179 | 0.854025 | -0.19287 | 0.282077 |
| DoxX family protein | 1.033287 | 0.979843 | -0.07662 | 0.282253 |
| hypothetical protein ABUW_3422 | 1.043024 | 1.008604 | -0.04841 | 0.282832 |
| ABC-type nitrate/sulfonate/bicarbonate transport system per | 0.999178 | 1.107697 | 0.148751 | 0.283169 |
| transcriptional regulator, LysR family...327 | 1.09608 | 1.031066 | -0.08822 | 0.283338 |
| transcription antitermination factor NusB | 1.013596 | 0.975128 | -0.05582 | 0.284144 |
| hypothetical protein ABUW_2346 | 1.021686 | 0.950432 | -0.1043 | 0.284301 |
| carbon-nitrogen hydrolase | 0.973138 | 1.00681 | 0.049076 | 0.286196 |
| type VI secretion protein, EvpB/family | 0.989277 | 1.041975 | 0.074874 | 0.287546 |
| hypothetical protein ABUW_0984 | 0.996664 | 0.947924 | -0.07234 | 0.287595 |
| crispr-associated protein, Csy2 family protein | 0.956207 | 1.029782 | 0.106944 | 0.287839 |
| acetylornithine aminotransferase | 0.962453 | 0.990527 | 0.04148 | 0.287867 |
| hypothetical protein ABUW_1154 | 0.947665 | 1.235918 | 0.383134 | 0.28924 |
| transcriptional regulator EstR, HTH-type | 0.980168 | 1.004307 | 0.0351 | 0.289973 |
| catalase...1924 | 0.994786 | 0.946858 | -0.07124 | 0.290453 |
| UDP-N-acetylglucosamine 2-epimerase...1088 | 1.112387 | 0.963695 | -0.20701 | 0.290879 |
| 3-hydroxyphenylpropionic acid transporter | 0.903558 | 0.953195 | 0.077154 | 0.291171 |
| cation efflux system protein...965 | 1.003843 | 0.963103 | -0.05977 | 0.291579 |
| DNA gyrase, B subunit | 1.013879 | 0.995755 | -0.02602 | 0.291588 |
| two component transcriptional regulator, LuxR family | 0.955727 | 0.883154 | -0.11393 | 0.291797 |
| L-carnitine dehydrogenase | 1.056554 | 1.108802 | 0.069635 | 0.291972 |
| methylated-DNA--protein-cysteine methyltransferase transcri | 0.94557 | 1.00107 | 0.082288 | 0.29241 |
| transcriptional regulator, GntR family...321 | 1.011517 | 1.067333 | 0.07749 | 0.29254 |
| enoyl-CoA hydratase/isomerase...1654 | 1.021154 | 0.990619 | -0.0438 | 0.29297 |
| endonuclease/exonuclease/phosphatase...541 | 0.986195 | 1.039122 | 0.075421 | 0.293711 |
| OsmC-like protein | 1.027779 | 0.963024 | -0.09389 | 0.293737 |
| hypothetical protein ABUW_3267 | 0.947039 | 0.922652 | -0.03764 | 0.294336 |
| transcriptional regulator, IclR family...398 | 0.934923 | 1.009085 | 0.110129 | 0.295686 |
| ATP-dependent DNA helicase RecG | 1.008007 | 0.981072 | -0.03907 | 0.29614 |

| | | | | |
|---|---|---|---|---|
| tryptophan synthase, alpha subunit | 1.049403 | 1.012247 | -0.05201 | 0.296848 |
| signal transduction histidine kinase, nitrogen specific, NtrB | 0.983864 | 0.937643 | -0.06942 | 0.297152 |
| hypothetical protein ABUW_4087 (plasmid) | 1.031251 | 0.984991 | -0.06621 | 0.29733 |
| urease, gamma subunit | 0.954204 | 0.988603 | 0.051094 | 0.297922 |
| phosphoribosylaminoimidazole-succinocarboxamide synthase | 0.957066 | 1.004219 | 0.069384 | 0.297999 |
| lysophospholipase | 1.016313 | 1.057289 | 0.057025 | 0.29858 |
| glycine cleavage system transcriptional activator | 0.977139 | 1.042196 | 0.092991 | 0.29954 |
| hypothetical protein ABUW_0060 | 0.915907 | 0.977167 | 0.093405 | 0.299692 |
| extradiol ring-cleavage dioxygenase, class III enzyme, subunit | 0.991048 | 1.028513 | 0.053534 | 0.300074 |
| protein-export membrane protein SecD | 0.962737 | 0.987986 | 0.037349 | 0.300183 |
| hypothetical protein ABUW_2157 | 0.90957 | 1.030031 | 0.179432 | 0.300659 |
| transcriptional regulator, LysR family...387 | 0.975245 | 0.846091 | -0.20495 | 0.301001 |
| DNA-3-methyladenine glycosylase 1 | 0.980371 | 1.008693 | 0.041088 | 0.302148 |
| NAD(P)H oxidoreductase | 1.101815 | 1.028369 | -0.09952 | 0.302979 |
| multidrug efflux protein AdeI | 1.006469 | 0.976346 | -0.04384 | 0.303047 |
| hypothetical protein ABUW_4094 (plasmid) | 1.002763 | 1.113307 | 0.150871 | 0.304822 |
| oxidoreductase...1921 | 1.010721 | 0.925551 | -0.127 | 0.305337 |
| hypothetical protein ABUW_0826 | 1.061435 | 1.086777 | 0.034041 | 0.305746 |
| fructose-bisphosphate aldolase, class II, Calvin cycle subtype | 1.018417 | 0.985197 | -0.04784 | 0.306179 |
| TonB-dependent receptor...623 | 1.064134 | 0.981741 | -0.11626 | 0.308653 |
| 5-formyltetrahydrofolate cyclo-ligase | 0.881137 | 0.966978 | 0.134117 | 0.30881 |
| malonate utilization transcriptional regulator, LysR family | 1.049097 | 0.939636 | -0.15897 | 0.308852 |
| transcriptional regulator, LysR family...456 | 1.004541 | 0.9552 | -0.07266 | 0.308955 |
| hypothetical protein ABUW_0023 | 1.000404 | 0.919407 | -0.12181 | 0.309803 |
| hypothetical protein ABUW_0636 | 1.050276 | 0.973233 | -0.10991 | 0.309887 |
| monooxygenase, flavin-binding family...313 | 1.026126 | 1.095697 | 0.094642 | 0.310048 |
| Sel1 repeat family | 0.808804 | 0.94823 | 0.229447 | 0.310096 |
| DNA-directed RNA polymerase, omega subunit | 1.007489 | 0.972031 | -0.05169 | 0.310229 |
| phenazine biosynthesis protein PhzF family...17 | 1.376321 | 0.879462 | -0.64612 | 0.31035 |
| hypothetical protein ABUW_0793 | 0.979653 | 0.895763 | -0.12915 | 0.311009 |
| cytochrome O ubiquinol oxidase subunit IV | 0.976038 | 0.744007 | -0.39162 | 0.311485 |
| dihydroorotase, homodimeric type | 0.985832 | 1.024571 | 0.055606 | 0.313555 |
| DNA polymerase III delta prime subunit | 0.956152 | 0.992487 | 0.053809 | 0.314592 |
| hypothetical protein ABUW_1984 | 1.029078 | 0.967937 | -0.08837 | 0.315198 |
| short-chain dehydrogenase/reductase...1686 | 0.980355 | 0.994881 | 0.02122 | 0.316668 |
| hypothetical protein ABUW_1225 | 0.949376 | 0.829336 | -0.19502 | 0.316691 |
| extracellular solute-binding protein, family 5 | 0.925952 | 0.806585 | -0.19911 | 0.31676 |
| macrolide export ATP-binding/permease protein MacB | 1.021378 | 0.969672 | -0.07495 | 0.317149 |
| hypothetical protein ABUW_3748 | 0.988121 | 0.936226 | -0.07783 | 0.31723 |
| hypothetical protein ABUW_1357 | 0.982058 | 1.022221 | 0.057827 | 0.317244 |
| RNA helicase | 1.028023 | 1.012544 | -0.02189 | 0.317299 |
| EsvE2 | 1.016887 | 0.965019 | -0.07553 | 0.318472 |
| 3-demethylubiquinone-9 3-O-methyltransferase | 1.007724 | 1.094463 | 0.119123 | 0.318613 |
| formimidoylglutamase | 0.991586 | 0.93687 | -0.08189 | 0.31898 |
| valyl-tRNA synthetase | 0.992425 | 1.023406 | 0.044349 | 0.319242 |
| hypothetical protein ABUW_0751 | 0.923748 | 1.018837 | 0.141352 | 0.320189 |
| hypothetical protein ABUW_3436 | 0.929361 | 1.00692 | 0.115638 | 0.320456 |

| | | | | |
|---|---|---|---|---|
| glutamate--cysteine ligase | 1.065557 | 0.99887 | -0.09324 | 0.321041 |
| 4-hydroxythreonine-4-phosphate dehydrogenase | 0.955908 | 1.019928 | 0.093524 | 0.321098 |
| putative oxidoreductase | 1.097152 | 1.333673 | 0.281641 | 0.32132 |
| enoyl-CoA hydratase...1218 | 1.028868 | 0.978662 | -0.07217 | 0.322926 |
| hypothetical protein ABUW_3066 | 1.084401 | 1.024725 | -0.08166 | 0.323234 |
| crispr-associated protein, Csy4 family | 1.051044 | 1.080963 | 0.040494 | 0.324652 |
| DNA methylase N-4/N-6 | 1.031307 | 1.001242 | -0.04268 | 0.325313 |
| beta-ketoadipyl CoA thiolase...2305 | 0.9708 | 0.995425 | 0.036138 | 0.326506 |
| ribosomal protein L17 | 1.079942 | 1.011666 | -0.09422 | 0.326687 |
| transcriptional regulator, LysR family...526 | 0.933468 | 1.023663 | 0.133069 | 0.328382 |
| hypothetical protein ABUW_0952 | 1.015422 | 1.072553 | 0.07897 | 0.331255 |
| hypothetical protein ABUW_2448 | 1.005732 | 1.045264 | 0.055621 | 0.331516 |
| hypothetical protein ABUW_3524 | 1.03016 | 1.002553 | -0.03919 | 0.331664 |
| pili assembly chaperone | 0.925685 | 1.032228 | 0.157168 | 0.33174 |
| acetyl-CoA carboxylase, carboxyl transferase, alpha subunit | 1.03775 | 1.028057 | -0.01354 | 0.332063 |
| hypothetical protein ABUW_0479 | 1.071919 | 0.987257 | -0.1187 | 0.332115 |
| exonuclease V gamma chain | 0.880238 | 0.815455 | -0.11029 | 0.33235 |
| hypothetical protein ABUW_0985 | 0.959952 | 0.890214 | -0.10881 | 0.33287 |
| thioredoxin-disulfide reductase...678 | 1.080916 | 1.030209 | -0.06932 | 0.333407 |
| 2-C-methyl-D-erythritol 2,4-cyclodiphosphate synthase | 0.996119 | 1.025704 | 0.042223 | 0.336668 |
| putative transcriptional regulator, Cro/CI family | 1.024287 | 0.980843 | -0.06253 | 0.337348 |
| protein DedA | 1.170126 | 1.073606 | -0.1242 | 0.337784 |
| hypothetical protein ABUW_2047 | 0.302174 | 2.501766 | 3.049494 | 0.340799 |
| phage integrase | 0.987305 | 0.916475 | -0.1074 | 0.342445 |
| hypothetical protein ABUW_3085 | 1.082258 | 1.001292 | -0.11218 | 0.342494 |
| hypothetical protein ABUW_3459 | 1.105714 | 1.066456 | -0.05215 | 0.342782 |
| ATP synthase F1, delta subunit | 1.006929 | 1.065233 | 0.081207 | 0.343496 |
| hypothetical protein ABUW_3579 | 1.000773 | 0.986579 | -0.02061 | 0.344422 |
| catalase...2482 | 0.855986 | 0.954568 | 0.157261 | 0.345545 |
| P-hydroxybenzoate hydroxylase transcriptional activator | 1.103066 | 0.897095 | -0.29819 | 0.345576 |
| hypothetical protein ABUW_1139 | 0.949764 | 0.918174 | -0.0488 | 0.346307 |
| lipoate-protein ligase B | 0.939267 | 1.047971 | 0.157992 | 0.34719 |
| hypothetical protein ABUW_0949 | 1.003731 | 0.944565 | -0.08765 | 0.348317 |
| inositol-1-monophosphate | 1.064691 | 0.987305 | -0.10887 | 0.348361 |
| AdeS | 1.015205 | 1.054153 | 0.054314 | 0.349575 |
| hypothetical protein ABUW_2270 | 0.975323 | 0.938673 | -0.05526 | 0.349606 |
| FMN-binding protein | 1.002745 | 1.054611 | 0.072756 | 0.350092 |
| ABC-type dipeptide/oligopeptide/nickel transport system per | 0.982306 | 0.912543 | -0.10628 | 0.350173 |
| transcriptional regulator, IclR family...1036 | 0.979862 | 0.941391 | -0.05778 | 0.35019 |
| methionine aminopeptidase, type I...1731 | 1.053937 | 0.986587 | -0.09527 | 0.351158 |
| hypothetical protein ABUW_3528 | 1.067839 | 1.010718 | -0.07931 | 0.351397 |
| bifunctional adenosylcobalamin biosynthesis protein CobP | 0.979499 | 1.040071 | 0.086566 | 0.351471 |
| arginine N-succinyltransferase...61 | 0.965345 | 0.808243 | -0.25626 | 0.354709 |
| rhomboid family protein | 0.975615 | 0.846738 | -0.2044 | 0.355091 |
| alkaline lipase | 1.055089 | 0.958645 | -0.1383 | 0.358174 |
| hypothetical protein ABUW_1241 | 1.131542 | 1.050717 | -0.10692 | 0.359262 |
| monovalent cation transporter | 0.958057 | 0.980665 | 0.033649 | 0.359493 |

| | | | | |
|---|---|---|---|---|
| septum site-determining protein MinC | 0.920693 | 0.986441 | 0.099512 | 0.359907 |
| glyoxalase | 0.996843 | 0.966797 | -0.04415 | 0.362732 |
| general secretion pathway protein...108 | 1.161585 | 0.951603 | -0.28766 | 0.362864 |
| polysaccharide deacetylase...1243 | 0.865635 | 0.90083 | 0.057497 | 0.363493 |
| hypothetical protein ABUW_1657 | 0.907423 | 1.029609 | 0.182249 | 0.363587 |
| 8-amino-7-oxononanoate synthase | 0.974319 | 0.924632 | -0.07552 | 0.363802 |
| hypothetical protein ABUW_2784 | 0.921851 | 0.838922 | -0.136 | 0.364513 |
| oxidoreductase short-chain dehydrogenase/reductase family. | 0.998079 | 0.960114 | -0.05595 | 0.364859 |
| cyclic AMP phosphodiesterase | 1.059873 | 1.110304 | 0.067064 | 0.367017 |
| hypothetical protein ABUW_1193 | 0.973985 | 0.95579 | -0.02721 | 0.367483 |
| metalloprotease | 0.980174 | 0.929602 | -0.07642 | 0.369437 |
| phosphohistidine phosphatase | 0.964646 | 1.02 | 0.080498 | 0.370525 |
| tRNA-dihydrouridine synthase A | 0.992495 | 0.937501 | -0.08224 | 0.371074 |
| hypothetical protein ABUW_2139 | 1.042474 | 1.010121 | -0.04548 | 0.3711 |
| endoribonuclease L-PSP family protein | 1.112039 | 1.026683 | -0.11522 | 0.371726 |
| transcriptional regulator, LysR family...920 | 0.941195 | 0.986041 | 0.067154 | 0.371909 |
| hypothetical protein ABUW_3283 | 1.01296 | 1.050877 | 0.053016 | 0.372833 |
| D-isomer specific 2-hydroxyacid dehydrogenase, NAD-binding | 0.975655 | 1.01134 | 0.051825 | 0.374499 |
| hypothetical protein ABUW_3036 | 0.967292 | 0.906011 | -0.09442 | 0.377175 |
| hypothetical protein ABUW_1327 | 1.022868 | 1.056982 | 0.047331 | 0.377585 |
| 3-hydroxyisobutyrate dehydrogenase...2297 | 1.036261 | 1.002917 | -0.04718 | 0.377655 |
| glutamine amidotransferase | 1.029711 | 1.006317 | -0.03316 | 0.378639 |
| 2,4-dienoyl-CoA reductase | 0.921935 | 0.962467 | 0.062072 | 0.379097 |
| cobalamin adenosyltransferase | 1.06586 | 0.966169 | -0.14167 | 0.379218 |
| hypothetical protein ABUW_0346 | 1.03563 | 1.06234 | 0.036738 | 0.379372 |
| acetyltransferase, gnat family...703 | 1.04804 | 1.017848 | -0.04217 | 0.379386 |
| alanine racemase domain-containing protein | 0.986586 | 0.959115 | -0.04074 | 0.379756 |
| two component signal transduction system kinase sensor con | 0.943022 | 1.015838 | 0.107307 | 0.380025 |
| biotin biosynthesis protein BioH | 1.132991 | 1.07896 | -0.0705 | 0.381398 |
| outer membrane lipoprotein LolB | 0.963703 | 0.933613 | -0.04576 | 0.381664 |
| hypothetical protein ABUW_3003 | 0.98668 | 1.010264 | 0.034078 | 0.381956 |
| peptidyl-prolyl cis-trans isomerase...1216 | 1.023109 | 0.989289 | -0.0485 | 0.383021 |
| hypothetical protein ABUW_0303 | 0.970468 | 1.027549 | 0.082455 | 0.38346 |
| ABC transporter | 1.057176 | 1.118437 | 0.081269 | 0.383756 |
| 3'(2'),5'-bisphosphate nucleotidase | 0.982132 | 1.024422 | 0.060822 | 0.38578 |
| major facilitator superfamily MFS_1...981 | 0.976919 | 1.01068 | 0.049015 | 0.387169 |
| methylcrotonoyl-CoA carboxylase beta chain...2251 | 1.002632 | 0.993107 | -0.01377 | 0.387242 |
| oxidoreductase...2378 | 0.962491 | 0.983003 | 0.030422 | 0.387458 |
| long-chain-fatty-acid--CoA ligase | 1.006608 | 1.028569 | 0.031137 | 0.387515 |
| hypothetical protein ABUW_3366 | 0.896334 | 0.978951 | 0.127199 | 0.38788 |
| replicative DNA helicase...6 | 1.142299 | 1.022457 | -0.1599 | 0.388105 |
| isocitrate dehydrogenase, NADP-dependent...2379 | 1.100027 | 1.056789 | -0.05785 | 0.388268 |
| acyl-CoA synthase...1213 | 1.005477 | 0.938245 | -0.09984 | 0.389516 |
| type IV pilus assembly protein PilM | 2.379214 | 0.283854 | -3.06726 | 0.38988 |
| outer membrane efflux protein | 0.912241 | 0.973413 | 0.093637 | 0.391199 |
| hypothetical protein ABUW_1859 | 0.937376 | 1.035196 | 0.143205 | 0.392733 |
| ribosomal protein S7 | 1.012824 | 0.993364 | -0.02799 | 0.393493 |

| | | | | |
|---|---|---|---|---|
| non-canonical purine NTP pyrophosphatase, HAM1 family | 1.084093 | 0.975421 | -0.15239 | 0.394253 |
| carbamoyl-phosphate synthase | 1.01357 | 1.047488 | 0.047488 | 0.397773 |
| L-lactate utilization transcriptional repressor | 0.979509 | 0.879791 | -0.1549 | 0.398479 |
| aldo/keto reductase...865 | 0.978143 | 0.889031 | -0.13781 | 0.399373 |
| siderophore biosynthesis protein...42 | 1.277991 | 1.159484 | -0.1404 | 0.400268 |
| major facilitator superfamily MFS_1...35 | 0.971658 | 1.069389 | 0.138265 | 0.400799 |
| GMP synthase | 1.036166 | 0.979864 | -0.0806 | 0.403079 |
| transcriptional regulator, TetR family...1326 | 0.998396 | 0.977972 | -0.02982 | 0.403113 |
| hypothetical protein ABUW_2291 | 0.888777 | 1.027284 | 0.208943 | 0.404547 |
| Dyp-type peroxidase | 1.013886 | 0.950785 | -0.0927 | 0.40467 |
| hypothetical protein ABUW_1222 | 1.036008 | 1.062816 | 0.036856 | 0.405409 |
| thymidylate kinase | 0.989696 | 1.022852 | 0.04754 | 0.406997 |
| betaine aldehyde dehydrogenase...1863 | 1.008282 | 0.967363 | -0.05977 | 0.407554 |
| transcriptional regulator, AraC family...20 | 0.914832 | 0.799312 | -0.19475 | 0.408439 |
| thiosulfate-binding protein...130 | 1.036134 | 0.987101 | -0.06994 | 0.412141 |
| co-chaperone GrpE | 0.993689 | 1.038594 | 0.063765 | 0.412744 |
| hypothetical protein ABUW_0972 | 1.052402 | 0.979708 | -0.10326 | 0.413044 |
| glutathione synthase | 0.988973 | 1.01405 | 0.036124 | 0.413276 |
| hypothetical protein ABUW_1719 | 0.985954 | 1.060673 | 0.105388 | 0.413331 |
| transcriptional regulator, ArsR family...472 | 1.056877 | 1.096147 | 0.052634 | 0.413964 |
| hypothetical protein ABUW_1482 | 1.037406 | 1.001288 | -0.05112 | 0.413987 |
| acyl-CoA dehydrogenase...555 | 1.049375 | 0.992369 | -0.08058 | 0.414398 |
| phosphoglucosamine mutase | 1.036209 | 0.989811 | -0.06609 | 0.414586 |
| cation efflux system protein...791 | 0.946655 | 0.936672 | -0.0153 | 0.414729 |
| 5-methyltetrahydropteroyltriglutamate-homocysteine methy | 0.980093 | 1.028749 | 0.0699 | 0.418045 |
| hypothetical protein ABUW_4019 (plasmid) | 0.93071 | 0.968663 | 0.057663 | 0.418473 |
| phage-related membrane protein | 1.043849 | 0.992357 | -0.07298 | 0.420078 |
| glutamate N-acetyltransferase/amino-acid acetyltransferase | 0.9898 | 1.014968 | 0.036225 | 0.421592 |
| disulfide bond formation protein | 0.94624 | 0.883298 | -0.09931 | 0.422064 |
| alpha/beta fold family hydrolase...695 | 0.971401 | 1.025313 | 0.077925 | 0.425271 |
| zinc import ATP-binding protein ZnuC | 0.926794 | 0.988277 | 0.092667 | 0.42707 |
| NADH dehydrogenase I chain M membrane subunit | 0.98855 | 1.036035 | 0.067688 | 0.428271 |
| activator of HSP90 ATPase...1790 | 0.953315 | 0.99198 | 0.057359 | 0.428898 |
| preprotein translocase, YajC subunit | 0.987966 | 1.008706 | 0.029973 | 0.429241 |
| outer membrane protein CarO | 0.974071 | 0.915992 | -0.08869 | 0.432633 |
| pirin domain protein...1290 | 0.997168 | 1.045632 | 0.068467 | 0.433347 |
| hypothetical protein ABUW_0484 | 1.040317 | 0.992927 | -0.06726 | 0.433568 |
| hypothetical protein ABUW_3232 | 1.042129 | 0.995191 | -0.06649 | 0.434152 |
| transcriptional regulator, LysR-type | 0.932483 | 0.949557 | 0.026177 | 0.434342 |
| hypothetical protein ABUW_4005 (plasmid) | 1.003246 | 0.970585 | -0.04775 | 0.439853 |
| phage tail protein | 0.972474 | 0.899099 | -0.11318 | 0.440077 |
| phosphate transporter | 0.919111 | 0.969695 | 0.077291 | 0.440868 |
| bla(GES-14) (plasmid) | 0.919174 | 0.648297 | -0.50368 | 0.441481 |
| glutamate/aspartate transport system permease protein GltK | 0.970912 | 0.90236 | -0.10564 | 0.441599 |
| two-component system histidine kinase sensor component | 1.082162 | 1.007464 | -0.10319 | 0.441678 |
| acyl-CoA ligase | 1.087437 | 1.014269 | -0.10049 | 0.44389 |
| hypothetical protein ABUW_2154 | 0.922653 | 0.975797 | 0.080793 | 0.444144 |

| | | | | |
|---|---|---|---|---|
| RND efflux system, outer membrane lipoprotein, NodT family | 0.971476 | 1.015055 | 0.063307 | 0.444145 |
| oxidoreductase short chain dehydrogenase/reductase family. | 1.080563 | 1.006208 | -0.10285 | 0.445199 |
| hypothetical protein ABUW_2345 | 1.14674 | 1.225303 | 0.0956 | 0.44551 |
| peptidoglycan-associated lipoprotein | 1.015294 | 0.998887 | -0.0235 | 0.445869 |
| DEAD/DEAH box helicase...1763 | 1.002407 | 0.991564 | -0.01569 | 0.446113 |
| short chain dehydrogenase...52 | 1.04729 | 0.997398 | -0.07042 | 0.447048 |
| hypothetical protein ABUW_1600 | 0.924693 | 0.844267 | -0.13127 | 0.447199 |
| phosphoserine aminotransferase | 0.985481 | 1.005521 | 0.029044 | 0.450058 |
| peptidase M61 | 1.004882 | 1.072524 | 0.093984 | 0.452001 |
| succinylglutamic semialdehyde dehydrogenase | 0.951301 | 0.984907 | 0.050086 | 0.452373 |
| hypothetical protein ABUW_3901 | 1.009164 | 1.024789 | 0.022167 | 0.453968 |
| acetyltransferase, gnat family...380 | 0.902029 | 0.94286 | 0.06387 | 0.454578 |
| AFG1-family ATPase | 1.03383 | 1.018309 | -0.02182 | 0.454626 |
| hypothetical protein ABUW_1993 | 1.053137 | 0.999146 | -0.07593 | 0.454853 |
| hypothetical protein ABUW_2219 | 0.940941 | 1.019109 | 0.115133 | 0.454952 |
| hypothetical protein ABUW_1788 | 0.893887 | 0.622242 | -0.52262 | 0.456287 |
| transcriptional regulator, GntR family...308 | 1.059832 | 1.152027 | 0.12034 | 0.460828 |
| hypothetical protein ABUW_2162 | 0.977345 | 1.059225 | 0.116069 | 0.46157 |
| transcriptional regulator, MarR-family...916 | 1.016293 | 0.993312 | -0.033 | 0.462093 |
| phage replication protein | 0.938233 | 0.888628 | -0.07837 | 0.463496 |
| transcriptional regulator, AraC family...651 | 0.889054 | 0.912615 | 0.037735 | 0.463771 |
| oxidoreductase, FAD/FMN-binding | 1.007652 | 1.02477 | 0.024304 | 0.463863 |
| cytochrome bd ubiquinol oxidase, subunit I | 0.917419 | 0.949167 | 0.049081 | 0.463895 |
| hypothetical protein ABUW_0618 | 0.951589 | 0.972679 | 0.031625 | 0.464303 |
| hypothetical protein ABUW_0861 | 0.974561 | 0.918229 | -0.0859 | 0.464493 |
| serine/threonine protein kinase | 1.03901 | 0.958716 | -0.11603 | 0.465049 |
| hypothetical protein ABUW_3717 | 0.997739 | 0.932153 | -0.0981 | 0.465397 |
| polyphosphate-AMP phosphotransferase | 0.982171 | 1.014615 | 0.046886 | 0.465407 |
| putative ribonuclease | 1.023214 | 1.110208 | 0.117722 | 0.466817 |
| Tol-Pal system beta propeller repeat protein TolB | 1.00685 | 0.99123 | -0.02256 | 0.467894 |
| hypothetical protein ABUW_0459 | 1.053968 | 1.030598 | -0.03235 | 0.468436 |
| lipase foldase | 1.077136 | 1.014282 | -0.08674 | 0.469671 |
| DNA primase | 0.948256 | 0.932615 | -0.024 | 0.471241 |
| metallo-beta-lactamase superfamily protein | 0.892903 | 0.942248 | 0.077603 | 0.471487 |
| PpiC-type peptidyl-prolyl cis-trans isomerase | 0.987497 | 0.972854 | -0.02155 | 0.471501 |
| hypothetical protein ABUW_2553 | 1.082266 | 0.976368 | -0.14856 | 0.472107 |
| hypothetical protein ABUW_3612 | 0.983757 | 1.03106 | 0.067754 | 0.472836 |
| transcriptional regulator, XRE family...813 | 1.066747 | 0.987632 | -0.11117 | 0.472857 |
| hypothetical protein ABUW_2832 | 0.879708 | 0.77572 | -0.18149 | 0.474526 |
| LysR-family transcriptional regulator | 1.013504 | 1.031595 | 0.025525 | 0.476186 |
| transcription elongation factor GreB | 1.089578 | 1.060532 | -0.03898 | 0.476342 |
| phosphoglycerate mutase...1046 | 1.007451 | 0.97353 | -0.04941 | 0.476812 |
| iron-sulfur cluster assembly protein IscA | 1.018409 | 0.950222 | -0.09998 | 0.477834 |
| two-component system sensor protein...160 | 0.941783 | 1.012601 | 0.1046 | 0.478723 |
| catechol 1,2-dioxygenase | 1.031452 | 1.065206 | 0.046457 | 0.479471 |
| DNA polymerase III, epsilon subunit | 1.008084 | 1.048509 | 0.056723 | 0.480796 |
| hypothetical protein ABUW_3587 | 1.026528 | 0.985427 | -0.05895 | 0.481084 |

| | | | | |
|---|---|---|---|---|
| hypothetical protein ABUW_2874 | 1.00953 | 1.000122 | -0.01351 | 0.484767 |
| replication protein (plasmid) | 0.983725 | 0.947357 | -0.05435 | 0.485228 |
| 3-oxoacyl-[acyl-carrier-protein] synthase 1 | 1.041905 | 1.00552 | -0.05128 | 0.48541 |
| oxidoreductase alpha (molybdopterin) subunit | 0.970054 | 0.941256 | -0.04348 | 0.485818 |
| AraC family transcriptional regulator | 1.033794 | 0.999339 | -0.0489 | 0.489466 |
| cytochrome O ubiquinol oxidase subunit II | 0.973487 | 1.0141 | 0.058965 | 0.489953 |
| prolipoprotein diacylglyceryl transferase | 0.983329 | 0.966541 | -0.02484 | 0.490806 |
| LysR family transcriptional regulator | 1.006287 | 1.029472 | 0.032862 | 0.491793 |
| transcriptional regulator, ModE family | 0.959947 | 0.940952 | -0.02883 | 0.491799 |
| aldehyde dehydrogenase...633 | 0.987134 | 1.051587 | 0.091251 | 0.492608 |
| transcriptional regulator, GntR family...980 | 0.95566 | 0.931685 | -0.03666 | 0.492838 |
| glycosyl transferase, family 3 | 0.99082 | 1.00693 | 0.023269 | 0.494078 |
| polysaccharide export protein | 0.990067 | 0.975098 | -0.02198 | 0.494577 |
| urease accessory protein G | 0.940108 | 0.996222 | 0.083642 | 0.496339 |
| glycerophosphoryl diester phosphodiesterase | 0.999646 | 1.017065 | 0.024923 | 0.497224 |
| acyl-CoA dehydrogenase...97 | 1.065813 | 0.999533 | -0.09263 | 0.498527 |
| membrane-associated Zn-dependent protease | 1.061232 | 0.976949 | -0.11939 | 0.498879 |
| hypothetical protein ABUW_3102 | 1.138906 | 1.212177 | 0.089952 | 0.499029 |
| hypothetical protein ABUW_2677 | 1.052334 | 0.920959 | -0.19238 | 0.499303 |
| putative RND family drug transporter...1367 | 0.987797 | 0.944534 | -0.06461 | 0.499566 |
| anthranilate synthase component I | 0.935229 | 0.987695 | 0.078746 | 0.507149 |
| phospholipase D/transphosphatidylase | 0.994353 | 1.007895 | 0.019515 | 0.50816 |
| type IV / VI secretion system protein, DotU family | 0.950512 | 1.022237 | 0.104954 | 0.508535 |
| hypothetical protein ABUW_1086 | 0.96996 | 0.931924 | -0.05771 | 0.510698 |
| lysozyme-like domain-containing protein | 0.904801 | 1.091239 | 0.270296 | 0.513495 |
| isovaleryl-CoA dehydrogenase | 1.008554 | 1.026552 | 0.025518 | 0.515577 |
| 2-dehydro-3-deoxyphosphooctonate aldolase | 1.01007 | 0.978786 | -0.04539 | 0.516666 |
| glutamate 5-kinase | 1.040262 | 1.007023 | -0.04685 | 0.517703 |
| microcin B17 transport protein | 0.926964 | 0.954122 | 0.041661 | 0.518503 |
| histidine triad protein...1546 | 1.045719 | 1.023295 | -0.03127 | 0.520341 |
| trans-hexaprenyltranstransferase | 0.996474 | 1.017829 | 0.030591 | 0.521033 |
| pyridine nucleotide-disulfide oxidoreductase | 1.06428 | 1.003144 | -0.08535 | 0.521921 |
| succinylarginine dihydrolase | 1.004138 | 0.990983 | -0.01902 | 0.523405 |
| deoxyribodipyrimidine photo-lyase | 0.998168 | 1.098206 | 0.137794 | 0.524005 |
| hypothetical protein ABUW_1809 | 0.95405 | 0.913058 | -0.06336 | 0.525205 |
| acetyltransferase, gnat family...1760 | 1.01643 | 1.053714 | 0.051972 | 0.52534 |
| hypothetical protein ABUW_1414 | 1.249238 | 1.16937 | -0.09532 | 0.526037 |
| hypothetical protein ABUW_0822 | 0.922612 | 0.967621 | 0.068718 | 0.527256 |
| oxidoreductase short-chain dehydrogenase/reductase family. | 0.981902 | 0.942016 | -0.05983 | 0.527479 |
| ribonuclease G | 0.957859 | 0.968532 | 0.015986 | 0.528661 |
| lipoic acid synthetase...155 | 1.153219 | 0.973702 | -0.24411 | 0.528861 |
| hypothetical protein ABUW_1217 | 1.00675 | 0.997637 | -0.01312 | 0.529948 |
| hypothetical protein ABUW_0549 | 1.00648 | 1.057811 | 0.071764 | 0.534754 |
| hypothetical protein ABUW_2995 | 0.886632 | 0.922658 | 0.05746 | 0.535913 |
| heme oxygenase-like protein...1563 | 0.94403 | 0.978858 | 0.052267 | 0.536939 |
| transcriptional regulator, LysR family...1242 | 0.977942 | 0.959301 | -0.02777 | 0.537536 |
| glutamate dehydrogenase | 0.981588 | 1.007775 | 0.037984 | 0.539811 |

| | | | | |
|---|---|---|---|---|
| type VI secretion protein, family...784 | 0.929909 | 0.98126 | 0.077546 | 0.540291 |
| malate synthase G | 0.996741 | 1.013481 | 0.024028 | 0.541372 |
| two-component system sensor protein...816 | 1.011645 | 1.032665 | 0.029669 | 0.542995 |
| hypothetical protein ABUW_0863 | 1.021105 | 0.9869 | -0.04916 | 0.545697 |
| transketolase | 1.011607 | 0.98893 | -0.03271 | 0.548764 |
| hypothetical protein ABUW_1653 | 1.047196 | 1.021449 | -0.03591 | 0.550113 |
| ben operon transcriptional regulator BenM | 1.016501 | 0.996403 | -0.02881 | 0.552505 |
| hypothetical protein ABUW_4007 (plasmid) | 0.962006 | 0.931476 | -0.04653 | 0.553988 |
| hypothetical protein ABUW_0588 | 1.025113 | 1.002321 | -0.03244 | 0.554032 |
| ATP synthase F1, epsilon subunit | 1.007842 | 1.022923 | 0.021427 | 0.554037 |
| diaminobutyrate--2-oxoglutarate aminotransferase | 0.995659 | 1.021245 | 0.036605 | 0.555299 |
| acyl-CoA dehydrogenase...721 | 0.934424 | 0.999733 | 0.097465 | 0.557527 |
| undecaprenyl-diphosphatase UppP | 0.991968 | 0.948804 | -0.06418 | 0.558257 |
| fatty acid desaturase...256 | 1.11774 | 1.14128 | 0.030069 | 0.558379 |
| hypothetical protein ABUW_2163 | 0.997063 | 0.991306 | -0.00835 | 0.558508 |
| hypothetical protein ABUW_3623 | 0.978011 | 0.923703 | -0.08242 | 0.558902 |
| arsenical resistance protein ArsH | 0.97993 | 1.006053 | 0.037955 | 0.559872 |
| hypothetical protein ABUW_4106 (plasmid) | 1.016841 | 1.134579 | 0.158064 | 0.560769 |
| transcriptional regulator, TetR family...693 | 0.933412 | 0.967869 | 0.052297 | 0.561665 |
| hypothetical protein ABUW_1753 | 0.965904 | 1.006234 | 0.059015 | 0.56254 |
| transcriptional regulator, LysR family...943 | 1.013793 | 1.036525 | 0.031992 | 0.564229 |
| major capsid protein | 0.942221 | 1.098464 | 0.22135 | 0.565293 |
| transcriptional regulator, TetR family...89 | 1.043805 | 0.970383 | -0.10523 | 0.567839 |
| riboflavin biosynthesis protein RibF | 0.959837 | 0.981575 | 0.032309 | 0.568585 |
| toluene tolerance efflux transporter...1509 | 0.993385 | 0.977889 | -0.02268 | 0.569207 |
| beta-ketoadipyl CoA thiolase...930 | 1.057614 | 1.036857 | -0.0286 | 0.570022 |
| NADPH dehydrogenase | 1.034343 | 1.026114 | -0.01152 | 0.570192 |
| molybdopterin oxidoreductase...156 | 0.76285 | 0.903437 | 0.244025 | 0.570333 |
| phosphoenolpyruvate synthase | 1.021272 | 1.038701 | 0.024413 | 0.570837 |
| hypothetical protein ABUW_2940 | 0.969525 | 1.039116 | 0.100007 | 0.574269 |
| RND family drug transporter | 0.825933 | 0.855675 | 0.051039 | 0.57435 |
| alpha/beta hydrolase fold protein...38 | 0.86042 | 0.889643 | 0.048186 | 0.575904 |
| short-chain dehydrogenase/reductase...116 | 0.890481 | 0.832319 | -0.09745 | 0.576669 |
| hypothetical protein ABUW_3021 | 0.91074 | 0.951187 | 0.062691 | 0.579239 |
| ribosomal protein S8 | 1.022446 | 1.007615 | -0.02108 | 0.58011 |
| hypothetical protein ABUW_2549 | 0.959788 | 0.931739 | -0.04279 | 0.581164 |
| transcriptional regulator, AsnC family...898 | 0.987005 | 1.01659 | 0.042608 | 0.582795 |
| rhomboid family peptidase | 1.047778 | 1.004902 | -0.06028 | 0.583479 |
| NDP-sugar dehydrogenase | 0.979793 | 0.955815 | -0.03575 | 0.583864 |
| inosine-5'-monophosphate dehydrogenase | 0.980318 | 1.012731 | 0.04693 | 0.583878 |
| permease...1176 | 0.967926 | 0.980983 | 0.019332 | 0.584446 |
| hypothetical protein ABUW_6003 (plasmid) | 0.997309 | 0.964838 | -0.04775 | 0.584453 |
| succinyl-diaminopimelate desuccinylase | 1.034131 | 1.073021 | 0.053259 | 0.584906 |
| hypothetical protein ABUW_0737 | 1.077104 | 1.001663 | -0.10476 | 0.585599 |
| Peptidase M20D, amidohydrolase | 1.034922 | 1.104371 | 0.093703 | 0.587489 |
| hypothetical protein ABUW_1685 | 1.012493 | 1.028868 | 0.023146 | 0.58846 |
| lytic transglycosylase, catalytic | 1.009495 | 0.997246 | -0.01761 | 0.592284 |

| | | | | |
|---|---|---|---|---|
| DNA cytosine methyltransferase | 0.971074 | 0.996122 | 0.036741 | 0.592696 |
| succinate dehydrogenase, cytochrome b556 subunit | 0.977558 | 0.925427 | -0.07906 | 0.592878 |
| nicotinamide-nucleotide adenylyltransferase | 0.993913 | 0.98142 | -0.01825 | 0.594802 |
| hypothetical protein ABUW_2383 | 1.151779 | 1.054013 | -0.12797 | 0.595289 |
| indole-3-glycerol phosphate synthase | 0.955636 | 0.982423 | 0.039882 | 0.596072 |
| TRAP C4-dicarboxylate transport system permease | 0.943214 | 0.921692 | -0.0333 | 0.597224 |
| hypothetical protein ABUW_3701 | 0.98262 | 0.966313 | -0.02414 | 0.599768 |
| hypothetical protein ABUW_0438 | 0.945379 | 0.99695 | 0.076629 | 0.599817 |
| ribosomal protein L9 | 1.028763 | 1.010957 | -0.02519 | 0.599996 |
| long-chain fatty acid transport protein | 1.127265 | 1.04144 | -0.11425 | 0.600765 |
| oxidoreductase...140 | 1.19401 | 0.992939 | -0.26604 | 0.600976 |
| translation elongation factor P | 1.03277 | 0.995713 | -0.05272 | 0.601357 |
| orotidine 5'-phosphate decarboxylase | 1.002105 | 1.01541 | 0.019028 | 0.601923 |
| hypothetical protein ABUW_0837 | 0.909114 | 0.9416 | 0.050653 | 0.602996 |
| capsid scaffolding protein | 0.923966 | 0.952802 | 0.044337 | 0.603289 |
| MutT/NUDIX family protein | 0.798354 | 0.828693 | 0.05381 | 0.603362 |
| hypothetical protein ABUW_0243 | 0.946821 | 0.889267 | -0.09047 | 0.603666 |
| sigma-54 specific, transcriptional regulator, Fis family | 1.012051 | 1.033044 | 0.02962 | 0.603666 |
| hypothetical protein ABUW_1989 | 1.127511 | 1.021238 | -0.14282 | 0.603993 |
| ribose 5-phosphate isomerase | 1.019203 | 1.011507 | -0.01094 | 0.60446 |
| putative helicase | 0.986287 | 1.007602 | 0.030847 | 0.604602 |
| hypothetical protein ABUW_3090 | 1.057698 | 1.028422 | -0.0405 | 0.604973 |
| hypothetical protein ABUW_1471 | 0.872618 | 0.772974 | -0.17493 | 0.605288 |
| hypothetical protein ABUW_1324 | 1.181453 | 1.136438 | -0.05604 | 0.606478 |
| putative phosphodiesterase | 0.959225 | 0.947343 | -0.01798 | 0.606952 |
| hypothetical protein ABUW_2559 | 1.018779 | 1.000365 | -0.02631 | 0.608429 |
| FAD-dependent pyridine nucleotide- disulfide oxidoreductase | 0.932346 | 0.957127 | 0.037844 | 0.608967 |
| bacterioferritin...2226 | 0.975778 | 1.01526 | 0.057224 | 0.611632 |
| ferrochelatase | 0.995247 | 1.013904 | 0.026794 | 0.611697 |
| hypothetical protein ABUW_1349 | 0.981968 | 1.025697 | 0.062857 | 0.613598 |
| hypothetical protein ABUW_1903 | 1.042895 | 0.989986 | -0.07511 | 0.613853 |
| hypothetical protein ABUW_3027 | 0.898492 | 0.933092 | 0.054514 | 0.615245 |
| Putative tellurique resistant protein (plasmid) | 0.999554 | 0.954264 | -0.0669 | 0.616721 |
| NAD-dependent deacetylase regulatory protein | 1.010378 | 0.96686 | -0.06352 | 0.61964 |
| hypothetical protein ABUW_3543 | 0.972305 | 0.995202 | 0.033579 | 0.620372 |
| aspartate 1-decarboxylase | 1.063028 | 1.039295 | -0.03257 | 0.621176 |
| hypothetical protein ABUW_0787 | 0.95888 | 1.01249 | 0.078484 | 0.621279 |
| 3-oxoacyl-[acyl-carrier-protein] reductase...261 | 1.065221 | 1.031834 | -0.04594 | 0.621664 |
| DNA-binding protein (plasmid) | 0.989695 | 1.032947 | 0.06171 | 0.622202 |
| membrane protein involved in aromatic hydrocarbon degrada | 1.000975 | 1.017069 | 0.023012 | 0.622535 |
| OmpW family protein...1531 | 1.006297 | 1.038213 | 0.045047 | 0.622617 |
| NADP-dependent fatty aldehyde dehydrogenase...1507 | 0.961662 | 0.973804 | 0.018101 | 0.623084 |
| 2Fe-2S iron-sulfur cluster binding domain protein | 1.048829 | 1.067606 | 0.0256 | 0.626187 |
| potassium transport system low affinity (KUP family) | 0.961467 | 0.974807 | 0.019879 | 0.628166 |
| metallopeptidase, zinc binding | 0.974764 | 0.969684 | -0.00754 | 0.629379 |
| putative flavoprotein | 0.997549 | 0.979475 | -0.02638 | 0.629696 |
| deoxyguanosinetriphosphate triphosphohydrolase | 1.017537 | 0.992812 | -0.03549 | 0.629872 |

| | | | | |
|---|---|---|---|---|
| phosphoglycerate mutase...732 | 0.98234 | 0.969507 | -0.01897 | 0.629992 |
| TonB-dependent receptor protein (plasmid) | 1.024244 | 1.004834 | -0.0276 | 0.630191 |
| thioredoxin...1316 | 1.009223 | 0.9878 | -0.03095 | 0.634496 |
| sulfate adenylyltransferase subunit 1 | 1.01348 | 1.022245 | 0.012423 | 0.634793 |
| 5-dehydro-4-deoxyglucarate dehydratase | 1.005129 | 0.990434 | -0.02125 | 0.63484 |
| hypothetical protein ABUW_1240 | 0.882322 | 0.949908 | 0.106482 | 0.634931 |
| hypothetical protein ABUW_1844 | 1.03614 | 1.010822 | -0.03569 | 0.63538 |
| porin B...2388 | 0.995384 | 1.004315 | 0.012887 | 0.636992 |
| phage-associated protein, family | 1.007755 | 1.031126 | 0.033075 | 0.638011 |
| ubiquinone biosynthesis protein COQ7 | 0.995444 | 1.012886 | 0.02506 | 0.642765 |
| arsenical resistance operon repressor | 1.040991 | 1.107755 | 0.089682 | 0.643212 |
| transposition helper protein C | 0.993329 | 0.978089 | -0.02231 | 0.643356 |
| beta-ketoadipyl CoA thiolase...203 | 0.896945 | 0.925161 | 0.044685 | 0.645604 |
| GTP-binding protein...1376 | 0.977091 | 0.986896 | 0.014405 | 0.64565 |
| betaine aldehyde dehydrogenase...1043 | 0.985574 | 0.963125 | -0.03324 | 0.646208 |
| methionine adenosyltransferase | 1.009873 | 0.974755 | -0.05106 | 0.649001 |
| acyl-CoA N-acyltransferase | 0.986066 | 1.017433 | 0.045178 | 0.649254 |
| pantetheine-phosphate adenylyltransferase | 1.024675 | 0.995109 | -0.04224 | 0.652175 |
| beta-lactamase ADC7 | 1.01139 | 1.019009 | 0.010827 | 0.652543 |
| hypothetical protein ABUW_1718 | 0.88178 | 0.889852 | 0.013147 | 0.6528 |
| hydroxyethylthiazole kinase | 0.975327 | 0.958979 | -0.02439 | 0.653617 |
| thioesterase superfamily protein...709 | 1.075214 | 1.029005 | -0.06337 | 0.656072 |
| acetyl-CoA acetyltransferase...1657 | 1.034666 | 1.013693 | -0.02954 | 0.657171 |
| hypothetical protein ABUW_1631 | 1.008332 | 0.991545 | -0.02422 | 0.658658 |
| 3-methylglutaconyl-CoA hydratase | 0.994037 | 1.019485 | 0.036469 | 0.65874 |
| formate/nitrite transporter family | 0.935471 | 0.919031 | -0.02558 | 0.658988 |
| lytic murein transglycosylase B | 1.022771 | 1.016042 | -0.00952 | 0.662412 |
| thiamine-monophosphate kinase | 1.013361 | 1.056659 | 0.060361 | 0.664085 |
| DNA-3-methyladenine glycosylase | 0.984793 | 1.043301 | 0.083263 | 0.664122 |
| chorismate synthase | 0.981505 | 0.963776 | -0.0263 | 0.664725 |
| phenylacetate-CoA oxygenase, PaaH subunit | 0.89321 | 0.923536 | 0.048169 | 0.664781 |
| glycosyl transferase, group 1 | 0.981527 | 0.99469 | 0.01922 | 0.666831 |
| hypothetical protein ABUW_0460 | 1.017534 | 1.033506 | 0.02247 | 0.669106 |
| SPOUT methyltransferase | 1.026534 | 1.010846 | -0.02222 | 0.671022 |
| K+ uptake system component | 0.999941 | 1.03776 | 0.053558 | 0.671477 |
| tetraacyldisaccharide 4'-kinase | 1.024887 | 1.044874 | 0.027864 | 0.672591 |
| tyrosine recombinase XerD | 1.025706 | 1.012977 | -0.01802 | 0.673214 |
| ABC transporter permease protein | 0.988065 | 0.976976 | -0.01628 | 0.673972 |
| hypothetical protein ABUW_1083 | 0.988066 | 0.960905 | -0.04021 | 0.673986 |
| nitrogen metabolism transcriptional regulator, NtrC, Fis Famil | 0.981298 | 0.964683 | -0.02464 | 0.674632 |
| pseudouridine synthase | 0.974648 | 0.987786 | 0.019316 | 0.675762 |
| pyruvate decarboxylase E1 component | 1.023639 | 1.010715 | -0.01833 | 0.676795 |
| tRNA modification GTPase TrmE | 0.97481 | 0.996178 | 0.031282 | 0.678773 |
| acyl-CoA dehydrogenase...1841 | 0.949926 | 0.959457 | 0.014404 | 0.682379 |
| glutathione S-transferase family protein...688 | 0.995899 | 1.009536 | 0.019622 | 0.68319 |
| acyl carrier protein...44 | 0.972933 | 0.860915 | -0.17647 | 0.68674 |
| nitroreductase...1514 | 1.023947 | 1.034434 | 0.0147 | 0.687505 |

| | | | | |
|---|---|---|---|---|
| crispr-associated protein, Csy1 family | 0.927956 | 0.918643 | -0.01455 | 0.68868 |
| acyl-CoA dehydrogenase...519 | 0.934706 | 0.95599 | 0.032483 | 0.690751 |
| two-component system hybrid histidine kinase/response regu | 0.974499 | 0.997864 | 0.034182 | 0.69096 |
| transporter, drug/metabolite exporter family | 0.994394 | 0.963352 | -0.04576 | 0.692425 |
| acyl-CoA dehydrogenase...543 | 0.994532 | 0.968431 | -0.03837 | 0.694844 |
| hypothetical protein ABUW_2830 | 0.91363 | 0.945025 | 0.048742 | 0.695627 |
| multidrug efflux protein AdeJ | 0.963174 | 0.97071 | 0.011245 | 0.695632 |
| electron transfer flavoprotein-ubiquinone oxidoreductase (Es | 0.996303 | 1.00283 | 0.009421 | 0.69576 |
| GTP-binding protein TypA/BipA | 0.996364 | 1.026159 | 0.04251 | 0.696265 |
| hypothetical protein ABUW_0991 | 1.057248 | 1.042148 | -0.02075 | 0.696984 |
| 1-deoxy-D-xylulose 5-phosphate reductoisomerase | 1.036125 | 1.022582 | -0.01898 | 0.698625 |
| hypothetical protein ABUW_3494 | 1.008557 | 1.037608 | 0.040969 | 0.700429 |
| DnaK suppressor protein | 1.021848 | 1.003889 | -0.02558 | 0.700679 |
| ribosomal RNA small subunit methyltransferase C | 0.968753 | 0.935424 | -0.05051 | 0.701954 |
| HlyD family secretion protein | 1.167295 | 1.239432 | 0.086509 | 0.704186 |
| aromatic amino acid transport protein | 1.055293 | 1.08053 | 0.034096 | 0.706252 |
| oxidoreductase...43 | 1.16613 | 1.091096 | -0.09595 | 0.706682 |
| malonate decarboxylase, beta subunit | 1.118054 | 1.104978 | -0.01697 | 0.708757 |
| transcriptional regulator, LysR family...595 | 1.077366 | 1.086338 | 0.011964 | 0.709371 |
| 3-dehydroquinate synthase | 1.026297 | 1.035059 | 0.012265 | 0.709489 |
| superoxide dismutase (Fe) | 1.029609 | 1.019191 | -0.01467 | 0.710645 |
| glutaredoxin | 0.979039 | 0.99189 | 0.018813 | 0.710826 |
| dihydrolipoamide dehydrogenase...2456 | 1.001084 | 1.033037 | 0.045328 | 0.710911 |
| phosphoserine phosphatase...8 | 1.071472 | 1.125279 | 0.07069 | 0.711618 |
| inner membrane symporter YgjU | 0.948276 | 0.96659 | 0.027598 | 0.712279 |
| tRNA (uracil-5-)-methyltransferase | 0.959951 | 0.946915 | -0.01972 | 0.712435 |
| ribosomal protein S4 | 1.006429 | 0.998794 | -0.01099 | 0.717296 |
| hypothetical protein ABUW_1994 | 1.053228 | 1.026392 | -0.03724 | 0.717863 |
| ammonium transporter...7 | 1.039306 | 0.987971 | -0.07308 | 0.720067 |
| hypothetical protein ABUW_0518 | 1.003404 | 0.990975 | -0.01798 | 0.721783 |
| penicillin-binding protein 2 | 1.020753 | 0.9896 | -0.04472 | 0.722565 |
| aldo-keto reductase | 1.051873 | 1.104067 | 0.069868 | 0.723835 |
| transcriptional regulator, TetR family...637 | 0.953697 | 0.983429 | 0.04429 | 0.72502 |
| ISPpu12 transposase | 0.947011 | 0.912829 | -0.05304 | 0.725852 |
| flavohemoprotein | 0.989342 | 0.998188 | 0.012842 | 0.726567 |
| transglycosylase-associated protein | 1.012691 | 1.060182 | 0.066119 | 0.726971 |
| hypothetical protein ABUW_0728 | 1.00378 | 0.989388 | -0.02084 | 0.72827 |
| peptidase S24 S26A and S26B...191 | 1.081846 | 1.106292 | 0.032236 | 0.7284 |
| hypothetical protein ABUW_2050 | 0.969491 | 0.993468 | 0.035246 | 0.730148 |
| beta-hydroxylase | 1.058562 | 1.07886 | 0.027402 | 0.730612 |
| two-component system response regulator protein...2286 | 1.008233 | 0.9998 | -0.01212 | 0.732272 |
| glycerol-3-phosphate dehydrogenase | 0.971058 | 0.9547 | -0.02451 | 0.732285 |
| hypothetical protein ABUW_1503 | 0.961786 | 0.987464 | 0.038012 | 0.733102 |
| hypothetical protein ABUW_4038 (plasmid) | 0.947625 | 0.933112 | -0.02227 | 0.733311 |
| electron transfer flavoprotein subunit alpha | 1.01957 | 1.022029 | 0.003476 | 0.733519 |
| hypothetical protein ABUW_1591 | 0.994974 | 0.992589 | -0.00346 | 0.734407 |
| histidinol dehydrogenase | 0.953404 | 0.94676 | -0.01009 | 0.735295 |

| | | | | |
|---|---|---|---|---|
| cytochrome D ubiquinol oxidase, subunit II...235 | 1.014937 | 0.993403 | -0.03094 | 0.736638 |
| peptidase S24 S26A and S26B...1028 | 0.964497 | 0.977545 | 0.019386 | 0.73785 |
| bis(5'-nucleosyl)-tetraphosphatase | 0.97664 | 0.981689 | 0.00744 | 0.739778 |
| phenylacetate-CoA oxygenase/reductase, PaaK subunit | 0.940281 | 0.916985 | -0.03619 | 0.740868 |
| hypothetical protein ABUW_1695 | 0.959613 | 0.978227 | 0.027715 | 0.740965 |
| oligoribonuclease Orn | 0.9943 | 1.006765 | 0.017975 | 0.744047 |
| L-sorbosone dehydrogenase...2085 | 0.941002 | 0.932223 | -0.01352 | 0.744101 |
| 3-carboxy-cis,cis-muconate cycloisomerase | 1.047375 | 1.031825 | -0.02158 | 0.74434 |
| phage-related terminase | 1.012621 | 1.042414 | 0.041834 | 0.745354 |
| transcriptional regulator, LysR family...588 | 0.985997 | 0.979386 | -0.00971 | 0.746501 |
| hypothetical protein ABUW_0856 | 0.971355 | 0.947406 | -0.03602 | 0.747215 |
| non-ribosomal peptide synthetase...40 | 1.14608 | 1.10557 | -0.05192 | 0.75037 |
| muconolactone delta-isomerase | 1.014392 | 1.030971 | 0.023388 | 0.750712 |
| molybdopterin biosynthesis protein...1083 | 1.015288 | 1.019307 | 0.005699 | 0.75207 |
| methyltransferase type 12 | 0.877539 | 0.892268 | 0.024014 | 0.752597 |
| two-component system regulatory protein KdpE | 1.063524 | 1.046858 | -0.02279 | 0.752735 |
| methylcrotonoyl-CoA carboxylase beta chain...103 | 1.191998 | 1.148551 | -0.05357 | 0.753442 |
| hypothetical protein ABUW_1447 | 1.006116 | 0.995987 | -0.0146 | 0.753897 |
| 4'-phosphopantetheinyl transferase | 1.070627 | 0.917322 | -0.22295 | 0.753993 |
| auxin-responsive GH3-related protein | 1.002519 | 0.967768 | -0.0509 | 0.755912 |
| ribosomal protein L2 | 0.987496 | 0.998391 | 0.01583 | 0.756689 |
| ABC-1 domain protein...1828 | 0.960864 | 0.951893 | -0.01353 | 0.757927 |
| hypothetical protein ABUW_2063 | 0.972289 | 0.984165 | 0.017515 | 0.761376 |
| hypothetical protein ABUW_2495 | 0.98752 | 0.969495 | -0.02658 | 0.768163 |
| TonB-dependent siderophore receptor...630 | 1.028279 | 1.042825 | 0.020265 | 0.771061 |
| benzoate 1,2-dioxygenase, small subunit | 0.972845 | 0.940288 | -0.04911 | 0.771108 |
| xanthine dehydrogenase, molybdopterin binding subunit | 1.108521 | 1.057538 | -0.06793 | 0.771137 |
| hypothetical protein ABUW_1323 | 0.937814 | 0.953127 | 0.023368 | 0.772789 |
| TonB-dependent siderophore receptor...2197 | 0.985555 | 0.977059 | -0.01249 | 0.773677 |
| ABC transporter, ATP-binding protein...2066 | 0.973148 | 0.9647 | -0.01258 | 0.77431 |
| hypothetical protein ABUW_1632 | 0.791247 | 0.76312 | -0.05222 | 0.774713 |
| ribonuclease PH | 1.025517 | 1.040296 | 0.020642 | 0.776849 |
| hypothetical protein ABUW_2700 | 0.972248 | 0.957375 | -0.02224 | 0.776949 |
| hypothetical protein ABUW_0144 | 0.921369 | 0.932221 | 0.016893 | 0.779522 |
| acetyl-/propionyl-coenzyme A carboxylase alpha chain | 1.059368 | 1.047789 | -0.01585 | 0.781865 |
| methionine-S-sulfoxide reductase | 0.974106 | 0.966663 | -0.01107 | 0.782278 |
| putative competence protein | 1.008197 | 1.000347 | -0.01128 | 0.783576 |
| alpha/beta hydrolase...54 | 1.040503 | 1.022292 | -0.02547 | 0.785886 |
| hypothetical protein ABUW_1129 | 0.955243 | 0.970994 | 0.023595 | 0.789049 |
| nitrogen regulatory protein P-II | 1.00755 | 1.01285 | 0.00757 | 0.789358 |
| hypothetical protein ABUW_1311 | 0.905645 | 0.880374 | -0.04083 | 0.791155 |
| oxygenase | 1.006132 | 1.034388 | 0.039957 | 0.7955 |
| response regulator | 0.998869 | 1.007725 | 0.012736 | 0.803306 |
| hypothetical protein ABUW_3387 | 0.992193 | 0.997861 | 0.008219 | 0.804151 |
| hypothetical protein ABUW_0597 | 1.024236 | 1.005917 | -0.02604 | 0.805638 |
| ribonuclease D | 1.186252 | 1.171056 | -0.0186 | 0.806486 |
| hypothetical protein ABUW_3611 | 0.971065 | 0.98369 | 0.018636 | 0.806767 |

| | | | | |
|---|---|---|---|---|
| ABC transporter, methionine-binding protein | 0.982684 | 0.977193 | -0.00808 | 0.806902 |
| peptide deformylase...1954 | 0.999994 | 0.995287 | -0.00681 | 0.809705 |
| NADH dehydrogenase I chain A | 1.024777 | 1.004856 | -0.02832 | 0.810834 |
| hypothetical protein ABUW_0866 | 0.972451 | 0.962149 | -0.01537 | 0.811585 |
| heat shock protein 15 | 0.991157 | 0.964773 | -0.03893 | 0.811911 |
| phage protein...123 | 0.916639 | 0.933328 | 0.026029 | 0.813647 |
| translation elongation factor Ts | 1.015601 | 1.021196 | 0.007926 | 0.814786 |
| Fe(II) trafficking protein | 1.017742 | 1.022 | 0.006024 | 0.81557 |
| iron-sulfur cluster-binding protein, Rieske family | 0.979776 | 0.949261 | -0.04565 | 0.81666 |
| GMC oxidoreductase | 0.784649 | 0.764312 | -0.03789 | 0.818598 |
| short-chain dehydrogenase/reductase SDR | 0.94177 | 0.925645 | -0.02492 | 0.819306 |
| signal recognition particle-docking protein FtsY | 1.006004 | 0.998016 | -0.0115 | 0.819958 |
| secretion chaperone | 1.02483 | 1.012639 | -0.01726 | 0.823621 |
| hypothetical protein ABUW_0104 | 0.992004 | 1.002931 | 0.015805 | 0.824035 |
| phenazine biosynthesis protein | 0.988477 | 0.99477 | 0.009156 | 0.824866 |
| transcriptional regulator, GntR family...507 | 1.019681 | 1.027746 | 0.011365 | 0.825013 |
| DNA repair system | 1.04112 | 1.019833 | -0.0298 | 0.825324 |
| hypothetical protein ABUW_0336 | 1.007946 | 1.02621 | 0.025908 | 0.827481 |
| ubiquinone/menaquinone biosynthesis methyltransferase Ub | 0.996604 | 1.00263 | 0.008696 | 0.829538 |
| thiamine S/molybdopterin converting factor subunit 1 | 0.843042 | 0.850885 | 0.01336 | 0.83164 |
| thioredoxin-disulfide reductase...1897 | 0.984551 | 0.98868 | 0.006038 | 0.832136 |
| N-acetyl-beta-glucosaminidase | 1.007205 | 0.99751 | -0.01395 | 0.837248 |
| guanosine-3',5'-bis(diphosphate) 3'-pyrophosphohydrolase (( | 1.007326 | 0.981238 | -0.03786 | 0.838328 |
| hypothetical protein ABUW_1824 | 1.028224 | 1.015619 | -0.01779 | 0.838421 |
| tRNA (guanine-N(7)-)-methyltransferase | 0.954188 | 0.963927 | 0.014651 | 0.838838 |
| alginate biosynthesis protein | 0.89285 | 0.92156 | 0.045661 | 0.844316 |
| acetate kinase | 1.011858 | 1.018838 | 0.009919 | 0.845766 |
| molybdate ABC transporter, periplasmic molybdate-binding p | 1.088106 | 1.077952 | -0.01353 | 0.847486 |
| signal recognition particle protein | 0.993795 | 0.997563 | 0.00546 | 0.855226 |
| phosphate regulon sensor kinase PhoR | 0.975379 | 0.977385 | 0.002965 | 0.857675 |
| malonate decarboxylase, alpha subunit | 0.948266 | 0.958808 | 0.01595 | 0.858768 |
| isochorismate synthetase | 1.478284 | 1.510757 | 0.031348 | 0.859366 |
| transcriptional regulator, TetR family...1609 | 0.994801 | 0.987667 | -0.01038 | 0.864094 |
| metal-dependent hydrolase of the beta-lactamase superfamil | 1.010011 | 1.006955 | -0.00437 | 0.864526 |
| dihydroorotate oxidase | 1.015607 | 1.020173 | 0.006472 | 0.866132 |
| multidrug efflux protein AdeA | 1.037125 | 1.028799 | -0.01163 | 0.866313 |
| adenylate kinase | 1.011907 | 1.004923 | -0.00999 | 0.867325 |
| phosphocarrier protein HPr | 1.027808 | 1.031872 | 0.005693 | 0.868009 |
| hypothetical protein ABUW_0859 | 0.994606 | 0.986044 | -0.01247 | 0.869313 |
| beta-lactamase OXA-23 | 1.029189 | 1.024405 | -0.00672 | 0.871875 |
| multidrug efflux protein...539 | 1.136814 | 1.130405 | -0.00816 | 0.872804 |
| tRNA pseudouridine synthase A | 0.997212 | 1.003058 | 0.008433 | 0.878764 |
| D-and L-methionine transport protein | 0.972316 | 0.990955 | 0.027394 | 0.879234 |
| NAD-dependent malic enzyme | 1.06345 | 1.060559 | -0.00393 | 0.880237 |
| hypothetical protein ABUW_1140 | 0.85215 | 0.869469 | 0.029028 | 0.880594 |
| alcohol dehydrogenase | 1.050552 | 1.037273 | -0.01835 | 0.881915 |
| tRNA(Ile)-lysidine synthase | 0.93487 | 0.924881 | -0.0155 | 0.883657 |

| | | | | |
|---|---|---|---|---|
| polysaccharide deacetylase...694 | 0.968507 | 0.973656 | 0.00765 | 0.88388 |
| GTPase | 0.973655 | 0.98233 | 0.012797 | 0.88504 |
| haloacid dehydrogenase | 0.657662 | 0.669358 | 0.025432 | 0.888684 |
| YgfB/YecA family protein | 1.037134 | 1.031484 | -0.00788 | 0.891486 |
| glutathione-regulated potassium-efflux system protein KefB | 1.020351 | 1.014142 | -0.00881 | 0.891692 |
| ferredoxin--NADP(+) reductase | 1.01571 | 1.012561 | -0.00448 | 0.891729 |
| hypothetical protein ABUW_2321 | 0.898354 | 0.87958 | -0.03047 | 0.892351 |
| leucine-responsive regulatory protein | 1.095523 | 1.084074 | -0.01516 | 0.892537 |
| nucleoside diphosphate kinase | 1.036175 | 1.030965 | -0.00727 | 0.894185 |
| hypothetical protein ABUW_1604 | 0.96324 | 0.969415 | 0.009219 | 0.894673 |
| hypothetical protein ABUW_2502 | 1.0222 | 1.013715 | -0.01203 | 0.896316 |
| ribonuclease E | 0.98063 | 0.981607 | 0.001437 | 0.896533 |
| bacterioferritin...2280 | 0.99948 | 1.005294 | 0.008368 | 0.901366 |
| pH adaptation potassium efflux system protein G | 0.923411 | 0.941044 | 0.027289 | 0.90216 |
| phosphoenolpyruvate-protein phosphotransferase | 1.004908 | 1.007002 | 0.003003 | 0.902437 |
| hypothetical protein ABUW_3438 | 1.042318 | 1.057446 | 0.020789 | 0.90485 |
| transcriptional regulator, XRE family...24 | 1.085151 | 1.068775 | -0.02194 | 0.906303 |
| competence/damage-inducible protein CinA...2042 | 0.954054 | 0.960615 | 0.009888 | 0.9069 |
| protein TolR | 0.958715 | 0.950637 | -0.01221 | 0.90722 |
| FilC | 0.96813 | 0.978969 | 0.016063 | 0.907824 |
| hydrolase, NUDIX family protein | 0.930341 | 0.922245 | -0.01261 | 0.908729 |
| lipoprotein, putative...1678 | 0.995394 | 0.991347 | -0.00588 | 0.911513 |
| putative transport protein | 0.919918 | 0.91364 | -0.00988 | 0.912519 |
| hypothetical protein ABUW_2148 | 1.001947 | 1.004686 | 0.003939 | 0.913025 |
| carbon starvation protein A | 0.978502 | 0.974904 | -0.00531 | 0.914672 |
| hypothetical protein ABUW_3501 | 1.03759 | 1.029664 | -0.01106 | 0.915548 |
| transcriptional regulator, AsnC/Lrp family | 0.975077 | 0.979649 | 0.00675 | 0.918605 |
| D-methionine transport protein | 0.867713 | 0.872813 | 0.008454 | 0.9187 |
| transcriptional regulator, TetR family...478 | 0.919536 | 0.913772 | -0.00907 | 0.919307 |
| hypothetical protein ABUW_3358 | 1.021738 | 1.007686 | -0.01998 | 0.923932 |
| CobW/P47K family protein | 1.124047 | 1.116076 | -0.01027 | 0.92421 |
| transcriptional regulator, AraC family...1160 | 0.976857 | 0.972696 | -0.00616 | 0.92646 |
| hypothetical protein ABUW_4029 (plasmid) | 1.115026 | 1.102661 | -0.01609 | 0.928927 |
| hypothetical protein ABUW_3382 | 0.986577 | 0.990595 | 0.005864 | 0.929537 |
| peptidase M23/M37 family | 0.953091 | 0.948798 | -0.00651 | 0.933263 |
| phosphate ABC transporter, permease protein | 1.08603 | 1.089428 | 0.004507 | 0.934107 |
| hypothetical protein ABUW_0302 | 1.014711 | 1.019599 | 0.006934 | 0.934128 |
| phosphopyruvate hydratase (enolase) | 1.007891 | 1.013 | 0.007294 | 0.934792 |
| hypothetical protein ABUW_3111 | 1.018972 | 1.020159 | 0.00168 | 0.940149 |
| hypothetical protein ABUW_2705 | 0.938131 | 0.944333 | 0.009506 | 0.940224 |
| ribosomal protein S19 | 1.026276 | 1.025289 | -0.00139 | 0.94283 |
| sulfate ABC transporter, ATP-binding protein CysA | 0.955298 | 0.962836 | 0.011338 | 0.944434 |
| succinate-semialdehyde dehydrogenase (NADP+)...1602 | 1.014993 | 1.018308 | 0.004705 | 0.94677 |
| rod shape-determining protein MreB | 1.01205 | 1.013218 | 0.001664 | 0.946803 |
| hypothetical protein ABUW_1563 | 1.148606 | 1.154741 | 0.007686 | 0.948133 |
| Aminoglycoside phosphotransferase (plasmid) | 0.962841 | 0.958863 | -0.00597 | 0.948491 |
| type IV-A pilus assembly ATPase PilB | 1.051581 | 1.060686 | 0.012437 | 0.948784 |

| | | | | |
|---|---|---|---|---|
| hypothetical protein ABUW_1263 | 0.999758 | 1.002759 | 0.004324 | 0.950616 |
| acetoin dehydrogenase | 0.89235 | 0.889396 | -0.00478 | 0.950648 |
| type IV pilus response regulator protein PilH | 1.001577 | 0.99974 | -0.00265 | 0.952119 |
| outer membrane protein...2465 | 1.013468 | 0.999316 | -0.02029 | 0.952878 |
| L-serine dehydratase | 0.995373 | 0.992773 | -0.00377 | 0.954559 |
| methylated-DNA--protein-cysteine methyltransferase | 0.985105 | 0.982657 | -0.00359 | 0.956025 |
| cytosol aminopeptidase | 1.007174 | 1.0033 | -0.00556 | 0.956374 |
| hypothetical protein ABUW_0835 | 0.979571 | 0.986716 | 0.010485 | 0.958721 |
| hypothetical protein ABUW_0610 | 1.014067 | 1.011383 | -0.00382 | 0.958759 |
| hypothetical protein ABUW_0333 | 1.020614 | 1.021305 | 0.000977 | 0.959504 |
| transcriptional regulator, LysR family...251 | 0.957721 | 0.954742 | -0.00449 | 0.959807 |
| short-chain dehydrogenase/reductase...1803 | 1.016986 | 1.019126 | 0.003033 | 0.962728 |
| hydrolase...334 | 1.013032 | 1.011029 | -0.00286 | 0.964288 |
| hypothetical protein ABUW_0811 | 1.032701 | 1.030516 | -0.00306 | 0.964624 |
| 3-hydroxyacyl-CoA dehydrogenase...1561 | 0.985542 | 0.986373 | 0.001215 | 0.965543 |
| phage shock protein C | 1.019347 | 1.020783 | 0.00203 | 0.967096 |
| hypothetical protein ABUW_0841 | 1.001228 | 1.005895 | 0.006709 | 0.967304 |
| pca operon regulatory protein | 0.91442 | 0.917903 | 0.005484 | 0.967682 |
| hypothetical protein ABUW_1302 | 1.081162 | 1.074931 | -0.00834 | 0.971584 |
| short chain dehydrogenase...696 | 0.961669 | 0.964753 | 0.004618 | 0.972621 |
| phenylacetate-CoA oxygenase, PaaJ subunit | 0.942419 | 0.945779 | 0.005134 | 0.972717 |
| S-adenosyl-methyltransferase MraW | 1.016404 | 1.015943 | -0.00065 | 0.973568 |
| xanthine/uracil permease | 1.105956 | 1.109602 | 0.004749 | 0.974554 |
| two-component system sensor kinase | 1.001345 | 0.999303 | -0.00295 | 0.974889 |
| acetoin:2,6-dichlorophenolindophenol oxidoreductase subuni | 0.922722 | 0.921992 | -0.00114 | 0.975377 |
| putative transcriptional regulator...797 | 0.945724 | 0.947532 | 0.002756 | 0.975812 |
| transcriptional regulator, LysR family...31 | 1.022004 | 1.025309 | 0.004657 | 0.975948 |
| formyltetrahydrofolate deformylase | 1.024131 | 1.025092 | 0.001354 | 0.982478 |
| 3-deoxy-7-phosphoheptulonate synthase...356 | 0.942651 | 0.941137 | -0.00232 | 0.982535 |
| transcriptional regulator, TetR family...498 | 1.012025 | 1.008347 | -0.00525 | 0.983913 |
| thermostable carboxypeptidase 1 | 0.9729 | 0.973404 | 0.000746 | 0.985755 |
| tRNA (5-methylaminomethyl-2-thiouridylate)-methyltransfera | 0.966536 | 0.966913 | 0.000562 | 0.986697 |
| hypothetical protein ABUW_1443 | 0.950966 | 0.950151 | -0.00124 | 0.987723 |
| tol-pal system-associated acyl-CoA thioesterase | 1.033545 | 1.03287 | -0.00094 | 0.987958 |
| carbonate dehydratase | 1.030087 | 1.029705 | -0.00054 | 0.988155 |
| phospholipase D/Transphosphatidylase | 0.9963 | 0.996925 | 0.000904 | 0.988437 |
| ATP-dependent Clp protease, ATP-binding subunit ClpX | 0.988888 | 0.989096 | 0.000304 | 0.988895 |
| glutathione S-transferase...2276 | 1.021948 | 1.021409 | -0.00076 | 0.989614 |
| lipocalin family protein | 0.944418 | 0.943594 | -0.00126 | 0.99132 |
| transcription termination factor Rho | 0.973988 | 0.974287 | 0.000444 | 0.992183 |
| O-antigen polymerase family | 0.98193 | 0.982607 | 0.000995 | 0.992338 |
| hypothetical protein ABUW_0625 | 1.001969 | 1.001455 | -0.00074 | 0.993141 |
| thioesterase superfamily protein...1305 | 0.937049 | 0.937572 | 0.000804 | 0.99503 |
| hypothetical protein ABUW_2890 | 0.995042 | 0.995104 | 8.89E-05 | 0.995564 |
| acyl-CoA dehydrogenase domain protein | 0.989939 | 0.990197 | 0.000377 | 0.996521 |
| cysteine synthase A | 0.992641 | 0.992862 | 0.000321 | 0.996533 |
| tRNA-specific adenosine deaminase | 0.939401 | 0.939251 | -0.00023 | 0.997549 |

| | | | | |
|---|---|---|---|---|
| hypothetical protein ABUW_2059 | 1.01465 | 1.014525 | -0.00018 | 0.997634 |
| hypothetical protein ABUW_2582 | 1.003768 | 1.003741 | -4E-05 | 0.999301 |

| t-value |
|---|
| -68.3362 |
| -40.448 |
| -39.6778 |
| 34.85932 |
| 28.7545 |
| -28.2802 |
| -25.2076 |
| 59.40785 |
| -29.0584 |
| 26.35483 |
| -31.9184 |
| 23.90979 |
| -24.6109 |
| -25.0629 |
| 21.5959 |
| 33.22187 |
| -23.8242 |
| 21.17263 |
| 24.6772 |
| -25.7643 |
| 18.36062 |
| 20.94385 |
| -17.5566 |
| -20.7746 |
| -18.3795 |
| 16.83445 |
| -18.4153 |
| 17.98613 |
| 33.3525 |
| 22.3035 |
| 28.28861 |
| 17.30147 |
| -16.3461 |
| 15.37262 |
| 15.47778 |
| 21.15056 |
| 15.36354 |
| 15.56161 |
| -17.4985 |
| 19.40398 |
| 17.84768 |
| 20.26822 |
| 25.36731 |
| 16.76034 |
| 14.76379 |
| 15.56831 |

| |
|---|
| -28.0693 |
| 47.03158 |
| -13.6347 |
| 18.70469 |
| -12.9968 |
| -19.7068 |
| -13.6191 |
| 19.07127 |
| 13.28606 |
| 12.55773 |
| 24.29 |
| 17.64441 |
| 12.47738 |
| -12.3697 |
| -24.6041 |
| 23.28345 |
| 14.20859 |
| 37.85023 |
| 14.0602 |
| 13.08018 |
| -16.1202 |
| 12.88519 |
| -18.6993 |
| -20.7065 |
| 15.69072 |
| -12.3456 |
| 17.24948 |
| 11.49972 |
| -12.5352 |
| 12.05665 |
| 18.65814 |
| -19.8213 |
| 12.54292 |
| 16.22874 |
| -12.0189 |
| -41.3652 |
| 14.39797 |
| -20.7366 |
| -11.2249 |
| -10.8707 |
| 16.34819 |
| 29.18918 |
| 11.01757 |
| 11.17888 |
| 12.12017 |
| 11.18718 |
| -15.2291 |

| |
|---|
| 16.67392 |
| -12.3985 |
| 10.61922 |
| -11.7078 |
| -14.1231 |
| 13.44889 |
| 17.37478 |
| 15.4992 |
| 10.44584 |
| -10.4485 |
| -10.337 |
| 11.12664 |
| 22.18695 |
| 12.04772 |
| 14.77134 |
| 16.61557 |
| -10.0352 |
| 24.41997 |
| -17.4325 |
| -11.5416 |
| 10.50679 |
| -12.6678 |
| 13.53459 |
| 20.42819 |
| 9.861764 |
| -13.3767 |
| -10.2924 |
| 11.48218 |
| 15.63133 |
| 9.610592 |
| 13.16391 |
| 11.75226 |
| 15.88807 |
| 15.40714 |
| -9.60519 |
| 12.57958 |
| 15.93688 |
| -19.5211 |
| -9.18373 |
| 9.782606 |
| 9.1395 |
| -9.13537 |
| -9.58553 |
| -10.9705 |
| 15.31599 |
| 10.91891 |
| 9.066963 |

| |
|---|
| -12.539 |
| -8.89828 |
| 10.14803 |
| -11.7307 |
| 10.39636 |
| -8.99255 |
| 14.11081 |
| -10.046 |
| 12.7012 |
| 10.5363 |
| -9.34909 |
| 21.44311 |
| -11.258 |
| 8.410856 |
| 13.35574 |
| -8.75969 |
| 8.37545 |
| 12.81522 |
| 10.12765 |
| -8.29211 |
| -8.6373 |
| 8.340876 |
| -12.4296 |
| 8.266995 |
| 8.16303 |
| 8.119746 |
| 13.75153 |
| -8.07646 |
| 9.360874 |
| 7.994517 |
| -18.195 |
| -24.0698 |
| -8.08058 |
| 7.990278 |
| 11.32085 |
| 15.67767 |
| 10.54138 |
| -12.8157 |
| -8.26193 |
| 12.47345 |
| 15.29059 |
| -8.01566 |
| 8.170866 |
| -7.88539 |
| -7.79422 |
| 11.6655 |
| -7.69862 |

| |
|---|
| 8.052601 |
| 7.581636 |
| 9.073618 |
| 10.49049 |
| 9.627168 |
| 8.22478 |
| 8.369244 |
| -7.55843 |
| 7.424872 |
| 7.647675 |
| 9.426968 |
| -11.1523 |
| -7.50907 |
| 8.626275 |
| -7.36787 |
| 7.886582 |
| -10.7577 |
| 7.506721 |
| 18.66396 |
| 8.663741 |
| -7.14468 |
| 12.69362 |
| -10.7539 |
| 13.06975 |
| -7.8903 |
| -8.21777 |
| -8.86767 |
| 9.932147 |
| -10.3464 |
| 8.01655 |
| -10.8547 |
| -7.11165 |
| 7.727414 |
| 8.076063 |
| 9.321706 |
| -7.07089 |
| 8.063965 |
| -6.92249 |
| -11.968 |
| 7.651379 |
| 9.652001 |
| 8.761812 |
| -10.9313 |
| 7.700935 |
| 8.89869 |
| 8.670112 |
| -12.5393 |

| |
|---|
| 12.21966 |
| 6.817794 |
| 7.847166 |
| 9.014528 |
| 8.70081 |
| 8.164755 |
| 13.08678 |
| -9.34456 |
| 9.251849 |
| 8.016836 |
| 11.73273 |
| -7.52244 |
| -7.02966 |
| -6.8839 |
| 9.242607 |
| 10.06957 |
| 7.660069 |
| 8.964103 |
| -6.56311 |
| 7.132462 |
| -7.26923 |
| 17.16065 |
| -14.4315 |
| -14.6894 |
| 15.88923 |
| -6.72548 |
| 12.77375 |
| 10.10285 |
| 6.366901 |
| 6.296524 |
| 6.589579 |
| 6.37988 |
| 7.077664 |
| 6.919408 |
| -6.23278 |
| -10.2013 |
| 9.981254 |
| 6.617099 |
| -14.9665 |
| 9.563153 |
| 7.872945 |
| -6.52071 |
| 6.170437 |
| -10.8025 |
| -6.4851 |
| -7.19956 |
| -8.48204 |

| |
|---|
| -8.57068 |
| 8.89757 |
| 7.770869 |
| 13.91483 |
| 8.052923 |
| 6.952013 |
| 7.280284 |
| -9.92692 |
| -6.07859 |
| -7.96235 |
| 6.456862 |
| 10.77686 |
| 12.99093 |
| 7.854026 |
| 11.99933 |
| 5.943963 |
| 10.96203 |
| 5.93244 |
| -6.63069 |
| -12.6357 |
| -9.69135 |
| 7.298527 |
| 5.823539 |
| 6.154361 |
| 6.000272 |
| 12.84228 |
| 7.335326 |
| 11.90589 |
| -11.604 |
| 8.462466 |
| 10.59304 |
| -9.20322 |
| -8.60313 |
| 7.055431 |
| 7.10097 |
| -6.27795 |
| 9.773527 |
| 8.482663 |
| -7.86296 |
| -9.72411 |
| -10.441 |
| 10.63376 |
| 9.580653 |
| 5.577731 |
| 5.734581 |
| -7.04275 |
| 6.928798 |

| |
|---|
| -7.7055 |
| -10.3777 |
| -9.04719 |
| 8.416748 |
| 6.004201 |
| 8.465766 |
| 6.805474 |
| -6.67894 |
| 8.316331 |
| 5.833596 |
| 11.98416 |
| 10.15268 |
| 7.777461 |
| 5.455549 |
| -6.0172 |
| 10.40703 |
| 6.166248 |
| 5.53013 |
| -10.1952 |
| -5.47402 |
| -6.96943 |
| 6.401237 |
| -5.51269 |
| -11.652 |
| 7.383203 |
| 7.110634 |
| -10.4614 |
| 5.660632 |
| 5.326161 |
| 7.866751 |
| 9.585193 |
| -11.3564 |
| 6.324033 |
| 5.402173 |
| 11.24707 |
| -6.34778 |
| 5.299293 |
| 5.716877 |
| 5.87518 |
| -10.4403 |
| -5.55794 |
| -6.82726 |
| 5.235991 |
| 9.034872 |
| -7.76059 |
| -10.1127 |
| 12.0412 |

| |
|---|
| -5.73377 |
| -10.8408 |
| 6.455627 |
| 6.079897 |
| -8.89527 |
| 5.916829 |
| 9.351869 |
| 8.349187 |
| 5.845482 |
| -9.87549 |
| 5.45197 |
| 5.709228 |
| 10.01169 |
| -8.58848 |
| 7.143861 |
| 9.296184 |
| 11.37127 |
| 7.218872 |
| 5.225857 |
| -7.11114 |
| -8.59667 |
| 6.967666 |
| 5.225494 |
| 5.845632 |
| -5.79557 |
| -11.0369 |
| 6.588493 |
| 6.229817 |
| 10.40111 |
| 5.322651 |
| -5.28136 |
| 5.938575 |
| -9.18442 |
| 6.547823 |
| -8.72353 |
| 6.77947 |
| 10.32298 |
| -6.13642 |
| -8.54287 |
| -6.72985 |
| 5.757704 |
| 10.00015 |
| 5.581561 |
| 9.560785 |
| 5.410268 |
| 5.037389 |
| 6.158824 |

| |
|---|
| 5.358819 |
| -4.89011 |
| 10.17503 |
| -6.31585 |
| -4.87097 |
| -7.79365 |
| -7.51736 |
| 6.950074 |
| 8.411285 |
| -5.31505 |
| 7.022216 |
| -7.56898 |
| 9.638671 |
| -5.16822 |
| -5.26669 |
| -8.51533 |
| 4.810544 |
| 5.849591 |
| -6.87872 |
| -7.44878 |
| -6.91658 |
| 5.919647 |
| -8.47292 |
| 4.751776 |
| 6.284184 |
| -6.25461 |
| -5.12005 |
| -6.37344 |
| 7.139596 |
| 6.770666 |
| -8.47865 |
| -5.09705 |
| -9.19472 |
| -7.33962 |
| 7.902794 |
| 4.875446 |
| 5.004844 |
| -4.66446 |
| 8.717418 |
| -7.24741 |
| 5.301475 |
| 4.755479 |
| -9.08821 |
| 4.900798 |
| 6.879539 |
| -6.46844 |
| -5.63937 |

| |
|---|
| 6.482236 |
| 7.313592 |
| -6.6331 |
| 6.686858 |
| 7.978709 |
| -6.54899 |
| 5.080567 |
| 5.156079 |
| 4.761371 |
| -9.2276 |
| -4.56003 |
| -8.9534 |
| 6.496738 |
| -7.932 |
| 5.707907 |
| -4.52451 |
| 4.567789 |
| 7.11898 |
| 7.981623 |
| 7.816767 |
| -6.74498 |
| 4.492002 |
| 7.402148 |
| -4.54136 |
| -7.55362 |
| 5.347637 |
| 7.020302 |
| 7.035655 |
| 8.108153 |
| -4.52092 |
| 4.660808 |
| 4.424614 |
| -6.67365 |
| -4.78853 |
| -4.54271 |
| 8.925879 |
| 5.08172 |
| 5.229451 |
| -4.86006 |
| -7.43196 |
| -4.59026 |
| 4.938262 |
| -4.52477 |
| -5.7102 |
| -5.42821 |
| 5.378464 |
| 6.671122 |

| |
|---|
| -7.48934 |
| 4.875926 |
| -6.00407 |
| 4.944807 |
| 6.603974 |
| 8.347781 |
| -4.84067 |
| -5.91754 |
| 4.898304 |
| 8.39475 |
| 8.103952 |
| -8.68545 |
| 4.565293 |
| -4.9978 |
| 4.57308 |
| -4.48552 |
| 4.45773 |
| -4.62902 |
| -5.2889 |
| -8.05999 |
| -8.26776 |
| 8.420111 |
| -5.84984 |
| 5.864623 |
| -6.6892 |
| -7.86226 |
| 5.437713 |
| -4.25057 |
| 5.095118 |
| 6.026983 |
| -4.36655 |
| 8.064221 |
| -5.03713 |
| -7.01149 |
| -4.24133 |
| -7.38061 |
| -4.96325 |
| -7.73891 |
| -7.43099 |
| 6.226074 |
| -5.79762 |
| 7.739047 |
| -8.10075 |
| -5.23229 |
| -7.1726 |
| 5.360924 |
| 5.43684 |

| |
|---|
| 6.344787 |
| 4.217571 |
| 5.500924 |
| -7.31518 |
| 4.116018 |
| 7.937817 |
| -7.11315 |
| 5.486331 |
| -7.83287 |
| -5.64844 |
| 5.038634 |
| -6.92957 |
| -6.33156 |
| -6.24893 |
| 4.607914 |
| 6.821112 |
| 4.669998 |
| -4.03727 |
| 4.099752 |
| -4.87169 |
| -6.01804 |
| 4.848955 |
| 4.335428 |
| 7.266766 |
| 4.47606 |
| 7.742605 |
| -4.00713 |
| 4.416724 |
| 6.422984 |
| -4.11033 |
| 6.38952 |
| 4.000643 |
| 5.650594 |
| 6.368925 |
| 3.990111 |
| -7.03553 |
| -5.46857 |
| -4.07648 |
| 3.944029 |
| -6.16076 |
| -4.38792 |
| -4.19954 |
| 5.732063 |
| 5.033573 |
| 4.430307 |
| 4.431133 |
| -7.16409 |

| |
|---|
| -7.33747 |
| 4.302827 |
| -4.41638 |
| 6.820356 |
| 4.20385 |
| 7.119703 |
| -5.15625 |
| -5.92477 |
| 3.991366 |
| 4.612609 |
| -3.86508 |
| -5.28901 |
| 5.579052 |
| -4.34759 |
| -4.57993 |
| 4.768348 |
| -6.49246 |
| 5.082527 |
| -5.60218 |
| -6.58533 |
| -5.10687 |
| -3.80887 |
| -5.73226 |
| -5.81427 |
| -5.90553 |
| -5.87575 |
| -6.41656 |
| 6.847198 |
| 5.651227 |
| 4.579615 |
| 6.721156 |
| 6.514871 |
| 4.502622 |
| 6.536965 |
| 3.835433 |
| 6.060249 |
| 6.281901 |
| -3.91392 |
| -5.22893 |
| 5.613256 |
| 4.186427 |
| 3.819637 |
| 3.974252 |
| 3.909189 |
| -6.30041 |
| 3.685761 |
| -3.67597 |

| |
|---|
| -5.67366 |
| -4.58848 |
| 5.260434 |
| -3.73116 |
| -3.68358 |
| -6.46156 |
| -5.92804 |
| 3.96841 |
| 3.65902 |
| -3.6457 |
| 3.700683 |
| -3.63492 |
| -6.33351 |
| 4.54182 |
| 4.872173 |
| -5.1849 |
| -5.53465 |
| 4.836009 |
| 4.875519 |
| -5.52064 |
| 4.370828 |
| 4.797117 |
| 5.219709 |
| -6.41575 |
| -6.03309 |
| -5.94433 |
| 3.885953 |
| 4.03604 |
| -4.36712 |
| 3.614697 |
| 5.455738 |
| -3.58769 |
| 3.67301 |
| 4.303399 |
| 3.999935 |
| 3.707713 |
| 4.901493 |
| 3.636894 |
| -3.55546 |
| -3.5736 |
| -6.24398 |
| -3.99155 |
| 4.925418 |
| 4.562489 |
| -3.64766 |
| 4.034119 |
| 5.07339 |

| |
|---|
| -3.51596 |
| 5.317915 |
| -5.42254 |
| 5.439117 |
| -3.58743 |
| -5.44646 |
| 5.30251 |
| 5.578177 |
| -4.50238 |
| -4.09776 |
| 4.753559 |
| -4.89154 |
| 3.674562 |
| 4.507549 |
| -3.5435 |
| 3.472896 |
| 5.479295 |
| -5.03108 |
| -4.89111 |
| -5.81901 |
| -4.21819 |
| -4.77629 |
| -3.55161 |
| 5.006553 |
| 4.412609 |
| -3.50985 |
| -4.69077 |
| -3.71807 |
| -4.31633 |
| 5.171046 |
| 4.247129 |
| 5.665691 |
| -5.1972 |
| 4.702032 |
| -4.79221 |
| 3.447022 |
| -4.40401 |
| 3.989564 |
| -4.09982 |
| 4.951569 |
| -5.78705 |
| -4.51018 |
| 4.337561 |
| -5.3907 |
| -5.02274 |
| -4.44752 |
| -3.51207 |

| |
|---|
| 4.696387 |
| 3.372648 |
| -5.63676 |
| -3.3563 |
| -4.50578 |
| -3.97103 |
| 3.348959 |
| -4.07669 |
| -3.34601 |
| 3.902925 |
| 5.502025 |
| -5.69114 |
| 3.380704 |
| -3.49611 |
| -4.09083 |
| 3.357191 |
| 4.580492 |
| -3.83399 |
| 4.812348 |
| -4.58378 |
| -5.54874 |
| 3.296094 |
| -5.57516 |
| 4.653576 |
| -5.11023 |
| -5.26221 |
| -3.31988 |
| 5.321804 |
| 3.704838 |
| -4.55181 |
| -4.99112 |
| -3.25948 |
| -5.13012 |
| 3.571521 |
| -5.36343 |
| -3.62199 |
| -3.89066 |
| 4.952231 |
| 5.219829 |
| -3.27366 |
| -3.32556 |
| -4.58384 |
| 4.500187 |
| 3.410007 |
| 4.421169 |
| -4.07188 |
| -3.48882 |

| |
|---|
| -4.63355 |
| -3.18052 |
| 4.578719 |
| 3.318507 |
| 5.078179 |
| 4.300079 |
| 4.681119 |
| 4.058152 |
| 3.975301 |
| 4.732675 |
| -3.25629 |
| -4.44361 |
| -4.71475 |
| 4.98049 |
| 3.520368 |
| 3.879925 |
| 4.46242 |
| 3.245958 |
| -4.13028 |
| -4.11219 |
| 3.823036 |
| 3.631069 |
| 4.246105 |
| 5.116821 |
| 3.822459 |
| 5.083266 |
| 4.416 |
| 4.001484 |
| 3.187969 |
| -3.16864 |
| -3.20433 |
| 4.94019 |
| 3.988618 |
| 3.318007 |
| 3.162488 |
| 4.7066 |
| 4.55512 |
| 4.741418 |
| -4.31203 |
| 3.850063 |
| -4.62488 |
| 3.325056 |
| -3.05965 |
| 3.613819 |
| 3.84172 |
| 4.58234 |
| -4.52302 |

| |
|---|
| 4.755525 |
| -4.53009 |
| 4.125304 |
| 3.398161 |
| 3.067192 |
| -3.10202 |
| -4.25013 |
| 3.7085 |
| 3.160247 |
| -4.66002 |
| 4.059828 |
| 4.039881 |
| -4.6263 |
| 3.954159 |
| 3.337488 |
| -4.49758 |
| -3.7489 |
| 3.065142 |
| -3.43909 |
| 3.306592 |
| 4.415086 |
| -4.4564 |
| 3.348163 |
| -3.56828 |
| 4.047667 |
| -4.75807 |
| -4.00604 |
| -3.57319 |
| 4.581671 |
| 3.140464 |
| 3.530811 |
| -3.28475 |
| 2.954525 |
| -3.40356 |
| 4.429658 |
| 4.334946 |
| -3.1647 |
| 3.09291 |
| -4.1119 |
| 3.621921 |
| 3.457144 |
| -2.98938 |
| -4.43388 |
| 3.230194 |
| 3.31846 |
| 4.109083 |
| 3.781763 |

-3.20271
4.543384
3.439965
3.953725
3.656739
3.512672
3.257211
-3.67714
3.01536
-3.14618
-3.6914
3.198836
4.439182
2.916511
-3.5967
3.077973
3.169465
3.537135
4.254876
-3.68492
-4.2871
2.982605
4.207212
3.519114
-3.57274
-3.50689
-4.42459
-3.9934
4.324597
4.023041
3.183813
-4.05038
-3.9625
-4.0314
3.284772
-3.5661
-2.86002
4.349859
4.182484
3.600409
4.314282
-4.28884
3.448407
-4.28155
3.337086
4.024424
-4.0586

|  |
| ---: |
| 3.428964 |
| -4.24088 |
| -3.62132 |
| -2.9472 |
| -2.81596 |
| -3.4101 |
| 2.827733 |
| -3.96211 |
| 3.96858 |
| -3.90232 |
| 4.117226 |
| -2.88119 |
| -4.12739 |
| 3.695308 |
| 3.082667 |
| -4.12178 |
| -2.99098 |
| 2.735326 |
| 3.471795 |
| -2.73291 |
| -3.05133 |
| 3.57738 |
| -2.75634 |
| 3.261781 |
| -3.43846 |
| -3.48032 |
| 3.694335 |
| -4.10113 |
| 3.979984 |
| 3.376817 |
| 3.798916 |
| -3.39116 |
| -3.08651 |
| -2.73973 |
| 3.439813 |
| 2.857532 |
| -3.06102 |
| 3.726584 |
| 2.944783 |
| -3.71029 |
| -3.79345 |
| -3.72486 |
| 2.766756 |
| -3.96101 |
| -3.21152 |
| 3.760761 |
| 3.057684 |

| |
|---|
| 2.671631 |
| -2.79184 |
| -4.0225 |
| -3.74335 |
| 2.924449 |
| 3.710178 |
| 3.225425 |
| -3.81344 |
| 3.355393 |
| 3.601571 |
| 3.724129 |
| -2.662 |
| -3.66013 |
| -3.17474 |
| 2.954618 |
| 3.230807 |
| -3.53976 |
| -3.01899 |
| -2.75815 |
| 3.833331 |
| 2.982865 |
| -2.62553 |
| -3.81455 |
| 2.929279 |
| -3.86416 |
| -3.82093 |
| -3.62669 |
| 3.049792 |
| -3.70671 |
| -2.83373 |
| 2.741862 |
| 3.765462 |
| 3.311366 |
| -2.59998 |
| 3.518434 |
| -3.75637 |
| 3.371374 |
| 3.014853 |
| 3.388574 |
| -3.0885 |
| -2.6645 |
| 3.39278 |
| 2.789225 |
| 3.731604 |
| -3.75187 |
| 2.981939 |
| 3.3319 |

| |
|---|
| 3.84231 |
| 3.446353 |
| -3.46255 |
| -3.52287 |
| 3.719937 |
| -3.64427 |
| 3.236608 |
| 2.959555 |
| 3.576912 |
| -3.7125 |
| -2.57545 |
| 2.556365 |
| 2.661429 |
| 3.22516 |
| -2.60174 |
| 3.095287 |
| -3.69593 |
| 2.565303 |
| 3.100029 |
| 3.182459 |
| -3.51495 |
| 3.539001 |
| -3.52516 |
| -3.40736 |
| 3.007624 |
| -3.63208 |
| -3.63627 |
| 2.562724 |
| -3.391 |
| 2.653441 |
| 2.948239 |
| 3.324986 |
| 2.959391 |
| 2.773406 |
| 3.059903 |
| 2.825629 |
| -3.55572 |
| -2.63171 |
| 3.58864 |
| 3.177057 |
| 3.366741 |
| -3.30578 |
| -2.50816 |
| 2.480135 |
| -2.60089 |
| -3.39343 |
| 3.515122 |

| |
|---|
| 3.456188 |
| 2.607117 |
| -2.47769 |
| -3.50515 |
| 2.67572 |
| 2.982601 |
| 3.573892 |
| -2.72633 |
| 2.712811 |
| 2.472592 |
| -3.42195 |
| 2.607269 |
| -3.22147 |
| 3.224486 |
| -3.5322 |
| 2.493452 |
| -3.11745 |
| 2.435444 |
| -3.06854 |
| -3.5017 |
| -3.21141 |
| -3.03542 |
| 3.437778 |
| 3.492946 |
| 2.436468 |
| 2.72938 |
| -2.48658 |
| -2.69313 |
| 3.305833 |
| -2.92756 |
| -3.11116 |
| -3.16527 |
| 2.455546 |
| -3.4265 |
| 2.610129 |
| -2.41372 |
| -2.92243 |
| 2.645932 |
| -3.43948 |
| -3.35166 |
| -3.15904 |
| 2.717609 |
| 3.113096 |
| -2.43543 |
| 3.230534 |
| -3.35046 |
| -3.29136 |

| |
|---|
| 2.713733 |
| 2.844155 |
| 3.07484 |
| -3.31894 |
| -2.74727 |
| 3.391363 |
| 2.891352 |
| -3.22023 |
| -2.47943 |
| -2.67261 |
| -2.42831 |
| -3.13517 |
| -3.14561 |
| 3.221951 |
| -3.12483 |
| -2.92237 |
| 2.885254 |
| -3.30084 |
| -3.22288 |
| -3.23886 |
| -3.05549 |
| -3.16631 |
| 2.420678 |
| 2.820995 |
| -2.94462 |
| 2.552903 |
| -2.98277 |
| 2.33032 |
| 3.169127 |
| -3.24298 |
| 2.446137 |
| -2.78986 |
| 2.927688 |
| 2.702875 |
| -2.96489 |
| 2.294307 |
| -3.20411 |
| 2.319016 |
| -2.47033 |
| 2.302256 |
| -3.13014 |
| -3.17729 |
| 2.85757 |
| -2.75116 |
| -2.84296 |
| -2.91784 |
| -3.1733 |

| |
|---|
| 2.865837 |
| 2.671777 |
| 2.527863 |
| 3.029139 |
| -3.08753 |
| -3.07224 |
| 2.984308 |
| 2.679134 |
| 2.452642 |
| -2.29324 |
| -2.87039 |
| 2.521121 |
| -2.38772 |
| 3.053815 |
| -3.1341 |
| 2.720874 |
| -3.16127 |
| -2.95468 |
| -2.7406 |
| -3.08539 |
| 2.276033 |
| -2.45175 |
| -3.1262 |
| -3.09218 |
| 3.076911 |
| 3.070678 |
| -2.26816 |
| -3.05865 |
| 2.966783 |
| -2.70826 |
| 2.969694 |
| 2.988598 |
| -2.76859 |
| 2.637447 |
| -2.74703 |
| -2.93779 |
| 2.892723 |
| 2.608625 |
| 2.625657 |
| 2.561341 |
| 3.008771 |
| 2.227164 |
| 2.274613 |
| -3.04634 |
| -2.74215 |
| 2.29825 |
| -2.27206 |

| |
|---|
| -2.95447 |
| 2.40962 |
| 2.95673 |
| 2.985366 |
| 2.794074 |
| -2.4212 |
| -2.83351 |
| 2.654236 |
| -2.95249 |
| -2.44034 |
| -2.94595 |
| 2.77299 |
| -2.67775 |
| 2.691053 |
| -2.78692 |
| 2.327577 |
| -2.86242 |
| 2.859035 |
| -2.98508 |
| 2.367726 |
| 2.914093 |
| 2.649149 |
| 2.505525 |
| -2.29913 |
| -2.65662 |
| 2.604795 |
| -2.66209 |
| -2.85086 |
| 2.643031 |
| -2.37722 |
| -2.80231 |
| 2.585307 |
| 2.522672 |
| 2.436813 |
| 2.672871 |
| 2.418624 |
| -2.74523 |
| 2.643546 |
| -2.89032 |
| 2.752596 |
| 2.32895 |
| -2.79706 |
| -2.49796 |
| 2.493766 |
| 2.850803 |
| 2.183278 |
| -2.62712 |

| |
|---|
| 2.341194 |
| -2.4008 |
| -2.7379 |
| -2.82779 |
| 2.646377 |
| -2.6485 |
| -2.43437 |
| -2.60209 |
| 2.190796 |
| -2.77852 |
| -2.58446 |
| 2.447696 |
| 2.616316 |
| -2.77549 |
| -2.65462 |
| -2.47053 |
| 2.691544 |
| -2.07284 |
| 2.708963 |
| -2.77835 |
| 2.465377 |
| 2.6337 |
| 2.320394 |
| -2.19409 |
| -2.73502 |
| -2.06353 |
| -2.62189 |
| -2.72316 |
| -2.68041 |
| -2.66476 |
| 2.133918 |
| 2.691095 |
| 2.672687 |
| 2.700503 |
| -2.59251 |
| -2.53872 |
| -2.03848 |
| 2.509229 |
| -2.73133 |
| -2.03247 |
| 2.149257 |
| -2.05909 |
| -2.3597 |
| 2.172593 |
| -2.63107 |
| 2.034518 |
| 2.444774 |

| |
|---|
| -2.48875 |
| 2.143102 |
| 2.024363 |
| 2.249129 |
| -2.62546 |
| -2.02246 |
| 2.177858 |
| -2.54292 |
| 2.650113 |
| -2.60874 |
| 2.66569 |
| -2.32539 |
| 2.626857 |
| -2.20721 |
| 2.670057 |
| -2.22064 |
| -2.61826 |
| -2.65444 |
| -2.61952 |
| 2.604814 |
| 2.583513 |
| -2.21085 |
| 2.547718 |
| 2.096794 |
| 2.558471 |
| 2.026286 |
| -2.44967 |
| 2.002376 |
| -2.59779 |
| 2.555776 |
| -2.5867 |
| -2.11871 |
| -2.2417 |
| -2.08252 |
| -2.54881 |
| -2.22564 |
| -2.10887 |
| -2.56968 |
| -2.38948 |
| -2.56272 |
| -2.49989 |
| 2.392178 |
| -2.29536 |
| -1.95178 |
| -2.12889 |
| -2.49917 |
| -2.23622 |

| |
|---|
| 2.383478 |
| 2.069338 |
| -2.54624 |
| -2.52747 |
| -2.14493 |
| -2.55362 |
| 2.195173 |
| -2.16864 |
| -2.48148 |
| 2.139046 |
| -2.2987 |
| -2.54526 |
| -2.49772 |
| 2.204753 |
| -2.36458 |
| -2.50376 |
| 1.932344 |
| -1.95534 |
| 2.497085 |
| -2.10339 |
| 2.165357 |
| -2.38292 |
| 2.096718 |
| -2.03462 |
| 2.310442 |
| -1.93294 |
| 2.265611 |
| -2.4714 |
| 2.022539 |
| 2.385738 |
| 2.488338 |
| -2.08183 |
| -2.14611 |
| -2.39418 |
| 2.091267 |
| -1.95922 |
| -2.10422 |
| -2.45948 |
| 2.475418 |
| -2.0837 |
| 2.227198 |
| -2.40648 |
| 2.456304 |
| -2.39833 |
| 1.931128 |
| -1.94493 |
| 1.962494 |

| |
|---|
| 2.139815 |
| -2.37279 |
| 2.162207 |
| 1.881047 |
| -2.41966 |
| 2.01266 |
| -1.89673 |
| -2.11098 |
| 2.02291 |
| 2.349645 |
| 2.409274 |
| 2.327693 |
| -1.87903 |
| 2.026995 |
| 2.418774 |
| 2.06585 |
| -1.95874 |
| -2.41613 |
| 2.045981 |
| 2.395282 |
| 2.151985 |
| -2.18926 |
| 1.898258 |
| 2.16082 |
| -1.97429 |
| -1.92494 |
| 2.173066 |
| -2.05135 |
| -1.89156 |
| -2.21607 |
| -1.85379 |
| 2.238509 |
| 1.895857 |
| 2.255499 |
| 2.306516 |
| 1.830576 |
| 2.065845 |
| -2.16977 |
| -2.35915 |
| -2.20125 |
| -2.02866 |
| -2.33643 |
| -2.07042 |
| 2.070549 |
| -2.27257 |
| 2.232386 |
| -2.32278 |

| |
|---|
| -2.28079 |
| -2.20461 |
| 2.037655 |
| -2.07394 |
| -2.29146 |
| 2.124824 |
| -2.3167 |
| 1.797457 |
| 1.990153 |
| -2.20793 |
| -2.29101 |
| -1.90923 |
| 2.186777 |
| -2.24513 |
| -2.19596 |
| -2.09255 |
| 2.239277 |
| 2.190288 |
| -2.18756 |
| 1.82035 |
| 2.231632 |
| -2.14419 |
| -2.02759 |
| -2.16404 |
| 1.833622 |
| 2.205992 |
| -2.23658 |
| -2.21528 |
| -2.07573 |
| 1.755971 |
| -2.1909 |
| 2.172238 |
| -2.07215 |
| -1.80634 |
| -2.05574 |
| -2.05663 |
| -1.88521 |
| -2.04351 |
| -2.15771 |
| 1.864008 |
| -2.18674 |
| -2.19558 |
| 2.155842 |
| 1.801933 |
| -2.10138 |
| -2.03601 |
| -1.94967 |

| |
|---|
| -1.72104 |
| 2.160783 |
| 2.172239 |
| -2.18048 |
| 1.762043 |
| -2.17056 |
| 1.842759 |
| -1.8966 |
| 2.126357 |
| -1.7448 |
| 1.990309 |
| -1.95003 |
| 1.871033 |
| 1.946775 |
| -2.05671 |
| -2.14488 |
| -2.09533 |
| 2.012216 |
| -2.12794 |
| -1.76778 |
| -2.14637 |
| -2.1434 |
| -2.06267 |
| -2.03063 |
| 2.124843 |
| 1.695007 |
| -1.95163 |
| -2.12744 |
| -2.08445 |
| -2.02646 |
| -1.70938 |
| -1.85073 |
| -2.08943 |
| -2.05543 |
| -2.11444 |
| 1.725142 |
| 1.693788 |
| 1.784393 |
| -2.02205 |
| -1.67655 |
| -2.07991 |
| 1.767794 |
| 1.826739 |
| -2.08322 |
| -1.93123 |
| -1.82023 |
| 2.002493 |

| |
|---|
| 1.689927 |
| -1.96105 |
| 1.710028 |
| -1.97829 |
| -1.89207 |
| 1.939569 |
| -2.05739 |
| 1.644753 |
| 1.951389 |
| 1.89416 |
| -1.91127 |
| 1.746256 |
| 1.635727 |
| -2.01801 |
| -1.94712 |
| -1.63154 |
| -2.02635 |
| 1.835614 |
| -2.0106 |
| -2.02148 |
| 1.984132 |
| 1.988624 |
| 1.624681 |
| 1.847524 |
| -1.98607 |
| -1.94047 |
| 1.670902 |
| 1.849094 |
| -1.9978 |
| 1.72672 |
| 1.743446 |
| -1.85279 |
| -1.77273 |
| 1.848217 |
| 1.907786 |
| -1.93069 |
| -1.78118 |
| 1.743182 |
| -1.88829 |
| 1.718525 |
| 1.970343 |
| 1.807194 |
| 1.586005 |
| -1.73179 |
| 1.581125 |
| -1.93678 |
| -1.83444 |

| |
|---|
| 1.739476 |
| -1.62088 |
| -1.92311 |
| -1.61512 |
| 1.883537 |
| -1.81895 |
| 1.876738 |
| -1.83819 |
| -1.68058 |
| -1.83557 |
| 1.739341 |
| -1.55743 |
| -1.91088 |
| -1.77492 |
| 1.554259 |
| -1.87775 |
| -1.9081 |
| -1.67925 |
| 1.858805 |
| -1.56432 |
| -1.75703 |
| -1.88691 |
| 1.674971 |
| 1.593695 |
| -1.87552 |
| -1.86171 |
| -1.87559 |
| 1.563475 |
| -1.74878 |
| -1.69982 |
| 1.824447 |
| -1.73579 |
| -1.82998 |
| -1.52796 |
| 1.688695 |
| 1.520874 |
| 1.50971 |
| 1.835306 |
| 1.764793 |
| -1.63404 |
| -1.55571 |
| 1.720801 |
| 1.546886 |
| 1.79512 |
| -1.52065 |
| -1.54258 |
| 1.541369 |

| |
|---|
| -1.48299 |
| -1.69591 |
| -1.49536 |
| -1.62971 |
| 1.542769 |
| 1.788917 |
| 1.742315 |
| 1.479925 |
| 1.540949 |
| -1.62691 |
| 1.48023 |
| 1.771611 |
| 1.509041 |
| -1.77585 |
| -1.53999 |
| -1.75556 |
| 1.647445 |
| 1.605273 |
| -1.5738 |
| -1.77046 |
| 1.470377 |
| 1.773972 |
| -1.74986 |
| -1.69825 |
| -1.73272 |
| 1.474412 |
| 1.749944 |
| 1.716078 |
| 1.457716 |
| -1.75716 |
| -1.70434 |
| 1.543646 |
| 1.751196 |
| -1.71212 |
| 1.588642 |
| 1.463703 |
| -1.70312 |
| -1.57201 |
| -1.73549 |
| -1.67778 |
| -1.47829 |
| 1.611631 |
| -1.71199 |
| -1.69928 |
| 1.577213 |
| 1.558947 |
| -1.66481 |

| |
|---|
| 1.610134 |
| 1.637232 |
| -1.54144 |
| 1.454027 |
| 1.432459 |
| 1.622455 |
| -1.7048 |
| 1.646059 |
| -1.68088 |
| -1.46376 |
| 1.686055 |
| 1.573239 |
| -1.68075 |
| -1.53611 |
| -1.59361 |
| -1.57303 |
| 1.39652 |
| 1.647512 |
| 1.641794 |
| 1.572127 |
| -1.57599 |
| 1.658736 |
| 1.58881 |
| -1.41432 |
| -1.64464 |
| -1.64402 |
| -1.59597 |
| 1.457895 |
| -1.4365 |
| 1.646608 |
| -1.5628 |
| 1.478566 |
| -1.63794 |
| -1.42651 |
| -1.44509 |
| -1.5724 |
| 1.449218 |
| 1.612362 |
| 1.491596 |
| 1.624101 |
| 1.522032 |
| -1.60632 |
| 1.483843 |
| -1.3593 |
| -1.45431 |
| -1.48357 |
| 1.513367 |

| |
|---|
| 1.434135 |
| -1.58427 |
| -1.4718 |
| 1.591905 |
| 1.349352 |
| 1.5508 |
| 1.392325 |
| 1.522735 |
| 1.445277 |
| 1.539942 |
| 1.337773 |
| 1.335921 |
| 1.583991 |
| 1.51333 |
| 1.371828 |
| -1.36549 |
| -1.38008 |
| 1.571995 |
| 1.323323 |
| 1.422193 |
| 1.449986 |
| -1.41168 |
| -1.33427 |
| -1.37523 |
| 1.486761 |
| 1.497627 |
| -1.54685 |
| -1.39619 |
| 1.541177 |
| 1.381539 |
| -1.49113 |
| -1.5112 |
| -1.50247 |
| 1.332465 |
| 1.469386 |
| 1.512002 |
| -1.29165 |
| -1.29151 |
| 1.466229 |
| -1.28699 |
| -1.37608 |
| 1.279363 |
| 1.415167 |
| 1.480174 |
| -1.43787 |
| 1.413553 |
| 1.300796 |

| |
|---|
| -1.44079 |
| 1.453868 |
| -1.36251 |
| -1.26764 |
| 1.482234 |
| -1.44968 |
| 1.467923 |
| -1.40078 |
| -1.40996 |
| 1.38433 |
| -1.41554 |
| -1.32957 |
| -1.44861 |
| 1.347428 |
| 1.425257 |
| -1.30059 |
| 1.248307 |
| 1.40605 |
| -1.38748 |
| -1.38989 |
| -1.2586 |
| 1.361297 |
| -1.27449 |
| -1.33801 |
| -1.35046 |
| 1.439442 |
| 1.429309 |
| -1.33671 |
| 1.302722 |
| 1.409778 |
| 1.409729 |
| 1.322935 |
| -1.3581 |
| -1.41537 |
| 1.224594 |
| -1.36036 |
| -1.3907 |
| -1.37451 |
| 1.295802 |
| 1.214007 |
| 1.25981 |
| -1.24532 |
| 1.35378 |
| -1.403 |
| -1.28628 |
| 1.249926 |
| -1.31208 |

| |
|---|
| -1.37915 |
| -1.3271 |
| -1.21476 |
| 1.19649 |
| 1.34851 |
| 1.201085 |
| 1.272626 |
| 1.348804 |
| 1.205637 |
| 1.381993 |
| 1.310901 |
| -1.36214 |
| 1.249493 |
| -1.29458 |
| -1.33966 |
| 1.279474 |
| -1.35887 |
| 1.300853 |
| -1.33389 |
| -1.1806 |
| 1.257186 |
| -1.29821 |
| -1.16495 |
| -1.28332 |
| -1.20824 |
| 1.196801 |
| 1.324877 |
| -1.28355 |
| -1.33692 |
| -1.32136 |
| -1.31253 |
| 1.227488 |
| 1.158228 |
| -1.20438 |
| 1.230396 |
| -1.3155 |
| -1.31704 |
| -1.31517 |
| -1.294 |
| 1.166676 |
| -1.28321 |
| -1.22814 |
| 1.314363 |
| -1.15404 |
| 1.288099 |
| 1.280628 |
| 1.276522 |

| |
|---|
| -1.28113 |
| 1.305206 |
| 1.152301 |
| -1.30069 |
| -1.15169 |
| 1.156734 |
| -1.18147 |
| 1.215138 |
| -1.25303 |
| 1.270865 |
| 1.120412 |
| 1.168018 |
| -1.24226 |
| 1.24796 |
| -1.10915 |
| -1.26262 |
| -1.17027 |
| -1.25037 |
| -1.11037 |
| 1.177163 |
| -1.21208 |
| -1.18069 |
| 1.239699 |
| -1.12514 |
| -1.17751 |
| -1.10234 |
| 1.223338 |
| -1.08322 |
| 1.204369 |
| -1.20989 |
| -1.12722 |
| 1.180132 |
| -1.20053 |
| -1.2113 |
| 1.186335 |
| -1.08643 |
| 1.092534 |
| -1.20061 |
| -1.13598 |
| -1.20116 |
| -1.14216 |
| 1.156737 |
| -1.14829 |
| -1.06411 |
| -1.17577 |
| -1.11765 |
| 1.038017 |

| |
|---|
| 1.153304 |
| -1.04084 |
| -1.16124 |
| 1.127883 |
| 1.130818 |
| -1.07707 |
| -1.16052 |
| -1.04955 |
| 1.103341 |
| -1.01514 |
| -1.0659 |
| 1.134549 |
| -1.05669 |
| -1.02777 |
| -1.0766 |
| 1.073201 |
| 1.005073 |
| 1.128361 |
| -1.05381 |
| 1.02027 |
| -1.10167 |
| -0.99929 |
| 1.069016 |
| -1.10499 |
| 1.042528 |
| -1.0792 |
| -1.0353 |
| 1.077108 |
| -1.03304 |
| -1.03216 |
| 0.987388 |
| -1.05707 |
| 1.078563 |
| 0.979835 |
| 1.017712 |
| 0.976039 |
| -0.98186 |
| 0.988086 |
| 1.060791 |
| 0.978285 |
| -1.02653 |
| -1.08403 |
| -1.08164 |
| -1.08902 |
| 1.050506 |
| 1.066419 |
| -1.06372 |

| |
|---|
| -1.07469 |
| 1.064032 |
| -0.99502 |
| -1.03151 |
| -0.98703 |
| 0.953244 |
| -1.04562 |
| -1.01003 |
| 1.00596 |
| -0.99927 |
| 1.044868 |
| 0.959697 |
| -1.02708 |
| -0.99607 |
| -0.92586 |
| 0.980932 |
| -0.94672 |
| 0.976437 |
| 0.996934 |
| 0.912558 |
| -1.02248 |
| -1.00416 |
| -1.00639 |
| -0.9214 |
| 0.923854 |
| 0.928015 |
| -0.89971 |
| 0.957665 |
| -0.95547 |
| 0.983727 |
| 0.985052 |
| 0.95275 |
| 0.892449 |
| 0.963546 |
| -0.93405 |
| 0.902748 |
| -0.93229 |
| -0.95422 |
| 0.895253 |
| -0.93054 |
| -0.94327 |
| 0.889814 |
| -0.94885 |
| -0.89343 |
| -0.94197 |
| -0.91165 |
| 0.926538 |

| |
|---|
| 0.938843 |
| -0.87439 |
| 0.919002 |
| -0.94025 |
| -0.93489 |
| -0.8862 |
| -0.91292 |
| 0.9291 |
| 0.869352 |
| 0.897924 |
| 0.839264 |
| 0.908366 |
| -0.84012 |
| -0.84437 |
| 0.915501 |
| -0.91583 |
| 0.833404 |
| 0.813537 |
| -0.83519 |
| -0.82474 |
| 0.851537 |
| 0.817624 |
| 0.820465 |
| 0.829937 |
| -0.8531 |
| -0.80747 |
| -0.80702 |
| 0.883782 |
| 0.885934 |
| -0.80368 |
| -0.82352 |
| -0.86036 |
| -0.84271 |
| 0.805287 |
| -0.80888 |
| -0.85513 |
| 0.868982 |
| -0.84913 |
| -0.84218 |
| 0.786803 |
| -0.84307 |
| -0.8229 |
| -0.85956 |
| 0.850336 |
| 0.83761 |
| 0.855761 |
| -0.81868 |

| |
|---|
| -0.80275 |
| -0.80858 |
| -0.84728 |
| -0.84122 |
| -0.76801 |
| 0.833753 |
| -0.76866 |
| 0.799517 |
| -0.76028 |
| 0.804905 |
| -0.80742 |
| 0.7574 |
| -0.82544 |
| 0.80679 |
| 0.746341 |
| -0.78636 |
| -0.81535 |
| 0.754059 |
| -0.81042 |
| -0.80682 |
| 0.800198 |
| 0.729343 |
| 0.774905 |
| -0.78309 |
| 0.78389 |
| 0.750703 |
| -0.77415 |
| -0.77068 |
| 0.707143 |
| -0.72406 |
| 0.756093 |
| -0.73357 |
| -0.72491 |
| 0.729666 |
| -0.73564 |
| 0.748784 |
| -0.69674 |
| 0.697472 |
| -0.73904 |
| 0.713754 |
| -0.69135 |
| -0.68715 |
| 0.736835 |
| 0.709964 |
| 0.72254 |
| -0.71754 |
| 0.732405 |

0.728476
0.721052
0.691646
-0.70661
-0.71169
-0.70622
-0.65427
-0.67923
-0.69985
0.698897
0.676953
0.691229
-0.66239
0.673491
-0.64242
-0.68293
0.649754
0.633605
0.657006
0.658039
0.667109
0.672966
-0.66007
0.650272
-0.64004
-0.6181
-0.64182
0.660103
0.654292
0.641028
0.631885
0.610478
-0.61598
0.651089
-0.60979
-0.6387
0.60026
-0.61183
-0.627
0.645447
0.623954
-0.61493
0.636727
-0.63084
0.589587
0.588183
-0.60973

| |
|---|
| 0.594243 |
| -0.61487 |
| -0.62256 |
| -0.60714 |
| 0.619396 |
| -0.62067 |
| -0.56914 |
| 0.607594 |
| -0.60503 |
| -0.61513 |
| -0.6148 |
| -0.61368 |
| 0.593362 |
| 0.56433 |
| 0.601773 |
| 0.570504 |
| -0.60849 |
| 0.599809 |
| -0.60661 |
| -0.59415 |
| 0.563122 |
| -0.56068 |
| -0.59575 |
| -0.58889 |
| -0.57784 |
| -0.57036 |
| 0.56277 |
| 0.595154 |
| 0.573357 |
| 0.560568 |
| -0.55513 |
| 0.587826 |
| -0.56421 |
| -0.56227 |
| 0.536116 |
| -0.56683 |
| 0.575998 |
| -0.56499 |
| 0.561728 |
| 0.562207 |
| 0.536118 |
| 0.545189 |
| 0.526817 |
| 0.551629 |
| -0.54585 |
| -0.55558 |
| -0.5471 |

| |
|---|
| -0.55172 |
| -0.5599 |
| -0.54382 |
| 0.513726 |
| -0.53245 |
| 0.551254 |
| -0.55145 |
| 0.535135 |
| 0.533856 |
| 0.501034 |
| 0.513924 |
| -0.505 |
| 0.502267 |
| 0.49646 |
| -0.51379 |
| -0.52638 |
| 0.525732 |
| -0.50529 |
| 0.48577 |
| 0.492263 |
| -0.49116 |
| -0.51257 |
| -0.49564 |
| -0.50438 |
| 0.507111 |
| -0.49407 |
| -0.50267 |
| 0.497896 |
| 0.496073 |
| -0.4818 |
| 0.467743 |
| 0.465945 |
| 0.478411 |
| -0.49221 |
| 0.488573 |
| 0.466055 |
| -0.45442 |
| -0.45815 |
| -0.4792 |
| -0.46766 |
| 0.455196 |
| -0.4823 |
| 0.478132 |
| 0.440577 |
| 0.454512 |
| -0.45121 |
| 0.434782 |

-0.43564
0.451523
0.452514
-0.43063
-0.44164
0.426419
0.421474
0.438517
0.442714
-0.44635
-0.42018
0.413534
-0.44009
-0.42311
0.415999
0.406748
-0.43333
-0.41733
0.410294
0.408953
-0.41878
0.422144
0.42634
0.401201
0.416909
-0.40862
-0.41482
-0.39225
-0.38472
-0.38701
-0.40802
0.397845
0.397862
-0.37696
0.400552
0.388387
-0.39043
0.377614
0.386282
0.39299
-0.39032
-0.39103
0.382063
-0.37215
0.380556
-0.36397
-0.387

| |
|---|
| -0.38406 |
| 0.358875 |
| 0.374618 |
| -0.36141 |
| 0.370526 |
| 0.36319 |
| -0.37382 |
| -0.36871 |
| 0.364784 |
| -0.35149 |
| -0.35722 |
| -0.36225 |
| 0.359492 |
| 0.346665 |
| 0.344441 |
| -0.338 |
| -0.34369 |
| -0.34413 |
| -0.35791 |
| -0.35211 |
| 0.353354 |
| -0.34674 |
| 0.336699 |
| -0.33408 |
| 0.32119 |
| -0.31236 |
| -0.33201 |
| 0.313588 |
| -0.32545 |
| -0.32683 |
| -0.30806 |
| 0.319499 |
| -0.3057 |
| 0.299839 |
| -0.30787 |
| -0.30753 |
| -0.31306 |
| -0.29647 |
| 0.286176 |
| 0.298873 |
| -0.29671 |
| 0.286913 |
| 0.27919 |
| 0.26815 |
| -0.26626 |
| -0.27602 |
| 0.277438 |

| |
|---|
| -0.26625 |
| -0.26108 |
| -0.25886 |
| -0.26871 |
| -0.26756 |
| 0.252528 |
| 0.263848 |
| 0.249528 |
| -0.25432 |
| -0.24626 |
| -0.25956 |
| -0.25858 |
| -0.24629 |
| 0.237381 |
| 0.237657 |
| 0.250351 |
| -0.2481 |
| 0.242113 |
| 0.240603 |
| 0.236044 |
| 0.237646 |
| -0.2324 |
| -0.22869 |
| -0.23138 |
| 0.221448 |
| 0.21486 |
| 0.21636 |
| -0.21777 |
| 0.197117 |
| 0.201954 |
| 0.199707 |
| 0.189495 |
| -0.18931 |
| -0.19154 |
| 0.190243 |
| -0.18771 |
| -0.18622 |
| 0.177511 |
| -0.18018 |
| -0.1807 |
| -0.17134 |
| 0.164729 |
| 0.167032 |
| -0.16244 |
| 0.166051 |
| -0.16724 |
| -0.15591 |

| |
|---|
| 0.155861 |
| 0.162966 |
| 0.149109 |
| -0.14554 |
| -0.15295 |
| -0.152 |
| -0.147 |
| -0.15281 |
| -0.14921 |
| 0.14657 |
| -0.14314 |
| 0.13894 |
| 0.139486 |
| 0.137516 |
| 0.137361 |
| 0.133763 |
| -0.12568 |
| 0.12535 |
| -0.12785 |
| 0.12619 |
| -0.12573 |
| -0.12538 |
| -0.12367 |
| 0.119369 |
| -0.11412 |
| -0.1189 |
| 0.115416 |
| 0.115138 |
| -0.10972 |
| -0.1039 |
| -0.10197 |
| -0.10418 |
| -0.0955 |
| 0.098714 |
| -0.09456 |
| 0.09315 |
| 0.092357 |
| 0.092372 |
| 0.083624 |
| 0.082512 |
| -0.0782 |
| 0.078282 |
| 0.071487 |
| 0.071024 |
| 0.070977 |
| -0.06929 |
| 0.070508 |

| |
|---|
| 0.066879 |
| -0.06901 |
| -0.06588 |
| -0.06663 |
| -0.06255 |
| -0.06082 |
| -0.06135 |
| 0.056172 |
| -0.05783 |
| 0.054149 |
| -0.05674 |
| 0.052705 |
| -0.04844 |
| -0.04942 |
| 0.046444 |
| 0.044116 |
| 0.046253 |
| 0.043262 |
| -0.04016 |
| 0.038664 |
| 0.038577 |
| -0.03536 |
| 0.03433 |
| -0.03353 |
| -0.03284 |
| 0.034108 |
| 0.034003 |
| 0.024764 |
| -0.02329 |
| -0.02262 |
| 0.019732 |
| 0.018103 |
| -0.01665 |
| -0.01702 |
| -0.01591 |
| 0.015484 |
| 0.014827 |
| -0.01426 |
| -0.01158 |
| 0.01078 |
| 0.010217 |
| -0.00962 |
| 0.006956 |
| 0.005967 |
| 0.004812 |
| 0.004895 |
| -0.00333 |

-0.00318
-0.00099
