## Supplemental Table 2 for "Evidence for high-frequency parallel evolution in virulent A. *baumannii* cultures"

| Protein | Parental | Opaque | Log2FC | p-value | t-value |
|---|---|---|---|---|---|
| peptidase M48 family protein | 1.053808 | 0.940081736 | -0.16475 | 9.81E-06 | -29.0865 |
| NAD(P) transhydrogenase sub | 0.815779 | 1.483661267 | 0.862911 | 1.47E-05 | 48.97277 |
| nitroreductase family protein | 1.077242 | 0.980104451 | -0.13633 | 2.55E-05 | -22.1086 |
| adenosylmethionine-8-amino | 1.152658 | 0.946782193 | -0.28386 | 2.69E-05 | -21.7259 |
| NAD(P) transhydrogenase sub | 0.661442 | 1.471203263 | 1.153309 | 3.14E-05 | 21.4618 |
| hypothetical protein ABUW_2 | 1.202753 | 0.858372427 | -0.48667 | 3.17E-05 | -38.3595 |
| toluene tolerance protein | 1.076256 | 0.902854374 | -0.25346 | 4.05E-05 | -35.1159 |
| hypothetical protein ABUW_3 | 1.573988 | 0.654522676 | -1.26591 | 4.25E-05 | -19.4279 |
| acyl-CoA dehydrogenase...172 | 0.889293 | 1.071638721 | 0.269087 | 4.71E-05 | 24.9832 |
| D-amino acid dehydrogenase | 1.152646 | 0.72332394 | -0.67224 | 5.34E-05 | -22.6401 |
| urocanate hydratase | 1.092899 | 0.97938713 | -0.15821 | 7E-05 | -23.7484 |
| putative metal-dependent hy | 0.873425 | 1.493659564 | 0.774096 | 8.14E-05 | 17.28753 |
| alkyl hydroperoxide reductase | 1.097743 | 0.97699887 | -0.16811 | 9.3E-05 | -17.3068 |
| trehalose-6-phosphate syntha | 0.598205 | 1.102187653 | 0.881657 | 0.000117 | 22.20465 |
| TonB-dependent copper recep | 0.796835 | 1.008339989 | 0.339628 | 0.000142 | 14.39598 |
| cytochrome O ubiquinol oxida | 0.900745 | 1.02481289 | 0.18617 | 0.00015 | 16.5392 |
| peptidase S45, penicillin amid | 1.014454 | 0.914863981 | -0.14907 | 0.000154 | -16.3935 |
| porphobilinogen deaminase | 1.022203 | 0.946733375 | -0.11065 | 0.000173 | -14.2243 |
| NAD-dependent malic enzyme | 1.06345 | 0.778628167 | -0.44975 | 0.000175 | -14.9823 |
| NAD(P) transhydrogenase sub | 0.654728 | 1.736672811 | 1.407359 | 0.000197 | 23.00105 |
| fructose-1,6-bisphosphatase | 1.86743 | 0.599911842 | -1.63823 | 0.000254 | -40.2144 |
| polysaccharide deacetylase...1 | 0.907838 | 1.178614916 | 0.376586 | 0.000292 | 13.62946 |
| phosphoribosylamine--glycine | 0.970952 | 0.927438116 | -0.06615 | 0.000359 | -13.2289 |
| integration host factor, alpha | 1.1308 | 0.926290844 | -0.28781 | 0.000365 | -13.7185 |
| lytic murein transglycosylase | 1.022771 | 0.964382454 | -0.08481 | 0.000403 | -13.5733 |
| homocysteine S-methyltransf | 1.044475 | 0.91544996 | -0.19023 | 0.000403 | -13.9793 |
| L-asparaginase 2 | 1.164188 | 0.986161603 | -0.23943 | 0.000443 | -10.6335 |
| modulator of drug activity B | 1.195831 | 0.857864751 | -0.47919 | 0.000455 | -10.8233 |
| C4-dicarboxylate transport pr | 0.973388 | 0.880686859 | -0.14439 | 0.000468 | -11.2818 |
| high affinity gluconate perme | 0.938148 | 1.105048329 | 0.236222 | 0.000499 | 10.3665 |
| hypothetical protein ABUW_1 | 1.09868 | 0.86828291 | -0.33953 | 0.000506 | -13.3344 |
| biotin synthase | 1.255587 | 0.861996333 | -0.54261 | 0.000521 | -10.4547 |
| acetyl-CoA carboxylase, carbo | 1.03775 | 0.898990007 | -0.20708 | 0.000554 | -11.7149 |
| hypothetical protein ABUW_1 | 0.953831 | 1.389966327 | 0.543244 | 0.000568 | 11.60975 |
| fatty acid desaturase...479 | 0.89948 | 1.064878521 | 0.243526 | 0.000691 | 9.503611 |
| hypothetical protein ABUW_0 | 1.142787 | 0.876181045 | -0.38326 | 0.000697 | -10.092 |
| acetyl-CoA carboxylase, carbo | 1.065488 | 0.839651871 | -0.34365 | 0.000734 | -9.34839 |
| hypothetical protein ABUW_2 | 0.63754 | 1.185234582 | 0.894586 | 0.000745 | 9.651705 |
| preprotein translocase, SecY s | 0.828511 | 1.017297664 | 0.29615 | 0.00075 | 9.28577 |
| ribosomal protein L29 | 1.055713 | 0.94088493 | -0.16613 | 0.000847 | -9.08267 |
| isocitrate dehydrogenase, NA | 0.999556 | 0.851733251 | -0.23089 | 0.000863 | -10.3057 |
| lipoic acid synthetase...1311 | 1.056312 | 0.945719595 | -0.15955 | 0.000964 | -8.82215 |
| putative integrase/recombina | 0.900671 | 1.309792226 | 0.540266 | 0.000964 | 8.741093 |
| fatty oxidation complex, alpha | 1.074211 | 0.890424264 | -0.27071 | 0.000993 | -8.72355 |
| chaperonin GroS | 1.084276 | 0.876732133 | -0.30652 | 0.001004 | -10.8192 |
| hypothetical protein ABUW_1 | 1.180516 | 0.78508609 | -0.5885 | 0.00103 | -8.56125 |

| | | | | | |
|---|---|---|---|---|---|
| acyl-CoA dehydrogenase...167 | 1.059447 | 0.98048451 | -0.11174 | 0.001043 | -8.84942 |
| L-sorbosone dehydrogenase.. | 0.983957 | 1.104101031 | 0.166205 | 0.001085 | 9.162476 |
| NADPH-dependent fmn reduc | 0.896084 | 1.116270162 | 0.316981 | 0.001104 | 17.24202 |
| hypothetical protein ABUW_1 | 0.948569 | 1.131055712 | 0.253845 | 0.001132 | 9.797805 |
| tyrosyl-tRNA synthetase | 0.963168 | 0.919164915 | -0.06746 | 0.001146 | -10.387 |
| hypothetical protein ABUW_1 | 0.994974 | 1.054012902 | 0.083162 | 0.001171 | 8.321994 |
| undecaprenyl diphosphate sy | 1.172047 | 0.952955648 | -0.29855 | 0.001224 | -8.19296 |
| thioesterase superfamily prot | 1.297893 | 0.826114475 | -0.65176 | 0.001229 | -8.18384 |
| acetyl-CoA carboxylase, biotir | 1.091196 | 0.903344921 | -0.27256 | 0.001238 | -19.1657 |
| threonyl-tRNA synthetase | 0.98643 | 0.935948762 | -0.07579 | 0.001261 | -10.1359 |
| hypothetical protein ABUW_1 | 1.047196 | 0.947578584 | -0.14421 | 0.001338 | -9.51015 |
| benzoate transport porin Ben | 0.957213 | 0.836766747 | -0.19401 | 0.001338 | -15.6817 |
| dethiobiotin synthase | 1.102959 | 0.932915126 | -0.24156 | 0.001372 | -10.883 |
| hypothetical protein ABUW_2 | 0.871427 | 1.115460246 | 0.356188 | 0.001396 | 9.098906 |
| ATP phosphoribosyltransferas | 1.026074 | 0.939275911 | -0.12751 | 0.001415 | -15.5425 |
| transcriptional regulator, Ara( | 1.135186 | 0.870358648 | -0.38325 | 0.001469 | -16.2424 |
| hypothetical protein ABUW_0 | 0.939014 | 1.175686654 | 0.324286 | 0.001568 | 9.961073 |
| peptidase M16 domain protei | 1.123473 | 0.979017931 | -0.19856 | 0.001622 | -13.9475 |
| electron transfer flavoprotein | 1.01957 | 0.926829225 | -0.13759 | 0.001686 | -14.2673 |
| metallo-beta-lactamase famil | 0.670771 | 1.265290449 | 0.915576 | 0.001724 | 10.23584 |
| peptidase S1C, Do | 1.058665 | 1.001379703 | -0.08026 | 0.001839 | -10.2238 |
| putative anti-anti-sigma facto | 0.714768 | 1.205204896 | 0.753732 | 0.001846 | 15.00386 |
| bifunctional protein wax ester | 0.946164 | 1.036201476 | 0.131143 | 0.001877 | 7.816674 |
| methyltransferase | 0.956924 | 0.784209983 | -0.28716 | 0.001945 | -9.59191 |
| efflux transporter, RND family | 1.1177 | 0.889833683 | -0.32893 | 0.001995 | -11.4822 |
| hypothetical protein ABUW_0 | 1.034212 | 0.951794014 | -0.11981 | 0.002152 | -7.87894 |
| glutamine synthetase, type I | 0.821895 | 1.090034312 | 0.407347 | 0.002167 | 7.03991 |
| FAD-dependent pyridine nucl | 0.917815 | 1.078034456 | 0.232128 | 0.002293 | 6.916133 |
| transcription elongation facto | 1.056464 | 0.944843436 | -0.1611 | 0.00233 | -7.71589 |
| 3-hydroxybutyrate dehydroge | 1.025169 | 0.844612585 | -0.2795 | 0.002338 | -9.85493 |
| ribosomal large subunit pseud | 0.867869 | 1.052461319 | 0.278217 | 0.002381 | 8.569463 |
| carbonate dehydratase | 1.030087 | 0.901555181 | -0.19228 | 0.002405 | -6.86377 |
| hypothetical protein ABUW_3 | 1.084959 | 0.959961386 | -0.17659 | 0.002499 | -12.8861 |
| acyl-CoA synthase...1459 | 0.927721 | 1.065716433 | 0.200061 | 0.002502 | 8.561712 |
| quinoprotein glucose dehydro | 0.929492 | 1.052121876 | 0.178787 | 0.002554 | 8.954607 |
| NADH dehydrogenase I chain | 0.939555 | 1.012892751 | 0.108433 | 0.002557 | 7.187263 |
| dihydroxy-acid dehydratase... | 1.156939 | 0.892338178 | -0.37465 | 0.002576 | -6.83863 |
| 2-methylcitrate synthase | 0.893626 | 1.004940762 | 0.169367 | 0.002619 | 6.845888 |
| dihydroxy-acid dehydratase... | 0.939833 | 0.974184362 | 0.05179 | 0.00264 | 8.837876 |
| transcriptional regulator, Ara( | 1.105759 | 0.955081566 | -0.21134 | 0.002651 | -6.7648 |
| preprotein translocase iisp far | 0.989236 | 1.126358419 | 0.187279 | 0.002655 | 9.24596 |
| formate/nitrite transporter fa | 0.935471 | 1.102690781 | 0.237264 | 0.002668 | 7.47061 |
| trehalose-phosphatase | 0.679279 | 0.997217109 | 0.553903 | 0.002671 | 11.5225 |
| aldehyde dehydrogenase...23 | 1.006084 | 1.079585456 | 0.101727 | 0.002774 | 7.968087 |
| C4-dicarboxylate transport pr | 0.906342 | 0.63095912 | -0.52251 | 0.002823 | -7.0145 |
| hypothetical protein ABUW_0 | 0.895801 | 1.215925849 | 0.440805 | 0.002829 | 9.471343 |
| hypothetical protein ABUW_1 | 0.904092 | 1.072086622 | 0.24588 | 0.002837 | 7.38061 |

| | | | | | |
|---|---|---|---|---|---|
| hypothetical protein ABUW_2 | 0.898074 | 1.112868301 | 0.309377 | 0.002879 | 6.614422 |
| trigger factor | 1.095802 | 0.923221852 | -0.24724 | 0.00298 | -8.69233 |
| NADP-specific glutamate dehy | 0.855478 | 0.996450456 | 0.220068 | 0.003055 | 7.219971 |
| TonB protein | 0.798785 | 0.938110199 | 0.231951 | 0.003163 | 6.343937 |
| hypothetical protein ABUW_0 | 1.443132 | 0.731998226 | -0.97929 | 0.003179 | -9.34338 |
| TonB dependent outer memb | 0.801769 | 0.726641208 | -0.14194 | 0.003197 | -6.37504 |
| diaminopimelate epimerase | 1.079174 | 0.927036118 | -0.21923 | 0.003205 | -6.7513 |
| hypothetical protein ABUW_3 | 1.209631 | 0.885531403 | -0.44995 | 0.003225 | -6.32473 |
| Fe(II) trafficking protein | 1.017742 | 0.907413203 | -0.16554 | 0.003249 | -6.77099 |
| cystathionine beta-lyase PLP-( | 0.895122 | 1.081383341 | 0.272721 | 0.003257 | 8.785221 |
| ribosome-binding factor A | 1.045305 | 0.973684531 | -0.1024 | 0.003362 | -7.11058 |
| 3-oxoacyl-(acyl-carrier-proteir | 1.077636 | 0.882008273 | -0.28901 | 0.0034 | -10.4982 |
| transcriptional regulator, Moc | 0.959947 | 1.123907473 | 0.227496 | 0.003437 | 6.374895 |
| translation initiation factor IF- | 1.019469 | 0.965855375 | -0.07794 | 0.003674 | -8.30283 |
| hypothetical protein ABUW_1 | 1.045897 | 1.314538176 | 0.329815 | 0.003703 | 6.090434 |
| histidine triad protein...1597 | 1.129093 | 0.900093906 | -0.32702 | 0.003748 | -8.72309 |
| RHS family protein | 0.930686 | 1.067681395 | 0.198114 | 0.003839 | 6.181173 |
| nonribosomal peptide synthe | 1.070435 | 0.868186609 | -0.30212 | 0.003845 | -6.13746 |
| ribosomal protein L19 | 1.052054 | 0.913651659 | -0.20349 | 0.003938 | -6.3498 |
| RDD family protein | 0.949147 | 1.072940733 | 0.176867 | 0.003983 | 5.962417 |
| dihydrofolate reductase (plas | 0.903813 | 0.982346035 | 0.120206 | 0.004101 | 6.028112 |
| thioesterase domain protein | 0.76874 | 1.003839243 | 0.384961 | 0.004203 | 6.011597 |
| ErfK/YbiS/YcfS/YnhG family | 0.837531 | 1.104531946 | 0.399221 | 0.004227 | 6.338942 |
| ribosome recycling factor | 1.013062 | 0.966805494 | -0.06742 | 0.004319 | -6.71574 |
| ribosomal RNA large subunit r | 1.052319 | 0.939633476 | -0.1634 | 0.004331 | -9.35585 |
| transcriptional regulator, IclR | 0.884929 | 1.094263695 | 0.306327 | 0.004415 | 6.004227 |
| hypothetical protein ABUW_0 | 0.865878 | 1.134632631 | 0.38999 | 0.004472 | 6.700778 |
| histidyl-tRNA synthetase | 0.95053 | 1.026159567 | 0.110452 | 0.004609 | 6.203331 |
| heme oxygenase-like protein. | 0.707304 | 1.181546558 | 0.740274 | 0.004713 | 11.56137 |
| hypothetical protein ABUW_0 | 0.836456 | 1.158196272 | 0.469519 | 0.004775 | 5.697192 |
| phosphopantothenoylcysteine | 0.826164 | 1.166290028 | 0.497426 | 0.004784 | 8.5738 |
| hypothetical protein ABUW_3 | 0.918846 | 1.177528999 | 0.357868 | 0.004868 | 12.10372 |
| phage shock protein C | 1.019347 | 0.787941528 | -0.37149 | 0.004879 | -5.75001 |
| short-chain fatty acid transpo | 0.769682 | 0.9793288 | 0.347531 | 0.004892 | 6.625103 |
| thiamine biosynthesis protein | 1.537656 | 0.584595874 | -1.39522 | 0.004956 | -8.27359 |
| glutamyl-tRNA(Gln) amidotrai | 0.94812 | 0.971297592 | 0.034844 | 0.005008 | 6.966196 |
| pseudouridine synthase, Rsu | 0.922563 | 1.018499709 | 0.142726 | 0.005226 | 5.585701 |
| sporulation initiation inhibitor | 1.053586 | 0.921096492 | -0.19388 | 0.005443 | -8.27425 |
| poly(A) polymerase | 0.873884 | 1.083224168 | 0.309819 | 0.00569 | 6.141372 |
| glutathione S-transferase...59 | 1.011351 | 0.966426245 | -0.06555 | 0.005731 | -5.75581 |
| ribosomal protein L36 | 1.542741 | 0.786633908 | -0.97173 | 0.005871 | -8.771 |
| ribonuclease E | 0.98063 | 1.036883734 | 0.080473 | 0.005945 | 6.48585 |
| hypothetical protein ABUW_0 | 0.878696 | 1.175125445 | 0.419379 | 0.005955 | 11.30075 |
| hypothetical protein ABUW_3 | 1.09035 | 0.938361459 | -0.21657 | 0.006048 | -5.47174 |
| sulfate transporter | 0.880236 | 0.985476031 | 0.162931 | 0.006106 | 7.024265 |
| ribosomal protein S6 | 1.029077 | 0.975568811 | -0.07703 | 0.00619 | -6.24582 |
| guanylate kinase | 1.11363 | 1.037551686 | -0.10209 | 0.006297 | -5.79587 |

| | | | | | |
|---|---|---|---|---|---|
| hypothetical protein ABUW_1 | 1.146205 | 0.82328569 | -0.4774 | 0.006351 | -6.83789 |
| D-serine/D-alanine/glycine tra | 0.912298 | 1.095879271 | 0.264512 | 0.006377 | 5.231792 |
| hypothetical protein ABUW_2 | 1.067311 | 0.960708317 | -0.15181 | 0.006437 | -5.51306 |
| hypothetical protein ABUW_3 | 0.627074 | 1.354927329 | 1.111508 | 0.006509 | 6.595032 |
| ferric acinetobactin receptor | 0.986306 | 0.891816483 | -0.14529 | 0.006523 | -9.57437 |
| hypothetical protein ABUW_1 | 1.060625 | 0.911338616 | -0.21886 | 0.006553 | -5.1939 |
| hypothetical protein ABUW_0 | 1.007158 | 1.473284833 | 0.548747 | 0.006717 | 5.7485 |
| L-serine dehydratase | 0.995373 | 1.25938932 | 0.339415 | 0.006755 | 5.30861 |
| ribosomal protein L35 | 1.146016 | 0.935678394 | -0.29254 | 0.006755 | -5.79051 |
| hypothetical protein ABUW_3 | 1.057698 | 0.751976551 | -0.49217 | 0.006835 | -5.24258 |
| hypoxanthine phosphoribosyl | 1.077212 | 1.003806608 | -0.10182 | 0.006859 | -5.14833 |
| signal peptide peptidase SppA | 0.925729 | 1.041440111 | 0.169918 | 0.006872 | 5.992787 |
| multidrug resistance protein E | 1.219702 | 0.911576773 | -0.42009 | 0.006949 | -5.19085 |
| RNA polymerase sigma factor | 1.024387 | 0.939569006 | -0.12469 | 0.006971 | -6.74376 |
| hypothetical protein ABUW_0 | 0.974558 | 1.023798971 | 0.071113 | 0.007068 | 5.131727 |
| phospholipid/glycerol acyltrar | 0.972935 | 0.899526001 | -0.11318 | 0.007091 | -5.77231 |
| nudix hydrolase...683 | 0.927435 | 1.064071816 | 0.198277 | 0.007131 | 6.397079 |
| ABC-type nitrate/sulfonate/bi | 0.999178 | 0.817760921 | -0.28906 | 0.007146 | -5.60771 |
| ribosomal L25 | 1.058089 | 0.907998012 | -0.2207 | 0.007172 | -5.14695 |
| imidazole glycerol phosphate | 0.943585 | 1.048001086 | 0.151416 | 0.007179 | 5.078332 |
| acyl-CoA dehydrogenase...232 | 0.98918 | 1.204970715 | 0.284694 | 0.007243 | 8.103097 |
| PGAP1 family protein | 0.993177 | 1.088818019 | 0.132641 | 0.007546 | 5.09152 |
| hypothetical protein ABUW_0 | 1.061435 | 0.859798013 | -0.30395 | 0.007595 | -10.7143 |
| transcriptional regulator, TetR | 1.055953 | 0.938304292 | -0.17042 | 0.007629 | -4.98668 |
| Fe-S protein assembly co-chap | 1.038331 | 0.940170193 | -0.14327 | 0.007642 | -5.7013 |
| hypothetical protein ABUW_3 | 1.147605 | 0.896602862 | -0.35608 | 0.007727 | -9.21672 |
| putative UDP-galactose phosp | 1.015869 | 1.221160888 | 0.265539 | 0.007739 | 5.669633 |
| long-chain fatty-acid-CoA ligas | 1.002882 | 0.89942436 | -0.15708 | 0.007839 | -7.47867 |
| phosphoribosylanthranilate is | 0.965052 | 1.051355558 | 0.123572 | 0.007882 | 5.14759 |
| family 1 glycosyl transferase.. | 1.032032 | 1.075839184 | 0.059975 | 0.008025 | 5.708605 |
| invasion protein expression u | 0.793335 | 1.026916943 | 0.372317 | 0.008377 | 10.23342 |
| hydrolase, alpha/beta fold far | 1.065542 | 1.176554244 | 0.142981 | 0.008403 | 5.806195 |
| amino acid transport protein | 0.832522 | 1.062056912 | 0.3513 | 0.008428 | 5.847601 |
| polysaccharide deacetylase...: | 0.865635 | 1.205946894 | 0.478336 | 0.008531 | 5.73239 |
| glutamate-1-semialdehyde-2, | 0.982233 | 0.942175405 | -0.06007 | 0.008702 | -7.17256 |
| acetyl-CoA C-acyltransferase I | 1.03656 | 0.91157664 | -0.18537 | 0.008742 | -7.77195 |
| phospholipase C, phosphocho | 1.187926 | 0.884045874 | -0.42625 | 0.008901 | -6.44313 |
| hypothetical protein ABUW_2 | 0.997063 | 0.95091566 | -0.06837 | 0.008965 | -5.36703 |
| Cu(I)-responsive transcription | 1.142888 | 0.950304834 | -0.26622 | 0.009037 | -4.89283 |
| saccharopine dehydrogenase | 1.059102 | 1.029964313 | -0.04025 | 0.00906 | -4.78097 |
| putative RNA methylase | 0.924187 | 1.0036462 | 0.118994 | 0.00908 | 4.827998 |
| hypothetical protein ABUW_2 | 1.990196 | 1.006897394 | -0.98299 | 0.00912 | -4.8083 |
| chaperonin GroL | 1.109916 | 0.918480052 | -0.27313 | 0.009151 | -6.44508 |
| maleylacetoacetate isomerase | 0.926042 | 0.99442469 | 0.102785 | 0.009202 | 5.041444 |
| glucose dehydrogenase | 0.985426 | 1.146493919 | 0.21841 | 0.009218 | 8.022169 |
| DNA topoisomerase I | 0.984804 | 1.036621149 | 0.07398 | 0.00932 | 5.757096 |
| alpha/beta hydrolase fold pro | 1.136326 | 0.869620874 | -0.38592 | 0.009347 | -4.86928 |

| | | | | | |
|---|---|---|---|---|---|
| OmpA/MotB domain protein. | 0.869968 | 1.327669165 | 0.609862 | 0.009414 | 5.401961 |
| excinuclease ABC, C subunit (I | 1.08821 | 1.052710325 | -0.04785 | 0.009539 | -4.68091 |
| cytochrome bd ubiquinol oxic | 0.917419 | 1.220482353 | 0.411798 | 0.009659 | 9.349607 |
| putative VGR-related protein | 0.911683 | 1.110087547 | 0.284069 | 0.009766 | 7.415845 |
| Heavy metal transport/detoxi | 1.123177 | 0.918597269 | -0.29008 | 0.009863 | -6.43773 |
| hypothetical protein ABUW_( | 1.003404 | 0.876480496 | -0.19511 | 0.009902 | -4.71937 |
| peptidase M24 | 1.082368 | 1.011091585 | -0.09828 | 0.009985 | -4.64875 |
| TatD-related deoxyribonuclea | 1.040793 | 0.909412837 | -0.19468 | 0.010097 | -4.927 |
| hypothetical protein ABUW_( | 0.925666 | 1.075174031 | 0.216006 | 0.010261 | 4.641225 |
| hypothetical protein ABUW_( | 1.078889 | 0.913897029 | -0.23944 | 0.010573 | -6.12091 |
| NADH-dependent enoyl-ACP r | 1.057397 | 0.8827611 | -0.26042 | 0.010778 | -9.42376 |
| MFS permease | 0.958007 | 1.180992904 | 0.301892 | 0.010881 | 6.416509 |
| phosphoenolpyruvate carbox | 0.871378 | 1.08438856 | 0.315511 | 0.010922 | 6.052917 |
| 3-deoxy-D-manno-octulosona | 0.967142 | 1.046389599 | 0.11362 | 0.010943 | 4.489544 |
| delta-1-pyrroline-5-carboxylat | 0.901551 | 1.078999234 | 0.259213 | 0.01096 | 5.694113 |
| hypothetical protein ABUW_( | 0.889605 | 1.004972857 | 0.175919 | 0.010978 | 6.809989 |
| hypothetical protein ABUW_1 | 1.00675 | 0.944515923 | -0.09206 | 0.01115 | -5.72773 |
| translation initiation factor IF- | 1.024668 | 0.9782835 | -0.06683 | 0.011192 | -4.95696 |
| ribosomal protein L3 | 1.030325 | 0.923901234 | -0.15729 | 0.011207 | -4.98118 |
| hypothetical protein ABUW_( | 1.150575 | 0.859492964 | -0.4208 | 0.011277 | -7.80176 |
| cell division protein ZipA | 0.830188 | 0.933362105 | 0.168999 | 0.011508 | 4.420916 |
| ornithine carbamoyltransfera | 1.14481 | 1.03364178 | -0.14737 | 0.011534 | -6.97136 |
| putative phosphodiesterase | 0.959225 | 1.109375173 | 0.209807 | 0.011732 | 4.689536 |
| 2-nitropropane dioxygenase.. | 1.068761 | 0.932402555 | -0.19691 | 0.011773 | -4.39282 |
| CsuD | 0.870939 | 1.016447829 | 0.222892 | 0.011821 | 8.640859 |
| iron-containing alcohol dehyd | 1.003541 | 1.05708452 | 0.074991 | 0.011881 | 4.400019 |
| cell division protein FtsA | 0.911436 | 1.01622715 | 0.15701 | 0.012247 | 4.389706 |
| hypothetical protein ABUW_( | 0.98074 | 1.062655431 | 0.115731 | 0.012289 | 4.468926 |
| ATP-dependent dsDNA exonu | 0.891998 | 1.410580844 | 0.661177 | 0.012456 | 6.273138 |
| hypothetical protein ABUW_4 | 1.802691 | 0.703873289 | -1.35676 | 0.012593 | -7.38864 |
| hypothetical protein ABUW_3 | 0.919729 | 0.962741649 | 0.06594 | 0.012663 | 4.349925 |
| ATP-dependent chaperone Cl | 1.113424 | 1.000801077 | -0.15385 | 0.012699 | -6.99381 |
| hypothetical protein ABUW_1 | 1.208003 | 0.90747015 | -0.4127 | 0.012781 | -8.42749 |
| malonyl CoA-acyl carrier prot | 1.095974 | 0.89687744 | -0.28923 | 0.012904 | -7.81924 |
| hypothetical protein ABUW_1 | 0.791247 | 1.460910567 | 0.884667 | 0.012923 | 4.833947 |
| peptidase | 0.987955 | 1.094179586 | 0.147333 | 0.012961 | 6.531914 |
| membrane spanning export p | 1.015875 | 1.065648186 | 0.069008 | 0.013107 | 5.996134 |
| formamidopyrimidine-DNA gl | 1.070624 | 0.951790187 | -0.16974 | 0.013142 | -4.50704 |
| seryl-tRNA synthetase | 0.98402 | 0.937475139 | -0.06991 | 0.013196 | -4.26424 |
| chloramphenicol resistance p | 1.065341 | 0.973663447 | -0.12982 | 0.013239 | -4.33752 |
| nudix hydrolase...687 | 0.986904 | 0.943439569 | -0.06498 | 0.0133 | -5.20353 |
| hypothetical protein ABUW_3 | 0.953775 | 1.017993786 | 0.094008 | 0.013425 | 5.898248 |
| phenylacetic acid degradatior | 0.8246 | 1.016435021 | 0.301751 | 0.013746 | 5.377186 |
| [protein-PII] uridylyltransfera | 0.94344 | 0.987077116 | 0.065232 | 0.013945 | 4.651996 |
| glutathione S-transferase...18 | 1.048662 | 0.995688605 | -0.07478 | 0.01396 | -4.87958 |
| beta-ketoadipyl CoA thiolase. | 1.057614 | 0.905454634 | -0.2241 | 0.014034 | -6.15395 |
| thymidylate synthase | 0.939828 | 1.02639508 | 0.127118 | 0.014318 | 6.387947 |

| | | | | | |
|---|---|---|---|---|---|
| peroxidase | 1.087724 | 0.981336149 | -0.14849 | 0.01447 | -4.15997 |
| chromosome segregation pro | 0.804839 | 1.313220054 | 0.706337 | 0.014641 | 4.151161 |
| chaperone protein HtpG | 1.154684 | 0.912355853 | -0.33983 | 0.014728 | -5.72792 |
| Holliday junction DNA helicase | 1.269212 | 0.983832095 | -0.36745 | 0.014737 | -5.19432 |
| 3-oxoacid CoA-transferase, su | 1.036377 | 0.820297089 | -0.33733 | 0.014754 | -6.75767 |
| helicase domain-containing te | 1.139249 | 0.81436003 | -0.48434 | 0.014967 | -4.2653 |
| peptide deformylase...1954 | 0.999994 | 0.916759041 | -0.12538 | 0.014985 | -4.41127 |
| oxidoreductase short chain de | 1.217599 | 1.043974995 | -0.22195 | 0.015119 | -7.16943 |
| PTS system fructose-specific E | 1.109621 | 0.96402199 | -0.20293 | 0.015295 | -4.22929 |
| hypothetical protein ABUW_2 | 0.702711 | 1.344816875 | 0.936407 | 0.015691 | 4.802729 |
| L-sorbosone dehydrogenase.. | 0.941002 | 1.122246139 | 0.254119 | 0.015829 | 7.715428 |
| alanyl-tRNA synthetase | 0.954443 | 1.010391052 | 0.082183 | 0.015935 | 5.279276 |
| ribose 5-phosphate isomerase | 1.019203 | 0.929152435 | -0.13345 | 0.015979 | -6.45917 |
| hypothetical protein ABUW_3 | 1.019925 | 0.943486501 | -0.11239 | 0.016 | -6.28821 |
| ribosomal protein S14 | 1.149 | 0.896255788 | -0.3584 | 0.016104 | -6.76688 |
| delta-aminolevulinic acid deh | 0.894263 | 0.967142625 | 0.11303 | 0.016165 | 4.142672 |
| NADPH:quinone oxidoreducta | 0.976034 | 0.915660425 | -0.09212 | 0.016195 | -6.52422 |
| putative lysozyme | 1.162399 | 0.979669904 | -0.24674 | 0.01621 | -7.0408 |
| ribonuclease Z | 1.02501 | 1.167242851 | 0.187466 | 0.016229 | 4.399525 |
| transcriptional regulator, HxlR | 1.091383 | 0.941382991 | -0.2133 | 0.016343 | -4.40628 |
| acyl-CoA dehydrogenase...476 | 0.754832 | 1.058801975 | 0.488206 | 0.016363 | 3.982949 |
| hypothetical protein ABUW_0 | 1.142569 | 0.977186619 | -0.22558 | 0.016446 | -5.71386 |
| D-alanine--D-alanine ligase B | 1.001644 | 0.961486656 | -0.05903 | 0.016526 | -4.42613 |
| hydroxypyruvate isomerase | 1.057504 | 0.981055314 | -0.10826 | 0.016542 | -4.12726 |
| methylcrotonoyl-CoA carboxy | 1.048185 | 0.938190364 | -0.15994 | 0.016546 | -4.82832 |
| preprotein translocase, SecA s | 0.942704 | 0.961413619 | 0.028352 | 0.016547 | 4.548479 |
| citrate transporter | 0.702013 | 0.568625331 | -0.30402 | 0.016665 | -3.98119 |
| thiol:disulfide interchange pro | 1.003879 | 1.103056348 | 0.13592 | 0.016693 | 6.888007 |
| D-methionine transport prote | 0.867713 | 1.105695343 | 0.349664 | 0.016738 | 3.972026 |
| two-component system respo | 1.022221 | 0.943725151 | -0.11527 | 0.016882 | -4.00349 |
| protein-(glutamine-N5) methy | 1.010494 | 0.948335566 | -0.09159 | 0.016939 | -3.94158 |
| TonB-dependent siderophore | 1.053456 | 0.88414474 | -0.25278 | 0.01709 | -4.05674 |
| glycosyltransferase...577 | 0.898597 | 1.145705479 | 0.350489 | 0.017121 | 6.055379 |
| Csy3 family crispr-associated p | 0.966035 | 1.016872399 | 0.073991 | 0.017182 | 6.23231 |
| aminopeptidase N | 0.883055 | 1.092501249 | 0.30706 | 0.0172 | 4.835756 |
| hypothetical protein ABUW_0 | 0.93862 | 1.084005785 | 0.20776 | 0.01734 | 4.816959 |
| xanthine phosphoribosyltrans | 1.12819 | 0.967892822 | -0.22109 | 0.017373 | -4.69107 |
| hypothetical protein ABUW_3 | 0.934609 | 0.966659755 | 0.048645 | 0.01741 | 4.426026 |
| TonB-dependent receptor...22 | 0.928989 | 0.850631129 | -0.12713 | 0.01745 | -4.69537 |
| phosphomannomutase...1567 | 0.885014 | 1.086991302 | 0.296568 | 0.017454 | 6.627826 |
| putative L-asparaginase I | 1.022361 | 0.943388392 | -0.11598 | 0.017687 | -4.2328 |
| hypothetical protein ABUW_0 | 0.795035 | 1.034482813 | 0.379819 | 0.017703 | 4.104163 |
| phosphoribosylformimino-5-a | 0.985224 | 0.924902876 | -0.09115 | 0.017724 | -4.46051 |
| monooxygenase | 1.027612 | 0.958614033 | -0.10027 | 0.017764 | -5.63266 |
| hypothetical protein ABUW_2 | 0.892205 | 1.0556408 | 0.242671 | 0.017853 | 4.576181 |
| chaperone protein | 0.921488 | 1.011824227 | 0.134922 | 0.01796 | 5.030032 |
| Sua5/YciO/YrdC/YwlC family p | 1.051805 | 0.884849059 | -0.24936 | 0.01815 | -5.63111 |

| | | | | | |
|---|---|---|---|---|---|
| ribosomal protein S16 | 1.065097 | 0.956107745 | -0.15574 | 0.018281 | -5.07747 |
| 2-oxoglutarate dehydrogenas | 0.949615 | 0.904882449 | -0.06961 | 0.018382 | -4.89835 |
| hypothetical protein ABUW_( | 1.03563 | 0.89639799 | -0.2083 | 0.018502 | -5.64016 |
| aspartate carbamoyltransfera | 0.903075 | 0.986919729 | 0.128087 | 0.018554 | 5.205259 |
| hypothetical protein ABUW_3 | 0.988759 | 0.861046766 | -0.19953 | 0.018603 | -4.74496 |
| hypothetical protein ABUW_3 | 1.009164 | 0.907279484 | -0.15354 | 0.018645 | -4.00055 |
| hypothetical protein ABUW_1 | 0.770213 | 1.164188446 | 0.595996 | 0.018764 | 5.828866 |
| protein in PtsN-ptsO intergen | 0.980818 | 0.948317913 | -0.04861 | 0.018789 | -3.90167 |
| integral membrane protein DI | 1.122916 | 0.98684094 | -0.18636 | 0.018875 | -5.15264 |
| ferredoxin, 2Fe-2S type | 1.091077 | 0.92478067 | -0.23857 | 0.019165 | -6.48062 |
| isocitrate dehydrogenase, NA | 1.100027 | 0.818946513 | -0.4257 | 0.019171 | -7.10842 |
| transcriptional regulator, Ara( | 0.965189 | 1.142735005 | 0.243607 | 0.019182 | 4.120474 |
| ribosomal protein S15 | 1.094606 | 0.977476084 | -0.16328 | 0.0192 | -3.87664 |
| hypothetical protein ABUW_( | 1.019514 | 0.94653621 | -0.10715 | 0.019467 | -4.98133 |
| CsuE | 0.873392 | 0.972846139 | 0.155582 | 0.019656 | 3.829031 |
| homoserine dehydrogenase... | 0.914217 | 0.850665878 | -0.10394 | 0.019697 | -3.76484 |
| succinate dehydrogenase, flav | 0.894669 | 0.93717329 | 0.066961 | 0.019931 | 4.400158 |
| outer-membrane receptor for | 0.853238 | 0.939979913 | 0.139681 | 0.02007 | 3.841038 |
| hypothetical protein ABUW_4 | 1.048452 | 0.970173267 | -0.11195 | 0.020144 | -4.95454 |
| arginine N-succinyltransferase | 0.938537 | 1.132668468 | 0.27124 | 0.020346 | 6.27018 |
| molybdenum cofactor biosynt | 0.90742 | 1.043229488 | 0.201214 | 0.020418 | 4.742474 |
| transcriptional regulator, Ten/ | 1.028551 | 0.959373226 | -0.10045 | 0.020447 | -3.82044 |
| hypothetical protein ABUW_1 | 0.645648 | 1.267237596 | 0.972868 | 0.020943 | 4.520594 |
| phospholipid/glycerol acyltrai | 1.077021 | 0.938788071 | -0.19818 | 0.021016 | -4.10695 |
| immunoreactive 21 kD antige | 0.889502 | 1.019205055 | 0.196374 | 0.021114 | 4.162901 |
| nitrogen regulatory protein P- | 1.00755 | 0.92271705 | -0.12689 | 0.021159 | -4.42637 |
| 17 kDa surface antigen | 1.088703 | 0.964074169 | -0.17539 | 0.021281 | -4.27912 |
| UspA domain protein | 1.250018 | 0.952136072 | -0.39271 | 0.021321 | -4.68977 |
| hypothetical protein ABUW_2 | 0.884281 | 1.017928124 | 0.203059 | 0.021353 | 3.699601 |
| thiol:disulfide interchange prc | 1.002361 | 0.971210661 | -0.04555 | 0.021469 | -3.87023 |
| oxidoreductase short-chain d( | 0.998079 | 1.106773109 | 0.149133 | 0.021551 | 3.725028 |
| L-aspartate oxidase | 1.005136 | 0.95674285 | -0.07119 | 0.021684 | -5.91714 |
| isocitrate lyase | 1.049813 | 0.942643523 | -0.15535 | 0.021747 | -3.99643 |
| transcriptional regulatory pro | 1.163567 | 0.858409309 | -0.43882 | 0.021827 | -4.35843 |
| OsmC family protein...1516 | 1.047345 | 0.95790412 | -0.12878 | 0.021869 | -3.88558 |
| cysteine desulfurase IscS | 1.200594 | 0.931440411 | -0.36621 | 0.022054 | -3.73678 |
| acyl-CoA dehydrogenase...124 | 1.187247 | 0.863602068 | -0.45918 | 0.022398 | -5.84044 |
| tRNA-dihydrouridine synthase | 1.183536 | 0.885212441 | -0.41901 | 0.02242 | -4.98997 |
| aconitate hydratase 1 | 1.091914 | 0.956143002 | -0.19156 | 0.022533 | -4.53842 |
| queuine tRNA-ribosyltransfera | 1.032864 | 0.955296181 | -0.11263 | 0.022579 | -5.25185 |
| transcriptional regulator NrdR | 1.041362 | 0.963803182 | -0.11166 | 0.022642 | -4.31057 |
| putative two-component resp | 1.144695 | 0.981102267 | -0.22249 | 0.022679 | -5.60057 |
| lipid A biosynthesis acyltransf | 0.860976 | 0.971555952 | 0.174324 | 0.022862 | 4.351361 |
| enoyl-CoA hydratase/isomera | 1.021154 | 0.923474671 | -0.14506 | 0.022865 | -3.96277 |
| adenylosuccinate lyase | 0.914237 | 0.955187073 | 0.063215 | 0.022876 | 5.047028 |
| competence/damage-inducib | 1.029205 | 0.913607608 | -0.17188 | 0.022901 | -3.59301 |
| integration host factor, beta s | 1.121744 | 0.9568902 | -0.22932 | 0.023008 | -4.14325 |

| | | | | | |
|---|---|---|---|---|---|
| 16S rRNA processing protein I | 1.046963 | 0.910362393 | -0.2017 | 0.02305 | -3.59846 |
| hypothetical protein ABUW_1 | 1.057714 | 0.946242402 | -0.16067 | 0.023556 | -4.03037 |
| phosphate acetyltransferase | 1.165716 | 0.840765657 | -0.47144 | 0.023589 | -6.24878 |
| hypothetical protein ABUW_0 | 1.045752 | 0.864894489 | -0.27395 | 0.023678 | -5.55778 |
| benzoate 1,2-dioxygenase, lar | 0.85191 | 1.118760103 | 0.393128 | 0.024033 | 4.574158 |
| hypothetical protein ABUW_0 | 1.079899 | 0.827888133 | -0.38339 | 0.024149 | -3.7388 |
| hypothetical protein ABUW_2 | 1.053191 | 0.975268131 | -0.1109 | 0.024163 | -4.03826 |
| biopolymer transport protein | 1.162579 | 0.989887173 | -0.23199 | 0.024328 | -3.54222 |
| hypothetical protein ABUW_1 | 0.472962 | 1.747905475 | 1.885831 | 0.02443 | 4.689864 |
| general secretion pathway pr | 1.049995 | 0.965377248 | -0.12122 | 0.024437 | -3.53259 |
| GTP cyclohydrolase II | 0.994005 | 1.031979194 | 0.054089 | 0.024448 | 3.54429 |
| oxidoreductase short-chain de | 0.981902 | 1.230290128 | 0.325348 | 0.024499 | 4.283432 |
| 6-pyruvoyl tetrahydrobiopter | 0.987123 | 1.107344514 | 0.165803 | 0.024555 | 4.817999 |
| adenylate cyclase | 1.022068 | 0.938811761 | -0.12258 | 0.024735 | -4.4289 |
| hypothetical protein ABUW_1 | 1.148606 | 0.962442146 | -0.25511 | 0.024869 | -3.50661 |
| protein-export membrane pro | 0.962737 | 1.053853952 | 0.130461 | 0.024903 | 4.628034 |
| ribulose-phosphate 3-epimera | 0.969583 | 1.057470938 | 0.125181 | 0.024962 | 3.562832 |
| Spore coat polysaccharide bio | 1.014099 | 1.085679602 | 0.0984 | 0.025214 | 3.71341 |
| hypothetical protein ABUW_1 | 0.960644 | 1.250205194 | 0.380092 | 0.025225 | 4.596669 |
| sodium-and chloride-depende | 1.035709 | 1.210850795 | 0.225402 | 0.025453 | 4.542525 |
| thiamine pyrophosphate enzy | 1.021176 | 0.882436186 | -0.21067 | 0.025474 | -3.77291 |
| acyl-(acyl-carrier-protein)--UD | 0.989778 | 0.921551746 | -0.10304 | 0.025574 | -4.92024 |
| beta-lactamase OXA-23 | 1.029189 | 0.873340867 | -0.23689 | 0.025595 | -6.01182 |
| glyceraldehyde-3-phosphate c | 0.924179 | 1.069159376 | 0.210232 | 0.025627 | 5.714162 |
| glucose sorbosone dehydroge | 1.033567 | 0.946070628 | -0.12761 | 0.02563 | -4.51115 |
| hypothetical protein ABUW_0 | 1.055763 | 0.958016671 | -0.14016 | 0.02566 | -3.59702 |
| transcriptional regulator AcrR | 1.053792 | 0.996935641 | -0.08002 | 0.025666 | -3.66696 |
| cytidylate kinase | 1.071963 | 0.965002773 | -0.15165 | 0.025759 | -4.29482 |
| hypothetical protein ABUW_0 | 1.028427 | 0.933610478 | -0.13955 | 0.025926 | -4.12522 |
| permease...646 | 0.946128 | 1.071042097 | 0.178908 | 0.025981 | 3.782867 |
| fumarylacetoacetase | 0.893201 | 1.019103367 | 0.190243 | 0.025997 | 4.207581 |
| hypothetical protein ABUW_1 | 0.881929 | 1.083942178 | 0.297554 | 0.026152 | 4.821378 |
| ribonuclease III | 0.987637 | 1.091445068 | 0.144187 | 0.026178 | 4.576784 |
| hypothetical protein ABUW_2 | 1.107037 | 0.846624905 | -0.38691 | 0.026404 | -4.08868 |
| ABC transporter, ATP-binding | 1.141803 | 1.240372706 | 0.11946 | 0.026619 | 3.512487 |
| copper resistance protein A | 1.076109 | 1.019286813 | -0.07826 | 0.026645 | -3.47914 |
| NADPH-dependent fmn reduc | 0.966086 | 1.158210059 | 0.261673 | 0.026658 | 4.369546 |
| phosphoserine phosphatase.. | 0.932826 | 1.004355486 | 0.10659 | 0.026745 | 4.200519 |
| hypothetical protein ABUW_2 | 0.864857 | 1.108896524 | 0.358592 | 0.026763 | 3.685529 |
| methyltransferase type 11 | 0.967339 | 1.047739009 | 0.115186 | 0.02689 | 5.230883 |
| hypothetical protein ABUW_2 | 0.952975 | 1.174045465 | 0.300979 | 0.027125 | 3.995153 |
| lipoprotein, putative...1678 | 0.995394 | 1.137314241 | 0.192292 | 0.027211 | 3.557688 |
| chorismate mutase...711 | 1.130213 | 0.895089187 | -0.33649 | 0.027461 | -3.48097 |
| acyl-CoA dehydrogenase...519 | 0.934706 | 1.198971879 | 0.359213 | 0.027539 | 5.786729 |
| hypothetical protein ABUW_2 | 0.999501 | 0.90216745 | -0.14781 | 0.027604 | -4.22801 |
| 2-isopropylmalate synthase | 0.92617 | 0.996733411 | 0.10593 | 0.027891 | 4.15556 |
| hypothetical protein ABUW_3 | 1.023117 | 1.091515618 | 0.093362 | 0.028012 | 4.30621 |

| | | | | | |
|---|---|---|---|---|---|
| isoleucyl-tRNA synthetase | 0.957624 | 0.987762558 | 0.044705 | 0.028043 | 3.894535 |
| biofilm-associated protein...2: | 0.946417 | 1.233898654 | 0.382676 | 0.028256 | 5.63403 |
| thioredoxin-disulfide reductas | 0.984551 | 0.911237425 | -0.11164 | 0.028331 | -3.5673 |
| acetyl-CoA carboxylase, biotir | 1.157332 | 0.814201114 | -0.50735 | 0.028383 | -5.3384 |
| hypothetical protein ABUW_2 | 0.865065 | 1.053928078 | 0.284896 | 0.028521 | 3.556681 |
| general secretion pathway pr | 0.918176 | 1.015434113 | 0.145254 | 0.028574 | 3.373624 |
| secretion protein HlyD...2182 | 0.981533 | 1.047185605 | 0.093409 | 0.028673 | 4.710017 |
| short-chain dehydrogenase/re | 1.058313 | 0.946773794 | -0.16067 | 0.02891 | -4.27392 |
| pirin family protein | 1.097847 | 0.956915153 | -0.19821 | 0.028921 | -5.01242 |
| DNA-directed RNA polymeras | 1.027656 | 0.958189661 | -0.10097 | 0.028924 | -5.30512 |
| phosphoribosylglycinamide fc | 0.997878 | 0.932151689 | -0.0983 | 0.028994 | -3.96777 |
| oxidoreductase...1707 | 1.044809 | 0.908642727 | -0.20145 | 0.029001 | -4.69627 |
| fumarate lyase | 1.147777 | 0.945080846 | -0.28033 | 0.029041 | -4.16629 |
| ribosomal protein S8 | 1.022446 | 0.926760457 | -0.14176 | 0.029176 | -3.32755 |
| hypothetical protein ABUW_2 | 1.042474 | 1.140008641 | 0.129033 | 0.029278 | 3.444927 |
| endoribonuclease L-PSP famil | 1.112039 | 0.711202801 | -0.64487 | 0.029524 | -5.3469 |
| tryptophan synthase, beta sul | 1.150961 | 1.025434153 | -0.1666 | 0.029618 | -5.55259 |
| dioxygenase alpha subunit | 1.093321 | 0.91671829 | -0.25417 | 0.029789 | -3.55439 |
| peptidoglycan glycosyltransfe | 0.95768 | 1.028798619 | 0.103346 | 0.029978 | 3.841709 |
| hypothetical protein ABUW_3 | 0.901837 | 1.050534285 | 0.220185 | 0.030024 | 3.495402 |
| hypothetical protein ABUW_0 | 1.11394 | 0.840576058 | -0.40622 | 0.030034 | -5.36199 |
| alpha/beta hydrolase...54 | 1.040503 | 0.784325921 | -0.40776 | 0.030322 | -3.28794 |
| hypothetical protein ABUW_1 | 1.029107 | 0.982070185 | -0.0675 | 0.030341 | -3.41748 |
| ribosomal protein L7/L12 | 1.052941 | 0.91953475 | -0.19545 | 0.030548 | -4.54707 |
| 3-hydroxyacyl-CoA dehydroge | 1.11868 | 0.981574494 | -0.18863 | 0.030552 | -5.17208 |
| hypothetical protein ABUW_0 | 1.020639 | 0.970628325 | -0.07248 | 0.030559 | -4.17782 |
| hypothetical protein ABUW_0 | 0.956726 | 1.16871319 | 0.288743 | 0.030653 | 4.047963 |
| L-carnitine dehydrogenase | 1.056554 | 0.887237448 | -0.25197 | 0.03068 | -4.28412 |
| chaperone protein DnaK | 1.051915 | 1.008117562 | -0.06135 | 0.030715 | -3.29818 |
| NAD(+) kinase | 0.910679 | 1.047054733 | 0.201322 | 0.030857 | 3.63528 |
| tryptophanyl-tRNA synthetase | 0.944914 | 0.960870991 | 0.02416 | 0.030939 | 3.491696 |
| dihydroorotase | 0.841807 | 1.001514381 | 0.250622 | 0.031316 | 4.576371 |
| phenylalanyl-tRNA synthetase | 0.915623 | 0.962943925 | 0.072697 | 0.031318 | 3.34996 |
| hypothetical protein ABUW_2 | 1.65802 | 0.711589978 | -1.22034 | 0.031426 | -4.37434 |
| aspartyl-tRNA synthetase | 0.996909 | 0.967293163 | -0.04351 | 0.031445 | -3.38003 |
| homoserine O-acetyltransfera | 0.988197 | 0.959790641 | -0.04208 | 0.031567 | -3.26605 |
| hypothetical protein ABUW_1 | 0.621823 | 1.044157311 | 0.747762 | 0.031644 | 4.191837 |
| two-component system DNA- | 0.932004 | 0.996518776 | 0.096561 | 0.031732 | 5.090837 |
| glyoxalase | 0.996843 | 1.10213926 | 0.144869 | 0.031832 | 4.112039 |
| hypothetical protein ABUW_1 | 1.12378 | 0.940548942 | -0.25678 | 0.031888 | -5.36728 |
| amidase | 0.949664 | 1.087819865 | 0.19595 | 0.031907 | 3.327719 |
| hypothetical protein ABUW_2 | 1.17674 | 0.969099615 | -0.28008 | 0.032198 | -5.09567 |
| 4-hydroxy-3-methylbut-2-en-: | 0.931183 | 1.028344629 | 0.143187 | 0.032299 | 3.22301 |
| copper/zinc superoxide dismu | 1.147285 | 0.96034527 | -0.2566 | 0.032568 | -4.2772 |
| hypothetical protein ABUW_0 | 0.852168 | 0.979253445 | 0.200545 | 0.032635 | 3.402602 |
| transcriptional regulator, LysR | 0.977942 | 1.106058915 | 0.177607 | 0.032734 | 5.077255 |
| esterase...1086 | 0.918502 | 1.087825079 | 0.244091 | 0.032754 | 4.04064 |

| | | | | | |
|---|---|---|---|---|---|
| GTP-binding protein Era | 0.983607 | 1.058842052 | 0.106334 | 0.032763 | 4.497965 |
| 3-oxoacid CoA-transferase, su | 0.989196 | 0.839887218 | -0.23606 | 0.032919 | -4.48677 |
| exodeoxyribonuclease VII | 0.94532 | 1.075089843 | 0.185583 | 0.032964 | 3.843075 |
| thioredoxin-disulfide reductas | 1.080916 | 0.9307559 | -0.21578 | 0.033023 | -3.40688 |
| ribosomal protein L14 | 1.054056 | 0.933689822 | -0.17494 | 0.033073 | -3.52177 |
| aspartate-semialdehyde dehy | 1.04359 | 0.99849594 | -0.06373 | 0.033423 | -4.18323 |
| paraquat-inducible protein B | 0.971949 | 1.060302376 | 0.125523 | 0.033483 | 3.502087 |
| primosomal protein N` | 0.90902 | 1.127873444 | 0.311222 | 0.033702 | 4.439552 |
| nucleoprotein/polynucleotide | 0.86729 | 0.9766413 | 0.171314 | 0.033733 | 3.623341 |
| TonB-dependent receptor...2 | 0.905126 | 0.877043855 | -0.04547 | 0.033817 | -3.22862 |
| hypothetical protein ABUW_3 | 1.16466 | 0.795026121 | -0.55084 | 0.033893 | -3.24759 |
| UDP-3-O-(3-hydroxymyristoyl | 0.918431 | 0.995960653 | 0.116917 | 0.033959 | 3.168351 |
| NADPH dehydrogenase | 1.034343 | 0.948795391 | -0.12455 | 0.033985 | -4.52659 |
| major facilitator family transp | 0.843427 | 1.036198788 | 0.296966 | 0.034012 | 3.775575 |
| lysyl-tRNA synthetase...1441 | 0.971087 | 1.041728907 | 0.101308 | 0.03402 | 3.197316 |
| MscS Mechanosensitive ion c | 0.911772 | 1.057795307 | 0.214315 | 0.034081 | 4.741046 |
| hypothetical protein ABUW_0 | 0.978352 | 1.110655211 | 0.182985 | 0.034103 | 5.226529 |
| hypothetical protein ABUW_1 | 1.093481 | 0.900177111 | -0.28065 | 0.034509 | -4.95252 |
| hypothetical protein ABUW_5 | 1.143786 | 0.984420873 | -0.21647 | 0.034629 | -3.74681 |
| ribosomal protein L23 | 1.090241 | 0.941888436 | -0.21102 | 0.034764 | -3.7997 |
| site-specific recombinase | 0.926646 | 1.081119705 | 0.222436 | 0.035266 | 4.310789 |
| hypothetical protein ABUW_1 | 1.092307 | 1.038603909 | -0.07273 | 0.035336 | -4.22145 |
| 2-methylisocitrate dehydratas | 0.887862 | 1.028851849 | 0.212628 | 0.035336 | 5.108811 |
| CsuA | 0.912856 | 0.950311515 | 0.058013 | 0.035361 | 3.351556 |
| phosphoribosyl (-ATP, -AMP) | 0.868371 | 1.034698128 | 0.252826 | 0.03543 | 4.671163 |
| copper-translocating P-type A | 1.036001 | 0.991478723 | -0.06337 | 0.03544 | -3.13527 |
| 3-deoxy-7-phosphoheptulona | 0.942651 | 0.751525953 | -0.3269 | 0.035461 | -3.53845 |
| glutamyl-tRNA(Gln) amidotrar | 0.919838 | 0.996159034 | 0.114997 | 0.035464 | 3.277977 |
| phospholipase D | 0.943003 | 1.224648685 | 0.377034 | 0.035628 | 3.188286 |
| RNA methyltransferase, TrmH | 1.072012 | 1.153294489 | 0.10544 | 0.035799 | 3.770263 |
| outer membrane lipoprotein | 1.046658 | 0.944647507 | -0.14794 | 0.035806 | -4.04271 |
| hypothetical protein ABUW_3 | 1.076299 | 0.928289451 | -0.21343 | 0.03611 | -3.45893 |
| type VI secretion ATPase, ClpV | 0.971633 | 1.066524468 | 0.134434 | 0.036298 | 3.343241 |
| hypothetical protein ABUW_3 | 1.03667 | 0.994668939 | -0.05967 | 0.036604 | -3.67797 |
| NADH dehydrogenase | 1.067767 | 1.18614696 | 0.151686 | 0.036626 | 3.342866 |
| transcriptional regulator...957 | 1.077555 | 0.966622608 | -0.15674 | 0.036628 | -3.09224 |
| hemerythrin | 0.933171 | 1.04381378 | 0.16165 | 0.036747 | 3.715721 |
| transcriptional regulator, Gntl | 1.016457 | 0.872963934 | -0.21956 | 0.036844 | -3.19041 |
| hypothetical protein ABUW_1 | 0.9083 | 1.048349561 | 0.206879 | 0.036879 | 3.528046 |
| dimethylmenaquinone methy | 0.803864 | 0.934485402 | 0.217221 | 0.036912 | 4.146124 |
| hypothetical protein ABUW_3 | 1.091958 | 1.182296601 | 0.114674 | 0.037224 | 4.531355 |
| 3-oxoacyl-(acyl carrier protein | 0.939054 | 1.052416781 | 0.164426 | 0.037224 | 3.098266 |
| hypothetical protein ABUW_2 | 1.408924 | 0.768231635 | -0.87498 | 0.037488 | -4.29333 |
| ABC-1 domain protein...1828 | 0.960864 | 1.059714379 | 0.141272 | 0.03758 | 3.439863 |
| glutathione S-transferase...85 | 0.974902 | 1.038060931 | 0.090561 | 0.037582 | 3.733913 |
| superoxide dismutase (Fe) | 1.029609 | 0.932167954 | -0.14343 | 0.037869 | -3.70719 |
| putative outer membrane pro | 0.895706 | 1.061438963 | 0.244925 | 0.037871 | 3.196786 |

| | | | | | |
|---|---|---|---|---|---|
| histidine triad protein...1359 | 1.020412 | 0.959050417 | -0.08947 | 0.037987 | -3.53489 |
| lipoprotein, putative...1053 | 1.227401 | 0.95639355 | -0.35993 | 0.038006 | -4.30932 |
| glutathione S-transferase fam | 0.999639 | 1.076069212 | 0.106291 | 0.038182 | 3.061655 |
| adenosine deaminase...1836 | 1.128149 | 0.943247296 | -0.25825 | 0.038298 | -3.75538 |
| hypothetical protein ABUW_1 | 0.976179 | 1.339795223 | 0.456795 | 0.038384 | 3.040515 |
| hypothetical protein ABUW_3 | 1.142315 | 0.634850127 | -0.84747 | 0.038397 | -3.82929 |
| hypothetical protein ABUW_2 | 1.585364 | 0.825449602 | -0.94156 | 0.038403 | -4.80365 |
| acyl-CoA dehydrogenase...892 | 1.170587 | 1.009803061 | -0.21316 | 0.03862 | -3.07931 |
| acetyl-coenzyme A synthetase | 1.045478 | 0.965447476 | -0.11489 | 0.038655 | -3.0351 |
| ribosomal protein L27 | 1.113706 | 0.872665411 | -0.35187 | 0.03889 | -4.0049 |
| DNA translocase FtsK | 0.933675 | 1.111088974 | 0.250981 | 0.039211 | 4.860081 |
| hypothetical protein ABUW_3 | 0.90194 | 1.046199165 | 0.214055 | 0.039362 | 3.092247 |
| muconate cycloisomerase | 0.839061 | 0.988366979 | 0.236271 | 0.039548 | 4.173813 |
| phage small terminase subuni | 0.829788 | 0.986480435 | 0.249548 | 0.039553 | 4.471061 |
| aconitate hydratase 2 | 0.958484 | 0.913067908 | -0.07003 | 0.039554 | -3.01626 |
| crispr-associated protein Cas1 | 1.076821 | 0.842357089 | -0.35427 | 0.039708 | -3.00695 |
| ATP-dependent Clp protease, | 1.034848 | 0.930681369 | -0.15306 | 0.039737 | -3.56704 |
| ribosomal protein S3 | 1.047746 | 0.952584623 | -0.13737 | 0.039901 | -3.57295 |
| ribosomal protein L33 | 1.062882 | 0.96383249 | -0.14113 | 0.03996 | -3.55551 |
| hypothetical protein ABUW_0 | 1.021105 | 0.9118744 | -0.16322 | 0.040034 | -3.69215 |
| ribosomal protein L10 | 1.05832 | 0.94468868 | -0.16387 | 0.040125 | -4.26746 |
| hypothetical protein ABUW_0 | 1.130192 | 1.017003423 | -0.15224 | 0.04015 | -3.87293 |
| lipid-A-disaccharide synthase | 0.813652 | 1.033257627 | 0.344716 | 0.040382 | 4.184741 |
| transglutaminase | 0.933592 | 1.222636895 | 0.389133 | 0.040439 | 3.236084 |
| tRNA/rRNA methyltransferase | 1.128068 | 0.834270041 | -0.43527 | 0.04054 | -4.80873 |
| hypothetical protein ABUW_1 | 1.076927 | 0.982333169 | -0.13264 | 0.040885 | -4.09222 |
| poly(R)-hydroxyalkanoic acid s | 1.135248 | 0.951079871 | -0.25537 | 0.0409 | -4.19055 |
| sulfate adenylyltransferase su | 1.01348 | 0.969448115 | -0.06408 | 0.040926 | -3.37784 |
| hypothetical protein ABUW_3 | 0.896334 | 1.203621536 | 0.425273 | 0.041022 | 2.992041 |
| aromatic-amino-acid aminotra | 0.854222 | 0.936544178 | 0.132736 | 0.041248 | 2.972122 |
| rod shape-determining protei | 1.01205 | 0.957019801 | -0.08066 | 0.041295 | -2.99381 |
| glycerol kinase | 1.033485 | 0.920787866 | -0.16658 | 0.041434 | -4.48227 |
| transcriptional regulator, Mar | 1.016293 | 0.924799876 | -0.1361 | 0.041467 | -3.31328 |
| cytochrome B561...551 | 1.086154 | 1.199276523 | 0.142935 | 0.041505 | 3.251584 |
| phosphoglycolate phosphatas | 0.963986 | 0.842626503 | -0.19412 | 0.041526 | -3.3104 |
| transcription antitermination | 1.013596 | 0.89436975 | -0.18054 | 0.041647 | -4.38969 |
| ribosomal protein L18 | 1.045129 | 0.944524829 | -0.14602 | 0.0419 | -3.35425 |
| metallo-beta-lactamase super | 0.892903 | 1.110145367 | 0.314173 | 0.041923 | 4.127153 |
| single-strand binding protein | 0.982631 | 0.901227939 | -0.12476 | 0.041968 | -3.10955 |
| hypothetical protein ABUW_2 | 1.00953 | 1.058636393 | 0.068524 | 0.042158 | 2.978631 |
| isochorismatase hydrolase...4 | 0.849974 | 0.94957736 | 0.159867 | 0.042428 | 3.222884 |
| putative transferase | 1.240404 | 1.034708436 | -0.26159 | 0.042526 | -3.39962 |
| glycerol-3-phosphate acyltran | 0.937391 | 0.999364488 | 0.09236 | 0.04257 | 3.718606 |
| tryptophan synthase, alpha su | 1.049403 | 0.932239113 | -0.1708 | 0.042777 | -4.30115 |
| hypothetical protein ABUW_2 | 1.119204 | 1.174850529 | 0.070004 | 0.042969 | 2.971045 |
| hypothetical protein ABUW_3 | 1.028328 | 0.885421104 | -0.21587 | 0.042991 | -4.62509 |
| hypothetical protein ABUW_1 | 1.054933 | 1.022768611 | -0.04467 | 0.04304 | -3.32957 |

| | | | | | |
|---|---|---|---|---|---|
| hypothetical protein ABUW_2 | 0.812155 | 1.165088662 | 0.520613 | 0.043098 | 3.952472 |
| radical SAM enzyme, Cfr fami | 0.905048 | 1.051290919 | 0.216096 | 0.04324 | 3.1298 |
| glutathione S-transferase...16 | 1.118223 | 1.040745153 | -0.10359 | 0.043375 | -2.99724 |
| Rhs element Vgr protein, puta | 0.968425 | 1.075784807 | 0.151677 | 0.043444 | 3.237438 |
| pantoate--beta-alanine ligase | 0.943261 | 0.984515614 | 0.061757 | 0.043477 | 3.024261 |
| DNA polymerase III subunit ta | 1.01067 | 0.962496987 | -0.07046 | 0.043554 | -3.79376 |
| hypothetical protein ABUW_2 | 0.793135 | 1.103098463 | 0.475922 | 0.043634 | 4.060693 |
| hypothetical protein ABUW_1 | 0.803873 | 1.005715463 | 0.323182 | 0.043919 | 4.579241 |
| glutathione S-transferase...16 | 0.942709 | 0.986014045 | 0.064796 | 0.04394 | 3.279442 |
| O-methyl transferase | 1.012437 | 0.955229819 | -0.08391 | 0.043947 | -3.80704 |
| ribosomal protein L30 | 1.047051 | 0.964996765 | -0.11774 | 0.044063 | -3.12262 |
| NADH dehydrogenase I chain | 0.899889 | 1.013608414 | 0.171682 | 0.044335 | 3.966238 |
| hypothetical protein ABUW_1 | 0.846395 | 0.965921806 | 0.190575 | 0.044396 | 2.980872 |
| metallopeptidase, zinc binding | 0.974764 | 0.929993175 | -0.06783 | 0.044551 | -3.01008 |
| adenosylhomocysteinase | 1.105074 | 0.962202461 | -0.19973 | 0.044742 | -4.34802 |
| hypothetical protein ABUW_3 | 1.246119 | 1.359126401 | 0.125238 | 0.044904 | 2.945628 |
| ribosomal protein L1 | 1.057948 | 0.934789922 | -0.17855 | 0.045144 | -3.90561 |
| senescence marker protein-30 | 1.069104 | 0.942285261 | -0.18217 | 0.04518 | -3.75961 |
| malate dehydrogenase...2484 | 0.891935 | 0.947102929 | 0.086583 | 0.045263 | 4.36326 |
| integral membrane protein Te | 1.138597 | 0.93826762 | -0.27919 | 0.04542 | -2.93882 |
| cyclic nucleotide-binding dom | 0.989726 | 0.937977596 | -0.07748 | 0.045518 | -3.00331 |
| acyl-CoA dehydrogenase...241 | 1.075413 | 0.899325482 | -0.25798 | 0.046061 | -4.25816 |
| multidrug efflux protein AdeJ | 0.963174 | 1.019447092 | 0.081919 | 0.046214 | 3.985854 |
| transcriptional regulator, Fis-t | 0.872866 | 0.94554403 | 0.115385 | 0.046269 | 2.903881 |
| transcriptional regulator, TetF | 1.135837 | 0.89016085 | -0.35162 | 0.046323 | -4.29749 |
| nucleoside diphosphate kinas | 1.036175 | 0.911255162 | -0.18534 | 0.046329 | -3.012 |
| Peptidase M20D, amidohydro | 1.034922 | 0.742986998 | -0.47811 | 0.046367 | -2.92035 |
| GGDEF domain protein | 0.982866 | 0.87477968 | -0.16807 | 0.046546 | -3.74154 |
| beta-lactamase...5 | 1.179046 | 1.037360523 | -0.1847 | 0.04658 | -3.14996 |
| 2,3-dihydroxybenzoate-AMP l | 1.093796 | 0.869876067 | -0.33046 | 0.046627 | -3.05413 |
| monooxygenase, flavin-bindir | 0.791529 | 1.034355108 | 0.386017 | 0.046658 | 3.155529 |
| exonuclease V beta chain | 1.085741 | 0.938335051 | -0.21051 | 0.046755 | -2.84375 |
| hypothetical protein ABUW_2 | 0.919819 | 1.107069059 | 0.267323 | 0.046899 | 2.920936 |
| aminoglycoside phosphotrans | 0.952918 | 1.04820318 | 0.137494 | 0.047035 | 3.343994 |
| deoxyuridine 5'-triphosphate | 0.95323 | 0.885339716 | -0.10659 | 0.047193 | -3.94972 |
| carbon starvation protein A | 0.978502 | 1.062028429 | 0.118176 | 0.047245 | 3.021657 |
| phosphoenolpyruvate syntha: | 1.021272 | 0.919013025 | -0.15221 | 0.047303 | -3.84709 |
| peptidyl-prolyl cis-trans isome | 1.052937 | 1.001876212 | -0.07172 | 0.047391 | -2.9344 |
| ribosomal protein S10 | 1.051028 | 0.977911158 | -0.10403 | 0.047402 | -2.83116 |
| phenylacetic acid degradation | 1.090415 | 0.938884025 | -0.21586 | 0.04763 | -3.19239 |
| hypothetical protein ABUW_2 | 0.894781 | 1.090676936 | 0.285617 | 0.047639 | 3.354208 |
| ubiquinone/menaquinone bic | 0.996604 | 0.906057933 | -0.13742 | 0.047834 | -2.84469 |
| succinyl-CoA synthase, beta su | 0.946556 | 0.869267496 | -0.12289 | 0.047844 | -3.59945 |
| phosphoenolpyruvate carbox | 0.941796 | 0.887295658 | -0.086 | 0.047987 | -3.77628 |
| methyltransferase type 12 | 0.877539 | 1.039555228 | 0.244432 | 0.048009 | 4.287954 |
| protein secretion ABC efflux s | 0.978095 | 1.108632527 | 0.180734 | 0.048379 | 3.209112 |
| molybdopterin biosynthesis p | 0.918452 | 0.992046387 | 0.111203 | 0.048528 | 2.809674 |

| | | | | | |
|---|---|---|---|---|---|
| transcriptional regulator, TetF | 0.94525 | 0.924202832 | -0.03249 | 0.048612 | -2.98675 |
| transcriptional regulator, GntI | 0.925367 | 1.025017894 | 0.147552 | 0.048699 | 2.998041 |
| short chain dehydrogenase...2 | 1.105596 | 0.954449288 | -0.21208 | 0.048714 | -3.54257 |
| Ser/Thr protein phosphatase | 0.857302 | 1.028097431 | 0.262101 | 0.048752 | 3.881976 |
| lipoprotein | 0.977585 | 0.900561029 | -0.1184 | 0.048945 | -4.11224 |
| DNA-3-methyladenine glycosy | 0.980371 | 1.074858719 | 0.132747 | 0.04923 | 2.859029 |
| transcriptional regulator, Mar | 0.860946 | 0.948818932 | 0.140211 | 0.049305 | 3.075376 |
| type I secretion outer membr | 1.10756 | 1.042269549 | -0.08766 | 0.049349 | -2.97216 |
| short chain dehydrogenase...1 | 1.085983 | 1.009667972 | -0.10512 | 0.049512 | -3.02491 |
| Mating pair stabilization TraN | 1.169797 | 0.963240888 | -0.28029 | 0.049744 | -3.5546 |
| ribonuclease R | 0.936018 | 0.996050527 | 0.089682 | 0.050033 | 3.672064 |
| glycosyltransferase...487 | 1.011758 | 1.136339522 | 0.16753 | 0.050073 | 3.819153 |
| chaperone protein DnaJ | 0.938542 | 1.005651368 | 0.099637 | 0.050104 | 2.952104 |
| transcriptional regulator, LysF | 0.872815 | 1.138132417 | 0.382921 | 0.050153 | 3.480273 |
| hypothetical protein ABUW_0 | 0.991089 | 0.942239215 | -0.07292 | 0.050207 | -3.67778 |
| twitching motility protein | 1.214004 | 1.008749462 | -0.26721 | 0.050257 | -3.60271 |
| nitroreductase...1095 | 1.09671 | 1.043064942 | -0.07235 | 0.050265 | -3.14483 |
| hypothetical protein ABUW_1 | 0.845315 | 1.146759976 | 0.440003 | 0.050421 | 2.933326 |
| aminodeoxychorismate lyase. | 0.899374 | 0.991672268 | 0.140942 | 0.050486 | 3.088508 |
| transcriptional regulator, Pad | 1.070207 | 0.982023023 | -0.12406 | 0.050604 | -3.43333 |
| ribosomal protein S7 | 1.012824 | 0.948756058 | -0.09427 | 0.050688 | -3.13932 |
| 6-O-methylguanine-DNA metl | 1.277543 | 0.971651611 | -0.39486 | 0.050692 | -4.19725 |
| ribosomal RNA small subunit | 0.908566 | 0.961241638 | 0.081308 | 0.050792 | 3.263813 |
| alginate biosynthesis regulato | 1.077435 | 0.95523624 | -0.17367 | 0.051002 | -2.88481 |
| ribosomal protein L4/L1e | 1.095656 | 0.927848197 | -0.23983 | 0.051243 | -3.87068 |
| glutamate/aspartate ABC trar | 1.181256 | 0.956457461 | -0.30455 | 0.051318 | -3.60489 |
| DNA-binding protein HU | 1.054303 | 0.878916658 | -0.26249 | 0.051472 | -4.22104 |
| riboflavin biosynthesis proteir | 0.959837 | 1.083072292 | 0.174268 | 0.051608 | 3.844492 |
| cation efflux system protein (I | 0.928541 | 1.023391301 | 0.14032 | 0.051874 | 2.940094 |
| carboxy- protease | 1.020377 | 1.059706127 | 0.054562 | 0.051948 | 2.901617 |
| succinate dehydrogenase, hyd | 0.780919 | 1.05825576 | 0.438444 | 0.051958 | 3.876586 |
| helix-turn-helix- domain conta | 1.054918 | 0.934900102 | -0.17425 | 0.051977 | -3.56927 |
| tail tape measure protein...14 | 1.138547 | 1.03808765 | -0.13327 | 0.052328 | -2.8646 |
| phosphocarrier protein HPr | 1.027808 | 0.966440745 | -0.08882 | 0.052534 | -2.86042 |
| rhodanese domain protein...1 | 1.152369 | 0.868341461 | -0.40827 | 0.052786 | -2.97325 |
| fmn-dependent NADH-azored | 1.22617 | 0.71780155 | -0.7725 | 0.052841 | -3.79965 |
| hypothetical protein ABUW_2 | 0.931077 | 0.996958999 | 0.098633 | 0.053083 | 2.719591 |
| ribosomal protein L6 | 1.101186 | 0.923535722 | -0.25382 | 0.053195 | -4.02903 |
| UDP-N-acetylglucosamine dip | 0.882478 | 1.003094193 | 0.184825 | 0.053454 | 3.991293 |
| small GTP-binding protein | 0.970729 | 1.029619429 | 0.084971 | 0.053472 | 3.222044 |
| Zeta toxin family protein (plas | 1.053423 | 1.141236878 | 0.115514 | 0.053582 | 3.492836 |
| CsuA/B | 0.82877 | 0.904403259 | 0.125995 | 0.053612 | 3.517534 |
| hypothetical protein ABUW_1 | 0.992812 | 1.03436152 | 0.059148 | 0.053612 | 3.420878 |
| ribosomal protein L11 | 1.091607 | 0.91146531 | -0.26019 | 0.053699 | -3.88719 |
| NADH dehydrogenase I chain | 0.926112 | 1.033716315 | 0.158582 | 0.053818 | 3.274954 |
| N-ethylmaleimide reductase | 1.167987 | 0.866051838 | -0.4315 | 0.053853 | -3.312 |
| HAD-superfamily subfamily IB | 0.953289 | 1.018647553 | 0.095669 | 0.053871 | 3.633713 |

| | | | | | |
|---|---|---|---|---|---|
| putative periplasmic carboxyl- | 0.92032 | 1.164667072 | 0.33971 | 0.054451 | 3.506953 |
| replication initiation inhibitor | 1.05255 | 0.883581764 | -0.25245 | 0.054579 | -2.70953 |
| nucleotidyl transferase family | 1.044531 | 0.967313093 | -0.1108 | 0.054584 | -3.02134 |
| lipoprotein, putative...1648 | 0.891105 | 0.959010589 | 0.105952 | 0.054588 | 2.774265 |
| ErfK/YbiS/YcfS/YnhG family pr | 0.931227 | 1.018497211 | 0.129237 | 0.054686 | 3.039671 |
| hypothetical protein ABUW_0 | 1.060245 | 0.883284931 | -0.26345 | 0.054766 | -3.52035 |
| polyphosphate kinase...928 | 0.967087 | 1.046191253 | 0.113429 | 0.05515 | 2.704983 |
| flavin binding monooxygenase | 0.927561 | 1.002399715 | 0.111944 | 0.05535 | 3.152119 |
| translation elongation factor T | 1.015601 | 0.936953382 | -0.11629 | 0.055465 | -3.67557 |
| hypothetical protein ABUW_0 | 1.051037 | 0.822350611 | -0.35399 | 0.055544 | -3.51486 |
| biotin-[acetyl-CoA-carboxylas | 0.957259 | 1.121975419 | 0.22906 | 0.055891 | 3.256452 |
| monofunctional biosynthetic | 0.908477 | 1.129809814 | 0.314559 | 0.056176 | 3.256463 |
| hydroxyacylglutathione hydro | 1.067822 | 0.920645567 | -0.21395 | 0.056245 | -2.89428 |
| transcriptional regulator, TetR | 1.088766 | 0.945673811 | -0.20328 | 0.056268 | -2.73843 |
| aspartate kinase | 0.921749 | 0.990959937 | 0.104453 | 0.056448 | 3.424784 |
| glutamate-5-semialdehyde de | 1.034425 | 0.972802839 | -0.08861 | 0.056588 | -2.65873 |
| aldehyde dehydrogenase...15 | 0.893126 | 1.061795371 | 0.24957 | 0.056836 | 2.851657 |
| aminodeoxychorismate lyase. | 1.139285 | 0.873935326 | -0.38253 | 0.057183 | -3.65698 |
| acyl-CoA ligase | 1.087437 | 0.811289965 | -0.42264 | 0.057367 | -2.64917 |
| hypothetical protein ABUW_1 | 1.06232 | 0.974828255 | -0.124 | 0.057603 | -2.76773 |
| UDP-N-acetylenolpyruvoylglu | 0.980919 | 0.93934528 | -0.06248 | 0.057648 | -2.92249 |
| peptidyl-prolyl cis-trans isome | 1.054756 | 0.924726028 | -0.18981 | 0.057786 | -3.16519 |
| cysteine synthase B | 0.976592 | 0.950732921 | -0.03872 | 0.058048 | -2.68607 |
| acyl-CoA dehydrogenase...184 | 0.949926 | 1.063588569 | 0.163054 | 0.058052 | 3.131542 |
| P-hydroxyphenylacetate hydr | 0.739702 | 0.826445208 | 0.159975 | 0.058056 | 3.154086 |
| ribosomal protein S21 | 1.192555 | 0.929187259 | -0.36001 | 0.058248 | -3.8586 |
| hypothetical protein ABUW_0 | 0.996944 | 1.063031791 | 0.0926 | 0.058316 | 2.754183 |
| peptide chain release factor 1 | 0.920971 | 0.873592007 | -0.0762 | 0.058382 | -3.03402 |
| tyrosine recombinase XerD | 1.025706 | 0.948850375 | -0.11236 | 0.058535 | -2.62459 |
| gamma-glutamyltransferase | 1.123894 | 0.98850467 | -0.18519 | 0.058678 | -3.81465 |
| fumarylacetoacetate hydrolas | 0.939873 | 0.923193197 | -0.02583 | 0.058813 | -3.15582 |
| ATP synthase F0, A subunit | 0.881702 | 0.971578354 | 0.14004 | 0.059255 | 2.968397 |
| hypothetical protein ABUW_3 | 0.900693 | 1.072107345 | 0.251341 | 0.059339 | 2.946106 |
| cytochrome D ubiquinol oxida | 1.014937 | 1.229353862 | 0.276511 | 0.059347 | 3.840289 |
| D-lactate dehydrogenase | 0.862607 | 0.98563346 | 0.192347 | 0.059491 | 2.808783 |
| 3-oxoacyl-[acyl-carrier-protein | 1.041905 | 0.885719437 | -0.2343 | 0.059535 | -3.52623 |
| lipoprotein, putative...469 | 0.803062 | 1.071869009 | 0.416545 | 0.059701 | 2.663494 |
| ribosomal protein L24 | 1.095683 | 0.905224906 | -0.27548 | 0.059911 | -3.29236 |
| hypothetical protein ABUW_3 | 1.018972 | 0.973559279 | -0.06577 | 0.05992 | -2.89333 |
| formyltetrahydrofolate defor | 1.024131 | 0.878503644 | -0.22128 | 0.059934 | -3.69668 |
| hypothetical protein ABUW_2 | 0.973612 | 1.095477577 | 0.170141 | 0.059956 | 3.086821 |
| peptidase S16, lon domain pro | 0.838223 | 0.935598976 | 0.158556 | 0.060052 | 2.744079 |
| hypothetical protein ABUW_0 | 0.879757 | 0.962675649 | 0.129945 | 0.060131 | 2.620242 |
| thioesterase superfamily prot | 0.937049 | 1.188136794 | 0.342504 | 0.060263 | 3.182546 |
| pyruvate dehydrogenase com | 1.041071 | 0.93554986 | -0.15418 | 0.060379 | -3.87987 |
| hypothetical protein ABUW_3 | 1.143126 | 0.787796423 | -0.53709 | 0.060454 | -3.8504 |
| transcriptional regulator, LysR | 1.062521 | 1.798713012 | 0.759474 | 0.0606 | 3.128937 |

| | | | | | |
|---|---|---|---|---|---|
| MotA/TolQ/ExbB proton char | 0.873977 | 0.986065341 | 0.174088 | 0.0607 | 2.876575 |
| histidinol dehydrogenase | 0.953404 | 1.009397434 | 0.082335 | 0.060757 | 2.754564 |
| hypothetical protein ABUW_( | 1.285855 | 0.839224641 | -0.6156 | 0.06078 | -3.7774 |
| GTP-binding protein...1376 | 0.977091 | 0.926112976 | -0.0773 | 0.061091 | -2.58529 |
| amidohydrolase...1413 | 1.044113 | 1.113388093 | 0.092679 | 0.061106 | 3.083316 |
| ferredoxin--NADP(+) reductas | 1.01571 | 0.939770193 | -0.11211 | 0.061581 | -3.75436 |
| DNA topoisomerase IV, A sub | 1.039738 | 0.99485822 | -0.06366 | 0.061648 | -2.62433 |
| short chain dehydrogenase...! | 1.04729 | 1.329780777 | 0.344527 | 0.061766 | 3.421375 |
| heme oxygenase-like protein. | 0.94403 | 1.116072741 | 0.241527 | 0.062012 | 3.633822 |
| hypothetical protein ABUW_1 | 0.802614 | 1.097950238 | 0.452035 | 0.062072 | 2.688717 |
| integrase | 0.811871 | 0.964077826 | 0.247899 | 0.062118 | 2.853265 |
| phosphoribosylaminoimidazo | 0.969386 | 0.936537805 | -0.04973 | 0.062231 | -2.83851 |
| ferrous iron transport protein | 0.880708 | 0.975963455 | 0.148163 | 0.062532 | 2.741866 |
| transcriptional regulator, LysF | 0.850983 | 1.339245179 | 0.654218 | 0.062717 | 3.207329 |
| hypothetical protein ABUW_( | 0.912143 | 1.036073862 | 0.183795 | 0.062855 | 3.305852 |
| cytochrome D ubiquinol oxida | 0.813384 | 1.134877175 | 0.480528 | 0.06289 | 2.606443 |
| AdeR | 0.951807 | 1.071983795 | 0.171541 | 0.063074 | 3.194535 |
| hypothetical protein ABUW_2 | 0.842502 | 1.211514282 | 0.524059 | 0.063496 | 2.644987 |
| putative transcriptional regul | 1.500494 | 1.06022558 | -0.50107 | 0.063729 | -2.60335 |
| hypothetical protein ABUW_1 | 1.017028 | 0.869860705 | -0.2255 | 0.063817 | -3.31385 |
| peptidase S1 and S6 | 0.920935 | 1.025714154 | 0.155458 | 0.063839 | 2.702005 |
| hypothetical protein ABUW_( | 1.060668 | 0.841253224 | -0.33436 | 0.064177 | -2.92759 |
| 3-hydroxyisobutyrate dehydr( | 1.058121 | 1.011766274 | -0.06463 | 0.064192 | -2.55095 |
| non-ribosomal peptide synthe | 1.154765 | 0.967454659 | -0.25533 | 0.064339 | -2.53676 |
| vanillate O-demethylase oxyg | 0.905912 | 1.187340438 | 0.390291 | 0.064425 | 3.407159 |
| hypothetical protein ABUW_3 | 0.906822 | 1.072632616 | 0.242265 | 0.064551 | 3.468684 |
| ABC transporter, ATP-binding | 0.985267 | 0.891329337 | -0.14456 | 0.064609 | -3.11814 |
| hypothetical protein ABUW_( | 1.118552 | 0.926556755 | -0.27168 | 0.064715 | -2.52936 |
| 5-dehydro-4-deoxyglucarate ( | 1.005129 | 0.923878992 | -0.1216 | 0.064786 | -2.6621 |
| tRNA-dihydrouridine synthase | 0.992495 | 1.223239893 | 0.301576 | 0.064872 | 2.681425 |
| FeS cluster assembly scaffold | 1.151406 | 0.942688637 | -0.28854 | 0.065089 | -2.89481 |
| hypothetical protein ABUW_1 | 0.95405 | 1.140499342 | 0.257529 | 0.065122 | 3.463992 |
| ribosomal protein L20 | 1.090418 | 0.842828403 | -0.37157 | 0.065125 | -3.41021 |
| betaine aldehyde dehydroger | 0.929527 | 0.976927477 | 0.071755 | 0.065208 | 2.522348 |
| hypothetical protein ABUW_( | 1.020614 | 1.062484741 | 0.058005 | 0.065351 | 2.58158 |
| amino acid adenylation | 0.844838 | 1.136968585 | 0.428445 | 0.065466 | 2.604691 |
| thiol:disulfide interchange pr( | 0.888623 | 1.049772452 | 0.240433 | 0.065466 | 2.6609 |
| cold-shock DNA-binding dom: | 0.863415 | 1.033806966 | 0.25984 | 0.06548 | 3.526624 |
| hypothetical protein ABUW_3 | 1.063129 | 1.00174815 | -0.0858 | 0.065515 | -3.37056 |
| acyl-CoA dehydrogenase...237 | 1.067868 | 0.876798107 | -0.28442 | 0.065737 | -3.43228 |
| ribosomal protein S11 | 1.044064 | 0.962779644 | -0.11693 | 0.065898 | -3.56572 |
| 3-hydroxyacyl-CoA dehydroge | 0.985542 | 0.937450511 | -0.07218 | 0.066158 | -2.60661 |
| alanine racemase...1478 | 0.90357 | 0.962654623 | 0.091383 | 0.066285 | 3.380708 |
| transcriptional regulator, LysF | 0.948182 | 1.021498264 | 0.107451 | 0.066303 | 3.562879 |
| choline dehydrogenase | 0.958869 | 1.001651096 | 0.062975 | 0.066335 | 3.420167 |
| hypothetical protein ABUW_1 | 0.969198 | 1.180899859 | 0.285023 | 0.066362 | 2.700288 |
| glycine hydroxymethyltransfe | 0.879128 | 0.937849273 | 0.093283 | 0.066431 | 2.671643 |

| | | | | | |
|---|---|---|---|---|---|
| lipoprotein, putative...997 | 0.896675 | 0.97146078 | 0.11557 | 0.066474 | 2.507701 |
| hypothetical protein ABUW_0 | 1.151263 | 0.927473758 | -0.31184 | 0.066616 | -3.4428 |
| lipolytic enzyme | 1.075474 | 0.892363497 | -0.26927 | 0.066652 | -3.26524 |
| TetR family transcriptional reg | 1.086367 | 1.014470698 | -0.09878 | 0.066658 | -3.50272 |
| hypothetical protein ABUW_1 | 0.840419 | 1.201634843 | 0.515818 | 0.066743 | 3.623157 |
| 5,10-methylenetetrahydrofola | 1.033588 | 0.953949189 | -0.11568 | 0.066921 | -2.52335 |
| hypothetical protein ABUW_1 | 0.973985 | 0.932471371 | -0.06284 | 0.066937 | -2.54927 |
| transcriptional regulator, TetR | 0.933412 | 1.090176397 | 0.223975 | 0.06715 | 3.056481 |
| ribosomal protein S4 | 1.006429 | 0.945390946 | -0.09026 | 0.067346 | -3.18588 |
| phosphoglucosamine mutase | 1.036209 | 0.876213761 | -0.24196 | 0.067369 | -3.46845 |
| O-acetylhomoserine/O-acetyl | 0.842268 | 1.049489191 | 0.317336 | 0.067389 | 3.538817 |
| catechol 1,2-dioxygenase | 1.031452 | 0.893162964 | -0.20768 | 0.06776 | -3.53759 |
| 4-carboxymuconolactone dec | 1.170995 | 0.942737303 | -0.31281 | 0.068219 | -3.05214 |
| glyoxalase/bleomycin resistar | 0.806635 | 1.00992561 | 0.324262 | 0.06837 | 2.62496 |
| putrescine importer | 1.089506 | 0.937426445 | -0.2169 | 0.068606 | -2.49717 |
| ribosomal protein S2 | 1.057078 | 0.947789106 | -0.15744 | 0.068883 | -3.08259 |
| phosphatidate cytidylyltransfe | 0.876822 | 0.951863868 | 0.118472 | 0.068887 | 3.085097 |
| glyoxalase I | 1.244735 | 0.875074353 | -0.50836 | 0.069056 | -3.48615 |
| ABC-type transporter, peripla | 0.999532 | 0.846264789 | -0.24014 | 0.069282 | -2.97427 |
| thiazole biosynthesis protein T | 1.038561 | 0.923601549 | -0.16924 | 0.069499 | -3.24579 |
| hypothetical protein ABUW_3 | 0.942125 | 1.011798096 | 0.102932 | 0.069659 | 3.398493 |
| transcriptional regulator, AsnC | 1.02703 | 0.94299835 | -0.12315 | 0.069772 | -2.88964 |
| thiamine S/molybdopterin co | 0.843042 | 1.057169285 | 0.326529 | 0.069944 | 2.854017 |
| Protein incC (plasmid) | 1.096769 | 1.027580591 | -0.09401 | 0.070288 | -3.03165 |
| hypothetical protein ABUW_4 | 0.915555 | 1.033922331 | 0.17541 | 0.070318 | 2.973226 |
| integral membrane protein M | 0.918421 | 1.016374214 | 0.146204 | 0.070426 | 3.159501 |
| fatty acid desaturase...256 | 1.11774 | 0.989758432 | -0.17544 | 0.070453 | -3.17509 |
| prophage LambdaCh01, coat p | 1.086881 | 1.019075582 | -0.09293 | 0.070484 | -3.27275 |
| muconolactone delta-isomera | 1.014392 | 0.875071814 | -0.21314 | 0.071014 | -2.54928 |
| BolA family protein | 1.140101 | 0.907575302 | -0.32907 | 0.071077 | -3.15974 |
| hypothetical protein ABUW_2 | 1.197706 | 0.966168097 | -0.30993 | 0.071126 | -3.23843 |
| serine/threonine protein kina | 1.03901 | 0.801851866 | -0.3738 | 0.071135 | -2.46444 |
| hypothetical protein ABUW_0 | 1.14916 | 0.907647385 | -0.34038 | 0.071397 | -3.51404 |
| phosphate starvation-inducib | 0.933639 | 0.998072525 | 0.09628 | 0.071658 | 2.506871 |
| ribosomal protein S5 | 1.060063 | 0.97951099 | -0.11402 | 0.071676 | -3.37342 |
| succinylglutamic semialdehyd | 0.951301 | 1.062133242 | 0.158991 | 0.071749 | 2.551154 |
| hypothetical protein ABUW_1 | 0.957722 | 1.10336332 | 0.20423 | 0.071771 | 3.141011 |
| hypothetical protein ABUW_0 | 1.017865 | 0.915008579 | -0.15369 | 0.071774 | -2.76967 |
| thiosulfate-binding protein...5 | 1.087024 | 0.924724252 | -0.23329 | 0.071835 | -3.01327 |
| glutathione S-transferase...22 | 1.021948 | 0.915003111 | -0.15947 | 0.072043 | -2.71154 |
| DNA polymerase IV | 0.998461 | 0.904984102 | -0.14181 | 0.072172 | -2.50769 |
| hypothetical protein ABUW_1 | 1.215921 | 0.921073866 | -0.40066 | 0.072231 | -3.50407 |
| lipid A biosynthesis acyltransf | 0.938977 | 1.049918621 | 0.161115 | 0.072444 | 2.860797 |
| GTP-binding protein...896 | 0.941001 | 1.0167417 | 0.111685 | 0.072674 | 2.620275 |
| anthranilate phosphoribosyltr | 0.978481 | 1.040352766 | 0.088458 | 0.072693 | 2.992101 |
| enoyl-CoA hydratase...223 | 0.726258 | 1.27450414 | 0.811382 | 0.072847 | 2.504809 |
| sodium/glutamate symporter | 0.846143 | 0.979687566 | 0.211421 | 0.072859 | 2.694106 |

| | | | | | |
|---|---|---|---|---|---|
| GTP cyclohydrolase I | 0.996605 | 0.944015784 | -0.07821 | 0.072882 | -3.17366 |
| hypothetical protein ABUW_3 | 1.105714 | 1.201627612 | 0.120011 | 0.0729 | 3.000088 |
| glutamate/aspartate transpor | 1.008581 | 1.088732055 | 0.110321 | 0.073017 | 2.65069 |
| hypothetical protein ABUW_3 | 0.992193 | 0.933541585 | -0.08791 | 0.073255 | -3.17449 |
| hypothetical protein ABUW_1 | 1.107292 | 0.853875898 | -0.37494 | 0.073291 | -3.46488 |
| glycyl-tRNA synthetase, alpha | 0.99687 | 0.952305073 | -0.06598 | 0.073334 | -2.66905 |
| NADH dehydrogenase I chain | 0.920285 | 1.012966437 | 0.138433 | 0.073583 | 2.422858 |
| glutathione-dependent forma | 1.150197 | 1.006814383 | -0.19208 | 0.073589 | -2.50394 |
| ribosomal protein S9 | 1.110847 | 0.91924088 | -0.27315 | 0.073788 | -3.39795 |
| putative transport protein | 0.919918 | 1.0898488 | 0.244551 | 0.074333 | 3.340307 |
| AraC family transcriptional re | 1.033794 | 1.144190531 | 0.146379 | 0.074334 | 2.671807 |
| phage-related membrane pro | 1.043849 | 0.883000806 | -0.24143 | 0.074445 | -2.41831 |
| molybdate ABC transporter, p | 1.088106 | 0.92822426 | -0.22927 | 0.074857 | -3.44355 |
| DNA-binding protein H-NS | 1.0647 | 0.962102453 | -0.14618 | 0.074995 | -3.3169 |
| glutathione S-transferase fam | 0.995899 | 0.960051807 | -0.05289 | 0.075096 | -2.56987 |
| hypothetical protein ABUW_3 | 0.864873 | 1.049147368 | 0.278657 | 0.07511 | 3.345895 |
| biopolymer transport protein | 0.913697 | 0.971886193 | 0.089072 | 0.075118 | 2.874195 |
| YD repeat protein | 0.905947 | 1.010781325 | 0.157973 | 0.075169 | 2.497332 |
| hypothetical protein ABUW_3 | 0.939541 | 0.954990787 | 0.023531 | 0.075213 | 2.413326 |
| transcriptional regulator, LysR | 1.022004 | 1.110197589 | 0.119416 | 0.075688 | 3.243275 |
| S-adenosyl-L-methionine-dep | 1.02435 | 0.930553451 | -0.13855 | 0.075769 | -3.04516 |
| hypothetical protein ABUW_0 | 0.928906 | 0.967200731 | 0.058283 | 0.075823 | 2.979559 |
| ribosomal protein L5 | 1.062911 | 0.900747767 | -0.23883 | 0.075828 | -3.10844 |
| 3-oxoacyl-[acyl-carrier-protei | 1.28211 | 0.758587755 | -0.75713 | 0.075949 | -3.31009 |
| 2-octaprenyl-6-methoxyphyn | 0.912624 | 0.967150201 | 0.08372 | 0.076008 | 3.201572 |
| enoyl-CoA hydratase/isomera | 1.031641 | 1.088884905 | 0.07791 | 0.076156 | 3.331069 |
| molybdopterin biosynthesis p | 1.015288 | 0.946681428 | -0.10094 | 0.076423 | -2.75064 |
| hypothetical protein ABUW_3 | 1.036298 | 0.991405457 | -0.06389 | 0.076467 | -2.52776 |
| protein YrdC | 1.043012 | 0.938880854 | -0.15174 | 0.076536 | -2.42367 |
| transcriptional regulator, TetR | 0.998396 | 0.940924525 | -0.08553 | 0.076654 | -2.52124 |
| hypothetical protein ABUW_0 | 0.951518 | 1.032803661 | 0.118263 | 0.076725 | 2.499325 |
| transcriptional factor | 1.048674 | 1.021910307 | -0.0373 | 0.07721 | -2.9098 |
| hypothetical protein ABUW_3 | 0.98262 | 0.918165551 | -0.09788 | 0.077244 | -2.93847 |
| phage-associated protein, fan | 1.007755 | 0.878355557 | -0.19827 | 0.077266 | -2.38756 |
| hypothetical protein ABUW_2 | 1.201345 | 0.667418469 | -0.84799 | 0.077316 | -3.38103 |
| phospholipid/glycerol acyltrai | 0.87493 | 1.010960933 | 0.208488 | 0.077454 | 2.739277 |
| sensory box protein | 1.008824 | 1.142124752 | 0.179046 | 0.077676 | 2.584598 |
| signal peptidase I | 0.950176 | 0.965811324 | 0.023546 | 0.07768 | 2.437959 |
| hypothetical protein ABUW_2 | 1.226387 | 0.871286383 | -0.4932 | 0.077787 | -2.72395 |
| succinate-semialdehyde dehy | 1.018183 | 1.075014204 | 0.078359 | 0.078195 | 3.053086 |
| type III restriction enzyme, re | 1.019328 | 1.067453472 | 0.066554 | 0.078262 | 2.458489 |
| uroporphyrin-III C-methyltran | 0.96303 | 1.00299288 | 0.058659 | 0.078318 | 2.434122 |
| hypothetical protein ABUW_1 | 0.860359 | 0.983577058 | 0.1931 | 0.078332 | 2.821265 |
| ribosomal protein L22 | 1.101845 | 0.919213386 | -0.26145 | 0.078335 | -3.2671 |
| phosphate regulon transcripti | 1.024484 | 0.984016008 | -0.05814 | 0.078514 | -2.83913 |
| NADH dehydrogenase fad-cor | 1.077623 | 0.848607058 | -0.34468 | 0.078536 | -2.59901 |
| type VI secretion-associated p | 0.888693 | 0.985378172 | 0.148992 | 0.078945 | 2.659446 |

| | | | | | |
|---|---|---|---|---|---|
| TonB-dependent siderophore | 0.881042 | 0.903750387 | 0.036713 | 0.078951 | 2.979927 |
| major facilitator superfamily I | 0.976919 | 1.062247228 | 0.120808 | 0.079312 | 2.352415 |
| porin...414 | 1.111584 | 0.95375694 | -0.22092 | 0.079362 | -2.57838 |
| homoserine kinase | 1.080129 | 0.827462989 | -0.38444 | 0.079381 | -3.10268 |
| CBS domain containing protei | 1.111049 | 0.97176538 | -0.19324 | 0.079419 | -3.0496 |
| leucine-responsive transcripti | 1.160977 | 1.076106713 | -0.10952 | 0.079922 | -3.00916 |
| succinyl-CoA synthetase, alph | 0.960245 | 0.861004516 | -0.15738 | 0.080009 | -2.83202 |
| tolerance to group A colicins s | 0.943341 | 1.068273427 | 0.17943 | 0.080065 | 2.458536 |
| endonuclease/exonuclease/p | 0.872165 | 1.064128803 | 0.286999 | 0.080227 | 3.021228 |
| 4-hydroxythreonine-4-phosph | 0.955908 | 1.101213069 | 0.20415 | 0.080436 | 2.721964 |
| short-chain dehydrogenase/re | 1.142453 | 1.056925511 | -0.11226 | 0.080564 | -3.00043 |
| glycine cleavage system H pro | 1.083839 | 0.887064212 | -0.28904 | 0.080577 | -3.28159 |
| DNA repair protein RecN | 0.994818 | 0.940195211 | -0.08147 | 0.080715 | -2.42492 |
| thymidylate kinase | 0.989696 | 0.890657979 | -0.15211 | 0.080872 | -3.22061 |
| hypothetical protein ABUW_0 | 1.096386 | 0.88046234 | -0.31642 | 0.081087 | -2.60226 |
| ribonuclease PH | 1.025517 | 0.879404944 | -0.22175 | 0.081386 | -3.22826 |
| hypothetical protein ABUW_1 | 0.907423 | 1.232801403 | 0.442093 | 0.081488 | 3.012024 |
| ribosomal protein L15 | 1.181659 | 0.884743711 | -0.41748 | 0.081508 | -3.24632 |
| hypothetical protein ABUW_2 | 1.01465 | 1.091586998 | 0.105445 | 0.081514 | 2.317438 |
| electron transfer flavoprotein | 1.079598 | 0.875443774 | -0.30241 | 0.081609 | -3.24238 |
| transcription elongation facto | 1.03794 | 0.971028658 | -0.09614 | 0.082331 | -2.44281 |
| hypothetical protein ABUW_1 | 1.131542 | 1.352629151 | 0.257476 | 0.08269 | 2.302755 |
| universal stress protein family | 1.077372 | 0.995137205 | -0.11455 | 0.082804 | -2.77776 |
| hypothetical protein ABUW_0 | 1.099798 | 0.953320876 | -0.2062 | 0.0833 | -2.85343 |
| hypothetical protein ABUW_2 | 1.332384 | 0.828179485 | -0.68599 | 0.083567 | -3.13576 |
| phosphatidylserine decarboxy | 0.923089 | 0.954017389 | 0.047546 | 0.083654 | 2.312434 |
| luciferase family monooxyger | 1.011833 | 1.068345126 | 0.078406 | 0.083662 | 2.571396 |
| IscR-regulated protein YhgI | 0.987763 | 0.877036654 | -0.17153 | 0.083696 | -3.01823 |
| VacJ family lipoprotein | 0.974026 | 0.929165061 | -0.06802 | 0.084303 | -2.3208 |
| hypothetical protein ABUW_0 | 1.396131 | 0.89711702 | -0.63807 | 0.084644 | -2.29642 |
| hypothetical protein ABUW_2 | 0.916553 | 0.989266933 | 0.110141 | 0.084836 | 2.411294 |
| 3-oxoadipate enol-lactonase.. | 0.986153 | 1.129282527 | 0.195524 | 0.08489 | 3.041339 |
| transcriptional regulator, TetR | 0.992069 | 0.922380079 | -0.10508 | 0.084908 | -2.44992 |
| lipopolysaccharide transport p | 0.977036 | 1.019237028 | 0.061006 | 0.085283 | 2.577457 |
| hypothetical protein ABUW_0 | 1.01177 | 1.097545493 | 0.117399 | 0.085416 | 3.050694 |
| hypothetical protein ABUW_0 | 1.127417 | 1.052667856 | -0.09897 | 0.085544 | -2.27317 |
| NADH dehydrogenase I chain | 0.9184 | 0.973372309 | 0.08387 | 0.085642 | 2.403005 |
| thioredoxin...1904 | 1.141112 | 0.882480575 | -0.3708 | 0.085969 | -3.1699 |
| O-succinylhomoserine sulfhyd | 0.878694 | 0.979159293 | 0.156183 | 0.086006 | 3.171151 |
| hypothetical protein ABUW_0 | 1.006464 | 1.091357832 | 0.116828 | 0.086358 | 2.295994 |
| isovaleryl-CoA dehydrogenase | 1.008554 | 0.93859096 | -0.10372 | 0.086409 | -3.09426 |
| intracellular septation protein | 0.991738 | 1.1011382 | 0.150964 | 0.086414 | 2.26664 |
| hypothetical protein ABUW_2 | 0.936191 | 1.007922788 | 0.10651 | 0.086516 | 3.123496 |
| thioesterase superfamily prot | 0.98224 | 1.092948179 | 0.154077 | 0.086522 | 2.59382 |
| hypothetical protein ABUW_2 | 1.383999 | 0.903367262 | -0.61546 | 0.086559 | -2.81779 |
| CDP-diacylglycerol--glycerol-3 | 0.703656 | 1.088739012 | 0.629716 | 0.086619 | 3.116269 |
| hypothetical protein ABUW_2 | 1.004413 | 0.948816213 | -0.08215 | 0.086627 | -2.263 |

| | | | | | |
|---|---|---|---|---|---|
| oligoribonuclease Orn | 0.9943 | 1.085251716 | 0.126277 | 0.086685 | 2.371366 |
| ribosomal protein S13 | 1.11322 | 0.863516681 | -0.36644 | 0.086825 | -3.08368 |
| hypothetical protein ABUW_1 | 1.095716 | 0.985374277 | -0.15313 | 0.086919 | -2.26483 |
| hypothetical protein ABUW_0 | 1.003731 | 0.864559258 | -0.21534 | 0.086938 | -2.61152 |
| hypothetical protein ABUW_2 | 1.04709 | 0.964142795 | -0.11907 | 0.087036 | -2.43342 |
| peptidyl-prolyl cis-trans isome | 1.023109 | 0.934055048 | -0.13138 | 0.087067 | -2.79999 |
| histidinol-phosphate aminotra | 0.962513 | 1.012582266 | 0.073162 | 0.087166 | 2.812134 |
| histidine kinase | 0.956423 | 0.997642104 | 0.060874 | 0.087186 | 2.440001 |
| hypothetical protein ABUW_2 | 0.956491 | 1.173222549 | 0.294654 | 0.087311 | 2.255101 |
| transcriptional regulator, AraC | 0.947961 | 1.015279765 | 0.098978 | 0.087444 | 2.389588 |
| AFG1-family ATPase | 1.03383 | 0.97318166 | -0.08722 | 0.087486 | -2.3191 |
| tol-pal system-associated acyl | 1.033545 | 0.909546281 | -0.18438 | 0.087489 | -3.11944 |
| hypothetical protein ABUW_0 | 1.114401 | 0.868858598 | -0.35907 | 0.087529 | -2.63022 |
| hypothetical protein ABUW_0 | 0.936368 | 1.009460369 | 0.108437 | 0.087767 | 2.458515 |
| 6,7-dimethyl-8-ribityllumazine | 1.039825 | 0.881165623 | -0.23886 | 0.088808 | -2.9184 |
| phosphoadenosine phosphosu | 1.073152 | 0.968871596 | -0.14748 | 0.088927 | -2.95573 |
| carbohydrate kinase family | 0.948355 | 1.000324786 | 0.076969 | 0.089115 | 2.2351 |
| hypothetical protein ABUW_3 | 1.067839 | 0.935571988 | -0.19077 | 0.089387 | -2.34536 |
| hypothetical protein ABUW_2 | 0.616487 | 1.357549181 | 1.138862 | 0.089513 | 2.842263 |
| transcriptional regulator, LysR | 1.004541 | 0.918743173 | -0.1288 | 0.090032 | -2.50604 |
| toluene tolerance efflux trans | 0.993385 | 0.930315848 | -0.09463 | 0.090114 | -2.76115 |
| carbamoyl-phosphate synthas | 1.01357 | 0.918815736 | -0.1416 | 0.090231 | -2.91067 |
| ferrichrome-iron receptor | 0.902536 | 0.994185747 | 0.139531 | 0.090707 | 2.400115 |
| hypothetical protein ABUW_3 | 0.927316 | 1.246656196 | 0.42693 | 0.090827 | 2.707329 |
| acetyltransferase...922 | 0.909167 | 1.00434054 | 0.143631 | 0.090839 | 2.350371 |
| alpha/beta hydrolase fold pro | 0.86042 | 1.037603293 | 0.270142 | 0.091023 | 2.514789 |
| hypothetical protein ABUW_1 | 1.246507 | 1.457030509 | 0.22514 | 0.091305 | 2.939495 |
| hypothetical protein ABUW_0 | 0.987963 | 0.944766545 | -0.0645 | 0.091417 | -2.75562 |
| 3,4-dihydroxy-2-butanone 4-p | 0.992387 | 0.971850736 | -0.03017 | 0.091791 | -2.21543 |
| hypothetical protein ABUW_0 | 1.000421 | 0.953110671 | -0.06989 | 0.091973 | -2.20721 |
| hypothetical protein ABUW_1 | 0.929129 | 0.89052614 | -0.06122 | 0.092327 | -2.30288 |
| translation elongation factor T | 1.04776 | 0.916325464 | -0.19338 | 0.092734 | -2.90931 |
| LemA | 1.110973 | 0.834580244 | -0.4127 | 0.092848 | -2.9302 |
| hypothetical protein ABUW_3 | 1.259476 | 0.940309495 | -0.42162 | 0.093587 | -2.4964 |
| hypothetical protein ABUW_0 | 0.883851 | 1.047888477 | 0.245611 | 0.093782 | 2.914733 |
| ribosomal protein S20 | 1.100139 | 0.93636434 | -0.23254 | 0.093794 | -2.65485 |
| RNA methyltransferase, TrmH | 1.059675 | 0.964226084 | -0.13618 | 0.093811 | -2.44533 |
| excinuclease ABC, B subunit | 0.995695 | 0.971220164 | -0.03591 | 0.093927 | -2.19278 |
| hypothetical protein ABUW_2 | 0.765213 | 1.025311318 | 0.422128 | 0.093976 | 2.646853 |
| fumarylacetoacetate hydrolas | 1.177415 | 0.896542191 | -0.39318 | 0.094378 | -2.76674 |
| transcriptional regulator...16 | 0.991821 | 1.337939202 | 0.431861 | 0.094462 | 2.337514 |
| hypothetical protein ABUW_0 | 0.959952 | 1.156901907 | 0.269232 | 0.094621 | 2.215771 |
| hypothetical protein ABUW_2 | 0.302174 | 0.535284421 | 0.824925 | 0.094746 | 2.878347 |
| major facilitator superfamily M | 1.04632 | 0.910451064 | -0.20067 | 0.095419 | -2.39784 |
| hypothetical protein ABUW_0 | 0.942378 | 0.835390219 | -0.17386 | 0.095454 | -2.26113 |
| type IV pilin structural subuni | 1.23268 | 1.102812149 | -0.16061 | 0.095622 | -2.95099 |
| glyceropghosphoryl diester ph | 1.202574 | 1.014530876 | -0.24531 | 0.09567 | -2.24359 |

| | | | | | |
|---|---|---|---|---|---|
| phage protein...602 | 1.034247 | 0.728629001 | -0.50532 | 0.09588 | -2.44609 |
| crispr-associated helicase Cas | 1.157343 | 0.869794771 | -0.41207 | 0.096007 | -2.88699 |
| dienelactone hydrolase...227( | 0.9813 | 0.946403529 | -0.05224 | 0.096233 | -2.16665 |
| peptidyl-prolyl cis-trans isome | 0.971343 | 0.952139668 | -0.02881 | 0.096367 | -2.77227 |
| ribosomal protein L31 | 1.111366 | 0.864698935 | -0.36206 | 0.09697 | -2.33705 |
| dihydroneopterin aldolase | 0.953072 | 1.002685824 | 0.073213 | 0.09705 | 2.948463 |
| flavodoxin/nitric oxide syntha | 0.937673 | 0.99604788 | 0.087129 | 0.097175 | 2.669604 |
| membrane protein involved ir | 1.000975 | 0.921447899 | -0.11943 | 0.097465 | -2.87315 |
| type VI secretion protein, fam | 0.929909 | 1.118623359 | 0.266564 | 0.09793 | 2.252475 |
| ribosomal protein L17 | 1.079942 | 0.936434906 | -0.2057 | 0.098489 | -2.49095 |
| hypothetical protein ABUW_3 | 0.966931 | 0.89607237 | -0.1098 | 0.09853 | -2.55611 |
| hypothetical protein ABUW_2 | 0.940576 | 0.998573342 | 0.086324 | 0.098894 | 2.908448 |
| multidrug efflux protein...701 | 1.117471 | 1.040310868 | -0.10322 | 0.098928 | -2.14601 |
| histidine acid phosphatase far | 1.063796 | 1.029106505 | -0.04783 | 0.098977 | -2.66181 |
| bifunctional succinylornithine | 1.018953 | 0.945329224 | -0.1082 | 0.099527 | -2.59287 |
| ABC-1 domain protein...2113 | 1.031496 | 0.993840615 | -0.05365 | 0.099894 | -2.15393 |
| protein-export membrane pro | 0.985491 | 1.058552081 | 0.103178 | 0.099941 | 2.65012 |
| tRNA pseudouridine synthase | 0.948645 | 1.055947351 | 0.154598 | 0.099989 | 2.390905 |
| hypothetical protein ABUW_2 | 1.060829 | 0.998714872 | -0.08705 | 0.100083 | -2.53403 |
| NADH:flavin oxidoreductase/r | 1.127243 | 1.04266878 | -0.11252 | 0.100141 | -2.2161 |
| ribosomal protein L21 | 1.076242 | 0.930980283 | -0.20918 | 0.100456 | -2.84295 |
| putative RND family drug tran | 1.055861 | 1.102786083 | 0.062733 | 0.100598 | 2.247994 |
| ribosomal protein L13 | 1.149074 | 0.870339216 | -0.40082 | 0.100635 | -2.65557 |
| NAD dependent epimerase/de | 1.122741 | 0.918240733 | -0.29008 | 0.100817 | -2.86122 |
| glycosyltransferase...1269 | 0.910091 | 0.980453744 | 0.107438 | 0.10088 | 2.565986 |
| FHA domain protein | 1.163866 | 0.776906536 | -0.58311 | 0.10099 | -2.83708 |
| inner membrane protein | 1.056562 | 0.895819919 | -0.2381 | 0.10115 | -2.41157 |
| porin B...946 | 1.025105 | 0.934643609 | -0.13328 | 0.101156 | -2.12234 |
| hypothetical protein ABUW_2 | 1.073909 | 0.80485203 | -0.41608 | 0.101192 | -2.87217 |
| cell division protein FtsQ | 0.93781 | 1.0342456 | 0.141212 | 0.101243 | 2.223346 |
| 2-amino-4-hydroxy-6-hydroxy | 0.918992 | 1.010201779 | 0.136519 | 0.101343 | 2.1236 |
| nucleoside-diphosphate-sugar | 0.99556 | 1.11987187 | 0.169753 | 0.101647 | 2.39335 |
| hypothetical protein ABUW_0 | 1.085551 | 0.945755889 | -0.19889 | 0.101778 | -2.74848 |
| Na+/H+ antiporter NhaC | 0.992031 | 1.041397489 | 0.070064 | 0.101785 | 2.127638 |
| type VI secretion protein, Evp | 0.989277 | 1.086174944 | 0.134809 | 0.102165 | 2.226033 |
| ATP synthase F1, epsilon subu | 1.007842 | 0.947482999 | -0.0891 | 0.102237 | -2.79884 |
| hypothetical protein ABUW_2 | 1.005732 | 0.924938187 | -0.12082 | 0.10342 | -2.43182 |
| hypothetical protein ABUW_1 | 0.799372 | 0.979843298 | 0.293684 | 0.103552 | 2.552608 |
| glutamine amidotransferase | 1.029711 | 1.105311229 | 0.102213 | 0.103631 | 2.199265 |
| hypothetical protein ABUW_0 | 1.053968 | 0.987814078 | -0.09352 | 0.103844 | -2.5114 |
| hypothetical protein ABUW_3 | 0.979137 | 1.003067856 | 0.034837 | 0.104142 | 2.813151 |
| general substrate transporter | 0.848212 | 0.880272225 | 0.053524 | 0.104329 | 2.09606 |
| hypothetical protein ABUW_1 | 0.959613 | 1.090421646 | 0.184361 | 0.104378 | 2.664784 |
| site-specific DNA-methyltrans | 1.039139 | 1.194364541 | 0.200854 | 0.104386 | 2.09969 |
| ThiJ/PfpI domain protein | 1.166786 | 0.985407302 | -0.24375 | 0.104426 | -2.53691 |
| anthranilate synthase compon | 0.935229 | 1.119456528 | 0.259407 | 0.104524 | 2.79401 |
| 2,4-dienoyl-CoA reductase | 0.921935 | 1.097945277 | 0.252069 | 0.104877 | 2.289765 |

| | | | | | |
|---|---|---|---|---|---|
| acyl-CoA dehydrogenase...555 | 1.049375 | 0.894273587 | -0.23074 | 0.104904 | -2.75097 |
| glutamyl-tRNA(Gln) amidotrai | 1.186941 | 0.859978942 | -0.46488 | 0.105041 | -2.68095 |
| ribosomal protein L28 | 1.081188 | 0.951266911 | -0.18469 | 0.105364 | -2.26093 |
| auxin efflux Carrier | 1.206231 | 1.039537342 | -0.21456 | 0.1054 | -2.09453 |
| anhydro-N-acetylmuramic aci | 1.092597 | 1.15193945 | 0.076303 | 0.105582 | 2.171253 |
| preprotein translocase, SecE s | 0.923999 | 0.997730766 | 0.110759 | 0.105603 | 2.665486 |
| putative transcriptional regula | 0.932596 | 1.086300355 | 0.220099 | 0.105836 | 2.456651 |
| histidine ammonia-lyase | 1.00048 | 1.042228172 | 0.058979 | 0.10594 | 2.165986 |
| DNA polymerase III, epsilon su | 1.008084 | 1.147357559 | 0.186699 | 0.106084 | 2.079614 |
| histidine triad protein...1546 | 1.045719 | 0.973180242 | -0.10372 | 0.106155 | -2.536 |
| methionine-S-sulfoxide reduc | 0.974106 | 0.914120666 | -0.09169 | 0.106657 | -2.48043 |
| GCN5-related N-acetyltransfe | 1.024866 | 0.96782561 | -0.08262 | 0.106812 | -2.50795 |
| hypothetical protein ABUW_2 | 0.887137 | 1.012340742 | 0.190466 | 0.107026 | 2.388912 |
| anti-sigma factor antagonist | 0.984315 | 1.037938449 | 0.076529 | 0.107242 | 2.573933 |
| biotin biosynthesis protein Bi | 1.282987 | 1.026175082 | -0.32223 | 0.107257 | -2.07787 |
| transcriptional regulator, TetF | 0.980854 | 1.016035432 | 0.050841 | 0.10742 | 2.076301 |
| FilC | 0.96813 | 1.1732091 | 0.277187 | 0.10752 | 2.548056 |
| peptidase M23/M37 family | 0.953091 | 1.076626278 | 0.175832 | 0.107736 | 2.675202 |
| glutathione S-transferase fam | 1.050524 | 1.190967228 | 0.181025 | 0.108205 | 2.404405 |
| hypothetical protein ABUW_3 | 1.213987 | 1.016035932 | -0.2568 | 0.108505 | -2.73893 |
| hypothetical protein ABUW_2 | 1.118835 | 1.000114084 | -0.16183 | 0.108922 | -2.63413 |
| 2-oxoglutarate dehydrogenas | 0.966809 | 0.888812094 | -0.12135 | 0.109098 | -2.65561 |
| hypothetical protein ABUW_2 | 1.107373 | 0.951693831 | -0.21857 | 0.109631 | -2.19061 |
| thioesterase superfamily | 1.173841 | 0.920437858 | -0.35084 | 0.110099 | -2.2951 |
| hypothetical protein ABUW_3 | 1.207923 | 0.822715939 | -0.55406 | 0.110133 | -2.06237 |
| transcriptional regulator, LysF | 1.068137 | 1.012124273 | -0.07771 | 0.110648 | -2.35571 |
| ABC transporter ATP-binding | 0.915036 | 1.043104512 | 0.188984 | 0.110725 | 2.533377 |
| glutathione import ATP-bindir | 1.088615 | 1.184031668 | 0.121214 | 0.111206 | 2.615139 |
| oligopeptidase A | 0.95992 | 1.022235609 | 0.090741 | 0.111301 | 2.507042 |
| coenzyme PQQ synthesis prot | 1.054973 | 1.190170569 | 0.173963 | 0.111407 | 2.615011 |
| permease...1066 | 1.058367 | 0.84404558 | -0.32645 | 0.111412 | -2.4651 |
| cold-shock domain protein | 0.924108 | 1.055769619 | 0.192162 | 0.111433 | 2.210731 |
| hypothetical protein ABUW_0 | 1.075896 | 1.172081537 | 0.123534 | 0.111443 | 2.179838 |
| threonine synthase | 0.8924 | 0.92912635 | 0.058185 | 0.111485 | 2.515054 |
| GTP-binding protein Obg/CgtA | 1.097876 | 1.052760658 | -0.06054 | 0.111649 | -2.13307 |
| indole-3-glycerol phosphate s | 0.955636 | 1.070387906 | 0.1636 | 0.112093 | 2.659306 |
| hypothetical protein ABUW_3 | 0.789308 | 1.00356078 | 0.346468 | 0.112171 | 2.307573 |
| hypothetical protein ABUW_1 | 0.876593 | 0.968159267 | 0.143337 | 0.112293 | 2.032571 |
| Smr protein/MutS2 | 0.934456 | 1.012404489 | 0.115588 | 0.112307 | 2.333891 |
| orotate phosphoribosyltransf | 0.942755 | 0.867262428 | -0.12041 | 0.112528 | -2.66993 |
| methylisocitrate lyase | 0.897206 | 0.944394081 | 0.073949 | 0.112679 | 2.331286 |
| ribonuclease | 1.110316 | 0.954383065 | -0.21833 | 0.113077 | -2.41419 |
| hypothetical protein ABUW_1 | 1.009785 | 1.048464478 | 0.05423 | 0.113146 | 2.037335 |
| peptidase M1, alanyl aminope | 0.961192 | 0.985395491 | 0.035878 | 0.113282 | 2.151905 |
| phosphoenolpyruvate-proteir | 1.004908 | 0.967367048 | -0.05493 | 0.113623 | -2.32332 |
| metalloprotease | 0.980174 | 1.089429807 | 0.152463 | 0.113738 | 2.377839 |
| ribosomal protein L9 | 1.028763 | 0.954289645 | -0.10841 | 0.113923 | -2.5538 |

| | | | | | |
|---|---|---|---|---|---|
| ketol-acid reductoisomerase | 1.005912 | 0.912118501 | -0.14121 | 0.114369 | -2.63922 |
| hypothetical protein ABUW_5 | 0.977218 | 0.929219826 | -0.07266 | 0.11454 | -2.10565 |
| transcriptional regulator, TetF | 0.894553 | 1.004725281 | 0.167562 | 0.114747 | 2.231104 |
| multidrug efflux protein AdeB | 1.041857 | 0.996147261 | -0.06473 | 0.114897 | -2.61343 |
| methylcrotonoyl-CoA carboxy | 1.191998 | 0.914970848 | -0.38158 | 0.11528 | -2.14391 |
| S-adenosylmethionine:tRNA r | 0.972175 | 0.90191568 | -0.10822 | 0.115405 | -2.10482 |
| pirin domain protein...1437 | 1.165933 | 0.985441326 | -0.24264 | 0.11544 | -2.66432 |
| transcriptional regulator, AraC | 1.042841 | 0.9614742 | -0.1172 | 0.115468 | -2.04177 |
| putative class A beta-lactama: | 1.085756 | 1.022315005 | -0.08686 | 0.11569 | -2.47995 |
| hypothetical protein ABUW_1 | 0.935164 | 1.03169278 | 0.141722 | 0.115697 | 2.066115 |
| two-component system senso | 0.838775 | 0.904578186 | 0.108961 | 0.115986 | 2.5077 |
| putative prophage protein | 1.012668 | 1.111375473 | 0.134185 | 0.116148 | 2.129284 |
| tRNA nucleotidyl transferase | 0.955851 | 1.027158841 | 0.103802 | 0.116885 | 2.111006 |
| hypothetical protein ABUW_3 | 0.978011 | 1.170057413 | 0.258657 | 0.117061 | 2.30189 |
| oxygen-insensitive NAD(P)H n | 1.069018 | 0.959444015 | -0.15602 | 0.117309 | -2.56362 |
| DoxX family protein | 1.033287 | 1.136286628 | 0.137085 | 0.117339 | 1.991028 |
| 3-methyl-2-oxobutanoate hyc | 0.946672 | 0.911794651 | -0.05416 | 0.11736 | -2.23294 |
| secretion protein HlyD...1783 | 1.012101 | 0.966558287 | -0.06642 | 0.117592 | -2.57323 |
| D-amino acid dehydrogenase | 0.868933 | 1.056714378 | 0.282268 | 0.118049 | 2.531562 |
| zinc-binding dehydrogenase | 1.042489 | 0.957170056 | -0.12319 | 0.11847 | -2.14204 |
| two-component system respc | 1.008233 | 0.955536398 | -0.07745 | 0.118476 | -2.30293 |
| putative arsenate reductase | 1.190941 | 0.95353754 | -0.32074 | 0.118544 | -2.57102 |
| ATP synthase F1, delta subuni | 1.006929 | 0.882066946 | -0.191 | 0.118606 | -2.63116 |
| methionine aminopeptidase, | 1.053937 | 0.913610924 | -0.20614 | 0.118725 | -2.40513 |
| lipid A phosphoethanolamine | 1.107891 | 0.98057731 | -0.17611 | 0.118748 | -2.10327 |
| YCII-related protein...752 | 0.870076 | 1.014570177 | 0.221655 | 0.118858 | 2.589997 |
| 3-oxoadipate CoA-transferase | 0.894045 | 1.011563765 | 0.178168 | 0.11886 | 2.574229 |
| cysteine desulfurase | 0.842552 | 1.199041761 | 0.509044 | 0.118908 | 2.179049 |
| glutamyl-tRNA synthetase...43 | 1.104957 | 0.780310227 | -0.50187 | 0.118957 | -2.2266 |
| general secretory pathway pr | 1.186145 | 1.010448703 | -0.23128 | 0.119007 | -2.54037 |
| nitrogen metabolism transcrip | 0.981298 | 1.062113615 | 0.114175 | 0.119181 | 2.350739 |
| non-canonical purine NTP pyr | 1.084093 | 0.81961515 | -0.40347 | 0.119272 | -2.61377 |
| CobW/P47K family protein | 1.124047 | 0.981603753 | -0.19549 | 0.119353 | -1.99722 |
| 2-C-methyl-D-erythritol 2,4-cy | 0.996119 | 0.938191657 | -0.08644 | 0.119407 | -2.30804 |
| rubredoxin | 0.863503 | 1.076073033 | 0.317504 | 0.119995 | 2.319733 |
| transcriptional regulator, AraC | 0.889054 | 0.951891288 | 0.098526 | 0.120285 | 2.255652 |
| rieske 2Fe-2S family protein | 1.078728 | 0.897033002 | -0.2661 | 0.120696 | -2.36754 |
| 3-oxoadipate CoA-transferase | 1.003727 | 0.905744391 | -0.14819 | 0.120889 | -2.51366 |
| hypothetical protein ABUW_2 | 0.938131 | 1.109877035 | 0.242539 | 0.120907 | 2.284509 |
| 3-dehydroquinate dehydratas | 1.256793 | 0.883621623 | -0.50825 | 0.121324 | -2.56892 |
| monooxygenase, NtaA/SnaA/ | 1.084861 | 0.972115551 | -0.15831 | 0.121332 | -2.48657 |
| cytochrome B561...383 | 1.001602 | 1.103049518 | 0.139189 | 0.121355 | 1.967569 |
| protein YegH | 0.985358 | 1.014093067 | 0.04147 | 0.122086 | 2.559002 |
| hypothetical protein ABUW_3 | 0.949455 | 1.024869034 | 0.110267 | 0.122205 | 2.023438 |
| hypothetical protein ABUW_0 | 0.889362 | 0.951227978 | 0.097021 | 0.122775 | 1.956511 |
| hypothetical protein ABUW_0 | 1.015422 | 1.115098914 | 0.135092 | 0.122902 | 2.18191 |
| sodium/proline symporter | 0.955023 | 1.106580685 | 0.212501 | 0.123153 | 2.456048 |

| | | | | | |
|---|---|---|---|---|---|
| peptidoglycan-associated lipo | 1.015294 | 0.973446814 | -0.06072 | 0.123226 | -2.18348 |
| N-acetylmuramoyl-L-alanine a | 1.01033 | 0.961004619 | -0.07221 | 0.123334 | -2.11633 |
| glutathione peroxidase...2311 | 1.097276 | 0.943905969 | -0.21721 | 0.123423 | -2.35504 |
| acyl-CoA dehydrogenase...20( | 1.01632 | 0.976792851 | -0.05723 | 0.12377 | -2.0401 |
| riboflavin synthase, alpha sub | 0.878721 | 0.975618408 | 0.150911 | 0.123941 | 2.449441 |
| transcriptional regulator-like | 1.078365 | 0.93870744 | -0.2001 | 0.123945 | -1.94511 |
| ureidoglycolate hydrolase | 1.006761 | 1.077523664 | 0.097999 | 0.123961 | 1.942923 |
| hypothetical protein ABUW_1 | 0.680815 | 0.971778232 | 0.513363 | 0.124524 | 2.459394 |
| exonuclease V alpha subunit | 1.083009 | 0.967746427 | -0.16234 | 0.124546 | -2.39726 |
| hypothetical protein ABUW_1 | 1.371443 | 0.912856794 | -0.58723 | 0.124868 | -2.45725 |
| transglycosylase | 0.982477 | 1.022161922 | 0.057128 | 0.124908 | 1.941704 |
| beta-lactamase ADC7 | 1.01139 | 0.977413309 | -0.0493 | 0.124943 | -1.95204 |
| outer membrane protein A | 1.057225 | 0.971274792 | -0.12233 | 0.125274 | -2.42741 |
| ribosomal protein S17 | 1.189543 | 0.865570077 | -0.45868 | 0.125337 | -2.52882 |
| hypothetical protein ABUW_1 | 0.872618 | 1.495751735 | 0.777449 | 0.125825 | 2.031022 |
| hypothetical protein ABUW_1 | 0.961786 | 1.113458715 | 0.21126 | 0.125828 | 2.050659 |
| phosphogluconate dehydrata | 0.961232 | 0.993446703 | 0.047558 | 0.12585 | 2.205026 |
| 3-hydroxyisobutyrate dehydr | 1.036261 | 0.966089093 | -0.10116 | 0.126372 | -2.13163 |
| pyruvate decarboxylase E1 co | 1.023639 | 0.956998277 | -0.09712 | 0.126834 | -2.46239 |
| inositol-1-monophosphate | 1.064691 | 0.912913832 | -0.22188 | 0.126836 | -2.21626 |
| deoxyguanosinetriphosphate | 1.017537 | 1.119436561 | 0.137692 | 0.126995 | 2.225597 |
| Zn-dependent oligopeptidase | 1.046168 | 1.098319336 | 0.070183 | 0.127018 | 2.454923 |
| UDP-N-acetylmuramate--alan | 0.974953 | 1.009298124 | 0.049948 | 0.1271 | 2.039594 |
| hypothetical protein ABUW_( | 0.919551 | 0.961977595 | 0.065074 | 0.127412 | 2.375893 |
| hypothetical protein ABUW_( | 1.000569 | 0.935146268 | -0.09756 | 0.127696 | -2.25123 |
| ribosomal subunit interface p | 1.022095 | 0.869775626 | -0.23281 | 0.128196 | -2.43053 |
| TonB family protein | 0.845189 | 1.029296605 | 0.284313 | 0.128223 | 2.182199 |
| enoyl-CoA hydratase/isomera | 0.985319 | 1.009671007 | 0.035222 | 0.128456 | 1.973488 |
| hypothetical protein ABUW_( | 1.04812 | 0.876300863 | -0.25831 | 0.128695 | -2.00431 |
| TonB-dependent receptor prc | 0.952743 | 0.912333328 | -0.06253 | 0.128852 | -2.23895 |
| transcriptional regulator, IclR | 1.042989 | 0.994620693 | -0.06851 | 0.129239 | -1.91261 |
| hypothetical protein ABUW_2 | 1.082904 | 1.037869554 | -0.06128 | 0.129266 | -2.45389 |
| hypothetical protein ABUW_3 | 1.125486 | 1.051369034 | -0.09828 | 0.129366 | -1.98876 |
| transcriptional regulator, Gntl | 1.059832 | 0.895496445 | -0.24308 | 0.129441 | -1.91157 |
| hypothetical protein ABUW_1 | 0.96324 | 1.056650953 | 0.133532 | 0.129629 | 2.044574 |
| fructose-bisphosphate aldolas | 1.018417 | 0.961215958 | -0.0834 | 0.129802 | -2.12613 |
| GTP-binding protein LepA | 0.961305 | 0.933158764 | -0.04287 | 0.130082 | -2.13939 |
| ribosomal large subunit pseu | 0.885206 | 0.969338971 | 0.130988 | 0.130236 | 2.479332 |
| S-formylglutathione hydrolase | 1.154991 | 0.892361998 | -0.37218 | 0.130737 | -2.44879 |
| ribosomal protein L34 | 1.481878 | 0.850696914 | -0.80071 | 0.131073 | -2.35712 |
| translation elongation factor | 1.03277 | 0.886411449 | -0.22047 | 0.131782 | -2.3937 |
| ribosomal protein S18 | 1.145549 | 0.863885607 | -0.40713 | 0.131941 | -2.44642 |
| transcriptional regulator, LysF | 1.060909 | 0.98236538 | -0.11097 | 0.131945 | -1.89661 |
| alcohol dehydrogenase | 1.050552 | 0.866118391 | -0.27851 | 0.13282 | -2.24007 |
| nitroreductase family protein | 1.066646 | 0.981569065 | -0.11992 | 0.133108 | -2.14638 |
| stringent starvation protein B | 0.87022 | 1.033570229 | 0.248185 | 0.133214 | 2.296509 |
| carbapenem-hydrolyzing oxa( | 1.028534 | 0.991782496 | -0.05249 | 0.133793 | -2.11204 |

| | | | | | |
|---|---|---|---|---|---|
| ribosomal protein L16 | 1.14352 | 0.89933594 | -0.34655 | 0.134107 | -2.40902 |
| ribosomal protein S1 | 1.059758 | 0.946507217 | -0.16305 | 0.134273 | -2.43651 |
| protein YaaA | 1.073743 | 0.965740575 | -0.15294 | 0.134763 | -2.13605 |
| hypothetical protein ABUW_2 | 0.881123 | 1.346104889 | 0.611376 | 0.134811 | 2.257669 |
| heat shock protein 33 | 1.040158 | 1.053223633 | 0.018009 | 0.135761 | 1.864324 |
| acetyl-CoA acetyltransferase.. | 0.991446 | 0.87745308 | -0.17621 | 0.135959 | -2.40185 |
| hypothetical protein ABUW_3 | 0.884173 | 0.814600949 | -0.11824 | 0.136268 | -2.23677 |
| CsuB | 0.916101 | 0.957261172 | 0.063406 | 0.136388 | 1.988954 |
| glycyl-tRNA synthetase, beta s | 1.014842 | 0.965958352 | -0.07122 | 0.136953 | -2.32554 |
| feruloyl-CoA synthetase | 1.118703 | 0.963667306 | -0.21522 | 0.137008 | -2.29799 |
| rhodanese domain protein...1 | 1.004633 | 1.03615379 | 0.044569 | 0.137133 | 2.070674 |
| Sec-independent protein tran | 1.40014 | 0.853745283 | -0.71369 | 0.137193 | -2.3756 |
| isochorismate hydrolase | 1.086188 | 0.917044531 | -0.24421 | 0.137527 | -2.34782 |
| hypothetical protein ABUW_1 | 1.03614 | 0.926457801 | -0.16142 | 0.137563 | -2.40254 |
| response regulator protein | 1.033265 | 0.958310639 | -0.10864 | 0.137586 | -1.95595 |
| glutamate N-acetyltransferase | 0.9898 | 0.93104381 | -0.08829 | 0.13803 | -2.34436 |
| peroxiredoxin | 1.092348 | 0.903110926 | -0.27446 | 0.138838 | -2.29952 |
| short-chain dehydrogenase/re | 0.980355 | 1.005349532 | 0.036321 | 0.138916 | 1.924837 |
| iron uptake factor | 0.973857 | 1.100056161 | 0.175795 | 0.139078 | 1.845326 |
| 3-deoxy-manno-octulosonate | 0.966792 | 0.990839333 | 0.035446 | 0.139164 | 1.856196 |
| TonB-dependent receptor pro | 1.024244 | 1.10450884 | 0.108845 | 0.139324 | 2.286089 |
| transcriptional regulator SoxR | 1.095024 | 1.016964822 | -0.10669 | 0.139647 | -1.92598 |
| pyridoxal phosphate biosynth | 0.998153 | 1.013349719 | 0.021799 | 0.139678 | 1.970243 |
| Glu-tRNA amidotransferase | 1.033175 | 0.975982681 | -0.08216 | 0.140634 | -2.05817 |
| dihydrodipicolinate reductase | 0.979045 | 0.949435586 | -0.04431 | 0.140647 | -2.08928 |
| hypothetical protein ABUW_0 | 1.026366 | 0.90648319 | -0.17919 | 0.141187 | -2.24317 |
| hypothetical protein ABUW_2 | 0.947916 | 1.014778628 | 0.098334 | 0.141714 | 1.845226 |
| oxidoreductase...2485 | 1.034595 | 0.930636538 | -0.15278 | 0.141829 | -2.14162 |
| amino-acid permease | 0.896495 | 0.998683457 | 0.155731 | 0.142512 | 1.974532 |
| UDP-glucose/GDP-mannose d | 0.957852 | 0.989080551 | 0.046286 | 0.142625 | 1.908032 |
| hypothetical protein ABUW_2 | 0.636585 | 0.566942468 | -0.16715 | 0.142778 | -2.11648 |
| crispr-associated protein, Csy | 0.956207 | 1.071114912 | 0.163719 | 0.142786 | 2.133588 |
| 1,6-dihydroxycyclohexa-2,4-d | 0.846691 | 1.089087481 | 0.363212 | 0.142847 | 2.015146 |
| transcriptional Regulator, Ara | 0.980435 | 1.057020773 | 0.10851 | 0.142882 | 2.018289 |
| outer membrane assembly lip | 1.029682 | 1.011327594 | -0.02595 | 0.143012 | -2.03622 |
| hypothetical protein ABUW_1 | 1.025828 | 0.889856067 | -0.20514 | 0.14332 | -2.33183 |
| hypothetical protein ABUW_0 | 1.173844 | 0.981991109 | -0.25746 | 0.143523 | -2.14364 |
| L-lactate dehydrogenase (cyto | 0.96516 | 0.942166804 | -0.03479 | 0.143702 | -1.83232 |
| hypothetical protein ABUW_2 | 0.948735 | 1.046101104 | 0.140945 | 0.143797 | 2.151843 |
| phage tail completion protein | 0.917258 | 1.125640698 | 0.295348 | 0.143918 | 2.333752 |
| hypothetical protein ABUW_2 | 0.964818 | 0.824041361 | -0.22754 | 0.144001 | -2.14899 |
| inner membrane protein OxaA | 1.08513 | 0.951564676 | -0.18949 | 0.144077 | -2.23557 |
| inorganic diphosphatase | 0.97373 | 0.929082922 | -0.06771 | 0.144087 | -2.18648 |
| hypothetical protein ABUW_1 | 1.249238 | 1.089812633 | -0.19697 | 0.144161 | -1.92214 |
| hypothetical protein ABUW_0 | 1.01534 | 1.092302988 | 0.105411 | 0.144207 | 1.819528 |
| hypothetical protein ABUW_0 | 0.979867 | 0.910805361 | -0.10544 | 0.14459 | -2.32335 |
| ribosomal-protein-alanine ace | 0.911792 | 0.960085168 | 0.074457 | 0.144619 | 1.880212 |

| | | | | | |
|---|---|---|---|---|---|
| peptide chain release factor 3 | 0.925536 | 0.957350031 | 0.048758 | 0.14487 | 2.003602 |
| hypothetical protein ABUW_( | 0.979571 | 0.716967509 | -0.45024 | 0.145056 | -2.32728 |
| short-chain dehydrogenase/r( | 1.016986 | 0.927175724 | -0.13338 | 0.145193 | -2.06629 |
| aldo/keto reductase...2262 | 1.072052 | 1.024534274 | -0.06541 | 0.145254 | -2.03024 |
| 3-oxoacyl-[acyl-carrier-protein | 1.065221 | 1.189994022 | 0.159802 | 0.145683 | 1.971685 |
| hypothetical protein ABUW_2 | 0.927062 | 1.056203537 | 0.18815 | 0.145795 | 2.152844 |
| hypothetical protein ABUW_2 | 0.90957 | 1.101973193 | 0.276833 | 0.147158 | 2.057996 |
| indole-3-pyruvate decarboxyl | 0.942092 | 1.072156141 | 0.186575 | 0.147203 | 1.92145 |
| iron-sulfur cluster assembly a | 0.794804 | 1.045486158 | 0.395502 | 0.147221 | 2.175016 |
| (di)nucleoside polyphosphate | 1.00945 | 0.961287985 | -0.07053 | 0.147865 | -2.13799 |
| activator of HSP90 ATPase...2 | 1.112042 | 1.017796928 | -0.12776 | 0.148538 | -2.08814 |
| hypothetical protein ABUW_( | 0.956142 | 1.069371579 | 0.161466 | 0.148763 | 1.816362 |
| RNA helicase | 1.028023 | 1.00134815 | -0.03793 | 0.14879 | -2.24799 |
| membrane-associated Zn-dep | 1.061232 | 0.829748037 | -0.355 | 0.148826 | -2.21236 |
| YgfB/YecA family protein | 1.037134 | 0.972901821 | -0.09224 | 0.149197 | -1.90533 |
| ATP synthase F1, beta subunit | 0.983767 | 0.917647235 | -0.10038 | 0.149594 | -2.11389 |
| ATP-dependent helicase HrpA | 0.87179 | 0.951023681 | 0.125501 | 0.149943 | 1.914221 |
| 3-oxoadipate CoA-transferase | 1.385965 | 0.720185906 | -0.94445 | 0.150206 | -2.26964 |
| uroporphyrinogen-III synthase | 0.945053 | 1.067905244 | 0.176316 | 0.150435 | 2.133312 |
| hypothetical protein ABUW_1 | 0.958824 | 0.926226006 | -0.0499 | 0.150568 | -1.96743 |
| NAD(P)H dehydrogenase, quir | 0.801685 | 1.126440403 | 0.490663 | 0.151138 | 2.230814 |
| heme-binding protein A...945 | 1.010528 | 0.963834877 | -0.06825 | 0.15159 | -1.77338 |
| flavodoxin/nitric oxide syntha | 1.017433 | 1.084161374 | 0.091646 | 0.151823 | 1.801241 |
| Na+/solute symporter | 1.00948 | 0.925487654 | -0.12533 | 0.152151 | -2.10682 |
| glutathione peroxidase...2012 | 1.082592 | 0.970679569 | -0.15742 | 0.153009 | -2.06176 |
| DNA-directed RNA polymeras | 1.002099 | 0.975654512 | -0.03858 | 0.153114 | -2.03966 |
| P-aminobenzoate synthetase | 1.197987 | 0.928386975 | -0.36781 | 0.15345 | -1.90987 |
| lipoprotein-releasing system t | 0.905773 | 0.95678244 | 0.079042 | 0.153723 | 1.979188 |
| transcriptional regulator, Asn( | 0.987005 | 0.898009195 | -0.13633 | 0.153973 | -1.77557 |
| DNA replication protein (plasr | 0.913502 | 1.02693574 | 0.168866 | 0.15403 | 2.019402 |
| FilD | 0.804554 | 1.048729533 | 0.382382 | 0.155292 | 2.029739 |
| transport protein | 1.057782 | 1.004205586 | -0.07499 | 0.155295 | -2.03331 |
| hypothetical protein ABUW_3 | 0.891107 | 0.986774439 | 0.147121 | 0.155673 | 1.749809 |
| outer membrane lipoprotein I | 0.963703 | 1.019958956 | 0.081851 | 0.155751 | 1.855794 |
| arabinose 5-phosphate isome | 0.976147 | 1.005092405 | 0.042158 | 0.156073 | 1.864484 |
| hypothetical protein ABUW_3 | 1.00654 | 0.982349592 | -0.0351 | 0.156285 | -1.80068 |
| malate dehydrogenase...2408 | 0.940396 | 0.888362489 | -0.08212 | 0.156852 | -2.14295 |
| hypothetical protein ABUW_4 | 1.104144 | 1.023334406 | -0.10965 | 0.157161 | -1.83478 |
| putative ATP-binding protein | 0.953825 | 0.991582742 | 0.056009 | 0.157403 | 1.771631 |
| alcohol dehydrogenase, iron-( | 1.135811 | 0.984309332 | -0.20654 | 0.157419 | -2.04713 |
| two component signal transdu | 1.000097 | 0.960116486 | -0.05886 | 0.157899 | -2.05339 |
| hypothetical protein ABUW_3 | 0.997739 | 1.113658241 | 0.158572 | 0.157913 | 2.046595 |
| translation initiation factor IF- | 1.174549 | 0.968734711 | -0.27793 | 0.157922 | -2.19626 |
| D-3-phosphoglycerate dehydr | 0.915933 | 0.94212028 | 0.04067 | 0.158442 | 1.923531 |
| D-isomer specific 2-hydroxyac | 0.975655 | 1.044437196 | 0.098283 | 0.158454 | 2.167107 |
| GatB/Yqey domain protein | 1.062013 | 1.023720906 | -0.05298 | 0.159298 | -1.93338 |
| hypothetical protein ABUW_1 | 1.022868 | 0.959844946 | -0.09175 | 0.159309 | -1.76035 |

| | | | | | |
|---|---|---|---|---|---|
| hypothetical protein ABUW_1 | 1.005525 | 0.959171615 | -0.06809 | 0.159709 | -1.73183 |
| uracil-DNA glycosylase | 1.027409 | 1.00571787 | -0.03079 | 0.160093 | -1.88306 |
| hypothetical protein ABUW_1 | 1.012493 | 0.973951683 | -0.05599 | 0.160177 | -1.87347 |
| ring hydroxylating dioxygenas | 0.945293 | 1.016443288 | 0.104696 | 0.160394 | 1.850625 |
| folylpolyglutamate synthetase | 0.963827 | 0.923324437 | -0.06194 | 0.160452 | -1.74956 |
| TonB-dependent receptor...13 | 1.017897 | 0.957076635 | -0.08888 | 0.160589 | -1.94171 |
| ABC transport system permea | 1.082446 | 1.197617619 | 0.145872 | 0.160665 | 1.839676 |
| pseudouridine synthase A | 1.030764 | 0.890595819 | -0.21087 | 0.16068 | -1.72162 |
| hypothetical protein ABUW_2 | 1.069874 | 1.198357356 | 0.163618 | 0.160722 | 1.819606 |
| high affinity Zn transport prot | 1.017436 | 1.080282141 | 0.086471 | 0.160825 | 2.088226 |
| esterase...2063 | 1.067227 | 1.094869065 | 0.036891 | 0.161447 | 1.71957 |
| translation elongation factor ( | 1.056409 | 0.932039204 | -0.18071 | 0.161756 | -2.04735 |
| capsid scaffolding protein | 0.923966 | 0.949579677 | 0.03945 | 0.161831 | 1.713563 |
| hypothetical protein ABUW_0 | 1.005605 | 0.939931044 | -0.09744 | 0.162353 | -1.95496 |
| hypothetical protein ABUW_0 | 1.000404 | 1.133110254 | 0.179706 | 0.162453 | 1.730863 |
| serine acetyltransferase | 0.923484 | 1.053442533 | 0.189953 | 0.163116 | 2.151605 |
| glutamine-dependent NAD+ s | 0.960334 | 0.977279947 | 0.025235 | 0.163468 | 1.70882 |
| hypothetical protein ABUW_3 | 1.084401 | 0.989923652 | -0.13151 | 0.163691 | -1.72999 |
| major facilitator superfamily I | 0.884508 | 0.837452989 | -0.07887 | 0.1638 | -1.70305 |
| imidazoleglycerol-phosphate | 0.92792 | 0.956926854 | 0.044408 | 0.163816 | 2.017838 |
| hypothetical protein ABUW_0 | 0.915907 | 1.009100138 | 0.139796 | 0.163994 | 1.768826 |
| DNA repair protein RadC | 0.904155 | 1.012588217 | 0.163405 | 0.164201 | 1.795485 |
| threonine dehydratase...1260 | 0.985646 | 0.951134696 | -0.05142 | 0.16431 | -1.7967 |
| ribosomal large subunit pseud | 0.944423 | 0.997631721 | 0.079074 | 0.16434 | 1.777624 |
| hypothetical protein ABUW_0 | 1.051092 | 1.107276637 | 0.075127 | 0.164694 | 1.841748 |
| hypothetical protein ABUW_3 | 0.885617 | 0.995521305 | 0.16877 | 0.164722 | 1.88186 |
| phosphate transporter | 0.919111 | 1.026915631 | 0.160006 | 0.164809 | 2.075448 |
| hypothetical protein ABUW_2 | 1.14674 | 0.994832278 | -0.20501 | 0.164964 | -2.00168 |
| guanosine-3',5'-bis(diphospha | 1.007326 | 1.238893901 | 0.298522 | 0.16524 | 2.024524 |
| putative enoyl-CoA hydratase | 1.037431 | 0.972051158 | -0.09391 | 0.165385 | -2.03772 |
| GMC oxidoreductase | 0.784649 | 0.924508234 | 0.236638 | 0.165841 | 1.745609 |
| oxidoreductase alpha (molybd | 0.970054 | 1.039405092 | 0.099622 | 0.16629 | 1.843978 |
| copper amine oxidase | 1.168496 | 1.055648116 | -0.14652 | 0.166508 | -1.71651 |
| hypothetical protein ABUW_0 | 1.205641 | 0.853289266 | -0.49869 | 0.166569 | -2.10506 |
| hypothetical protein ABUW_0 | 0.996664 | 1.068071638 | 0.099829 | 0.166615 | 2.050616 |
| lipid A export permease/ATP- | 0.95809 | 0.950576912 | -0.01136 | 0.167162 | -2.07538 |
| hypothetical protein ABUW_4 | 1.003246 | 0.933864245 | -0.10339 | 0.167198 | -1.90796 |
| glycoside hydrolase/deacetyla | 1.26922 | 0.69974835 | -0.85903 | 0.167255 | -2.11663 |
| hypothetical protein ABUW_2 | 1.310812 | 0.923955809 | -0.50457 | 0.167308 | -2.09793 |
| ribosomal protein S12 | 1.501345 | 0.759844417 | -0.98248 | 0.16731 | -2.10769 |
| long-chain-fatty-acid--CoA liga | 1.006608 | 0.964460953 | -0.06171 | 0.167375 | -2.09292 |
| methylated-DNA--protein-cys | 0.94557 | 1.052505841 | 0.154573 | 0.168007 | 1.773847 |
| methionine-R-sulfoxide reduc | 0.84825 | 1.021386821 | 0.267969 | 0.168649 | 1.998505 |
| hypothetical protein ABUW_3 | 1.082258 | 0.949277413 | -0.18914 | 0.168659 | -1.83605 |
| beta-ketoadipyl CoA thiolase. | 0.896945 | 0.992934326 | 0.146678 | 0.168926 | 1.761738 |
| hypothetical protein ABUW_3 | 1.037098 | 0.980923323 | -0.08034 | 0.168944 | -2.09978 |
| glycosyl transferase, group 1 | 0.981527 | 1.019995435 | 0.055463 | 0.169032 | 1.685613 |

| | | | | | |
|---|---|---|---|---|---|
| penicillin-binding protein 1A | 0.97598 | 1.020767766 | 0.064731 | 0.169161 | 1.899186 |
| enoyl-coA hydratase | 0.905328 | 1.007000935 | 0.153553 | 0.169211 | 1.897944 |
| tRNA (guanine-N(7)-)-methylt | 0.954188 | 1.036025182 | 0.118714 | 0.169222 | 2.09379 |
| ExsB protein | 1.224224 | 0.900996412 | -0.44227 | 0.169594 | -2.10343 |
| imidazolonepropionase | 1.047461 | 0.910719824 | -0.20182 | 0.170035 | -2.09321 |
| outer membrane protein Opr | 0.93243 | 0.96929508 | 0.055941 | 0.170274 | 1.818202 |
| hypothetical protein ABUW_4 | 1.218697 | 0.852061815 | -0.51631 | 0.17196 | -1.88863 |
| hypothetical protein ABUW_1 | 0.999758 | 1.062508461 | 0.087824 | 0.172736 | 1.682575 |
| hypothetical protein ABUW_3 | 0.860321 | 0.946255315 | 0.137355 | 0.1728 | 1.94725 |
| ribosomal protein L11 methyl | 1.084269 | 1.032952363 | -0.06995 | 0.173475 | -1.66694 |
| malonate decarboxylase, beta | 1.118054 | 1.054452821 | -0.0845 | 0.173589 | -1.66383 |
| ferric uptake regulation prote | 0.928861 | 0.880823397 | -0.07661 | 0.173677 | -2.02574 |
| glutathione synthase | 0.988973 | 0.940160151 | -0.07303 | 0.174259 | -1.95454 |
| 4-hydroxyphenylpyruvate dio | 1.075581 | 0.950972296 | -0.17764 | 0.174497 | -1.84569 |
| 2-dehydro-3-deoxyphosphoo | 1.01007 | 0.928968742 | -0.12075 | 0.175202 | -1.9905 |
| transcriptional regulator, Gntl | 0.897548 | 1.090826703 | 0.281362 | 0.175939 | 1.826894 |
| regulatory protein RecX | 1.452751 | 0.673685797 | -1.10864 | 0.176427 | -2.02458 |
| hypothetical protein ABUW_1 | 0.857423 | 0.946502242 | 0.142598 | 0.176689 | 1.639348 |
| ribosomal protein S19 | 1.026276 | 0.999802851 | -0.0377 | 0.176915 | -1.63728 |
| phenazine biosynthesis protei | 1.0624 | 0.983888111 | -0.11076 | 0.177091 | -1.77566 |
| arsenical resistance operon re | 1.040991 | 1.275326394 | 0.292909 | 0.177106 | 1.988528 |
| cell division topological specif | 1.094413 | 0.9863787 | -0.14994 | 0.177162 | -1.76378 |
| glutamate/aspartate transpor | 0.970912 | 1.108769402 | 0.191547 | 0.177441 | 1.756974 |
| threonine ammonia-lyase | 1.005624 | 0.962114343 | -0.06381 | 0.177584 | -1.83076 |
| lipocalin family protein | 0.944418 | 1.03989984 | 0.138948 | 0.177726 | 1.816802 |
| potassium transport system lo | 0.961467 | 1.022331491 | 0.088553 | 0.177967 | 1.649075 |
| dihydroorotate oxidase | 1.015607 | 0.965814441 | -0.07252 | 0.178461 | -1.65001 |
| hypothetical protein ABUW_2 | 1.043415 | 1.002999937 | -0.05699 | 0.178798 | -1.77701 |
| transcriptional regulator, TetR | 0.933391 | 1.065889435 | 0.191505 | 0.179313 | 1.631636 |
| zinc import ATP-binding prote | 0.926794 | 1.048770951 | 0.17838 | 0.179418 | 1.835542 |
| chromosome segregation and | 0.956381 | 1.041085911 | 0.122431 | 0.1797 | 1.733547 |
| nitroreductase...1514 | 1.023947 | 0.987317463 | -0.05256 | 0.179752 | -1.63027 |
| acyl-CoA thioesterase | 1.20694 | 0.763938694 | -0.65982 | 0.179805 | -2.00973 |
| polyphosphate-AMP phospho | 0.982171 | 1.054146804 | 0.10203 | 0.179871 | 1.941779 |
| alpha/beta hydrolase...1292 | 1.03658 | 0.971708658 | -0.09324 | 0.180183 | -1.86185 |
| cupin family protein | 0.976546 | 1.033857156 | 0.082277 | 0.18061 | 1.668086 |
| nicotinate-nucleotide diphosp | 0.914368 | 0.946095441 | 0.04921 | 0.18076 | 1.657613 |
| ATP-dependent DNA helicase | 1.008007 | 0.9634657 | -0.0652 | 0.181524 | -1.61556 |
| putative polysaccharide deace | 0.93536 | 1.010812602 | 0.111922 | 0.181586 | 1.616943 |
| alpha-methylacyl-CoA racema | 1.120763 | 1.014296471 | -0.144 | 0.181589 | -1.77584 |
| 3-dehydroshikimate dehydrat | 1.265001 | 1.111816595 | -0.18622 | 0.181677 | -1.88136 |
| hypothetical protein ABUW_2 | 1.218773 | 0.810206224 | -0.58907 | 0.182026 | -1.83114 |
| acetate kinase | 1.011858 | 0.951121439 | -0.08931 | 0.182364 | -1.9293 |
| cation efflux system protein... | 1.003843 | 1.067420523 | 0.088595 | 0.182569 | 1.612256 |
| short-chain dehydrogenase/re | 0.94177 | 1.06462338 | 0.176897 | 0.182732 | 1.951055 |
| nicotinamidase | 0.933215 | 1.046580814 | 0.165403 | 0.182816 | 1.923706 |
| hypothetical protein ABUW_2 | 0.979108 | 0.855941064 | -0.19396 | 0.183199 | -1.98615 |

| | | | | | |
|---|---|---|---|---|---|
| hypothetical protein ABUW_3 | 0.892337 | 1.183366416 | 0.407236 | 0.183203 | 1.893672 |
| dioxygenase beta subunit | 1.10494 | 0.959143118 | -0.20415 | 0.18326 | -1.98968 |
| hypothetical protein ABUW_0 | 1.809611 | 0.703921169 | -1.36219 | 0.183637 | -1.98984 |
| general secretion pathway pr | 1.161585 | 0.804321108 | -0.53025 | 0.183712 | -1.97608 |
| metalloendopeptidase | 1.022277 | 1.00028205 | -0.03138 | 0.184141 | -1.86936 |
| beta-hydroxyacyl-(acyl-carrier | 1.001242 | 0.90554221 | -0.14494 | 0.184335 | -1.94834 |
| hypothetical protein ABUW_1 | 1.053228 | 0.97985617 | -0.10418 | 0.184427 | -1.90231 |
| transcriptional regulator, GntR | 1.019681 | 1.084001754 | 0.088249 | 0.185248 | 1.958764 |
| FAD/FMN-binding/pyridine nu | 1.043316 | 0.993948597 | -0.06993 | 0.185471 | -1.72126 |
| transcriptional regulator, LysR | 1.099677 | 0.980373167 | -0.16568 | 0.185596 | -1.83759 |
| pyridine nucleotide transhydr | 0.943089 | 0.897858599 | -0.07091 | 0.185857 | -1.84787 |
| short-chain dehydrogenase/re | 0.931343 | 0.886009622 | -0.07199 | 0.186581 | -1.61278 |
| GTP-binding protein YchF | 0.972649 | 0.90820205 | -0.09891 | 0.186859 | -1.92131 |
| X-Pro aminopeptidase | 0.877069 | 0.955312235 | 0.123283 | 0.186938 | 1.840182 |
| hypothetical protein ABUW_2 | 0.797855 | 1.033154687 | 0.372858 | 0.18696 | 1.961248 |
| protein TolR | 0.958715 | 1.069921375 | 0.158331 | 0.187245 | 1.76806 |
| magnesium and cobalt transp | 0.848026 | 0.947273378 | 0.159673 | 0.18771 | 1.889339 |
| 3-dehydroquinate synthase | 1.026297 | 0.990090971 | -0.05181 | 0.187853 | -1.83269 |
| NADH dehydrogenase I chain | 0.879548 | 0.944197228 | 0.102327 | 0.188036 | 1.694268 |
| NAD dependent epimerase/de | 0.908244 | 0.959673303 | 0.079464 | 0.18811 | 1.60411 |
| cytochrome D ubiquinol oxida | 0.89159 | 1.01511744 | 0.187194 | 0.188112 | 1.609147 |
| acetolactate synthase, large s | 0.949356 | 0.977716596 | 0.042468 | 0.188498 | 1.685391 |
| Transposase (plasmid) | 1.350473 | 0.597926753 | -1.17542 | 0.189366 | -1.9517 |
| peptide deformylase...517 | 0.983343 | 1.005678084 | 0.032401 | 0.189393 | 1.717055 |
| 2-octaprenylphenol hydroxyla | 0.978221 | 0.999508478 | 0.031058 | 0.189772 | 1.677308 |
| short-chain dehydrogenase/re | 1.032414 | 0.972792118 | -0.08582 | 0.189819 | -1.84809 |
| hypothetical protein ABUW_4 | 1.704631 | 0.748479009 | -1.18743 | 0.189986 | -1.91775 |
| phenazine biosynthesis protei | 0.988477 | 0.94791594 | -0.06045 | 0.190151 | -1.70675 |
| hypothetical protein ABUW_3 | 0.971065 | 1.112254622 | 0.195847 | 0.190779 | 1.639153 |
| nucleotide sugar epimerase/d | 1.011019 | 1.059199418 | 0.067164 | 0.190879 | 1.70235 |
| tRNA-i(6)A37 thiotransferase | 1.08176 | 0.953896331 | -0.18148 | 0.190885 | -1.83844 |
| transcriptional regulator, LysR | 0.982157 | 0.961546192 | -0.0306 | 0.192023 | -1.61327 |
| ATP-dependent Clp protease A | 1.066336 | 0.970445364 | -0.13594 | 0.1922 | -1.73003 |
| hypothetical protein ABUW_2 | 1.072566 | 1.033386624 | -0.05369 | 0.192478 | -1.7391 |
| glycerol-3-phosphate dehydro | 0.971058 | 1.09913567 | 0.17874 | 0.192657 | 1.62831 |
| succinyl-CoA:coenzyme A trar | 1.154605 | 1.001234244 | -0.20562 | 0.192865 | -1.83145 |
| fimbrial protein | 1.082212 | 0.971558638 | -0.15561 | 0.192969 | -1.65344 |
| hypothetical protein ABUW_1 | 1.091782 | 0.901894795 | -0.27565 | 0.193039 | -1.76415 |
| putative transcriptional regula | 1.024287 | 0.958960328 | -0.09508 | 0.193357 | -1.67811 |
| endonuclease III | 1.086466 | 1.027871173 | -0.07998 | 0.193975 | -1.83301 |
| hypothetical protein ABUW_0 | 1.077104 | 0.857331027 | -0.32923 | 0.194327 | -1.86936 |
| NADH dehydrogenase I chain | 0.912504 | 0.982204473 | 0.106193 | 0.194575 | 1.662159 |
| alpha/beta fold family hydrola | 0.971401 | 1.086681732 | 0.16179 | 0.19467 | 1.557752 |
| transcriptional regulator, LysR | 1.036863 | 1.097505822 | 0.082003 | 0.194718 | 1.734351 |
| ABC transporter, ATP-binding | 1.024 | 0.965584462 | -0.08474 | 0.195095 | -1.73322 |
| polyprenyl synthetase | 0.868378 | 0.926003266 | 0.092694 | 0.195182 | 1.705114 |
| hypothetical protein ABUW_1 | 1.061603 | 1.240897767 | 0.22514 | 0.195474 | 1.819269 |

| | | | | | |
|---|---|---|---|---|---|
| DNA repair protein RadA | 0.924257 | 1.014508906 | 0.134415 | 0.195694 | 1.897931 |
| putative ATP-dependent DNA | 1.097417 | 1.031468653 | -0.08941 | 0.196179 | -1.68691 |
| methionine aminopeptidase, | 0.926828 | 0.969415102 | 0.064813 | 0.197016 | 1.548154 |
| 4-hydroxybenzoate transport | 0.843624 | 0.97581701 | 0.21001 | 0.197792 | 1.557219 |
| LemA family | 0.983358 | 1.092208796 | 0.15146 | 0.197973 | 1.853157 |
| acetylornithine aminotransfe | 0.962453 | 0.925290612 | -0.05681 | 0.198503 | -1.88744 |
| stearoyl-CoA 9-desaturase | 0.868496 | 0.945693211 | 0.122854 | 0.199949 | 1.782287 |
| hypothetical protein ABUW_2 | 1.0222 | 0.921570025 | -0.14951 | 0.200449 | -1.74161 |
| hypothetical protein ABUW_2 | 0.954226 | 0.991735844 | 0.055624 | 0.201038 | 1.759984 |
| hypothetical protein ABUW_( | 1.057248 | 0.995423647 | -0.08693 | 0.20122 | -1.81383 |
| cyclic AMP phosphodiesteras | 1.059873 | 0.980646758 | -0.11209 | 0.201564 | -1.684 |
| putative flavoprotein | 0.997549 | 1.055981522 | 0.082125 | 0.201607 | 1.786314 |
| oxidoreductase...2378 | 0.962491 | 1.01956974 | 0.083115 | 0.201724 | 1.608691 |
| porin...28 | 0.71365 | 0.802565012 | 0.169402 | 0.202086 | 1.553106 |
| hypothetical protein ABUW_3 | 1.067208 | 1.014857857 | -0.07256 | 0.202739 | -1.69685 |
| phenylacetic acide degradatic | 0.944798 | 1.036245708 | 0.133289 | 0.202867 | 1.566546 |
| RND efflux system, outer mer | 0.971476 | 1.06326646 | 0.130252 | 0.202868 | 1.530446 |
| DNA gyrase, B subunit | 1.013879 | 0.98878025 | -0.03616 | 0.202959 | -1.53656 |
| carbamoyl-phosphate synthas | 0.855272 | 0.837803191 | -0.02977 | 0.203483 | -1.60086 |
| rubredoxin reductase | 0.969938 | 0.907071047 | -0.09668 | 0.203897 | -1.84014 |
| DNA methylase N-4/N-6 | 1.031307 | 0.989848415 | -0.05919 | 0.204113 | -1.66425 |
| transcriptional regulator, Gntl | 1.044954 | 0.945989149 | -0.14354 | 0.204834 | -1.5309 |
| hypothetical protein ABUW_2 | 1.003919 | 0.895955311 | -0.16414 | 0.204959 | -1.62338 |
| UDP-N-acetylglucosamine 1-c | 1.024718 | 0.998777829 | -0.03699 | 0.206 | -1.67516 |
| phenazine biosynthesis protei | 1.376321 | 0.69888106 | -0.9777 | 0.206027 | -1.81443 |
| hypothetical protein ABUW_3 | 0.952062 | 1.011698309 | 0.087652 | 0.206052 | 1.660725 |
| nicotinamide phosphoribosylt | 1.001993 | 0.96685675 | -0.0515 | 0.206074 | -1.80754 |
| OmpA/MotB domain protein. | 1.086668 | 1.04274423 | -0.05953 | 0.206131 | -1.55559 |
| acyl coenzyme A reductase | 1.054748 | 1.00785506 | -0.06561 | 0.206156 | -1.66559 |
| DNA-directed RNA polymeras | 1.016678 | 0.983173328 | -0.04834 | 0.206159 | -1.71184 |
| siderophore biosynthesis prot | 0.630699 | 0.832798178 | 0.401015 | 0.20643 | 1.766661 |
| hypothetical protein ABUW_3 | 0.89149 | 0.97473457 | 0.128791 | 0.206444 | 1.788845 |
| hypothetical protein ABUW_3 | 0.86432 | 0.953140034 | 0.141122 | 0.206827 | 1.744752 |
| RNA polymerase sigma-54 fac | 1.016956 | 0.986556211 | -0.04378 | 0.207023 | -1.64468 |
| melanin biosynthesis protein | 0.949267 | 0.990085184 | 0.060739 | 0.207258 | 1.675685 |
| heat shock protein htpX | 1.000904 | 1.060458928 | 0.083385 | 0.207271 | 1.565612 |
| hypothetical protein ABUW_1 | 1.192854 | 1.305059229 | 0.129698 | 0.207478 | 1.517107 |
| penicillin-binding protein 6 (D | 0.971672 | 0.917096624 | -0.0834 | 0.207666 | -1.74388 |
| homoserine dehydrogenase... | 0.989753 | 1.041302655 | 0.073249 | 0.207674 | 1.501473 |
| catalase...2482 | 0.855986 | 0.999663672 | 0.223855 | 0.209532 | 1.638204 |
| protocatechuate 3,4-dioxyger | 0.966222 | 1.015199782 | 0.071336 | 0.20956 | 1.611085 |
| universal stress protein family | 1.057441 | 0.976596653 | -0.11474 | 0.210138 | -1.74988 |
| peptidoglycan-binding LysM.. | 1.04872 | 0.975666642 | -0.10417 | 0.21062 | -1.75914 |
| metallo-beta-lactamase famil | 0.860074 | 0.970349841 | 0.174044 | 0.211438 | 1.78884 |
| transcriptional regulator, LysF | 0.827539 | 1.016518416 | 0.296738 | 0.211561 | 1.794083 |
| stringent starvation protein A | 1.076152 | 1.140898755 | 0.084289 | 0.211583 | 1.798626 |
| ribosomal RNA large subunit r | 1.039034 | 0.990651088 | -0.06879 | 0.211656 | -1.48563 |

| | | | | | |
|---|---|---|---|---|---|
| hypothetical protein ABUW_3 | 0.972305 | 0.906506988 | -0.10109 | 0.212286 | -1.48647 |
| 2Fe-2S iron-sulfur cluster binc | 1.048829 | 0.857702139 | -0.29023 | 0.213443 | -1.72842 |
| multidrug ABC transporter pe | 2.084595 | 0.642511249 | -1.69797 | 0.213492 | -1.79314 |
| efflux ABC transporter, perme | 1.041701 | 0.923590379 | -0.17362 | 0.21415 | -1.53948 |
| outer membrane lipoprotein | 1.058227 | 1.024466536 | -0.04678 | 0.214457 | -1.49626 |
| hypothetical protein ABUW_( | 0.893496 | 0.948459664 | 0.086126 | 0.214533 | 1.490432 |
| OsmC-like protein | 1.027779 | 0.945629813 | -0.12018 | 0.214881 | -1.77972 |
| NADH pyrophosphatase | 0.978452 | 1.00707703 | 0.041601 | 0.215217 | 1.490117 |
| pyridoxamine 5'-phosphate o: | 0.959291 | 0.892424879 | -0.10424 | 0.215282 | -1.75908 |
| septum site-determining prot | 1.038479 | 1.004042466 | -0.04865 | 0.215516 | -1.65885 |
| phospho-N-acetylmuramoyl-p | 1.028866 | 1.141809064 | 0.150267 | 0.215863 | 1.563785 |
| phosphoribosylaminoimidazo | 0.894247 | 0.930984681 | 0.058084 | 0.216313 | 1.585806 |
| phage/plasmid-related protei | 0.783668 | 0.989315804 | 0.336188 | 0.216538 | 1.689148 |
| UDP-N-acetylglucosamine 2-e | 1.112387 | 0.928152592 | -0.26122 | 0.217359 | -1.72888 |
| hypothetical protein ABUW_3 | 1.026528 | 0.945033227 | -0.11934 | 0.217899 | -1.5681 |
| hypothetical protein ABUW_2 | 0.888777 | 1.121494606 | 0.33553 | 0.218252 | 1.518158 |
| transcriptional regulator, LysF | 1.069991 | 1.028182746 | -0.0575 | 0.218255 | -1.63789 |
| alanine racemase...1968 | 0.997361 | 0.871293196 | -0.19496 | 0.218436 | -1.65356 |
| patatin family phospholipase | 1.044895 | 0.953037805 | -0.13275 | 0.218926 | -1.63414 |
| enoyl-CoA hydratase/isomera | 1.238045 | 1.040244884 | -0.25114 | 0.21938 | -1.75093 |
| hypothetical protein ABUW_3 | 1.03016 | 0.992424871 | -0.05384 | 0.219476 | -1.57317 |
| transcriptional regulator, LysF | 0.985997 | 1.022636847 | 0.052638 | 0.219579 | 1.462412 |
| hypothetical protein ABUW_( | 0.775289 | 0.934048461 | 0.268763 | 0.221323 | 1.670082 |
| hypothetical protein ABUW_2 | 1.003768 | 1.052317215 | 0.068143 | 0.221703 | 1.527066 |
| acetyltransferase, gnat family | 0.902029 | 0.960774712 | 0.091024 | 0.222465 | 1.681444 |
| hypothetical protein ABUW_( | 1.051461 | 0.969831347 | -0.11659 | 0.224147 | -1.55839 |
| hypothetical protein ABUW_2 | 1.018779 | 0.959980163 | -0.08576 | 0.224329 | -1.43614 |
| betaine aldehyde dehydroger | 0.985574 | 0.917006748 | -0.10403 | 0.224907 | -1.54049 |
| pili assembly chaperone | 0.925685 | 1.068600392 | 0.207129 | 0.225076 | 1.601681 |
| hypothetical protein ABUW_2 | 1.455798 | 0.726621435 | -1.00253 | 0.225321 | -1.72134 |
| beta-lactamase domain prote | 1.051113 | 1.005751181 | -0.06364 | 0.225435 | -1.68431 |
| ferric acinetobactin transport | 0.956488 | 0.93736945 | -0.02913 | 0.225505 | -1.60083 |
| adenylate kinase | 1.011907 | 0.950394295 | -0.09048 | 0.226179 | -1.65448 |
| OsmC family protein...1447 | 0.959731 | 1.011046577 | 0.075148 | 0.226291 | 1.545719 |
| acetoin dehydrogenase | 0.89235 | 1.28428252 | 0.525282 | 0.226585 | 1.683542 |
| ATPase | 0.969006 | 0.924324425 | -0.06811 | 0.226825 | -1.42836 |
| transketolase | 1.011607 | 0.957985071 | -0.07857 | 0.226943 | -1.61899 |
| sodium:dicarboxylate sympor | 1.003974 | 1.037290143 | 0.047097 | 0.227687 | 1.503617 |
| aldehyde dehydrogenase...31 | 1.186816 | 0.98989843 | -0.26174 | 0.22849 | -1.60435 |
| UDP-glucose 4-epimerase...23 | 0.995481 | 0.950224117 | -0.06713 | 0.229168 | -1.54923 |
| 1-acyl-sn-glycerol-3-phosphat | 0.908682 | 0.953310928 | 0.069171 | 0.229567 | 1.527879 |
| hypothetical protein ABUW_( | 0.923748 | 1.065924779 | 0.206534 | 0.229841 | 1.41615 |
| malate synthase G | 0.996741 | 0.958485446 | -0.05646 | 0.232378 | -1.64197 |
| GntP family transporter | 0.923759 | 1.020243455 | 0.143325 | 0.232899 | 1.414848 |
| helix-turn-helix domain prote | 1.086687 | 1.044758359 | -0.05677 | 0.233345 | -1.41059 |
| lipoprotein releasing system, . | 0.942257 | 1.020882062 | 0.115623 | 0.233859 | 1.411132 |
| acetyltransferase, gnat family | 1.029391 | 1.00166521 | -0.03939 | 0.234073 | -1.51863 |

| | | | | | |
|---|---|---|---|---|---|
| curved DNA-binding protein | 1.022321 | 1.049256024 | 0.037519 | 0.234147 | 1.422181 |
| hypothetical protein ABUW_3 | 1.013986 | 0.984095846 | -0.04317 | 0.234195 | -1.48433 |
| response regulator | 0.998869 | 0.94823512 | -0.07505 | 0.235066 | -1.52284 |
| thioesterase family protein | 1.126023 | 0.975903665 | -0.20643 | 0.235233 | -1.63426 |
| AAA ATPase family protein | 1.080142 | 1.003639037 | -0.10598 | 0.23525 | -1.53924 |
| transcriptional regulator, ArsR | 1.056877 | 1.117027829 | 0.079858 | 0.235426 | 1.399797 |
| hypothetical protein ABUW_0 | 0.945259 | 0.984374311 | 0.058498 | 0.235911 | 1.573261 |
| putative ATP binding site | 0.999845 | 0.910053687 | -0.13575 | 0.235969 | -1.39422 |
| protein-export chaperone Sec | 0.964835 | 0.917126009 | -0.07316 | 0.236271 | -1.56565 |
| Tol-Pal system beta propeller | 1.00685 | 1.032491609 | 0.036282 | 0.236295 | 1.425939 |
| ribosomal protein L32 | 1.061148 | 0.936304183 | -0.18058 | 0.236949 | -1.66098 |
| arsenate reductase...113 | 0.728609 | 1.042554861 | 0.516907 | 0.237385 | 1.561215 |
| excinuclease ABC, A subunit | 0.958797 | 1.004385541 | 0.067016 | 0.237675 | 1.560631 |
| shikimate kinase...820 | 0.97762 | 1.048483897 | 0.100959 | 0.237788 | 1.387683 |
| hypothetical protein ABUW_1 | 1.229697 | 0.897602893 | -0.45415 | 0.238398 | -1.55667 |
| transcriptional regulator, ArsR | 1.045238 | 0.97488472 | -0.10053 | 0.238564 | -1.57205 |
| hypothetical protein ABUW_1 | 1.071182 | 1.020831132 | -0.06946 | 0.238573 | -1.63932 |
| beta-ketoadipyl CoA thiolase. | 0.913491 | 0.963312639 | 0.076613 | 0.238718 | 1.509874 |
| hypothetical protein ABUW_2 | 1.05514 | 1.188454395 | 0.171652 | 0.239804 | 1.505423 |
| surface antigen | 0.891602 | 0.942643612 | 0.080313 | 0.241104 | 1.392817 |
| hypothetical protein ABUW_1 | 1.136269 | 0.926431131 | -0.29455 | 0.241146 | -1.60917 |
| site-specific recombinase, pha | 0.941077 | 1.050634341 | 0.158876 | 0.241877 | 1.381193 |
| pirin domain protein...1290 | 0.997168 | 1.075242158 | 0.108753 | 0.242585 | 1.448523 |
| protocatechuate 3,4-dioxygen | 0.949369 | 0.988772265 | 0.05867 | 0.243987 | 1.447907 |
| NAD(P)H oxidoreductase | 1.101815 | 1.015003939 | -0.1184 | 0.244169 | -1.59083 |
| transporter, LysE family | 0.747656 | 1.021162342 | 0.449766 | 0.244254 | 1.611753 |
| hypothetical protein ABUW_0 | 0.9236 | 0.979266712 | 0.084433 | 0.24473 | 1.4182 |
| hypothetical protein ABUW_2 | 1.082266 | 0.888496022 | -0.28462 | 0.24495 | -1.61286 |
| hypothetical protein ABUW_1 | 0.977569 | 0.949230499 | -0.04244 | 0.245239 | -1.49089 |
| peptidyl-prolyl cis-trans isome | 0.977512 | 1.01884783 | 0.059752 | 0.245853 | 1.622961 |
| hypothetical protein ABUW_2 | 1.155474 | 1.053171435 | -0.13374 | 0.247139 | -1.48599 |
| hydrolase, NUDIX family | 0.928786 | 1.069482195 | 0.203495 | 0.247163 | 1.462825 |
| hypothetical protein ABUW_1 | 1.021419 | 1.147759861 | 0.168246 | 0.247328 | 1.601359 |
| hypothetical protein ABUW_0 | 1.009334 | 0.976348023 | -0.04794 | 0.247753 | -1.59693 |
| permease...1176 | 0.967926 | 0.999414168 | 0.046186 | 0.247853 | 1.449058 |
| ATPase, AAA family | 1.019447 | 0.99495864 | -0.03508 | 0.248812 | -1.41113 |
| hypothetical protein ABUW_2 | 0.670662 | 1.075266351 | 0.681037 | 0.249296 | 1.598142 |
| entericidin EcnAB | 1.085927 | 0.959948313 | -0.1779 | 0.249424 | -1.49013 |
| peptidyl-prolyl cis-trans isome | 1.030391 | 0.963556738 | -0.09675 | 0.250221 | -1.50993 |
| hypothetical protein ABUW_0 | 0.948379 | 0.995547071 | 0.070025 | 0.250359 | 1.565269 |
| hypothetical protein ABUW_3 | 0.986577 | 0.920638094 | -0.0998 | 0.250462 | -1.37161 |
| two component signal transdu | 0.943022 | 1.046368227 | 0.150028 | 0.250694 | 1.432587 |
| UDP-2,3-diacylglucosamine hy | 1.258769 | 1.150734238 | -0.12946 | 0.251813 | -1.3496 |
| DNA polymerase III subunit al | 0.942613 | 0.972262671 | 0.044681 | 0.252235 | 1.378789 |
| aminotransferase | 0.934842 | 0.986658938 | 0.077829 | 0.252737 | 1.577961 |
| 5-formyltetrahydrofolate cycl | 0.881137 | 0.921342854 | 0.064371 | 0.252776 | 1.557128 |
| shikimate transporter | 0.749687 | 0.885026491 | 0.239431 | 0.253624 | 1.376954 |

| | | | | | |
|---|---|---|---|---|---|
| 2-hydroxy-3-oxopropionate re | 1.064145 | 0.973258405 | -0.1288 | 0.253814 | -1.5439 |
| alternative sigma factor RpoH | 1.000977 | 1.066174644 | 0.091035 | 0.253938 | 1.404633 |
| coenzyme PQQ biosynthesis p | 1.059647 | 1.023153066 | -0.05056 | 0.255191 | -1.36932 |
| transcriptional regulator, LysR | 0.941195 | 1.004371682 | 0.093728 | 0.255389 | 1.361833 |
| ATP-dependent DNA helicase | 0.988897 | 0.904171889 | -0.12922 | 0.25558 | -1.54235 |
| transcriptional regulator, LysR | 1.019288 | 1.096431 | 0.105253 | 0.255939 | 1.415833 |
| metallo-beta-lactamase family | 0.96627 | 0.942929896 | -0.03528 | 0.255988 | -1.47595 |
| hypothetical protein ABUW_0 | 0.94198 | 1.088769425 | 0.20893 | 0.256277 | 1.518236 |
| outer membrane lipoprotein | 1.090837 | 1.050569679 | -0.05426 | 0.256953 | -1.34272 |
| hypothetical protein ABUW_0 | 0.992004 | 1.046424956 | 0.077051 | 0.257016 | 1.332539 |
| trans-hexaprenyltranstransfer | 0.996474 | 0.95392891 | -0.06295 | 0.257619 | -1.49896 |
| hypothetical protein ABUW_1 | 1.036008 | 0.994278244 | -0.05931 | 0.25834 | -1.34599 |
| putative DNA helicase | 0.946348 | 0.894579973 | -0.08116 | 0.259184 | -1.46062 |
| phenylacetate-CoA ligase | 0.898457 | 1.035325177 | 0.204563 | 0.259377 | 1.446704 |
| chorismate mutase family pro | 0.886812 | 1.045542394 | 0.237551 | 0.260408 | 1.430995 |
| 23S rRNA (uracil-5-)-methyltra | 0.910316 | 0.968401762 | 0.089239 | 0.261824 | 1.323367 |
| DNA gyrase, A subunit | 1.002608 | 0.989525575 | -0.01895 | 0.262244 | -1.33018 |
| acyl-CoA synthase...1213 | 1.005477 | 0.905429153 | -0.15121 | 0.26365 | -1.32215 |
| hypothetical protein ABUW_0 | 0.994606 | 1.06184586 | 0.094377 | 0.26369 | 1.361972 |
| thioesterase superfamily prot | 1.075214 | 1.21400598 | 0.175152 | 0.264397 | 1.343824 |
| ABC transporter, ATP-binding | 1.265129 | 1.112287193 | -0.18576 | 0.26448 | -1.39492 |
| polysaccharide export protein | 0.990067 | 0.960377575 | -0.04392 | 0.264553 | -1.31208 |
| adenylosuccinate synthetase | 0.97859 | 0.943767425 | -0.05227 | 0.264594 | -1.49658 |
| cell division protein FtsH | 0.950882 | 0.996404785 | 0.067465 | 0.264776 | 1.512722 |
| methenyltetrahydrofolate cyc | 0.949914 | 0.976382307 | 0.03965 | 0.26498 | 1.298362 |
| hypothetical protein ABUW_1 | 1.088382 | 1.020869725 | -0.09239 | 0.266997 | -1.30056 |
| ATP-dependent Clp protease a | 1.082352 | 0.996049284 | -0.11988 | 0.267452 | -1.40357 |
| transporter, major facilitator f | 0.963128 | 1.081282238 | 0.166943 | 0.268149 | 1.477897 |
| phosphohistidine phosphatas | 0.964646 | 1.037291852 | 0.10475 | 0.268439 | 1.427568 |
| hypothetical protein ABUW_2 | 0.972248 | 0.911706228 | -0.09276 | 0.268616 | -1.48544 |
| exonuclease V gamma chain | 0.880238 | 0.795742689 | -0.14559 | 0.269234 | -1.2885 |
| transcriptional regulator, TetR | 0.993005 | 0.952286267 | -0.06041 | 0.269284 | -1.28267 |
| hypothetical protein ABUW_0 | 1.07095 | 0.862915611 | -0.3116 | 0.269386 | -1.32574 |
| phenylacetate-CoA oxygenase | 0.897409 | 1.107514904 | 0.303488 | 0.26949 | 1.497083 |
| hypothetical protein ABUW_3 | 0.98668 | 0.958667347 | -0.04155 | 0.269647 | -1.40942 |
| hypothetical protein ABUW_2 | 0.940941 | 1.068360691 | 0.183223 | 0.269857 | 1.397194 |
| hypothetical protein ABUW_3 | 0.85907 | 1.125684006 | 0.389953 | 0.270254 | 1.492205 |
| phenylacetate-CoA oxygenase | 0.847207 | 1.095808336 | 0.37121 | 0.270312 | 1.50103 |
| guanine deaminase...565 | 0.985661 | 0.861032909 | -0.19502 | 0.270454 | -1.41006 |
| chorismate synthase | 0.981505 | 1.032356726 | 0.072874 | 0.270512 | 1.339053 |
| transcription elongation facto | 1.089578 | 1.036496644 | -0.07205 | 0.270719 | -1.29512 |
| tetraacyldisaccharide 4'-kinas | 1.024887 | 0.968053122 | -0.08231 | 0.271176 | -1.47881 |
| ATP phosphoribosyltransferas | 0.970583 | 0.926010584 | -0.06782 | 0.271181 | -1.47796 |
| acetyl-CoA acetyltransferase.. | 1.034666 | 0.97658377 | -0.08335 | 0.271323 | -1.47861 |
| transcriptional regulator, Fur f | 1.019184 | 0.997371987 | -0.03121 | 0.271733 | -1.407 |
| glutamate-ammonia-ligase ad | 0.983132 | 0.962051874 | -0.03127 | 0.271832 | -1.28385 |
| type IV pilus biogenesis/stabil | 1.041203 | 1.020737674 | -0.02864 | 0.272262 | -1.33858 |

| | | | | | |
|---|---|---|---|---|---|
| hypothetical protein ABUW_1 | 0.937814 | 1.034676209 | 0.141806 | 0.272366 | 1.299242 |
| universal stress protein | 1.029707 | 0.980958685 | -0.06997 | 0.273256 | -1.27938 |
| hydroxyethylthiazole kinase | 0.975327 | 0.915083028 | -0.09198 | 0.273366 | -1.28405 |
| hypothetical protein ABUW_3 | 0.985343 | 1.006126473 | 0.030114 | 0.274241 | 1.291881 |
| hypothetical protein ABUW_3 | 0.947039 | 0.921442409 | -0.03953 | 0.274498 | -1.46527 |
| tyrosine-protein kinase ptk | 0.989479 | 1.012968622 | 0.033848 | 0.274547 | 1.485006 |
| alpha/beta hydrolase...2175 | 1.034529 | 1.003941272 | -0.0433 | 0.275138 | -1.30381 |
| hypothetical protein ABUW_1 | 1.081162 | 1.309610219 | 0.276555 | 0.275149 | 1.415796 |
| transposition helper protein ( | 0.993329 | 1.032427162 | 0.055697 | 0.275231 | 1.286007 |
| transcriptional regulator, XRE | 1.066747 | 0.933255507 | -0.19287 | 0.275259 | -1.3318 |
| hypothetical protein ABUW_( | 0.807227 | 0.938879747 | 0.217967 | 0.275339 | 1.47475 |
| hypothetical protein ABUW_3 | 1.042318 | 0.878072905 | -0.24738 | 0.275491 | -1.34608 |
| catalase...1924 | 0.994786 | 1.044593682 | 0.070484 | 0.276342 | 1.400706 |
| signal recognition particle pro | 0.993795 | 0.956532399 | -0.05513 | 0.276588 | -1.28539 |
| KGG domain-containing prote | 0.99181 | 1.026833993 | 0.050068 | 0.277697 | 1.262466 |
| non-ribosomal peptide synthe | 0.937254 | 0.980254218 | 0.064716 | 0.278128 | 1.263981 |
| hypothetical protein ABUW_( | 0.96403 | 0.993506757 | 0.043452 | 0.278759 | 1.285244 |
| type IV pilus response regulat | 1.001577 | 0.963479737 | -0.05595 | 0.279367 | -1.32689 |
| hypothetical protein ABUW_( | 0.982051 | 0.901211111 | -0.12393 | 0.279634 | -1.25983 |
| hypothetical protein ABUW_( | 0.963575 | 0.929436368 | -0.05204 | 0.279763 | -1.25657 |
| RND family drug transporter | 0.825933 | 0.890547279 | 0.108668 | 0.280578 | 1.281197 |
| alpha/beta hydrolase fold pro | 0.899062 | 0.845607416 | -0.08843 | 0.280686 | -1.24625 |
| hypothetical protein ABUW_3 | 0.967292 | 0.88505099 | -0.12819 | 0.28109 | -1.27175 |
| hypothetical protein ABUW_3 | 1.082525 | 1.171798212 | 0.114324 | 0.281233 | 1.347483 |
| glutamyl-tRNA synthetase...2 | 0.956258 | 0.98985507 | 0.049818 | 0.281547 | 1.384712 |
| urease accessory protein F | 0.848452 | 1.062197677 | 0.324147 | 0.281713 | 1.378556 |
| 3-hydroxyacyl-CoA dehydroge | 0.670161 | 1.273719645 | 0.926468 | 0.281875 | 1.449616 |
| hydroxymethylglutaryl-CoA ly | 0.985358 | 0.956244483 | -0.04327 | 0.281912 | -1.27971 |
| amino-acid N-acetyltransfera | 0.941317 | 0.986895072 | 0.068215 | 0.28206 | 1.439992 |
| competence/damage-inducib | 0.954054 | 0.999366581 | 0.066944 | 0.282635 | 1.287498 |
| ferrochelatase | 0.995247 | 1.046374742 | 0.072273 | 0.282682 | 1.241236 |
| hypothetical protein ABUW_2 | 1.151779 | 0.931215779 | -0.30668 | 0.282996 | -1.4094 |
| glutamate--cysteine ligase | 1.065557 | 0.990689976 | -0.1051 | 0.283032 | -1.31075 |
| ABC transporter | 1.057176 | 0.987663179 | -0.09812 | 0.283225 | -1.30443 |
| guanine deaminase...1601 | 0.926895 | 0.968102273 | 0.062754 | 0.283468 | 1.252062 |
| tRNA modification GTPase Tr | 0.97481 | 0.910561212 | -0.09837 | 0.283571 | -1.34565 |
| glutaredoxin | 0.979039 | 1.021783477 | 0.061651 | 0.284897 | 1.37508 |
| transcriptional regulator, Lys | 1.09608 | 1.033908138 | -0.08424 | 0.285062 | -1.34348 |
| AdeS | 1.015205 | 1.073353719 | 0.080355 | 0.285097 | 1.422821 |
| diaminopimelate decarboxyla | 0.972457 | 0.997681041 | 0.036945 | 0.28521 | 1.327863 |
| pantetheine-phosphate aden | 1.024675 | 0.946095077 | -0.11511 | 0.285433 | -1.3067 |
| hypothetical protein ABUW_( | 1.074237 | 1.014641412 | -0.08234 | 0.285525 | -1.41617 |
| nitrate transport ATP-binding | 1.074654 | 0.970706015 | -0.14677 | 0.286274 | -1.23261 |
| nicotinate-nucleotide--dimetl | 0.957882 | 0.973478053 | 0.023301 | 0.287007 | 1.249734 |
| ben operon transcriptional re | 1.016501 | 1.05924924 | 0.059431 | 0.287803 | 1.264135 |
| hypothetical protein ABUW_1 | 1.048038 | 1.133656911 | 0.113293 | 0.287928 | 1.366759 |
| phosphopyruvate hydratase ( | 1.007891 | 0.928789175 | -0.11792 | 0.288064 | -1.42504 |

| | | | | | |
|---|---|---|---|---|---|
| cell division protein FtsZ | 0.979863 | 0.948209423 | -0.04737 | 0.288131 | -1.29545 |
| ATP synthase F0, B subunit | 0.988179 | 0.969619044 | -0.02735 | 0.288151 | -1.40517 |
| D-serine/D-alanine/glycine tra | 1.197565 | 1.035871418 | -0.20926 | 0.288951 | -1.2284 |
| CTP synthase | 0.952097 | 0.975664995 | 0.035277 | 0.289668 | 1.235651 |
| hypothetical protein ABUW_2 | 0.886632 | 0.8590757 | -0.04555 | 0.290109 | -1.253 |
| Holliday junction DNA helicase | 0.974996 | 1.013367821 | 0.05569 | 0.29021 | 1.316348 |
| aldo/keto reductase...2097 | 0.960398 | 0.931595864 | -0.04393 | 0.29058 | -1.24859 |
| hypothetical protein ABUW_3 | 1.042129 | 0.974098364 | -0.09739 | 0.291835 | -1.33986 |
| transcriptional regulator EstR, | 0.980168 | 0.956170515 | -0.03576 | 0.29216 | -1.3995 |
| DNA cytosine methyltransfera | 0.971074 | 0.916846829 | -0.0829 | 0.292761 | -1.24502 |
| OmpW family protein...1026 | 0.725247 | 0.748256478 | 0.04506 | 0.292913 | 1.33219 |
| hypothetical protein ABUW_2 | 0.965945 | 1.026462841 | 0.087668 | 0.293124 | 1.36377 |
| ABC transporter, periplasmic | 1.065 | 1.009720994 | -0.0769 | 0.295081 | -1.25118 |
| redox-sensitive transcriptiona | 0.946656 | 0.898551146 | -0.07524 | 0.295622 | -1.25932 |
| phosphate ABC transporter, A | 0.891057 | 1.014112534 | 0.186627 | 0.295839 | 1.251898 |
| zinc-binding alcohol dehydrog | 0.999709 | 1.143741895 | 0.194181 | 0.296206 | 1.361504 |
| quinate/shikimate dehydroge | 0.994692 | 1.078309712 | 0.11645 | 0.29643 | 1.2777 |
| transcriptional regulator, TetF | 0.894907 | 0.940734566 | 0.072051 | 0.296788 | 1.229587 |
| hypothetical protein ABUW_1 | 1.099006 | 1.060801918 | -0.05104 | 0.296826 | -1.36437 |
| 2,3-dihydro-2,3-dihydroxyber | 1.068449 | 1.015687671 | -0.07306 | 0.296852 | -1.35747 |
| glutamyl-tRNA reductase | 0.943732 | 0.983819171 | 0.060016 | 0.297601 | 1.382239 |
| hypothetical protein ABUW_1 | 0.932234 | 0.991131027 | 0.088384 | 0.298434 | 1.318047 |
| septum formation initiator | 1.173882 | 1.287794375 | 0.133614 | 0.298736 | 1.371271 |
| band 7 protein | 0.947679 | 0.959759581 | 0.018274 | 0.299202 | 1.217615 |
| ggdef domain/eal domain pro | 0.946036 | 0.90534216 | -0.06343 | 0.299337 | -1.19246 |
| transcriptional regulator, Gntl | 1.011517 | 0.980969399 | -0.04424 | 0.299352 | -1.27008 |
| methylmalonate-semialdehyd | 0.981778 | 0.958411266 | -0.03475 | 0.299511 | -1.19878 |
| hypothetical protein ABUW_2 | 1.126003 | 0.982363736 | -0.19688 | 0.299878 | -1.36 |
| iron-sulfur cluster-binding pro | 0.979776 | 0.828996614 | -0.24109 | 0.300103 | -1.28861 |
| hypothetical protein ABUW_2 | 0.997052 | 0.944846684 | -0.07759 | 0.300535 | -1.36649 |
| RNA methyltransferase, TrmH | 0.937436 | 0.962447239 | 0.037987 | 0.300685 | 1.202607 |
| short chain dehydrogenase...4 | 1.023345 | 1.143786777 | 0.160525 | 0.300946 | 1.312768 |
| type VI secretion protein IcmF | 0.994688 | 1.020152254 | 0.036469 | 0.301162 | 1.329625 |
| hypothetical protein ABUW_0 | 1.021615 | 0.962955893 | -0.08531 | 0.301371 | -1.217 |
| signal recognition particle-doc | 1.006004 | 1.048529812 | 0.059732 | 0.302199 | 1.2745 |
| transcriptional regulator, LysF | 0.954714 | 1.009550856 | 0.080574 | 0.302549 | 1.256115 |
| allantoicase | 0.962343 | 0.920142567 | -0.06469 | 0.302583 | -1.25944 |
| cold-shock DNA-binding doma | 0.647389 | 1.490821637 | 1.203403 | 0.302773 | 1.371198 |
| AcnD-accessory protein PrpF | 1.00849 | 0.993080078 | -0.02222 | 0.302878 | -1.22349 |
| hypothetical protein ABUW_1 | 1.053137 | 0.951676491 | -0.14615 | 0.303313 | -1.18636 |
| thiol:disulfide interchange pro | 0.995114 | 1.019660898 | 0.035156 | 0.303883 | 1.318108 |
| acyl-CoA dehydrogenase...193 | 1.02774 | 1.000314012 | -0.03902 | 0.304134 | -1.32513 |
| glycolate/propanediol utilizat | 1.024117 | 0.988466615 | -0.05112 | 0.304195 | -1.33512 |
| multidrug resistance protein V | 0.992483 | 1.0535629 | 0.086162 | 0.304258 | 1.353396 |
| UDP-N-acetylmuramyl-tripept | 0.951762 | 0.983622913 | 0.047504 | 0.304466 | 1.223441 |
| phosphomannomutase...2141 | 0.965412 | 0.977612388 | 0.018117 | 0.304645 | 1.239747 |
| khg/kdpg aldolase | 0.973483 | 0.932630592 | -0.06185 | 0.305979 | -1.28788 |

| | | | | | |
|---|---|---|---|---|---|
| alginate biosynthesis protein | 0.89285 | 1.056851013 | 0.243283 | 0.305996 | 1.322001 |
| alcohol dehydrogenase, zinc-b | 0.938088 | 0.904757113 | -0.05219 | 0.306539 | -1.31785 |
| glycerophosphoryl diester pho | 0.999646 | 0.971695719 | -0.04091 | 0.307232 | -1.17184 |
| putative esterase | 1.126231 | 0.960308602 | -0.22993 | 0.307951 | -1.16787 |
| lipoic acid synthetase...155 | 1.153219 | 0.881021644 | -0.38842 | 0.308232 | -1.28062 |
| SPOUT methyltransferase | 1.026534 | 0.982593439 | -0.06312 | 0.30833 | -1.22134 |
| prolyl-tRNA synthetase | 0.958513 | 0.992244688 | 0.049897 | 0.308719 | 1.1942 |
| triose-phosphate isomerase | 1.018117 | 1.00222544 | -0.0227 | 0.308742 | -1.16505 |
| bacterioferritin...2280 | 0.99948 | 0.943954642 | -0.08246 | 0.309305 | -1.32428 |
| acetoin:2,6-dichlorophenolinc | 0.922722 | 1.1668402 | 0.33864 | 0.309546 | 1.343967 |
| Transcriptional repressor prot | 1.057223 | 1.032872559 | -0.03362 | 0.309818 | -1.25271 |
| pirin domain protein...1396 | 0.970012 | 0.992359703 | 0.03286 | 0.309846 | 1.191265 |
| aspartate aminotransferase | 0.948551 | 0.962935821 | 0.021714 | 0.310879 | 1.182883 |
| long-chain fatty acid transpor | 1.127265 | 0.940959417 | -0.26062 | 0.31175 | -1.27801 |
| hypothetical protein ABUW_3 | 1.044508 | 1.127604185 | 0.110438 | 0.311902 | 1.307872 |
| aldehyde dehydrogenase...24 | 1.046713 | 0.959341254 | -0.12575 | 0.312141 | -1.18075 |
| DnaK suppressor protein | 1.021848 | 0.968195462 | -0.07781 | 0.312365 | -1.28846 |
| transcriptional regulator, LysR | 0.952273 | 0.984117204 | 0.047455 | 0.312461 | 1.332388 |
| hypothetical protein ABUW_1 | 0.981968 | 1.091056757 | 0.151979 | 0.312823 | 1.154422 |
| UDP-3-O-[3-hydroxymyristoyl | 0.957738 | 0.997450369 | 0.058614 | 0.31354 | 1.284999 |
| DnaA family protein | 0.958875 | 0.983761871 | 0.036966 | 0.314403 | 1.246265 |
| glutamine amidotransferase, | 0.970571 | 0.984392866 | 0.020401 | 0.315465 | 1.194459 |
| malate dehydrogenase...1909 | 0.933326 | 0.975737583 | 0.064111 | 0.315926 | 1.267396 |
| hypothetical protein ABUW_0 | 0.95888 | 1.082691574 | 0.1752 | 0.316336 | 1.212094 |
| two-component system senso | 0.941783 | 1.049489319 | 0.156221 | 0.316676 | 1.267616 |
| outer membrane protein...21 | 1.002659 | 0.975123932 | -0.04017 | 0.317068 | -1.18999 |
| MutT/NUDIX family protein | 0.798354 | 0.759829821 | -0.07135 | 0.317372 | -1.31227 |
| hypothetical protein ABUW_2 | 0.922653 | 0.996227074 | 0.110687 | 0.318995 | 1.313576 |
| cytochrome O ubiquinol oxida | 0.976038 | 0.748318563 | -0.38328 | 0.319006 | -1.30481 |
| alpha/beta hydrolase fold pro | 0.926029 | 0.968733689 | 0.065042 | 0.320546 | 1.255603 |
| hypothetical protein ABUW_1 | 0.965904 | 1.040314777 | 0.107069 | 0.320632 | 1.214471 |
| hypothetical protein ABUW_4 | 1.031251 | 0.97849078 | -0.07576 | 0.320926 | -1.17358 |
| hypothetical protein ABUW_0 | 1.11878 | 0.999583615 | -0.16253 | 0.321762 | -1.28401 |
| transcriptional regulator, TetR | 1.024338 | 0.988500576 | -0.05138 | 0.322028 | -1.1729 |
| thioredoxin...1316 | 1.009223 | 0.959917313 | -0.07226 | 0.322322 | -1.27098 |
| molybdopterin oxidoreductas | 0.76285 | 1.03206671 | 0.436065 | 0.322894 | 1.222541 |
| transcriptional regulator TetR | 1.175362 | 1.097636644 | -0.0987 | 0.323668 | -1.29648 |
| acyl carrier protein...44 | 0.972933 | 1.272437265 | 0.387182 | 0.323944 | 1.287414 |
| phospholipase C, phosphocho | 1.050852 | 0.902212639 | -0.22002 | 0.324832 | -1.12724 |
| xanthine/uracil permease | 1.105956 | 1.230839199 | 0.154348 | 0.324995 | 1.140326 |
| phage-related terminase | 1.012621 | 1.123004358 | 0.149269 | 0.325121 | 1.133659 |
| short chain dehydrogenase...6 | 0.961669 | 1.064145192 | 0.146082 | 0.325395 | 1.238297 |
| two-component system senso | 1.001345 | 1.054380508 | 0.074456 | 0.325452 | 1.163919 |
| multidrug efflux protein AdeK | 1.010304 | 1.026911511 | 0.023522 | 0.325608 | 1.123742 |
| RNA polymerase sigma-70 fac | 0.984641 | 1.020118929 | 0.051067 | 0.325719 | 1.255995 |
| catalase/peroxidase HPI | 0.943161 | 0.969553311 | 0.039816 | 0.325935 | 1.118923 |
| hypothetical protein ABUW_2 | 0.968404 | 0.946006603 | -0.03376 | 0.326343 | -1.17098 |

| | | | | | |
|---|---|---|---|---|---|
| hypothetical protein ABUW_3 | 1.043024 | 1.013051013 | -0.04207 | 0.326699 | -1.14653 |
| hypothetical protein ABUW_4 | 0.962006 | 0.905558108 | -0.08724 | 0.327299 | -1.15158 |
| succinate-semialdehyde dehy | 1.014993 | 0.963137656 | -0.07566 | 0.327714 | -1.13321 |
| magnesium and cobalt efflux | 0.929724 | 0.895680474 | -0.05382 | 0.328594 | -1.22662 |
| ribonucleoside-diphosphate r | 1.059536 | 1.019245848 | -0.05593 | 0.328616 | -1.27202 |
| secretion chaperone | 1.02483 | 0.965391951 | -0.0862 | 0.328668 | -1.19248 |
| short-chain dehydrogenase/re | 0.890481 | 0.788886413 | -0.17477 | 0.329304 | -1.22095 |
| hypothetical protein ABUW_2 | 1.059578 | 0.968155588 | -0.13018 | 0.329704 | -1.189 |
| hypothetical protein ABUW_0 | 0.983336 | 0.936104094 | -0.07102 | 0.332052 | -1.19663 |
| two-component system senso | 0.923907 | 0.97177482 | 0.072875 | 0.332296 | 1.252707 |
| two-component system respo | 1.004377 | 0.98222453 | -0.03218 | 0.332704 | -1.10642 |
| DNA polymerase III, delta sub | 0.951843 | 0.901486763 | -0.07842 | 0.332886 | -1.20373 |
| hypothetical protein ABUW_0 | 0.432131 | 2.099173766 | 2.280282 | 0.333204 | 1.265036 |
| DNA repair protein RecO | 0.934059 | 0.898782188 | -0.05554 | 0.334425 | -1.16073 |
| hypothetical protein ABUW_0 | 1.026395 | 1.00371148 | -0.03224 | 0.334633 | -1.13504 |
| hypothetical protein ABUW_2 | 0.9585 | 1.010167473 | 0.075744 | 0.335377 | 1.237772 |
| phage replication protein | 0.938233 | 1.039031349 | 0.147222 | 0.335387 | 1.114518 |
| transcriptional regulator, Mer | 0.898801 | 0.982649437 | 0.128675 | 0.33559 | 1.211532 |
| succinylglutamate desuccinyla | 1.03159 | 1.007386346 | -0.03425 | 0.335676 | -1.12271 |
| outer membrane protein CarO | 0.974071 | 1.060177974 | 0.122208 | 0.336154 | 1.094155 |
| dihydroxy-acid dehydratase... | 0.98845 | 0.930409262 | -0.0873 | 0.33667 | -1.20603 |
| sensor kinase CusS | 0.945196 | 0.988655705 | 0.064855 | 0.336925 | 1.095057 |
| 3-dehydroquinate dehydratas | 0.980444 | 0.96079789 | -0.0292 | 0.337523 | -1.1101 |
| hypothetical protein ABUW_2 | 0.942137 | 0.91911977 | -0.03568 | 0.337546 | -1.13891 |
| transcriptional regulator, Mar | 1.055846 | 0.983180572 | -0.10287 | 0.33816 | -1.20555 |
| ferredoxin-1 | 0.919193 | 1.035250197 | 0.17154 | 0.338509 | 1.159777 |
| hydrolase...287 | 1.003137 | 0.953089723 | -0.07383 | 0.338642 | -1.12516 |
| peptidyl-prolyl cis-trans isome | 1.059987 | 1.017167804 | -0.05949 | 0.338819 | -1.13697 |
| copper resistance D | 0.69177 | 1.746894011 | 1.336427 | 0.338935 | 1.245186 |
| hypothetical protein ABUW_0 | 1.017534 | 1.113224584 | 0.129667 | 0.339018 | 1.227746 |
| pH adaptation potassium efflu | 0.905527 | 0.87350458 | -0.05194 | 0.339405 | -1.0886 |
| phospholipase | 0.956759 | 0.938216977 | -0.02823 | 0.339441 | -1.22311 |
| D-and L-methionine transport | 0.972316 | 0.843631567 | -0.20481 | 0.33959 | -1.17893 |
| family 1 glycosyl transferase.. | 0.668474 | 0.715991885 | 0.099071 | 0.340038 | 1.19398 |
| glutamine-fructose-6-phospha | 0.969993 | 0.995790253 | 0.037867 | 0.340151 | 1.090735 |
| major capsid protein | 0.942221 | 1.004400928 | 0.092197 | 0.340614 | 1.115314 |
| dephospho-CoA kinase | 0.958325 | 0.927108895 | -0.04778 | 0.340823 | -1.08026 |
| glutaryl-CoA dehydrogenase | 0.937998 | 0.985590021 | 0.071403 | 0.342501 | 1.076281 |
| phosphate regulon sensor kin | 0.975379 | 0.99896555 | 0.034472 | 0.342519 | 1.122865 |
| GMP synthase | 1.036166 | 0.970133024 | -0.095 | 0.343111 | -1.17271 |
| 3-oxoadipate enol-lactonase.. | 0.893564 | 0.924083221 | 0.048452 | 0.343746 | 1.104289 |
| dihydrolipoamide dehydroger | 1.001084 | 0.909563628 | -0.13832 | 0.344833 | -1.19861 |
| hypothetical protein ABUW_2 | 0.286708 | 2.162910518 | 2.915318 | 0.344897 | 1.226118 |
| TrwC protein (plasmid) | 0.767476 | 0.892448403 | 0.217648 | 0.345082 | 1.075995 |
| alkaline lipase | 1.055089 | 1.157793137 | 0.134013 | 0.345821 | 1.113026 |
| transthyretin | 0.801833 | 0.90195856 | 0.169759 | 0.346932 | 1.214459 |
| general secretion pathway pr | 0.99211 | 1.1035845 | 0.153625 | 0.347312 | 1.190447 |

| | | | | | |
|---|---|---|---|---|---|
| shikimate kinase...825 | 0.926907 | 1.026472936 | 0.147199 | 0.347599 | 1.07852 |
| hypothetical protein ABUW_2 | 0.95072 | 0.927626153 | -0.03548 | 0.348136 | -1.07855 |
| tRNA(Ile)-lysidine synthase | 0.93487 | 0.876583357 | -0.09287 | 0.348205 | -1.08822 |
| transcriptional regulator, DUF | 0.49312 | 2.340492234 | 2.2468 | 0.348455 | 1.214573 |
| hypothetical protein ABUW_1 | 0.949376 | 0.840053923 | -0.1765 | 0.351369 | -1.18506 |
| NADH dehydrogenase I chain | 0.961839 | 0.975842949 | 0.020853 | 0.351557 | 1.113031 |
| inner membrane symporter Y | 0.948276 | 1.000635099 | 0.077538 | 0.351601 | 1.10074 |
| ribonucleoside-diphosphate r | 1.044135 | 1.002099026 | -0.05928 | 0.352234 | -1.0575 |
| NAD-dependent deacetylase r | 1.010378 | 0.918605948 | -0.13738 | 0.352388 | -1.08065 |
| type VI secretion protein, fam | 1.047542 | 1.020250488 | -0.03808 | 0.352513 | -1.19887 |
| hypothetical protein ABUW_1 | 0.949764 | 0.990287691 | 0.060279 | 0.353194 | 1.051695 |
| intracellular protease, PfpI far | 1.00836 | 0.960211601 | -0.07059 | 0.353915 | -1.19443 |
| short chain dehydrogenase...1 | 1.107316 | 0.899941469 | -0.29916 | 0.354756 | -1.18861 |
| thiamine biosynthesis protein | 0.9047 | 0.956447249 | 0.080246 | 0.355765 | 1.059915 |
| phosphatase domain-containi | 0.855965 | 0.928318577 | 0.117069 | 0.35603 | 1.173751 |
| hypothetical protein ABUW_5 | 1.001971 | 1.030637669 | 0.040697 | 0.356493 | 1.183149 |
| cold-shock DNA-binding doma | 0.399187 | 1.715052253 | 2.103116 | 0.356655 | 1.188376 |
| cytochrome b562 | 0.752825 | 1.049417919 | 0.479202 | 0.356964 | 1.161859 |
| hypothetical protein ABUW_0 | 1.016693 | 0.973073824 | -0.06326 | 0.35752 | -1.11002 |
| uroporphyrinogen decarboxyl | 0.945034 | 0.978386157 | 0.050038 | 0.357633 | 1.157193 |
| rod shape-determining protei | 1.083385 | 1.011230797 | -0.09943 | 0.358614 | -1.11972 |
| UDP-N-acetylmuramoylalanin | 0.925185 | 0.908755304 | -0.02585 | 0.358647 | -1.10858 |
| RND efflux transporter | 1.035108 | 0.989747655 | -0.06465 | 0.359707 | -1.13321 |
| hypothetical protein ABUW_1 | 1.06998 | 1.03602357 | -0.04653 | 0.359904 | -1.05827 |
| hypothetical protein ABUW_2 | 0.766814 | 0.690687129 | -0.15084 | 0.360145 | -1.03269 |
| crispr-associated protein, Csy | 0.927956 | 0.949108995 | 0.032517 | 0.360698 | 1.100848 |
| acetoin:2,6-dichlorophenolinc | 0.942821 | 1.216414543 | 0.367579 | 0.361683 | 1.163862 |
| bacterioferritin comigratory p | 0.875579 | 0.957943235 | 0.129704 | 0.36263 | 1.155076 |
| hypothetical protein ABUW_2 | 1.070433 | 1.14271491 | 0.09427 | 0.362663 | 1.045078 |
| hypothetical protein ABUW_0 | 0.945379 | 1.041963523 | 0.140341 | 0.362733 | 1.157602 |
| DEAD/DEAH box helicase...85 | 0.733783 | 1.473398232 | 1.005721 | 0.36464 | 1.162669 |
| urease, gamma subunit | 0.954204 | 0.996822629 | 0.06304 | 0.364678 | 1.049476 |
| protein-tyrosine-phosphatase | 1.03671 | 0.996316449 | -0.05734 | 0.364882 | -1.02209 |
| phenylacetate-CoA oxygenase | 0.940281 | 1.221278089 | 0.377227 | 0.365941 | 1.129844 |
| phosphate ABC transporter, s | 1.061436 | 1.032606775 | -0.03973 | 0.366202 | -1.12131 |
| hypothetical protein ABUW_3 | 0.882994 | 1.039824124 | 0.235863 | 0.366514 | 1.147724 |
| FAD linked oxidase domain pr | 0.931542 | 0.956160881 | 0.037633 | 0.366587 | 1.06028 |
| argininosuccinate synthase | 0.967256 | 0.945188181 | -0.0333 | 0.366794 | -1.02474 |
| undecaprenyldiphospho-mura | 0.953285 | 1.000726528 | 0.070068 | 0.36758 | 1.110145 |
| hypothetical protein ABUW_3 | 0.677104 | 1.606605615 | 1.246567 | 0.367679 | 1.153588 |
| hypothetical protein ABUW_0 | 0.968303 | 0.953906316 | -0.02161 | 0.367879 | -1.01418 |
| ribonuclease T | 0.96639 | 1.069076272 | 0.145688 | 0.36885 | 1.057255 |
| type IV / VI secretion system p | 0.950512 | 1.052832667 | 0.1475 | 0.369203 | 1.100953 |
| putative oxidoreductase | 1.097152 | 1.292087523 | 0.23594 | 0.369553 | 1.148036 |
| transcriptional regulator AraC | 0.792483 | 0.887889 | 0.163999 | 0.370761 | 1.007633 |
| ABC transporter ATP-binding p | 0.980441 | 0.966373808 | -0.02085 | 0.37242 | -1.03548 |
| nicotinamide-nucleotide ader | 0.993913 | 1.016650245 | 0.032632 | 0.372572 | 1.090075 |

| | | | | | |
|---|---|---|---|---|---|
| hypothetical protein ABUW_3 | 0.632069 | 1.99365313 | 1.65726 | 0.37279 | 1.138249 |
| hypothetical protein ABUW_0 | 1.049286 | 1.006985234 | -0.05937 | 0.374022 | -1.02951 |
| pyrroline-5-carboxylate reduc | 0.906921 | 0.892284808 | -0.02347 | 0.374323 | -1.07068 |
| aldose 1-epimerase | 1.078087 | 1.102662342 | 0.032517 | 0.375027 | 1.00384 |
| rhodanese domain protein...1 | 0.904942 | 1.155633451 | 0.352787 | 0.375995 | 1.124615 |
| Bacterial transferase hexapep | 1.192861 | 1.119907374 | -0.09105 | 0.37645 | -1.11394 |
| macrolide export ATP-binding | 1.021378 | 0.976325498 | -0.06508 | 0.377373 | -1.04496 |
| phospholipase D/transphosph | 0.994353 | 0.965318143 | -0.04275 | 0.377709 | -1.01938 |
| hypothetical protein ABUW_0 | 0.54855 | 2.131064494 | 1.95788 | 0.378012 | 1.123339 |
| sulfite reductase | 1.018278 | 0.984630282 | -0.04848 | 0.378088 | -1.05661 |
| phosphoserine aminotransfer | 0.985481 | 0.960792276 | -0.0366 | 0.378296 | -1.04325 |
| hypothetical protein ABUW_3 | 0.988121 | 1.032536185 | 0.063433 | 0.379616 | 1.063417 |
| 2-nitropropane dioxygenase.. | 1.117793 | 1.028226293 | -0.12049 | 0.380269 | -1.09473 |
| thermostable carboxypeptida | 0.9729 | 0.945888145 | -0.04062 | 0.381778 | -1.07258 |
| dihydrolipoamide acetyltrans | 0.984811 | 1.164342311 | 0.241596 | 0.382407 | 1.109288 |
| DNA polymerase III, beta subu | 0.97471 | 0.960135889 | -0.02173 | 0.38244 | -0.98209 |
| transcriptional regulator, Ara( | 1.114046 | 1.007098566 | -0.1456 | 0.383824 | -1.00803 |
| iron-sulfur cluster assembly p | 1.018409 | 0.93157666 | -0.12857 | 0.384487 | -1.09286 |
| 3'(2'),5'-bisphosphate nucleot | 0.982132 | 1.023280213 | 0.059213 | 0.384896 | 1.098711 |
| peptidase S24 S26A and S26B | 0.964497 | 0.994860542 | 0.044717 | 0.38526 | 0.988633 |
| acyl-CoA dehydrogenase dom | 0.989939 | 1.04605478 | 0.079547 | 0.3856 | 1.061596 |
| hypothetical protein ABUW_1 | 1.296679 | 1.047249127 | -0.30822 | 0.385916 | -0.9993 |
| transporter, anion:cation sym | 1.063806 | 0.941465876 | -0.17625 | 0.38598 | -1.00946 |
| type IV pilus assembly proteir | 2.379214 | 0.27423056 | -3.11702 | 0.388126 | -1.09402 |
| GTP-binding protein TypA/Bip | 0.996364 | 0.925190972 | -0.10692 | 0.388966 | -1.05473 |
| hypothetical protein ABUW_1 | 1.641942 | 8.255748035 | 2.329996 | 0.389055 | 1.083369 |
| thiosulfate-binding protein...1 | 1.036134 | 0.951561213 | -0.12284 | 0.391333 | -0.9882 |
| short chain dehydrogenase...1 | 1.031137 | 0.981526004 | -0.07114 | 0.392335 | -1.00713 |
| transcriptional activator prote | 0.955809 | 1.263951494 | 0.403147 | 0.39237 | 1.080317 |
| hypothetical protein ABUW_4 | 1.016841 | 1.16213181 | 0.19268 | 0.392626 | 1.000621 |
| hypothetical protein ABUW_2 | 1.052334 | 1.255866035 | 0.25509 | 0.393412 | 0.957607 |
| type IV pilus methyl-accepting | 1.166755 | 1.954758039 | 0.744489 | 0.394052 | 1.059999 |
| general secretion pathway pro | 0.919164 | 0.980855284 | 0.093718 | 0.394681 | 1.00305 |
| endonuclease/exonuclease/p | 0.986195 | 0.940694295 | -0.06815 | 0.395163 | -0.95959 |
| putative aminotransferase | 1.007377 | 1.036883329 | 0.04165 | 0.395225 | 1.023127 |
| cobalamin adenosyltransferas | 1.06586 | 0.970289685 | -0.13553 | 0.395878 | -1.05158 |
| NADH dehydrogenase I chain | 0.955669 | 0.972049091 | 0.024519 | 0.397756 | 1.021161 |
| single-stranded-DNA-specific | 0.907796 | 0.982778467 | 0.114498 | 0.399119 | 1.028504 |
| carbamoyl-phosphate synthas | 0.817752 | 0.801466051 | -0.02902 | 0.399209 | -0.97935 |
| glutamate racemase | 0.858934 | 0.919907454 | 0.098942 | 0.399427 | 0.996797 |
| peptidase S49 | 0.91008 | 0.968158497 | 0.08925 | 0.39955 | 0.957084 |
| hypothetical protein ABUW_0 | 0.971355 | 1.036501084 | 0.093651 | 0.402875 | 1.049967 |
| nitrite reductase [NAD(P)H], la | 1.04329 | 0.987973308 | -0.0786 | 0.404305 | -0.95612 |
| NADH dehydrogenase I chain | 1.024777 | 0.955502213 | -0.10098 | 0.405043 | -0.99376 |
| 5-methyltetrahydropteroyltrig | 0.980093 | 1.32045954 | 0.43005 | 0.405949 | 1.040845 |
| lysyl-tRNA synthetase...2233 | 0.932557 | 0.912004049 | -0.03215 | 0.406138 | -0.98469 |
| hypothetical protein ABUW_2 | 0.92269 | 1.486081482 | 0.687595 | 0.408282 | 1.032803 |

| | | | | | |
|---|---|---|---|---|---|
| hypothetical protein ABUW_( | 1.017229 | 1.079463393 | 0.08567 | 0.40869 | 0.986714 |
| copper resistance protein B... | 1.079056 | 1.011494427 | -0.09328 | 0.408756 | -1.01033 |
| large conductance mechanos( | 1.131607 | 1.036237144 | -0.12702 | 0.409141 | -1.01482 |
| hypothetical protein ABUW_3 | 0.929361 | 0.991945406 | 0.094021 | 0.409634 | 0.97582 |
| pseudouridine synthase | 0.974648 | 0.948901698 | -0.03862 | 0.409988 | -0.95575 |
| hypothetical protein ABUW_1 | 1.012599 | 0.974417569 | -0.05545 | 0.410381 | -1.03207 |
| hypothetical protein ABUW_1 | 1.028224 | 0.972135191 | -0.08093 | 0.411125 | -1.02039 |
| hypothetical protein ABUW_3 | 0.977385 | 0.929700573 | -0.07216 | 0.411495 | -0.93531 |
| hypothetical protein ABUW_4 | 0.93071 | 0.878599469 | -0.08313 | 0.411673 | -0.92028 |
| peptidyl-tRNA hydrolase | 1.079267 | 0.985895953 | -0.13054 | 0.412165 | -0.98322 |
| hypothetical protein ABUW_2 | 1.110505 | 1.180095004 | 0.087687 | 0.412683 | 0.92445 |
| transcriptional regulator, Crp/ | 1.152618 | 2.336314546 | 1.01932 | 0.413393 | 1.000488 |
| cysteine synthase A | 0.992641 | 0.946687481 | -0.06838 | 0.414852 | -1.01579 |
| hypothetical protein ABUW_2 | 1.062795 | 1.028629359 | -0.04714 | 0.414871 | -0.91005 |
| argininosuccinate lyase | 0.954143 | 0.933189542 | -0.03203 | 0.416833 | -0.93019 |
| pantothenate kinase, type III | 0.972805 | 0.949999126 | -0.03422 | 0.417255 | -0.99809 |
| DNA primase | 0.948256 | 0.930557346 | -0.02718 | 0.418393 | -0.96129 |
| DEAD/DEAH box helicase...17 | 1.002407 | 1.019527088 | 0.024431 | 0.419991 | 0.90469 |
| UDP-N-acetylmuramoylalanyl | 0.984597 | 1.061019889 | 0.107846 | 0.420723 | 0.958872 |
| hypothetical protein ABUW_1 | 0.954132 | 0.98904024 | 0.051841 | 0.421254 | 0.998476 |
| ABC transporter ATP-binding | 0.955426 | 0.966593382 | 0.016764 | 0.421566 | 0.973122 |
| C4-dicarboxylate transport pr | 0.920926 | 0.896223465 | -0.03923 | 0.421899 | -0.92395 |
| transaldolase | 0.991097 | 1.004939863 | 0.020012 | 0.422244 | 0.894026 |
| glutaredoxin 3 | 1.021958 | 0.991686604 | -0.04338 | 0.422981 | -0.95085 |
| multi-sensor hybrid histidine | 0.97046 | 0.985980456 | 0.022891 | 0.423732 | 0.890449 |
| hypothetical protein ABUW_1 | 1.034058 | 1.017569973 | -0.02319 | 0.424817 | -0.92283 |
| hypothetical protein ABUW_1 | 1.000962 | 1.059457885 | 0.081939 | 0.425215 | 0.952067 |
| apolipoprotein N-acyltransfer | 0.943097 | 0.961910518 | 0.028497 | 0.426019 | 0.895465 |
| hypothetical protein ABUW_3 | 1.072241 | 0.99432713 | -0.10884 | 0.426665 | -0.9426 |
| leucine-responsive regulatory | 1.095523 | 1.021249069 | -0.10128 | 0.428116 | -0.95725 |
| hypothetical protein ABUW_3 | 0.935162 | 0.954703608 | 0.029837 | 0.428746 | 0.944999 |
| penicillin-binding protein 2 | 1.020753 | 0.943143135 | -0.11408 | 0.42888 | -0.91848 |
| thiamine-monophosphate kin | 1.013361 | 0.927176845 | -0.12823 | 0.429155 | -0.92022 |
| hypothetical protein ABUW_( | 1.201828 | 1.088238699 | -0.14324 | 0.429705 | -0.88709 |
| phosphoribosylaminoimidazo | 0.957066 | 0.923713042 | -0.05117 | 0.430493 | -0.96933 |
| hypothetical protein ABUW_2 | 0.935652 | 0.962419058 | 0.040693 | 0.432197 | 0.948392 |
| molybdopterin oxidoreductas | 1.277316 | 1.112475896 | -0.19934 | 0.432253 | -0.89451 |
| putave Na+/H+ antiporter | 0.832487 | 0.797303108 | -0.0623 | 0.432482 | -0.88304 |
| FAD-dependent pyridine nucl( | 0.932346 | 1.006738527 | 0.110751 | 0.432899 | 0.904686 |
| hypothetical protein ABUW_1 | 1.008332 | 0.97643157 | -0.04638 | 0.433087 | -0.92527 |
| hypothetical protein ABUW_1 | 0.947665 | 1.144726375 | 0.272554 | 0.433623 | 0.958153 |
| tRNA--hydroxylase | 0.914309 | 0.979094696 | 0.098766 | 0.433648 | 0.880746 |
| hypothetical protein ABUW_2 | 0.898354 | 0.788598415 | -0.18799 | 0.433739 | -0.96717 |
| iron transporter | 1.00264 | 0.969242931 | -0.04887 | 0.434321 | -0.87288 |
| two-component system sensc | 1.011645 | 1.048686974 | 0.05188 | 0.434796 | 0.869195 |
| UDP-glucose 4-epimerase...2( | 1.021193 | 1.00478478 | -0.02337 | 0.437568 | -0.92813 |
| hypothetical protein ABUW_1 | 1.068475 | 1.100204796 | 0.042219 | 0.439185 | 0.886021 |

| | | | | | |
|---|---|---|---|---|---|
| hypothetical protein ABUW_( | 1.052733 | 1.095517272 | 0.057473 | 0.440849 | 0.85962 |
| nucleotide-binding protein | 0.95297 | 0.970621406 | 0.026477 | 0.442092 | 0.900256 |
| 2,3-bisphosphoglycerate-inde | 0.924697 | 0.933499278 | 0.013669 | 0.442579 | 0.926289 |
| K+ uptake system component | 0.999941 | 1.072411926 | 0.100944 | 0.443747 | 0.9404 |
| SsrA-binding protein | 0.961353 | 0.933285976 | -0.04275 | 0.444423 | -0.87073 |
| lipoprotein-releasing system / | 0.998703 | 1.013393597 | 0.021067 | 0.44486 | 0.874898 |
| transcriptional regulator, Gntl | 0.960351 | 0.988490464 | 0.041666 | 0.444877 | 0.878279 |
| glucose-6-phosphate isomera | 0.981079 | 0.992559622 | 0.016784 | 0.444929 | 0.85414 |
| phosphoribosylpyrophosphat | 0.965911 | 0.992128003 | 0.038635 | 0.445351 | 0.88077 |
| alkaline phosphatase | 0.907345 | 0.948763232 | 0.064397 | 0.44544 | 0.892063 |
| hypothetical protein ABUW_3 | 1.03759 | 0.974782791 | -0.09008 | 0.446227 | -0.89477 |
| riboflavin biosynthesis protei | 0.981855 | 0.965293563 | -0.02454 | 0.446755 | -0.87918 |
| putative dihydrodipicolinate s | 1.061997 | 1.028014656 | -0.04692 | 0.446811 | -0.86656 |
| major facilitator family transp | 0.788579 | 0.913781003 | 0.212593 | 0.447063 | 0.882916 |
| YCII-related protein...1512 | 0.906544 | 0.938338665 | 0.049732 | 0.447116 | 0.84302 |
| acyl-CoA dehydrogenase...72 | 0.934424 | 1.021854348 | 0.12904 | 0.447303 | 0.934994 |
| pH adaptation potassium efflu | 0.923411 | 0.805881652 | -0.19641 | 0.447407 | -0.92508 |
| tail tape measure protein...50 | 1.231295 | 1.145721864 | -0.10392 | 0.447426 | -0.91281 |
| hypothetical protein ABUW_3 | 1.065041 | 1.02325651 | -0.05774 | 0.449031 | -0.89175 |
| hypothetical protein ABUW_( | 1.03923 | 1.004853408 | -0.04853 | 0.449059 | -0.85956 |
| bla(GES-14) (plasmid) | 0.919174 | 1.184208725 | 0.365513 | 0.449828 | 0.929261 |
| enoyl-CoA hydratase...1218 | 1.028868 | 0.992427305 | -0.05202 | 0.45125 | -0.88774 |
| short-chain enoyl-CoA hydrat | 1.014545 | 1.001671361 | -0.01842 | 0.451431 | -0.88007 |
| aminoacyl-histidine dipeptida | 1.018078 | 0.99666668 | -0.03067 | 0.451742 | -0.87664 |
| hypothetical protein ABUW_( | 1.052668 | 1.005986356 | -0.06544 | 0.452667 | -0.89274 |
| hypothetical protein ABUW_( | 1.050276 | 0.998726239 | -0.07261 | 0.452851 | -0.9121 |
| multidrug efflux protein...539 | 1.136814 | 1.084949763 | -0.06737 | 0.453789 | -0.8736 |
| sulfur relay protein TusD/Dsrl | 0.956984 | 1.07835443 | 0.172265 | 0.454532 | 0.898466 |
| tRNA delta(2)-isopentenylpyr | 0.927907 | 0.965440587 | 0.057207 | 0.455868 | 0.869734 |
| mechanosensitive ion channe | 0.906443 | 0.946664143 | 0.062637 | 0.458507 | 0.896607 |
| hypothetical protein ABUW_6 | 0.997309 | 0.94205312 | -0.08223 | 0.45941 | -0.81775 |
| ATP synthase F1, alpha subun | 0.977394 | 0.967368892 | -0.01487 | 0.459852 | -0.89957 |
| Putative DNA binding protein | 1.058825 | 1.002546133 | -0.0788 | 0.460907 | -0.81913 |
| ribosomal protein L2 | 0.987496 | 0.958502199 | -0.04299 | 0.46132 | -0.84326 |
| phosphoribosylformylglycinar | 0.922183 | 0.89956995 | -0.03582 | 0.462734 | -0.89955 |
| endonuclease | 0.980307 | 0.956927599 | -0.03482 | 0.464181 | -0.84322 |
| hypothetical protein ABUW_2 | 1.127609 | 1.160865189 | 0.041933 | 0.464494 | 0.875243 |
| hypothetical protein ABUW_1 | 1.023688 | 1.000768314 | -0.03267 | 0.465212 | -0.81814 |
| isochorismate synthetase | 1.478284 | 1.595762679 | 0.110322 | 0.465614 | 0.8061 |
| hypothetical protein ABUW_( | 1.052402 | 1.114578421 | 0.082812 | 0.466621 | 0.847022 |
| radical SAM domain protein | 1.017753 | 1.051416115 | 0.046946 | 0.466692 | 0.817758 |
| hypothetical protein ABUW_( | 1.10456 | 1.014102655 | -0.12327 | 0.467618 | -0.85971 |
| hypothetical protein ABUW_1 | 1.181453 | 1.114605049 | -0.08403 | 0.468282 | -0.83703 |
| ATP-dependent protease La | 1.023273 | 0.999810705 | -0.03346 | 0.469201 | -0.80886 |
| hypothetical protein ABUW_1 | 1.008175 | 0.98621006 | -0.03178 | 0.469726 | -0.80553 |
| multidrug efflux protein AdeI | 1.006469 | 1.026296646 | 0.028145 | 0.470035 | 0.846118 |
| hypothetical protein ABUW_2 | 0.879708 | 0.983318581 | 0.160633 | 0.470958 | 0.87472 |

| | | | | | |
|---|---|---|---|---|---|
| TRAP C4-dicarboxylate transp | 0.943214 | 0.973643197 | 0.045809 | 0.471798 | 0.873352 |
| non-ribosomal peptide synthe | 1.14608 | 1.045368923 | -0.1327 | 0.471966 | -0.82812 |
| tryptophan synthase, beta sul | 0.991142 | 0.966030317 | -0.03702 | 0.473505 | -0.82507 |
| dienelactone hydrolase...1877 | 1.032599 | 1.069400073 | 0.050521 | 0.473751 | 0.831981 |
| ammonium transporter...7 | 1.039306 | 1.160347931 | 0.158937 | 0.473832 | 0.792967 |
| hypothetical protein ABUW_0 | 0.978332 | 1.036562923 | 0.083411 | 0.47438 | 0.835339 |
| Aminoglycoside phosphotrans | 0.962841 | 0.934317193 | -0.04338 | 0.475562 | -0.85092 |
| DNA polymerase V componer | 0.998446 | 0.981039196 | -0.02537 | 0.476686 | -0.80874 |
| hypothetical protein ABUW_2 | 0.921851 | 0.988461183 | 0.100651 | 0.476767 | 0.801917 |
| hypothetical protein ABUW_3 | 0.996121 | 1.477056522 | 0.568332 | 0.478364 | 0.855771 |
| arsenate reductase...599 | 0.973737 | 1.029901177 | 0.080902 | 0.478665 | 0.85213 |
| 1-deoxy-D-xylulose-5-phospha | 0.978389 | 0.952022377 | -0.03941 | 0.479474 | -0.82638 |
| fumarate hydratase | 0.923245 | 0.939463928 | 0.025124 | 0.481051 | 0.819414 |
| DSBA oxidoreductase | 0.984118 | 0.956555584 | -0.04098 | 0.482106 | -0.794 |
| chromosomal replication initi | 0.987372 | 1.002594637 | 0.022073 | 0.482663 | 0.775968 |
| putative peroxidase | 1.004526 | 1.028046012 | 0.03339 | 0.483309 | 0.774645 |
| acetyltransferase, gnat family | 1.04804 | 1.02071306 | -0.03812 | 0.484142 | -0.77199 |
| type VI secretion system effec | 0.707765 | 1.10195721 | 0.638726 | 0.484417 | 0.8503 |
| copper resistance protein B.... | 1.032064 | 1.061911446 | 0.041131 | 0.484588 | 0.803357 |
| protein TolQ | 0.964337 | 0.993156523 | 0.042484 | 0.485217 | 0.827214 |
| hypothetical protein ABUW_0 | 1.046719 | 1.002211586 | -0.06269 | 0.485695 | -0.82352 |
| tRNA-dihydrouridine synthase | 1.043275 | 1.088885608 | 0.061733 | 0.486493 | 0.778063 |
| hypothetical protein ABUW_1 | 0.85215 | 0.935675997 | 0.134902 | 0.487432 | 0.826414 |
| hypothetical protein ABUW_0 | 0.996759 | 1.008262825 | 0.016555 | 0.488557 | 0.765769 |
| hypothetical protein ABUW_4 | 1.002763 | 0.9336372 | -0.10305 | 0.48893 | -0.81193 |
| hypothetical protein ABUW_0 | 0.922612 | 1.014807259 | 0.13741 | 0.489535 | 0.787787 |
| hypothetical protein ABUW_2 | 0.972289 | 1.000220075 | 0.04086 | 0.489667 | 0.808983 |
| transcriptional regulator, LysF | 1.049512 | 1.120925442 | 0.094971 | 0.490821 | 0.784521 |
| hypothetical protein ABUW_2 | 0.933556 | 0.870080361 | -0.10159 | 0.49126 | -0.82272 |
| hypothetical protein ABUW_3 | 0.931236 | 0.962108289 | 0.047052 | 0.49157 | 0.821956 |
| acetate--CoA ligase | 1.039109 | 1.01035378 | -0.04049 | 0.491844 | -0.76223 |
| phosphoribosylaminoimidazo | 0.965854 | 0.954176342 | -0.01755 | 0.49205 | -0.81333 |
| hypothetical protein ABUW_2 | 1.05651 | 1.036061771 | -0.0282 | 0.492503 | -0.79026 |
| DNA mismatch repair protein | 0.981615 | 0.948587283 | -0.04938 | 0.49255 | -0.77302 |
| malonate decarboxylase, epsi | 0.987018 | 1.073149656 | 0.120704 | 0.493263 | 0.815799 |
| hypothetical protein ABUW_3 | 0.940444 | 0.976239413 | 0.053893 | 0.493905 | 0.769382 |
| 4-carboxymuconolactone dec | 1.000657 | 1.036024788 | 0.050111 | 0.494537 | 0.769225 |
| oxidoreductase, short chain d | 0.891955 | 0.937627338 | 0.072044 | 0.494691 | 0.790668 |
| 2'-aminoglycoside nucleotidyl | 1.050112 | 0.964259656 | -0.12305 | 0.495165 | -0.75912 |
| UDP-N-acetylglucosamine 2-e | 1.019934 | 1.040416902 | 0.028686 | 0.495302 | 0.750384 |
| hypothetical protein ABUW_2 | 1.071025 | 1.02913427 | -0.05756 | 0.495555 | -0.8138 |
| DNA-binding protein (plasmid | 0.989695 | 1.060953201 | 0.100305 | 0.496485 | 0.749767 |
| phosphoserine phosphatase/l | 0.909337 | 0.885492341 | -0.03834 | 0.496593 | -0.76463 |
| peptidase M23B | 1.017701 | 0.996908034 | -0.02978 | 0.496818 | -0.81989 |
| transcriptional regulator, LysF | 1.010494 | 0.938388096 | -0.1068 | 0.497582 | -0.81287 |
| hypothetical protein ABUW_1 | 1.034285 | 1.022052742 | -0.01716 | 0.497872 | -0.79994 |
| hypothetical protein ABUW_0 | 1.058069 | 1.072835539 | 0.019996 | 0.498644 | 0.754966 |

| | | | | | |
|---|---|---|---|---|---|
| hypothetical protein ABUW_( | 0.970591 | 0.954036 | -0.02482 | 0.499692 | -0.79082 |
| NDP-sugar dehydrogenase | 0.979793 | 1.009356234 | 0.042886 | 0.500157 | 0.812458 |
| malonate decarboxylase, alph | 0.948266 | 0.991912956 | 0.064922 | 0.500371 | 0.772688 |
| acetolactate synthase, small s | 0.901886 | 0.93172221 | 0.046955 | 0.50214 | 0.799747 |
| Dyp-type peroxidase | 1.013886 | 0.942245789 | -0.10572 | 0.502399 | -0.7951 |
| hypothetical protein ABUW_( | 0.951572 | 0.965218231 | 0.020543 | 0.503221 | 0.738449 |
| lipoate-protein ligase B | 0.939267 | 1.012150156 | 0.107816 | 0.503842 | 0.788252 |
| hypothetical protein ABUW_2 | 0.98752 | 0.944443151 | -0.06435 | 0.504578 | -0.79062 |
| enoyl-CoA hydratase/isomera | 0.968489 | 0.989054335 | 0.030314 | 0.50484 | 0.794358 |
| transcription termination/ant | 0.965845 | 0.993044151 | 0.040067 | 0.507815 | 0.792943 |
| hypothetical protein ABUW_3 | 0.91074 | 0.96202285 | 0.079032 | 0.507842 | 0.754733 |
| toluene tolerance protein Ttg | 0.918961 | 0.945253102 | 0.040697 | 0.507991 | 0.780802 |
| haloacid dehydrogenase | 0.657662 | 0.612181631 | -0.10339 | 0.508207 | -0.76979 |
| pilus assembly protein tip-ass | 1.061709 | 1.110485269 | 0.064802 | 0.508402 | 0.728208 |
| DNA repair system | 1.04112 | 0.972741582 | -0.09801 | 0.508676 | -0.76568 |
| OmpA/MotB...1765 | 1.011386 | 0.997553149 | -0.01987 | 0.508784 | -0.73869 |
| putative RND family drug trar | 0.987797 | 1.030685218 | 0.061317 | 0.5102 | 0.762125 |
| hypothetical protein ABUW_( | 1.014711 | 1.05564089 | 0.05705 | 0.511029 | 0.791929 |
| hypothetical protein ABUW_4 | 0.944663 | 0.967358498 | 0.034251 | 0.511688 | 0.724525 |
| 2-amino-4-hydroxy-6-hydroxy | 0.955541 | 1.042842193 | 0.126132 | 0.5123 | 0.730057 |
| sulfate adenylyltransferase su | 1.001324 | 0.991303113 | -0.01451 | 0.512932 | -0.73292 |
| aldehyde dehydrogenase...22 | 1.073605 | 1.034840322 | -0.05306 | 0.513111 | -0.77541 |
| hypothetical protein ABUW_( | 0.941117 | 0.983273057 | 0.063219 | 0.513649 | 0.741216 |
| P-hydroxybenzoate hydroxyla | 1.103066 | 0.964667674 | -0.19342 | 0.513966 | -0.74302 |
| glycine cleavage system trans | 0.977139 | 1.015552825 | 0.05563 | 0.515045 | 0.738889 |
| O-antigen polymerase family | 0.98193 | 1.027767619 | 0.065823 | 0.515516 | 0.713032 |
| high-affinity choline transport | 0.90847 | 0.932607303 | 0.037831 | 0.51616 | 0.757277 |
| PpiC-type peptidyl-prolyl cis-t | 0.987497 | 0.999777195 | 0.017831 | 0.516174 | 0.74798 |
| aspartate aminotransferase A | 0.916343 | 0.906386786 | -0.01576 | 0.516315 | -0.71895 |
| diaminobutyrate decarboxyla | 0.985212 | 0.992698153 | 0.01092 | 0.51673 | 0.715729 |
| hypothetical protein ABUW_( | 0.8824 | 0.927305022 | 0.071611 | 0.517948 | 0.710802 |
| monooxygenase, flavin-bindir | 1.026126 | 0.990082906 | -0.05159 | 0.518445 | -0.71253 |
| hypothetical protein ABUW_( | 0.921369 | 0.953528236 | 0.049497 | 0.519398 | 0.710326 |
| O-methyltransferase | 0.99517 | 0.963197017 | -0.04711 | 0.521239 | -0.72698 |
| hypothetical protein ABUW_3 | 0.950228 | 0.962168236 | 0.018015 | 0.522375 | 0.759626 |
| hypothetical protein ABUW_4 | 0.970966 | 0.937469416 | -0.05065 | 0.522454 | -0.71078 |
| tRNA (uracil-5-)-methyltransfe | 0.959951 | 0.937055839 | -0.03483 | 0.522762 | -0.73961 |
| hypothetical protein ABUW_1 | 0.950966 | 1.001540503 | 0.074755 | 0.52382 | 0.703173 |
| transcriptional regulator, TetF | 1.043805 | 0.962216075 | -0.11742 | 0.523821 | -0.76525 |
| ammonium transporter...748 | 0.924299 | 0.891120843 | -0.05274 | 0.524019 | -0.70809 |
| aquaporin Z | 1.080746 | 1.001670728 | -0.10962 | 0.524393 | -0.69859 |
| sulfate ABC transporter, ATP-l | 0.955298 | 1.001538579 | 0.068195 | 0.525262 | 0.747343 |
| hypothetical protein ABUW_( | 0.792954 | 0.83705311 | 0.078081 | 0.525468 | 0.697733 |
| hypothetical protein ABUW_2 | 0.904208 | 0.926538313 | 0.035195 | 0.526241 | 0.697691 |
| hypothetical protein ABUW_2 | 0.901383 | 0.868210002 | -0.0541 | 0.527097 | -0.7266 |
| NADH dehydrogenase I chain | 0.922348 | 0.968330329 | 0.070188 | 0.527625 | 0.754062 |
| cysteinyl-tRNA synthetase | 1.020471 | 1.007786988 | -0.01804 | 0.529454 | -0.68923 |

| | | | | | |
|---|---|---|---|---|---|
| hypothetical protein ABUW_2 | 1.008078 | 1.021519929 | 0.01911 | 0.531358 | 0.68446 |
| peptidase M61 | 1.004882 | 1.140183134 | 0.182239 | 0.532493 | 0.715611 |
| transcriptional regulator, GntI | 0.997484 | 1.025714795 | 0.040264 | 0.532961 | 0.684095 |
| DNA strand exchange and rec | 1.024685 | 1.013422356 | -0.01594 | 0.534376 | -0.68941 |
| hypothetical protein ABUW_2 | 0.93433 | 0.908208571 | -0.04091 | 0.534378 | -0.69196 |
| hypothetical protein ABUW_0 | 0.974561 | 1.077462134 | 0.144813 | 0.536145 | 0.702285 |
| GTP pyrophosphokinase (ppG | 1.02261 | 0.999145925 | -0.03349 | 0.536698 | -0.67497 |
| xanthine dehydrogenase, mol | 1.108521 | 1.23001335 | 0.150037 | 0.537881 | 0.688699 |
| glutamate synthase, small sub | 0.960011 | 0.975094242 | 0.022491 | 0.539394 | 0.713093 |
| coproporphyrinogen III oxidas | 0.968262 | 1.018550505 | 0.073049 | 0.54005 | 0.708161 |
| glycosyl transferase, family 3 | 0.99082 | 0.978291972 | -0.01836 | 0.541074 | -0.71896 |
| hypothetical protein ABUW_0 | 0.929513 | 1.024684933 | 0.140634 | 0.541361 | 0.712654 |
| transcription termination fact | 0.973988 | 0.992998318 | 0.027888 | 0.54307 | 0.716048 |
| EsvE2 | 1.016887 | 1.045442313 | 0.039954 | 0.543899 | 0.719165 |
| iscRSUA operon repressor | 1.103666 | 1.221825306 | 0.146734 | 0.544027 | 0.723603 |
| phenylacetate-CoA oxygenase | 0.89321 | 1.075841196 | 0.268394 | 0.545508 | 0.714619 |
| N-acetyl-beta-glucosaminidas | 1.007205 | 0.977332101 | -0.04344 | 0.545683 | -0.71977 |
| hypothetical protein ABUW_2 | 1.002701 | 0.979616175 | -0.0336 | 0.546978 | -0.71102 |
| methylenetetrahydrofolate re | 1.045854 | 0.987872725 | -0.08228 | 0.547033 | -0.7036 |
| dihydrofolate reductase | 0.885734 | 0.923151911 | 0.059694 | 0.548033 | 0.703151 |
| inosine-5'-monophosphate de | 0.980318 | 1.016266811 | 0.051958 | 0.548298 | 0.709899 |
| fatty acid desaturase...1337 | 1.026666 | 0.998230649 | -0.04052 | 0.549365 | -0.65421 |
| ubiquinone biosynthesis hydr | 0.965521 | 0.980823639 | 0.022686 | 0.552731 | 0.647276 |
| ubiquinone biosynthesis prote | 0.995444 | 0.975531758 | -0.02915 | 0.553538 | -0.65475 |
| glutamate 5-kinase | 1.040262 | 1.009948876 | -0.04266 | 0.553886 | -0.69081 |
| hypothetical protein ABUW_1 | 1.042895 | 1.105791556 | 0.084486 | 0.555607 | 0.65188 |
| heme-binding protein A...344 | 0.969814 | 1.007530722 | 0.055044 | 0.556706 | 0.65022 |
| FilF | 1.001741 | 1.018989539 | 0.02463 | 0.557617 | 0.669475 |
| protein DedA | 1.170126 | 1.104660774 | -0.08306 | 0.557806 | -0.63958 |
| host factor Hfq | 0.979755 | 0.964529631 | -0.0226 | 0.557972 | -0.68237 |
| methyltransferase GidB | 0.985826 | 0.973652747 | -0.01793 | 0.559617 | -0.63581 |
| succinyl-diaminopimelate des | 1.034131 | 0.992623168 | -0.0591 | 0.559668 | -0.69251 |
| YaeQ protein | 0.893864 | 0.926496337 | 0.05173 | 0.560502 | 0.689537 |
| RND type efflux pump | 1.089808 | 1.106910136 | 0.022464 | 0.562198 | 0.676646 |
| hypothetical protein ABUW_1 | 1.055092 | 1.041347461 | -0.01892 | 0.563181 | -0.636 |
| co-chaperone GrpE | 0.993689 | 1.023495293 | 0.042638 | 0.564148 | 0.67731 |
| hemerythrin/HHE cation-bind | 0.936222 | 1.033476108 | 0.142583 | 0.5648 | 0.637056 |
| GTPase | 0.973655 | 1.01807868 | 0.064366 | 0.565852 | 0.625578 |
| hydrolase, NUDIX family prote | 0.930341 | 0.883597333 | -0.07437 | 0.565966 | -0.62882 |
| tRNA (5-methylaminomethyl- | 0.966536 | 0.98434606 | 0.026342 | 0.566046 | 0.625956 |
| 1-phosphofructokinase | 1.031842 | 1.010119883 | -0.0307 | 0.566157 | -0.67989 |
| NADPH-dependent 7-cyano-7 | 0.956032 | 0.97871158 | 0.033825 | 0.566218 | 0.677018 |
| hypothetical protein ABUW_2 | 0.975323 | 1.000946004 | 0.037413 | 0.566369 | 0.624163 |
| hypothetical protein ABUW_1 | 0.982058 | 1.001720001 | 0.028599 | 0.567629 | 0.666525 |
| hypothetical protein ABUW_2 | 0.998656 | 1.02724853 | 0.040725 | 0.56789 | 0.621951 |
| transcriptional regulator, GntI | 0.99184 | 0.927818585 | -0.09626 | 0.570564 | -0.62092 |
| transcriptional regulator, LysR | 1.077366 | 1.093711663 | 0.021724 | 0.570591 | 0.61742 |

| | | | | | |
|---|---|---|---|---|---|
| multidrug efflux protein AdeC | 1.005302 | 1.034838306 | 0.041777 | 0.571409 | 0.615711 |
| aldo-keto reductase | 1.051873 | 0.967222014 | -0.12104 | 0.571573 | -0.66339 |
| ribosomal RNA small subunit | 0.968753 | 1.025395571 | 0.081979 | 0.572116 | 0.61656 |
| hypothetical protein ABUW_( | 0.979653 | 0.936163022 | -0.06551 | 0.573166 | -0.63766 |
| aldehyde dehydrogenase...63 | 0.987134 | 0.9361594 | -0.07649 | 0.573372 | -0.66378 |
| outer membrane efflux prote | 0.912241 | 0.952188225 | 0.061831 | 0.573647 | 0.628641 |
| crispr-associated protein, Csy | 1.051044 | 1.035813053 | -0.02106 | 0.575647 | -0.61762 |
| preprotein translocase, YajC s | 0.987966 | 0.973912344 | -0.02067 | 0.576064 | -0.65117 |
| hypothetical protein ABUW_3 | 1.00137 | 0.986817331 | -0.02112 | 0.576224 | -0.61012 |
| chorismate mutase...2047 | 0.952594 | 0.971833524 | 0.028848 | 0.576619 | 0.63302 |
| 4-aminobutyrate transaminas | 0.90784 | 0.971853365 | 0.0983 | 0.576912 | 0.657039 |
| phage head morphogenesis p | 1.028568 | 1.082739636 | 0.074049 | 0.578467 | 0.61679 |
| nonribosomal peptide synthe | 1.017675 | 1.075689955 | 0.079986 | 0.579272 | 0.623088 |
| transcriptional regulator, Asn | 0.975077 | 0.947169445 | -0.04189 | 0.580064 | -0.61523 |
| coenzyme PQQ biosynthesis p | 1.073131 | 1.090651661 | 0.023365 | 0.580687 | 0.634322 |
| transcriptional regulator, LysF | 1.019336 | 0.981669211 | -0.05432 | 0.582783 | -0.6115 |
| sulfurtransferase TusA | 1.006619 | 1.045349944 | 0.054469 | 0.583332 | 0.603433 |
| phospholipase D/Transphospl | 0.9963 | 0.975076886 | -0.03106 | 0.583688 | -0.6286 |
| acyl-CoA N-acyltransferase | 0.986066 | 0.937968988 | -0.07214 | 0.584164 | -0.5947 |
| regulator of ribonuclease acti | 1.006336 | 0.986665233 | -0.02848 | 0.585073 | -0.62426 |
| peptidase S24 S26A and S26B | 1.081846 | 1.044718582 | -0.05038 | 0.586871 | -0.60995 |
| transcriptional regulator, Bad | 1.053924 | 1.026251874 | -0.03839 | 0.58796 | -0.60921 |
| heat shock protein 15 | 0.991157 | 0.929566831 | -0.09256 | 0.588017 | -0.63591 |
| hypothetical protein ABUW_( | 1.001228 | 1.073507279 | 0.100561 | 0.588381 | 0.594561 |
| transcriptional regulator, LysF | 1.117299 | 1.047264646 | -0.09339 | 0.588634 | -0.59127 |
| Putative tellurique resistant p | 0.999554 | 0.94536589 | -0.08041 | 0.588924 | -0.59154 |
| phage tail protein | 0.972474 | 1.015783169 | 0.06286 | 0.591453 | 0.625359 |
| protease | 1.013618 | 1.060399227 | 0.065094 | 0.591599 | 0.62569 |
| dihydrodipicolinate synthase | 0.975573 | 0.957543699 | -0.02691 | 0.592008 | -0.61806 |
| 3-deoxy-7-phosphoheptulona | 0.944988 | 0.934436025 | -0.0162 | 0.593969 | -0.60453 |
| hypothetical protein ABUW_( | 1.071919 | 1.114155213 | 0.055754 | 0.595472 | 0.614754 |
| ribonuclease D | 1.186252 | 1.151218386 | -0.04325 | 0.597027 | -0.59522 |
| hypothetical protein ABUW_3 | 1.011162 | 1.039944836 | 0.040492 | 0.597351 | 0.608563 |
| 4-(cytidine 5'-diphospho)-2-C- | 0.973248 | 0.964681654 | -0.01275 | 0.598711 | -0.58678 |
| beta-ketoadipyl CoA thiolase. | 0.9708 | 0.958047315 | -0.01908 | 0.599045 | -0.58316 |
| citrate synthase I | 0.946059 | 0.908593558 | -0.05829 | 0.599669 | -0.60448 |
| Arc domain protein DNA bind | 0.925403 | 0.950266409 | 0.038251 | 0.601191 | 0.566804 |
| phosphoglycerate mutase...1( | 1.007451 | 1.032312315 | 0.03517 | 0.601261 | 0.58222 |
| Type IV secretory pathway, Vi | 0.989025 | 1.048298317 | 0.08397 | 0.602576 | 0.595626 |
| hypothetical protein ABUW_( | 0.909114 | 0.953747984 | 0.069148 | 0.603548 | 0.57492 |
| transcription-repair coupling f | 1.032945 | 1.024856029 | -0.01134 | 0.604386 | -0.56257 |
| hypothetical protein ABUW_2 | 0.997783 | 0.978089078 | -0.02876 | 0.605064 | -0.59626 |
| transcriptional regulator, TetF | 0.994801 | 0.970512222 | -0.03566 | 0.605321 | -0.56621 |
| rhomboid family peptidase | 1.047778 | 1.074772274 | 0.036698 | 0.605589 | 0.559923 |
| hypothetical protein ABUW_( | 1.101468 | 1.044034193 | -0.07726 | 0.605734 | -0.56125 |
| hypothetical protein ABUW_3 | 0.964583 | 0.97768433 | 0.019464 | 0.606035 | 0.5952 |
| ADP-ribose pyrophosphatase | 1.056479 | 0.996015754 | -0.08502 | 0.606875 | -0.56148 |

| | | | | | |
|---|---|---|---|---|---|
| Sel1 repeat family | 0.808804 | 0.876577739 | 0.116092 | 0.608164 | 0.567161 |
| arsenical resistance protein A | 0.97993 | 1.001669601 | 0.031656 | 0.60897 | 0.578693 |
| hypothetical protein ABUW_1 | 0.905645 | 0.971435494 | 0.101172 | 0.609027 | 0.554204 |
| transcriptional regulator, Gntl | 0.960871 | 0.985927876 | 0.037139 | 0.609889 | 0.594602 |
| DsrC family protein | 0.893404 | 0.91080399 | 0.027828 | 0.610855 | 0.551415 |
| hypothetical protein ABUW_2 | 0.985775 | 1.014290602 | 0.041141 | 0.611694 | 0.561189 |
| putative helicase | 0.986287 | 0.968763518 | -0.02586 | 0.613732 | -0.58556 |
| DNA topoisomerase IV, B subu | 1.024795 | 1.010406521 | -0.0204 | 0.613868 | -0.56364 |
| two-component system senso | 0.9723 | 0.983051298 | 0.015865 | 0.614347 | 0.556987 |
| deoxycytidine triphosphate de | 0.953422 | 0.981529888 | 0.041918 | 0.614393 | 0.572715 |
| succinylarginine dihydrolase | 1.004138 | 1.016773116 | 0.018041 | 0.614485 | 0.545371 |
| orotidine 5'-phosphate decarl | 1.002105 | 0.983754075 | -0.02666 | 0.616341 | -0.54448 |
| beta-ribbon domain peptidase | 0.993648 | 0.983954061 | -0.01414 | 0.61656 | -0.54656 |
| lipoprotein, putative...942 | 0.995161 | 0.968758867 | -0.03879 | 0.616987 | -0.5589 |
| transcriptional regulator OhrF | 0.972512 | 1.004910193 | 0.047278 | 0.617499 | 0.577151 |
| glutamate dehydrogenase | 0.981588 | 0.960575791 | -0.03122 | 0.617678 | -0.58012 |
| extracellular solute-binding pı | 0.925952 | 0.985460056 | 0.089859 | 0.617899 | 0.545926 |
| hypothetical protein ABUW_0 | 1.03442 | 1.058412035 | 0.03308 | 0.618218 | 0.574077 |
| transferase hexapeptide repe | 0.997711 | 1.01690922 | 0.027497 | 0.618986 | 0.577777 |
| alpha/beta fold family hydrola | 1.009184 | 1.033052605 | 0.033725 | 0.619545 | 0.557399 |
| hypothetical protein ABUW_0 | 1.002941 | 1.011049025 | 0.011616 | 0.619562 | 0.548539 |
| S-adenosyl-methyltransferase | 1.016404 | 1.009147599 | -0.01034 | 0.62015 | -0.53721 |
| alanine racemase domain-cor | 0.986586 | 0.966752166 | -0.0293 | 0.620254 | -0.53728 |
| ATP synthase F1, gamma subu | 0.947916 | 0.959013287 | 0.016792 | 0.622291 | 0.559207 |
| hypothetical protein ABUW_0 | 1.001969 | 1.033073315 | 0.044105 | 0.623481 | 0.554865 |
| hypothetical protein ABUW_3 | 0.983757 | 1.014759107 | 0.044763 | 0.623785 | 0.570969 |
| proline-specific permease Pro | 1.001927 | 0.942324345 | -0.08848 | 0.625116 | -0.5334 |
| imidazoleglycerol phosphate s | 1.005406 | 1.029029433 | 0.033506 | 0.625199 | 0.567573 |
| hydroxymethylbutenyl pyroph | 1.006004 | 0.989574195 | -0.02376 | 0.625843 | -0.52866 |
| succinate dehydrogenase, cyt | 0.977558 | 0.927269254 | -0.07619 | 0.626023 | -0.53806 |
| ABC transporter permease pro | 0.988065 | 0.964253724 | -0.03519 | 0.630344 | -0.54771 |
| multiphosphoryl transfer prot | 1.051715 | 1.042455922 | -0.01276 | 0.631707 | -0.53555 |
| hypothetical protein ABUW_1 | 0.893887 | 0.727688569 | -0.29677 | 0.631808 | -0.55919 |
| pH adaptation potassium efflu | 0.825847 | 0.809640635 | -0.02859 | 0.632358 | -0.55596 |
| hypothetical protein ABUW_3 | 0.940214 | 0.958593712 | 0.02793 | 0.632823 | 0.546655 |
| two-component heavy metal | 0.9965 | 1.035802055 | 0.055807 | 0.632987 | 0.544436 |
| aminoglycoside phosphotrans | 0.98525 | 0.974495016 | -0.01583 | 0.633339 | -0.54971 |
| transcriptional regulator, LysF | 0.933468 | 0.973823497 | 0.061059 | 0.634686 | 0.536866 |
| acetyl-CoA acyltransferase | 0.997947 | 0.992011762 | -0.00861 | 0.634858 | -0.52241 |
| cobyrinic acid a,c-diamide syn | 0.946462 | 0.974024095 | 0.041413 | 0.635477 | 0.525366 |
| hypothetical protein ABUW_1 | 0.96996 | 0.997878776 | 0.040939 | 0.635533 | 0.528178 |
| AAA ATPase superfamily prote | 0.940743 | 0.954092499 | 0.020329 | 0.635835 | 0.526672 |
| 3-isopropylmalate dehydratas | 1.007982 | 1.036698381 | 0.040527 | 0.637327 | 0.519983 |
| polyphosphate kinase...2069 | 0.979615 | 0.969344251 | -0.01521 | 0.63854 | -0.54764 |
| hypothetical protein ABUW_0 | 1.07818 | 1.050519733 | -0.0375 | 0.639027 | -0.54682 |
| toluene tolerance efflux trans | 0.970545 | 0.976667974 | 0.009072 | 0.639331 | 0.508854 |
| histidine utilization repressor | 0.94382 | 0.963776095 | 0.030186 | 0.641674 | 0.526838 |

| | | | | | |
|---|---|---|---|---|---|
| phage protein...123 | 0.916639 | 0.893136688 | -0.03747 | 0.643638 | -0.51555 |
| auxin-responsive GH3-related | 1.002519 | 1.056793222 | 0.076064 | 0.644402 | 0.516999 |
| flavohemoprotein | 0.989342 | 0.977287928 | -0.01769 | 0.645167 | -0.52284 |
| aspartate 1-decarboxylase | 1.063028 | 1.027526993 | -0.049 | 0.645944 | -0.50071 |
| pH adaptation potassium efflu | 0.940965 | 0.909401848 | -0.04922 | 0.646859 | -0.51036 |
| heme oxygenase-like protein. | 1.040097 | 1.055799228 | 0.021618 | 0.64805 | 0.504553 |
| hypothetical protein ABUW_2 | 1.071266 | 1.052390387 | -0.02565 | 0.648614 | -0.51235 |
| hypothetical protein ABUW_0 | 0.893518 | 0.902820464 | 0.014943 | 0.648692 | 0.500074 |
| hypothetical protein ABUW_0 | 1.040317 | 1.066990613 | 0.036524 | 0.650511 | 0.505778 |
| 5-carboxymethyl-2-hydroxym | 0.940266 | 0.967104342 | 0.040603 | 0.652793 | 0.498795 |
| multidrug efflux protein AdeA | 1.037125 | 1.01383612 | -0.03277 | 0.653034 | -0.50381 |
| putative MTA/SAH nucleosida | 1.005266 | 0.99343646 | -0.01708 | 0.656361 | -0.48187 |
| phosphoglycolate phosphatas | 0.894121 | 0.938997807 | 0.070652 | 0.657898 | 0.477842 |
| metal-dependent hydrolase o | 1.010011 | 1.018739472 | 0.012414 | 0.658389 | 0.486507 |
| UTP-glucose-1-phosphate urid | 1.039747 | 1.058852885 | 0.026269 | 0.660386 | 0.503915 |
| ABC transporter, methionine- | 0.982684 | 0.992007224 | 0.013623 | 0.660989 | 0.504301 |
| transcriptional regulator, TetR | 0.919536 | 0.943478997 | 0.037084 | 0.661044 | 0.497389 |
| hypothetical protein ABUW_0 | 0.951589 | 0.940377723 | -0.0171 | 0.662161 | -0.50697 |
| ISPpu12 transposase | 0.947011 | 0.991158183 | 0.065734 | 0.662752 | 0.470404 |
| putative transcriptional regula | 0.945724 | 0.975550644 | 0.044798 | 0.662801 | 0.476078 |
| 5'-nucleotidase | 0.990889 | 0.980377088 | -0.01539 | 0.663099 | -0.49764 |
| arginyl-tRNA synthetase | 1.00998 | 1.018815579 | 0.012567 | 0.66313 | 0.46957 |
| hypothetical protein ABUW_1 | 0.995455 | 1.019556617 | 0.034514 | 0.663392 | 0.469334 |
| hypothetical protein ABUW_2 | 0.961132 | 0.930738079 | -0.04636 | 0.66385 | -0.48093 |
| outer membrane protein E | 0.896641 | 0.918849435 | 0.035299 | 0.665712 | 0.482537 |
| acetyltransferase, gnat family | 1.01643 | 1.045706965 | 0.040968 | 0.667022 | 0.464819 |
| hypothetical protein ABUW_1 | 0.882322 | 0.944155574 | 0.097719 | 0.667132 | 0.488394 |
| oxidoreductase, FAD/FMN-bir | 1.007652 | 1.01665024 | 0.012826 | 0.670039 | 0.470914 |
| major facilitator superfamily I | 0.861627 | 0.91687673 | 0.089665 | 0.671575 | 0.481706 |
| transcriptional regulator, IclR | 0.934923 | 0.962973836 | 0.04265 | 0.672281 | 0.462469 |
| YecA family protein | 0.923682 | 0.94473569 | 0.032514 | 0.672344 | 0.468632 |
| hypothetical protein ABUW_2 | 0.989726 | 1.020738842 | 0.044513 | 0.673254 | 0.457094 |
| bifunctional hydroxy-methylp | 0.984991 | 0.991626301 | 0.009687 | 0.673532 | 0.483086 |
| MscS Mechanosensitive ion cl | 0.932054 | 0.956103287 | 0.036753 | 0.674645 | 0.476862 |
| phenylalanyl-tRNA synthetase | 0.962825 | 0.957265128 | -0.00836 | 0.677615 | -0.448 |
| transcriptional regulator, LysR | 0.957721 | 0.986748418 | 0.043078 | 0.677717 | 0.450701 |
| betaine aldehyde dehydroger | 1.008282 | 1.027448367 | 0.027166 | 0.678002 | 0.473575 |
| two-component system respo | 0.981062 | 1.002214551 | 0.030775 | 0.678046 | 0.44776 |
| flavin-containing monooxyger | 1.043404 | 1.037275032 | -0.0085 | 0.678627 | -0.44817 |
| hypothetical protein ABUW_2 | 0.969491 | 0.998509182 | 0.042548 | 0.681009 | 0.460366 |
| 3-deoxy-D-manno-2-octuloso | 1.018125 | 1.011006046 | -0.01012 | 0.682196 | -0.4563 |
| cytosol aminopeptidase | 1.007174 | 0.97737497 | -0.04333 | 0.683335 | -0.46459 |
| esterase...903 | 0.98599 | 0.997248807 | 0.01638 | 0.683834 | 0.439978 |
| hypothetical protein ABUW_2 | 1.079382 | 1.102203625 | 0.030185 | 0.684916 | 0.448461 |
| threonine dehydratase...2054 | 0.923132 | 0.910266339 | -0.02025 | 0.685731 | -0.45947 |
| formimidoylglutamase | 0.991586 | 1.013400895 | 0.031395 | 0.686107 | 0.439559 |
| hypothetical protein ABUW_2 | 0.91363 | 0.884490607 | -0.04676 | 0.686418 | -0.46652 |

| | | | | | |
|---|---|---|---|---|---|
| ABC-type dipeptide/oligopept | 0.982306 | 1.009338103 | 0.039164 | 0.687344 | 0.462204 |
| hypothetical protein ABUW_2 | 0.995042 | 1.003666366 | 0.01245 | 0.687414 | 0.444865 |
| fumarate hydratase, class II | 0.971878 | 0.967173237 | -0.007 | 0.688289 | -0.46196 |
| two-component system histic | 1.082162 | 1.045618405 | -0.04956 | 0.688654 | -0.45969 |
| adenosine deaminase...1044 | 1.054086 | 1.077964721 | 0.032317 | 0.689976 | 0.444758 |
| urease accessory protein G | 0.940108 | 0.904038407 | -0.05644 | 0.69283 | -0.42721 |
| prolipoprotein diacylglyceryl t | 0.983329 | 0.971599445 | -0.01731 | 0.692984 | -0.42513 |
| hypothetical protein ABUW_3 | 0.898492 | 0.926124423 | 0.0437 | 0.694592 | 0.438814 |
| quinoprotein glucose dehydro | 1.06665 | 1.053445908 | -0.01797 | 0.696725 | -0.43768 |
| acetyl-/propionyl-coenzyme A | 1.059368 | 1.040497832 | -0.02593 | 0.697138 | -0.42015 |
| peptidylprolyl isomerase, fkbp | 1.051337 | 1.034023324 | -0.02396 | 0.697583 | -0.44536 |
| DNA-3-methyladenine glycosy | 0.984793 | 0.933388629 | -0.07734 | 0.697692 | -0.44753 |
| transcriptional regulator, LysR | 0.995828 | 0.989558134 | -0.00911 | 0.698147 | -0.41695 |
| hypothetical protein ABUW_1 | 1.053713 | 1.031201028 | -0.03116 | 0.69971 | -0.44193 |
| protein tyrosine phosphatase | 0.974197 | 0.946941799 | -0.04094 | 0.701608 | -0.42336 |
| hypothetical protein ABUW_1 | 1.007284 | 1.017226859 | 0.014171 | 0.701639 | 0.427686 |
| hypothetical protein ABUW_4 | 0.947625 | 0.930504964 | -0.0263 | 0.702497 | -0.41401 |
| transcriptional repressor BetI | 0.918049 | 0.930452923 | 0.019362 | 0.702631 | 0.411267 |
| acetyltransferase...1307 | 0.99138 | 1.014888931 | 0.033812 | 0.703752 | 0.424125 |
| transcriptional regulator, TetR | 0.953697 | 0.98904315 | 0.052503 | 0.706334 | 0.409521 |
| pca operon regulatory protein | 0.91442 | 0.879569269 | -0.05606 | 0.709983 | -0.39953 |
| hypothetical protein ABUW_1 | 1.127511 | 1.201983684 | 0.092276 | 0.710231 | 0.427476 |
| transcriptional regulator, TetR | 1.04677 | 1.061835502 | 0.020616 | 0.710238 | 0.404535 |
| hypothetical protein ABUW_2 | 1.097305 | 1.086786581 | -0.0139 | 0.710254 | -0.41824 |
| septum site-determining prot | 0.920693 | 0.944427425 | 0.03672 | 0.713829 | 0.419232 |
| 3-isopropylmalate dehydratas | 0.987195 | 1.003690971 | 0.023908 | 0.714484 | 0.393114 |
| quinolinate synthetase compl | 0.978801 | 0.968681661 | -0.01499 | 0.714637 | -0.40362 |
| transcriptional regulator...858 | 1.045431 | 1.068549178 | 0.031555 | 0.714834 | 0.396799 |
| transcriptional regulator, IclR | 0.979862 | 0.966598203 | -0.01966 | 0.717412 | -0.41295 |
| ThiF family protein | 1.046942 | 1.03562461 | -0.01568 | 0.718615 | -0.39666 |
| enoyl-CoA hydratase...2343 | 0.916578 | 0.940614977 | 0.037347 | 0.71942 | 0.385609 |
| hypothetical protein ABUW_3 | 1.086656 | 1.065313283 | -0.02862 | 0.721475 | -0.3911 |
| hypothetical protein ABUW_1 | 0.937376 | 0.975082629 | 0.056897 | 0.722131 | 0.401668 |
| bis(5'-nucleosyl)-tetraphospha | 0.97664 | 0.966941363 | -0.0144 | 0.722519 | -0.40606 |
| hypothetical protein ABUW_4 | 1.115026 | 1.158311102 | 0.054945 | 0.722644 | 0.403058 |
| hypothetical protein ABUW_2 | 0.942296 | 0.933622792 | -0.01334 | 0.723021 | -0.38425 |
| putative benzoate transport p | 1.006686 | 0.996867924 | -0.01414 | 0.723317 | -0.3977 |
| chromosome partitioning pro | 0.982806 | 0.978693073 | -0.00605 | 0.723942 | -0.39064 |
| transcriptional regulator, LysR | 0.932483 | 0.923632276 | -0.01376 | 0.725952 | -0.38813 |
| peptidoglycan-binding LysM... | 1.036991 | 1.018951031 | -0.02532 | 0.726352 | -0.38567 |
| nucleoside-diphosphate-sugar | 1.051963 | 1.061462783 | 0.012969 | 0.727673 | 0.397707 |
| lysophospholipase | 1.016313 | 1.028670939 | 0.017437 | 0.729317 | 0.373243 |
| isochorismatase hydrolase...2 | 0.942734 | 0.954751017 | 0.018274 | 0.729532 | 0.388709 |
| 3-isopropylmalate dehydroge | 0.961799 | 0.980651741 | 0.028006 | 0.732312 | 0.38812 |
| glycoprotease | 0.973739 | 0.959436885 | -0.02135 | 0.732339 | -0.37143 |
| alcohol dehydrogenase, class | 0.981891 | 0.988510635 | 0.009694 | 0.733636 | 0.367265 |
| glutamate synthase, large sub | 0.984085 | 0.97528035 | -0.01297 | 0.738835 | -0.35787 |

| | | | | | |
|---|---|---|---|---|---|
| 3-methylglutaconyl-CoA hydra | 0.994037 | 0.975269569 | -0.0275 | 0.739756 | -0.37985 |
| hypothetical protein ABUW_1 | 0.998912 | 1.011475477 | 0.018032 | 0.740172 | 0.363162 |
| hypothetical protein ABUW_0 | 1.141041 | 1.063312451 | -0.10179 | 0.740197 | -0.36105 |
| hypothetical protein ABUW_2 | 0.971788 | 0.993521903 | 0.03191 | 0.741713 | 0.353738 |
| hypothetical protein ABUW_2 | 0.934907 | 0.952211575 | 0.026459 | 0.742013 | 0.376203 |
| LysR-family transcriptional reg | 1.013504 | 1.001599406 | -0.01705 | 0.742108 | -0.36099 |
| replicative DNA helicase...6 | 1.142299 | 1.190957294 | 0.060181 | 0.74246 | 0.353403 |
| RNA binding S1 domain prote | 1.037773 | 1.04736304 | 0.013271 | 0.744734 | 0.354173 |
| oxidoreductase short chain de | 1.080563 | 1.113207976 | 0.04294 | 0.747724 | 0.346317 |
| D-tyrosyl-tRNA(Tyr) deacylase | 0.950902 | 0.914402004 | -0.05647 | 0.747999 | -0.34424 |
| hypothetical protein ABUW_1 | 1.096237 | 1.084791681 | -0.01514 | 0.749713 | -0.35436 |
| hypothetical protein ABUW_1 | 1.08434 | 1.12718873 | 0.055912 | 0.75039 | 0.345151 |
| N-acetyl-gamma-glutamyl-pho | 0.929218 | 0.943418204 | 0.021881 | 0.751616 | 0.348149 |
| ATP synthase F0, C subunit | 0.918348 | 0.899800973 | -0.02944 | 0.753937 | -0.342 |
| methylcrotonoyl-CoA carboxy | 1.002632 | 1.005174367 | 0.003653 | 0.753992 | 0.335922 |
| exopolyphosphatase | 0.955645 | 0.948082963 | -0.01146 | 0.754359 | -0.35216 |
| amidohydrolase...1632 | 1.029269 | 1.036442931 | 0.01002 | 0.755017 | 0.352403 |
| hypothetical protein ABUW_0 | 0.955287 | 0.940923952 | -0.02186 | 0.755439 | -0.33374 |
| Glutamate synthase (NADPH) | 0.974688 | 0.966016971 | -0.01289 | 0.755828 | -0.33443 |
| oxidoreductase...43 | 1.16613 | 1.102352233 | -0.08114 | 0.756133 | -0.34554 |
| glutaminyl-tRNA synthetase | 0.968234 | 0.958557941 | -0.01449 | 0.757031 | -0.35375 |
| tRNA-specific adenosine dean | 0.939401 | 0.965719664 | 0.039863 | 0.757145 | 0.336555 |
| ParA family protein | 0.995574 | 1.003475409 | 0.011404 | 0.758533 | 0.338125 |
| dihydroorotase, homodimeric | 0.985832 | 0.975302631 | -0.01549 | 0.759071 | -0.34057 |
| taurine ABC transporter, perip | 1.122664 | 1.076161298 | -0.06103 | 0.759443 | -0.32841 |
| DNA ligase, NAD-dependent | 0.990525 | 0.980765582 | -0.01428 | 0.760774 | -0.32623 |
| hypothetical protein ABUW_2 | 1.016695 | 0.967193932 | -0.07201 | 0.760799 | -0.32978 |
| extradiol ring-cleavage dioxyg | 0.991048 | 0.997735242 | 0.009703 | 0.760977 | 0.336643 |
| Uridylate kinase | 0.964868 | 0.971209125 | 0.00945 | 0.761621 | 0.325881 |
| transcriptional regulator, LysR | 0.975245 | 0.94184156 | -0.05028 | 0.761716 | -0.33927 |
| polyribonucleotide nucleotidy | 1.01788 | 1.009529033 | -0.01189 | 0.762245 | -0.34319 |
| hypothetical protein ABUW_0 | 1.007946 | 1.035300587 | 0.038632 | 0.763548 | 0.32658 |
| hypothetical protein ABUW_0 | 1.027292 | 1.040746756 | 0.018772 | 0.764374 | 0.32653 |
| hypothetical protein ABUW_1 | 1.017178 | 1.002297257 | -0.02126 | 0.764722 | -0.32077 |
| 3-oxoadipate CoA-transferase | 0.921161 | 0.914807944 | -0.00998 | 0.765156 | -0.32482 |
| deoxyribodipyrimidine photo- | 0.998168 | 1.043720575 | 0.064382 | 0.765397 | 0.329093 |
| two component transcription | 0.955727 | 0.932287424 | -0.03582 | 0.766099 | -0.31843 |
| phosphoribosylformylglycinar | 0.951839 | 0.949068656 | -0.0042 | 0.767712 | -0.33395 |
| methionyl-tRNA synthetase | 1.014647 | 1.020185588 | 0.007854 | 0.768122 | 0.334889 |
| aromatic amino acid transpor | 1.055293 | 1.104918718 | 0.066296 | 0.770074 | 0.32611 |
| activator of HSP90 ATPase...1 | 0.953315 | 0.965613498 | 0.018494 | 0.771505 | 0.323432 |
| benzoate 1,2-dioxygenase, sm | 0.972845 | 1.004936853 | 0.046823 | 0.772688 | 0.310299 |
| biotin biosynthesis protein Bio | 1.132991 | 1.115221645 | -0.02281 | 0.773637 | -0.31057 |
| cation efflux system protein... | 0.946655 | 0.940699106 | -0.00911 | 0.774405 | -0.31321 |
| ribonuclease G | 0.957859 | 0.952007054 | -0.00884 | 0.775372 | -0.30529 |
| TonB-dependent receptor...15 | 0.974411 | 0.987131525 | 0.018712 | 0.775387 | 0.324082 |
| 8-amino-7-oxononanoate syn | 0.974319 | 0.957382478 | -0.0253 | 0.776176 | -0.30446 |

| | | | | | |
|---|---|---|---|---|---|
| hypothetical protein ABUW_2 | 1.017275 | 1.006737447 | -0.01502 | 0.777133 | -0.31136 |
| LmbE-like protein | 1.100447 | 1.079720701 | -0.02743 | 0.777219 | -0.30467 |
| L-lactate permease | 0.918932 | 0.908201182 | -0.01695 | 0.778503 | -0.31377 |
| tRNA pseudouridine synthase | 0.997212 | 0.979656829 | -0.02562 | 0.778626 | -0.31319 |
| aldehyde dehydrogenase...67 | 1.011653 | 0.997438855 | -0.02041 | 0.778908 | -0.31248 |
| beta-hydroxylase | 1.058562 | 1.041472494 | -0.02348 | 0.779499 | -0.3095 |
| aldo/keto reductase...865 | 0.978143 | 0.951750408 | -0.03946 | 0.782061 | -0.31579 |
| TonB-dependent siderophore | 1.028279 | 1.022524567 | -0.0081 | 0.783499 | -0.30923 |
| hypothetical protein ABUW_2 | 0.987769 | 1.001588545 | 0.020044 | 0.784516 | 0.305579 |
| methionine synthase | 0.900605 | 0.906039478 | 0.00868 | 0.784921 | 0.292817 |
| hypothetical protein ABUW_( | 1.024236 | 1.043507859 | 0.026893 | 0.786709 | 0.296752 |
| methionyl-tRNA formyltransfe | 1.005627 | 1.012161756 | 0.009345 | 0.787773 | 0.295241 |
| OmpW family protein...1531 | 1.006297 | 0.991155121 | -0.02187 | 0.789386 | -0.29655 |
| phage integrase | 0.987305 | 0.962586681 | -0.03658 | 0.792133 | -0.28235 |
| methionine adenosyltransfera | 1.009873 | 0.989508143 | -0.02939 | 0.793309 | -0.29136 |
| lysine exporter protein | 0.587175 | 0.598515951 | 0.027599 | 0.79497 | 0.278567 |
| para-aminobenzoate/anthran | 0.989853 | 1.000462918 | 0.015382 | 0.795094 | 0.281414 |
| hypothetical protein ABUW_1 | 1.037406 | 1.025341194 | -0.01688 | 0.795597 | -0.27889 |
| D-serine ammonia-lyase | 0.968198 | 0.973737656 | 0.008232 | 0.797623 | 0.282538 |
| lipase foldase | 1.077136 | 1.106613982 | 0.038952 | 0.798251 | 0.273549 |
| copper-translocating P-type A | 1.042367 | 1.050374529 | 0.011041 | 0.798745 | 0.289052 |
| alpha/beta fold family hydrola | 1.144449 | 1.125817407 | -0.02368 | 0.799196 | -0.27199 |
| DNA polymerase I | 0.961296 | 0.965224964 | 0.005884 | 0.800564 | 0.278583 |
| hypothetical protein ABUW_3 | 1.01296 | 1.023126716 | 0.014407 | 0.801391 | 0.269136 |
| endonuclease/exonuclease/p | 0.95389 | 0.972545791 | 0.027944 | 0.801856 | 0.275076 |
| hypothetical protein ABUW_2 | 0.977345 | 1.006338496 | 0.042175 | 0.801896 | 0.268335 |
| acetyltransferase, gnat family | 0.942752 | 0.951279931 | 0.012992 | 0.801979 | 0.275334 |
| hypothetical protein ABUW_( | 1.077544 | 1.09738165 | 0.026319 | 0.802292 | 0.278266 |
| lytic transglycosylase, catalyti | 1.009495 | 1.002779861 | -0.00963 | 0.802742 | -0.26717 |
| urea amidolyase | 1.081274 | 1.088184331 | 0.009191 | 0.803221 | 0.273207 |
| acetyltransferase, GNAT famil | 1.12276 | 1.143136581 | 0.025948 | 0.803658 | 0.265661 |
| methionine biosynthesis prot | 0.960246 | 0.965433567 | 0.007773 | 0.804154 | 0.268831 |
| hypothetical protein ABUW_3 | 1.000773 | 0.996540034 | -0.00611 | 0.804993 | -0.26409 |
| transcriptional regulator, Gntl | 1.011234 | 1.027348173 | 0.022808 | 0.806115 | 0.262277 |
| phosphoglycerate kinase | 0.955803 | 0.959846718 | 0.006091 | 0.808862 | 0.262309 |
| arginine N-succinyltransferase | 0.965345 | 0.925139252 | -0.06137 | 0.809031 | -0.26175 |
| signal transduction histidine k | 0.983864 | 0.974552931 | -0.01372 | 0.809042 | -0.27107 |
| hypothetical protein ABUW_( | 0.946821 | 0.919319257 | -0.04253 | 0.809183 | -0.26493 |
| dimethyladenosine transferas | 1.067655 | 1.076819438 | 0.012331 | 0.810171 | 0.259672 |
| hypothetical protein ABUW_1 | 1.041545 | 1.029903587 | -0.01622 | 0.810531 | -0.26254 |
| outer membrane protein...24 | 1.013468 | 0.954684769 | -0.0862 | 0.812179 | -0.26508 |
| acyl-CoA dehydrogenase...543 | 0.994532 | 1.010134112 | 0.022457 | 0.81311 | 0.262485 |
| 1-deoxy-D-xylulose 5-phosph | 1.036125 | 1.029662208 | -0.00903 | 0.814515 | -0.25054 |
| transporter, drug/metabolite | 0.994394 | 1.012923118 | 0.026635 | 0.816744 | 0.24914 |
| sugar kinase, ribokinase famil | 0.948166 | 0.971718411 | 0.035399 | 0.818627 | 0.24648 |
| 3-oxoadipate CoA-transferase | 1.046474 | 1.023537579 | -0.03197 | 0.821713 | -0.24689 |
| uracil phosphoribosyltransfer | 0.916312 | 0.919786047 | 0.00546 | 0.821845 | 0.242057 |

| | | | | | |
|---|---|---|---|---|---|
| putative ribonuclease | 1.023214 | 0.998293898 | -0.03557 | 0.822878 | -0.25348 |
| 2-ketogluconate reductase | 0.94749 | 0.95459325 | 0.010775 | 0.823875 | 0.242099 |
| CsuC | 0.929795 | 0.934469539 | 0.007234 | 0.825388 | 0.23557 |
| 2-C-methyl-D-erythritol 4-pho | 0.99686 | 1.008123592 | 0.01621 | 0.826271 | 0.244561 |
| glutamate/aspartate transpor | 1.092335 | 1.077821516 | -0.0193 | 0.826286 | -0.24558 |
| hypothetical protein ABUW_2 | 0.920801 | 0.928849253 | 0.012556 | 0.827598 | 0.23463 |
| peptide chain release factor 2 | 0.907948 | 0.922622757 | 0.023131 | 0.827916 | 0.238587 |
| general secretion pathway pr | 0.943749 | 0.929675074 | -0.02168 | 0.829838 | -0.2429 |
| aldehyde dehydrogenase...22 | 0.939793 | 0.946396013 | 0.010101 | 0.830939 | 0.241654 |
| hypothetical protein ABUW_3 | 1.138906 | 1.110825512 | -0.03602 | 0.831418 | -0.23266 |
| Glycerate kinase | 0.93691 | 0.950437292 | 0.02068 | 0.833142 | 0.238397 |
| acyl carrier protein...2241 | 0.944701 | 0.956733911 | 0.01826 | 0.833312 | 0.234913 |
| transcriptional regulator, Ara( | 0.976857 | 0.986856985 | 0.014694 | 0.83341 | 0.231585 |
| biofilm-associated protein...2 | 1.045381 | 1.060790656 | 0.021111 | 0.833724 | 0.235029 |
| pyridine nucleotide-disulfide | 1.06428 | 1.085005107 | 0.027824 | 0.834563 | 0.225045 |
| adenylate/guanylate cyclase | 1.036514 | 1.046546854 | 0.013897 | 0.837525 | 0.230913 |
| diaminobutyrate--2-oxoglutar | 0.995659 | 0.987355064 | -0.01208 | 0.838001 | -0.23074 |
| oxidoreductase...1921 | 1.010721 | 1.025044521 | 0.020302 | 0.839934 | 0.229198 |
| hypothetical protein ABUW_0 | 0.972451 | 0.981088757 | 0.012758 | 0.84162 | 0.224306 |
| OmpA/MotB...2467 | 1.085329 | 1.094414162 | 0.012026 | 0.842123 | 0.225026 |
| putative adenylyltransferase | 0.983784 | 0.971552694 | -0.01805 | 0.843208 | -0.22402 |
| 2,3,4,5-tetrahydropyridine-2,( | 0.928206 | 0.92475406 | -0.00538 | 0.844318 | -0.20967 |
| ATP-dependent Clp protease, | 0.988888 | 0.995407362 | 0.009479 | 0.844356 | 0.218256 |
| porin B...2388 | 0.995384 | 0.991646395 | -0.00543 | 0.845108 | -0.21365 |
| leucyl-tRNA synthetase | 0.969764 | 0.967451416 | -0.00344 | 0.846679 | -0.2101 |
| hypothetical protein ABUW_3 | 1.021738 | 0.995002702 | -0.03825 | 0.847504 | -0.21599 |
| Hca operon transcriptional ac | 0.98128 | 0.98544833 | 0.006116 | 0.847519 | 0.208832 |
| hypothetical protein ABUW_3 | 1.026988 | 1.03436639 | 0.010329 | 0.847595 | 0.207363 |
| heme oxygenase-like protein. | 1.082753 | 1.066110191 | -0.02235 | 0.848107 | -0.21642 |
| rhomboid family protein | 0.975615 | 0.999096131 | 0.034312 | 0.850136 | 0.205305 |
| TonB-dependent siderophore | 0.968629 | 0.96365641 | -0.00743 | 0.850458 | -0.20659 |
| HlyD family secretion protein | 1.167295 | 1.201416072 | 0.041566 | 0.852726 | 0.20273 |
| 5'-methylthioadenosine/S-ade | 1.017912 | 1.032194714 | 0.020103 | 0.853315 | 0.206623 |
| transcriptional regulator, Ara( | 0.914832 | 0.889113828 | -0.04114 | 0.854007 | -0.19881 |
| hypothetical protein ABUW_1 | 0.924693 | 0.94238019 | 0.027335 | 0.854704 | 0.20759 |
| FMN-binding protein | 1.002745 | 0.993040034 | -0.01403 | 0.857689 | -0.19253 |
| lysozyme-like domain-contain | 0.904801 | 0.895084639 | -0.01558 | 0.857957 | -0.19336 |
| hypothetical protein ABUW_2 | 1.021686 | 1.031924135 | 0.014385 | 0.858709 | 0.198063 |
| aldehyde dehydrogenase...13 | 0.960541 | 0.967272487 | 0.010076 | 0.859152 | 0.199729 |
| transglycosylase-associated p | 1.012691 | 1.042848142 | 0.042336 | 0.861634 | 0.185819 |
| exodeoxyribonuclease III | 1.01475 | 1.011909811 | -0.00404 | 0.862312 | -0.18834 |
| 5'/3'-nucleotidase SurE | 0.906482 | 0.902267558 | -0.00672 | 0.863588 | -0.18339 |
| hypothetical protein ABUW_0 | 0.998013 | 1.006075651 | 0.011608 | 0.864517 | 0.183373 |
| two-component system hybri | 0.974499 | 0.990971124 | 0.024182 | 0.866094 | 0.181462 |
| aromatic-ring-hydroxylating d | 0.985869 | 0.979938453 | -0.00871 | 0.866258 | -0.17978 |
| hypothetical protein ABUW_1 | 1.029078 | 1.024152016 | -0.00692 | 0.867093 | -0.18059 |
| tetratricopeptide repeat dom | 1.042208 | 1.051874743 | 0.01332 | 0.868497 | 0.181691 |

| | | | | | |
|---|---|---|---|---|---|
| hypothetical protein ABUW_2 | 0.928129 | 0.939380701 | 0.017384 | 0.87107 | 0.173239 |
| putative acetyltransferase | 1.087737 | 1.094344517 | 0.008737 | 0.871695 | 0.173111 |
| transcriptional regulator, TetF | 1.012025 | 0.98304457 | -0.04192 | 0.874042 | -0.17863 |
| shikimate 5-dehydrogenase | 0.973275 | 0.967804979 | -0.00813 | 0.874235 | -0.17766 |
| hydrolase...334 | 1.013032 | 1.007884981 | -0.00735 | 0.874426 | -0.1686 |
| polysaccharide deacetylase...( | 0.968507 | 0.954456787 | -0.02108 | 0.874997 | -0.17539 |
| Fe-S protein assembly chaper | 1.004061 | 0.999926577 | -0.00595 | 0.87679 | -0.17132 |
| hypothetical protein ABUW_1 | 0.927751 | 0.939345463 | 0.017918 | 0.877225 | 0.17258 |
| TonB-dependent siderophore | 0.985555 | 0.990107325 | 0.006649 | 0.87862 | 0.169544 |
| DNA replication protein (plasr | 0.973359 | 0.97098933 | -0.00352 | 0.879638 | -0.16988 |
| hypothetical protein ABUW_1 | 1.049375 | 1.045198637 | -0.00575 | 0.880136 | -0.1609 |
| beta-lactamase...991 | 1.042172 | 1.02084734 | -0.02983 | 0.880887 | -0.16009 |
| hypothetical protein ABUW_2 | 1.001947 | 0.995847193 | -0.00881 | 0.881331 | -0.16062 |
| methylated-DNA--protein-cys | 0.985105 | 0.978538678 | -0.00965 | 0.881944 | -0.16471 |
| methyltransferase superfamil | 0.822977 | 0.828932104 | 0.010401 | 0.882803 | 0.157433 |
| disulfide bond formation prot | 0.94624 | 0.957952413 | 0.017748 | 0.883253 | 0.158007 |
| resolvase (plasmid) | 1.115754 | 1.107630016 | -0.01054 | 0.883383 | -0.15639 |
| 3-carboxy-cis,cis-muconate cy | 1.047375 | 1.040519935 | -0.00947 | 0.883955 | -0.16358 |
| nicotinate phosphoribosyltrar | 0.957951 | 0.955021993 | -0.00442 | 0.884139 | -0.15555 |
| transcriptional regulator, TetF | 1.029084 | 1.025705365 | -0.00474 | 0.886068 | -0.15459 |
| oxygenase | 1.006132 | 0.99123825 | -0.02152 | 0.886765 | -0.16016 |
| DNA polymerase III delta prim | 0.956152 | 0.952196029 | -0.00598 | 0.887446 | -0.16014 |
| guanosine-3,5-bis(diphosphat | 1.033476 | 1.029867437 | -0.00505 | 0.887654 | -0.15298 |
| phosphoribosylglycinamide fc | 1.044287 | 1.038151613 | -0.0085 | 0.887765 | -0.1556 |
| phenylacetate-CoA oxygenase | 0.942419 | 0.962780221 | 0.030838 | 0.888522 | 0.149637 |
| organic solvent tolerance pro | 1.021822 | 1.019850225 | -0.00279 | 0.8886 | -0.15673 |
| tannase/feruloyl esterase fam | 1.106188 | 1.089822612 | -0.0215 | 0.889068 | -0.15637 |
| phosphate transport system r | 1.021815 | 1.019516424 | -0.00325 | 0.890981 | -0.14733 |
| 3-demethylubiquinone-9 3-O- | 1.007724 | 1.017632298 | 0.014116 | 0.894375 | 0.150097 |
| UDP-N-acetylmuramate:L-alar | 0.963983 | 0.969584559 | 0.008358 | 0.894522 | 0.144988 |
| two-component system regul | 1.063524 | 1.070013953 | 0.008777 | 0.894968 | 0.140629 |
| hypothetical protein ABUW_( | 1.007784 | 1.015119013 | 0.010463 | 0.895959 | 0.140942 |
| 3-hydroxyphenylpropionic aci | 0.903558 | 0.896457164 | -0.01138 | 0.898784 | -0.13603 |
| hypothetical protein ABUW_1 | 0.928664 | 0.936199629 | 0.011659 | 0.898855 | 0.140873 |
| monovalent cation transporte | 0.958057 | 0.96122285 | 0.00476 | 0.899137 | 0.134995 |
| thiamine-phosphate pyropho: | 0.968602 | 0.961706502 | -0.01031 | 0.90083 | -0.14067 |
| hypothetical protein ABUW_( | 1.019138 | 1.031458378 | 0.017336 | 0.9029 | 0.131424 |
| NADH dehydrogenase I chain | 0.98855 | 0.995235057 | 0.009724 | 0.902952 | 0.136409 |
| hypothetical protein ABUW_3 | 1.008557 | 0.998172413 | -0.01493 | 0.9031 | -0.1301 |
| sigma-54 specific, transcriptio | 1.012051 | 1.01670661 | 0.006622 | 0.904271 | 0.135179 |
| acyl-CoA dehydrogenase...97 | 1.065813 | 1.054670898 | -0.01516 | 0.905032 | -0.13134 |
| hypothetical protein ABUW_1 | 0.847314 | 0.837471367 | -0.01686 | 0.906825 | -0.12532 |
| DNA-directed RNA polymeras | 1.007489 | 1.00289872 | -0.00659 | 0.907385 | -0.12388 |
| glyceraldehyde 3-phosphate ( | 0.947003 | 0.949675757 | 0.004066 | 0.909721 | 0.121165 |
| hypothetical protein ABUW_( | 1.059309 | 1.061088059 | 0.002421 | 0.909902 | 0.127179 |
| dihydrolipoamide dehydroger | 1.020929 | 1.030826093 | 0.013918 | 0.910304 | 0.127111 |
| hypothetical protein ABUW_( | 1.025113 | 1.029252667 | 0.005814 | 0.911161 | 0.124499 |

| | | | | | |
|---|---|---|---|---|---|
| allophanate hydrolase | 0.977223 | 0.97442356 | -0.00414 | 0.911228 | -0.12042 |
| septum formation protein Ma | 0.983433 | 0.989978069 | 0.009569 | 0.912081 | 0.117606 |
| hypothetical protein ABUW_3 | 1.008133 | 1.004757555 | -0.00484 | 0.913453 | -0.11573 |
| hypothetical protein ABUW_1 | 1.039871 | 1.043520714 | 0.005055 | 0.914511 | 0.118812 |
| hypothetical protein ABUW_1 | 0.952886 | 0.956374303 | 0.005272 | 0.915313 | 0.119097 |
| electron transport complex, r | 0.982793 | 0.994842478 | 0.017581 | 0.916742 | 0.112592 |
| NADP-dependent fatty aldehy | 0.961662 | 0.959356649 | -0.00346 | 0.9182 | -0.11523 |
| hypothetical protein ABUW_C | 1.025557 | 1.022936973 | -0.00369 | 0.919552 | -0.1138 |
| hypothetical protein ABUW_3 | 0.974936 | 0.978454555 | 0.005198 | 0.920819 | 0.106056 |
| putative methyltransferase | 0.970324 | 0.973060356 | 0.004063 | 0.920917 | 0.107649 |
| putative competence protein | 1.008197 | 1.011070061 | 0.004105 | 0.922447 | 0.107113 |
| dihydropteroate synthase | 0.930709 | 0.926382324 | -0.00672 | 0.922795 | -0.10319 |
| malonate utilization transcrip | 1.049097 | 1.040202605 | -0.01228 | 0.923717 | -0.10723 |
| major facilitator superfamily I | 0.971658 | 0.961675707 | -0.0149 | 0.92459 | -0.10133 |
| transcriptional regulator, XRE | 1.126609 | 1.136581615 | 0.012715 | 0.926578 | 0.103511 |
| vitamin B12 receptor | 0.967024 | 0.963473389 | -0.00531 | 0.927167 | -0.10233 |
| fimbrial assembly protein PilC | 0.957786 | 0.954102601 | -0.00556 | 0.927614 | -0.09963 |
| ATP-dependent dsDNA exonu | 0.952467 | 0.955449966 | 0.004512 | 0.930955 | 0.097762 |
| hypothetical protein ABUW_1 | 0.939225 | 0.93730147 | -0.00296 | 0.934897 | -0.08948 |
| acetyl-CoA acetyltransferase.. | 0.986896 | 0.98520762 | -0.00247 | 0.935069 | -0.09074 |
| hypothetical protein ABUW_1 | 0.9573 | 0.956695024 | -0.00091 | 0.936198 | -0.08829 |
| hypothetical protein ABUW_1 | 0.966664 | 0.96418146 | -0.00371 | 0.936316 | -0.08667 |
| valyl-tRNA synthetase | 0.992425 | 0.990286254 | -0.00311 | 0.936327 | -0.09 |
| acetylglutamate kinase | 0.963726 | 0.962292147 | -0.00215 | 0.936803 | -0.08646 |
| magnesium Mg(2+)/cobalt Co | 1.048189 | 1.061006233 | 0.017534 | 0.937014 | 0.085067 |
| ferredoxin-NADP reductase | 0.990876 | 0.993842679 | 0.004312 | 0.940312 | 0.080878 |
| exodeoxyribonuclease X, puta | 0.923427 | 0.925710877 | 0.003564 | 0.940538 | 0.080371 |
| siderophore biosynthesis prot | 1.277991 | 1.267000753 | -0.01246 | 0.940855 | -0.07922 |
| DNA mismatch repair protein | 0.958479 | 0.956274315 | -0.00332 | 0.941803 | -0.07903 |
| oxidoreductase...140 | 1.19401 | 1.224210421 | 0.036037 | 0.942242 | 0.07794 |
| electron transfer flavoprotein | 0.996303 | 0.995167835 | -0.00164 | 0.942718 | -0.08109 |
| putative hemolysin | 1.11415 | 1.109448353 | -0.0061 | 0.943453 | -0.07871 |
| hypothetical protein ABUW_2 | 0.959788 | 0.95608399 | -0.00558 | 0.943756 | -0.07637 |
| type IV pilus signal transducti | 1.046911 | 1.050932965 | 0.005532 | 0.944183 | 0.074888 |
| putative cold shock protein | 0.974421 | 0.979008177 | 0.006776 | 0.944396 | 0.078211 |
| ankyrin repeat protein | 0.880352 | 0.875309024 | -0.00829 | 0.944748 | -0.07739 |
| hypothetical protein ABUW_2 | 0.955761 | 0.957990788 | 0.003361 | 0.946129 | 0.072067 |
| ABC transporter, ATP-binding | 0.973148 | 0.97120848 | -0.00288 | 0.946767 | -0.07537 |
| glycerol-3-phosphate dehydrc | 0.97471 | 0.973853857 | -0.00127 | 0.947127 | -0.07096 |
| multidrug efflux protein...158 | 1.02708 | 1.025502174 | -0.00222 | 0.947243 | -0.07464 |
| glutathione S-transferase...13 | 1.000761 | 0.998372691 | -0.00345 | 0.947808 | -0.07321 |
| hypothetical protein ABUW_C | 0.990181 | 0.986715568 | -0.00506 | 0.94887 | -0.07039 |
| hypothetical protein ABUW_1 | 0.955243 | 0.95189439 | -0.00507 | 0.948945 | -0.06824 |
| hypothetical protein ABUW_2 | 1.0364 | 1.044687123 | 0.01149 | 0.949792 | 0.067045 |
| TonB-dependent receptor...62 | 1.064134 | 1.058620008 | -0.00749 | 0.951182 | -0.0652 |
| hypothetical protein ABUW_C | 1.014067 | 1.017279043 | 0.004562 | 0.951816 | 0.066675 |
| recombination protein RecR | 0.960395 | 0.958773958 | -0.00244 | 0.952358 | -0.06517 |

| | | | | | |
|---|---|---|---|---|---|
| bifunctional adenosylcobalam | 0.979499 | 0.983808216 | 0.006333 | 0.954156 | 0.061174 |
| replication protein (plasmid) | 0.983725 | 0.980880871 | -0.00418 | 0.954816 | -0.0618 |
| microcin B17 transport protei | 0.926964 | 0.932032509 | 0.007867 | 0.955984 | 0.060936 |
| transcriptional regulator, XRE | 1.085151 | 1.079164296 | -0.00798 | 0.956154 | -0.05881 |
| L-lactate utilization transcript | 0.979509 | 0.97421824 | -0.00781 | 0.9658 | -0.0458 |
| transcriptional regulator, TetF | 1.033362 | 1.036531124 | 0.004418 | 0.966264 | 0.045113 |
| hypothetical protein ABUW_1 | 0.985954 | 0.989484847 | 0.005158 | 0.966438 | 0.046723 |
| hypothetical protein ABUW_3 | 0.97183 | 0.969902556 | -0.00286 | 0.967939 | -0.04349 |
| amidophosphoribosyltransfer | 0.945979 | 0.945503071 | -0.00073 | 0.968385 | -0.04267 |
| MaoC domain protein dehydr | 1.032229 | 1.03070009 | -0.00214 | 0.968468 | -0.04311 |
| carbon storage regulator | 0.980481 | 0.979928858 | -0.00081 | 0.969523 | -0.04081 |
| bacterioferritin...2226 | 0.975778 | 0.978520065 | 0.004048 | 0.970836 | 0.041129 |
| transcriptional regulator, TetF | 1.028843 | 1.026989578 | -0.0026 | 0.971502 | -0.03801 |
| replicative DNA helicase...111 | 0.940426 | 0.938876603 | -0.00238 | 0.972198 | -0.03727 |
| LysR family transcriptional reg | 1.006287 | 1.004828006 | -0.00209 | 0.975803 | -0.03241 |
| dihydrodipicolinate synthetas | 0.983747 | 0.98257703 | -0.00172 | 0.976577 | -0.03142 |
| ribosomal protein L11 methyl | 1.049335 | 1.050176637 | 0.001157 | 0.976699 | 0.031965 |
| hypothetical protein ABUW_1 | 0.95025 | 0.951243019 | 0.001506 | 0.977605 | 0.03065 |
| hypothetical protein ABUW_3 | 0.96245 | 0.961409221 | -0.00156 | 0.978257 | -0.03054 |
| phosphate ABC transporter, p | 1.08603 | 1.083730016 | -0.00306 | 0.97873 | -0.02895 |
| hypothetical protein ABUW_1 | 1.006116 | 1.007266982 | 0.00165 | 0.979873 | 0.027877 |
| branched-chain amino acid ar | 0.956367 | 0.955954953 | -0.00062 | 0.98055 | -0.02727 |
| outer membrane protein asse | 0.993555 | 0.993839395 | 0.000413 | 0.982111 | 0.024347 |
| hypothetical protein ABUW_( | 0.970468 | 0.971753121 | 0.001909 | 0.98303 | 0.023425 |
| undecaprenyl-diphosphatase | 0.991968 | 0.993385815 | 0.002061 | 0.983756 | 0.022422 |
| hypothetical protein ABUW_2 | 0.969525 | 0.964709639 | -0.00718 | 0.98427 | -0.02142 |
| cytochrome O ubiquinol oxida | 0.973487 | 0.972464665 | -0.00152 | 0.985072 | -0.02109 |
| putative RND family drug tran | 1.036204 | 1.036982342 | 0.001084 | 0.985401 | 0.020595 |
| acyl-CoA dehydrogenase, mid | 0.873693 | 0.872852228 | -0.00139 | 0.985712 | -0.01968 |
| phosphoserine phosphatase.. | 1.071472 | 1.073714454 | 0.003017 | 0.98638 | 0.01877 |
| succinate dehydrogenase iror | 0.88368 | 0.883953354 | 0.000446 | 0.987147 | 0.017225 |
| phosphoglycerate mutase...73 | 0.98234 | 0.982846228 | 0.000744 | 0.987311 | 0.016936 |
| type IV-A pilus assembly ATPa | 1.051581 | 1.049179074 | -0.0033 | 0.987623 | -0.01664 |
| transcriptional regulator, Gntl | 0.95566 | 0.956169471 | 0.000769 | 0.988414 | 0.015724 |
| hypothetical protein ABUW_( | 1.102312 | 1.103072564 | 0.000996 | 0.988961 | 0.014771 |
| NADP-dependent fatty aldehy | 0.997662 | 0.997166848 | -0.00072 | 0.989099 | -0.0149 |
| glutathione-regulated potassi | 1.020351 | 1.019500871 | -0.0012 | 0.989104 | -0.01453 |
| hypothetical protein ABUW_( | 1.041978 | 1.041338554 | -0.00089 | 0.990617 | -0.01253 |
| alcohol dehydrogenase, zinc-l | 0.983416 | 0.983005525 | -0.0006 | 0.991092 | -0.01226 |
| hypothetical protein ABUW_1 | 0.988066 | 0.987426067 | -0.00093 | 0.991865 | -0.01139 |
| hypothetical protein ABUW_( | 1.032701 | 1.032163301 | -0.00075 | 0.992539 | -0.00999 |
| transcriptional regulator, LysF | 1.013793 | 1.0140841 | 0.000414 | 0.993782 | 0.00871 |
| hypothetical protein ABUW_( | 1.00648 | 1.005985247 | -0.00071 | 0.994997 | -0.007 |
| D-alanyl-D-alanine carboxype | 0.990921 | 0.990436332 | -0.00071 | 0.995834 | -0.00569 |
| hypothetical protein ABUW_1 | 0.88178 | 0.881481682 | -0.00049 | 0.996278 | -0.00524 |
| 4'-phosphopantetheinyl trans | 1.070627 | 1.069145346 | -0.002 | 0.997545 | -0.00347 |
| glutaredoxin-related protein | 0.953933 | 0.953715674 | -0.00033 | 0.997638 | -0.00316 |

| | | | | | |
|---|---|---|---|---|---|
| carbon-nitrogen hydrolase | 0.973138 | 0.97319152 | 7.97E-05 | 0.998456 | 0.00212 |
| hypothetical protein ABUW_( | 1.00378 | 1.003828327 | 6.98E-05 | 0.99906 | 0.001298 |
